# A Thermodynamically Favoured Molecular Computer: Robust, Fast, Renewable, Scalable

**DOI:** 10.1101/2025.07.16.664196

**Authors:** Tristan Stérin, Abeer Eshra, Janet Adio, Constantine Glen Evans, Damien Woods

**Affiliations:** Hamilton Institute and Department of Computer Science; Department of Biology, Maynooth University, Co. Kildare, Ireland; prgm.dev, 9 rue des colonnes, Paris, France; Evans Foundation for Molecular Medicine, Pasadena, CA, USA

## Abstract

Like life, computers are out-of-equilibrium.^1,2^ Thermodynamically favoured error states are thwarted by energetically-costly processes such as kinetic proofreading of biological polymers, error-correcting codes in computer data storage, and redundancy in molecular programming. Decades of theoretical work shows that unlike life thermodynamic computers can operate by drifting naturally to equilibrium.^3,4^ Similar ideas underlie machine learning models^5^ and search algorithms such as simulated annealing,^6^ although executed on non-equilibrium architectures at enormous energy cost.^7^ Physically implementing thermodynamically favoured computation is a decades-long challenge that could reduce dependence on fuel-consuming error-correction and precise kinetic control. Here, we demonstrate a thermodynamically favoured Scaffolded DNA Computer (SDC) on ten programs including MULTIPLICATION-by-3, Division-by-2, 8-bit Parity-detection, and Addition of up to 25-bit numbers. SDC algorithms have simple experimental protocols, can be reused dozens of times and small instances run in under a minute. The SDC is grounded in mathematical, physical and computer science principles that explain why the output is thermodynamically favoured, why no error-correction nor precise step-by-step kinetic control are required and how it is programmable and scalable. This work creates a new way to think about equilibrium computation in all manner of synthetic systems.

---

Like living systems that consume fuel to stave off thermodynamic heat death, computers are typically out-of-equilibrium, consuming energy to enforce data integrity. The massive gap between theoretical energy lower bounds,^1,2,4,8^ and what modern computing systems consume,^9^ leaves significant space to rethink computing infrastructure and algorithms.^4,10,11^ To reduce this gap a perhaps counter-intuitive avenue is to design a programmable physical system whose equilibrium encodes the output of an algorithm, then allow the computer to simply drift to the right answer (Fig. 1, Supplementary Note S1). This notion has been presented in various forms^1,3,4,11^ and stands in stark contrast to popular computing methods, so why is it not widely adopted?

**Fig. 1.**
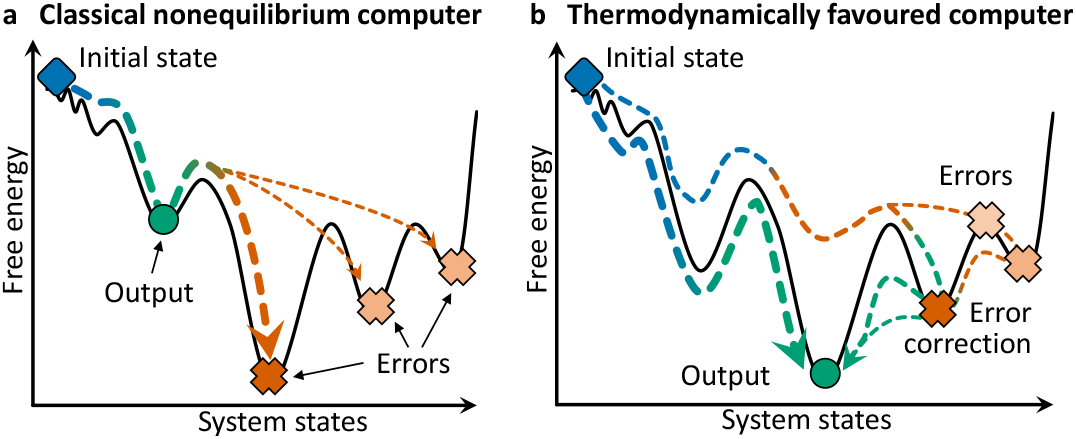
Classical versus thermodynamically favoured computation. **a.** Classical molecular computers have an intended target/final state that is out-of-equilibrium. Errors show up as unintended nucleation of incorrect structures in algorithmic self-assembly^13^ or leak in strand displacement circuits.^16^ Digital electronic computers have similar issues, where decades of research have lowered error rates but with large energetic costs.^9^ **b.** Thermodynamically favoured computation has the output state being the energetically most favoured. Also, the input state should be easy to prepare. Hence, computation happens by the system naturally and automatically going to equilibrium (Supplementary Notes S1, S2).

The first challenge is finding a physical implementation that is computationally expressive yet suited to energy-landscape engineering and analysis. The second challenge is thermodynamic: the input state should be easy to prepare and the output state should be highly favoured, meanin the design avoids equilibria polluted by multiple off-target states. The final challenge is speed: the system needs reasonably good kinetics, evolving along a smooth energy landscape.

Using DNA^12^ researchers have built self-assembling 6-bit programs,^13^ pattern recognisers,^14^ information-based self-replicators,^15^ 4-bit square root calculators,^16^ timers in cell culture,^17^ analog dynamical systems,^18,19^ Boolean circuits and surface-running walkers,^20–23^ and DNA data storage.^24^ Despite being among the most impressive molecular computers to date, there remain numerous difficulties due to the prevailing use of out-of-equilibrium architectures: logical errors and unintended nucleation for algorithmic self-assembly, signal leak to error states for strand displacement systems, tedious manual preparation and sensitivity to experimental conditions, and most are discarded as waste after one run (Supplementary Note S2).

However, there has been progress. One theory-driven approach shows how to encode computations in intentionally simplified chemical equilibria.^3,25^ Also, small equilibrium Boolean circuits were demonstrated using weak molecular affinities.^26^ Thermodynamic penalties to logical errors in DNA circuits were proposed.^27^ DNA Origami^28^ provides inspiration by exploiting a beautiful scaffolded principle to create complex bespoke nanostructures^29^ with a huge thermodynamic drive, but lacking any notion of computation (Supplementary Note S2). To address the first challenge, we propose and experimentally demonstrate a thermodynamically favoured, programmable, Scaffolded DNA Computer (SDC). The SDC has thermodynamic costs but unlike typical forms of computation, avoids needing extra error-correction subsystems by naturally evolving to the desired output state at equilibrium (Fig. 1b). Our approach to the second challenge is principled: mathematically, in simulation, and by molecular design, we show that the SDC output state is highly-favoured, outcompeting exponentially many other configurations in theory (but also verified experimentally). Finally, the SDC design has highly-parallel kinetics, with multiple potential pathways to the output, a novel thermodynamic form of molecular computing that, provides a well-structured and traversable energy landscape, but without tightly controling kinetics. By rethinking computation to be thermodynamically favoured, we get a conceptually simple system that is programmable, robust to errors, fast, and reusable.

## Thermodynamic computing design principles

An *N*-position *ℓ*-bit SDC has a one-dimensional scaffold with *N* unique binding domains called *scaffold positions* (Fig. 2a). A set of *computing tiles* is assigned to each scaffold position. No special out-of-equilibrium input state preparation is required. During a computation, computing tiles *bind* by their bottom *position domain* to a matching scaffold position. Information is processed by the binding of adjacent color-matching tile sides, or *compute domains*. Mismatching adjacent compute domains are permitted, but are an *algorithmic error*, implying an energetic cost that will be resolved by the future *replacement* of one or both tiles (Fig. 2b).

**Fig. 2.**
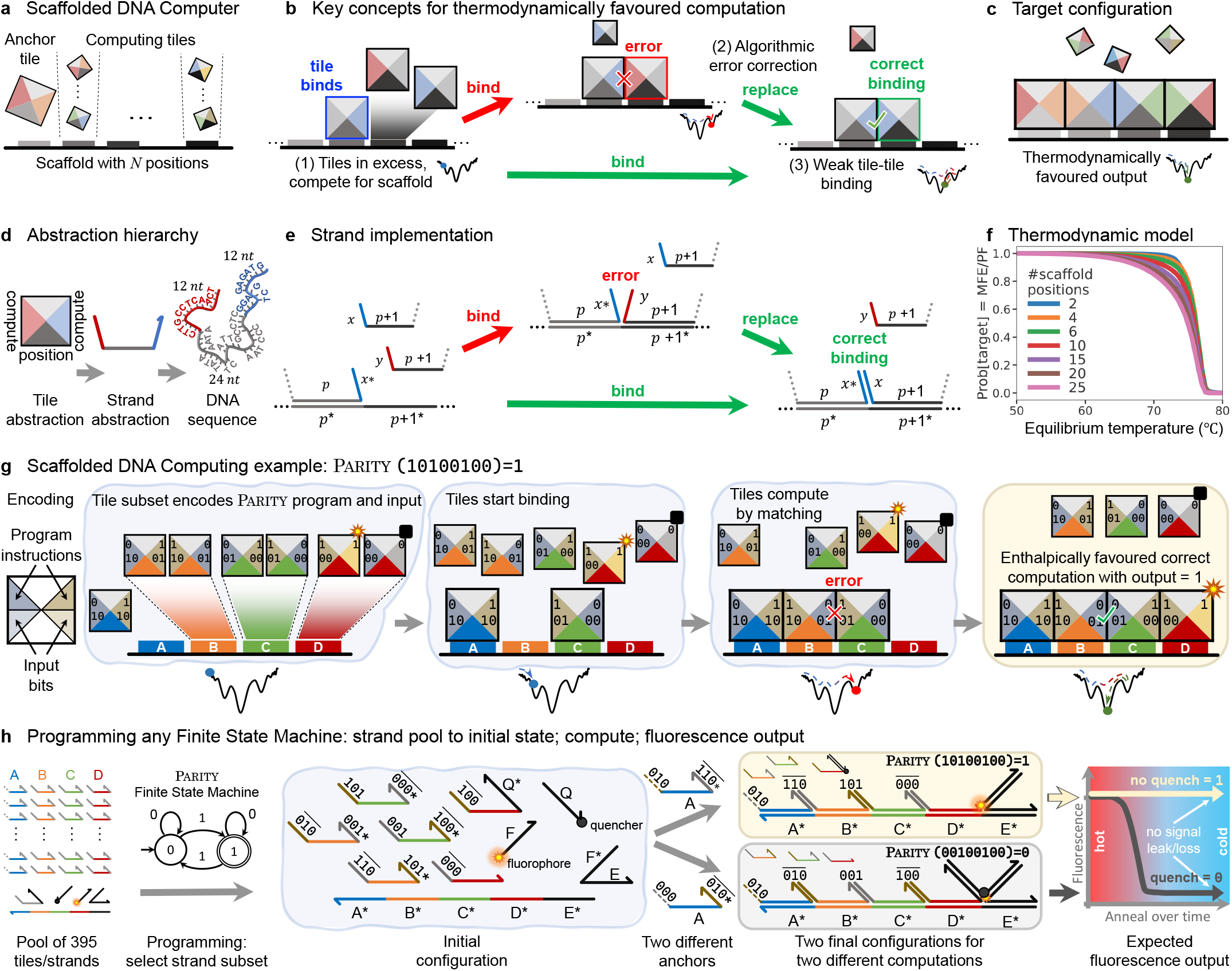
Scaffolded DNA computer (SDC) design principles. **a.** Reprogrammable SDC architecture: scaffold with *N* positions, each position has a set of competing tiles with compute domains on left/right sides. **b.** Key concepts: (1) Tiles at high excess concentration over the scaffold, ensuring each position gets a tile. (2) Errors (mismatching-colours) have an enthalpic penalty that leads to their correction via replacement. (3) Weak-affinity compute domains, to minimise unintended interactions such as off-scaffold assembly. **c.** From an easily-prepared start state the computation eventually leads to an enthalpically favoured fully-bound target configuration. **d.** Abstraction levels. **e.** Strand-level abstraction of bind and replace primitives. **f.** Probability of target configuration at equilibrium for a given temperature via a partition function (PF) algorithm.^30^ **g.** Compute domains encode program instructions (top) and input bits (bottom). Parity program computes whether an 8-bit input `10100100`has an odd or even number of 1’s using program/parity bits `0`or `1`(top) and 4 input bit-pairs (`10, 10, 01, 00`, bottom). **h.** Reprogrammability by choosing strands from a large pool. Two Parity examples emphasise that output depends on every bit of input (orange panel: input `10100100`reports output `1`; gray: input `00100100`reports output `0`). Right: intended fluorescence readout cartoon, with no signal loss.

For programming and algorithmic information processing, tiles encode ≤ *ℓ* bits per compute domain. Setting *ℓ* = 3 gives 2^3^ × 2^3^ = 64 possible computing tiles per scaffold position.

SDC programming means selecting a subset of competing tiles per scaffold position that determine the left-to-right instructions computed per position. A typical *deterministic* computation has a single *anchor tile* at position *A*, multiple competing tiles at other positions but only one is correct per position. The molecular process of choosing the correct tile per position executes the computation.

A sequence of colour-matching tiles bound to the scaffold is a valid computation, or *target configuration*. Fig. 2g shows an example Parity program that computes whether the number of 1s in an 8-bit input is odd or even. There are 2^8^ = 256 possible input strings each corresponding to a different subset of 7 tiles. Each of four compute domains encodes two input bits, and a variable parity bit computed by correct attachments. In general, a target configuration either outputs a single bit at the last position (e.g. Parity), or an *N*-bit output along the entire sequence of tiles (e.g. Addition, below). Supplementary Note S4 expands on SDC computational theory, in particular S4.2 and S4.4 argue why mixing input and program bits on the same domain is computationally reasonable.

Fig. 2b shows three key principles yielding the target configuration (output) being energetically favourable: (1) Inspired by DNA Origami,^28^ computing tiles are at concentration excess (typically 10 *×*) over scaffold, to ensure every scaffold position gets a tile. (2) Correct compute domain bindings are enthalpically favoured over algorithmic errors. (3) Compute domain duplex binding strength is weaker than scaffold position domain. Custom^30^ minimum free energy (MFE) and partition function algorithms show the target having arbitrarily high probability over all other scaffolded configurations (Fig. 2f, Supplementary Note S3.5.1), using energetics estimated from our designed DNA sequences. Mathematically, the system scales well: compute domain strengths need only be logarithmic in *N* (Supplementary Note S3.5.2). The SDC model meets the abstract goal in Fig. 1b.

At the more concrete strand-level of abstraction, the *scaffold strand* has *N* +1 position domains (extra domain for reporting), and each *compute strand* has two 12-base compute domains flanking a 24-base position domain (Fig. 2d). Fig. 2e shows the tile *bind* operation as hybridisation of the compute strand to a scaffold position and any adjacent matching compute domains. In Fig. 2e, we intentionally leave DNA base-level kinetic details of *replace* unspecified: it can be *any* pathway that swaps scaffold-binding strands. We expect there to be many plausible, temperature-dependent, pathways.

Under bind/replace kinetics the expected computation time is upper-bounded by *O*(*N*^2^), the time for the leftmost compute domain mismatch (error) to disappear by a random walk to scaffold position *N* assuming a single anchor tile (Supplementary Note S3.3). However, although we defined one reasonable kinetic model, we intentionally do not experimentally enforce any particular kinetics, a *laissez-faire* approach that stands in contrast to DNA computing using toehold-mediated strand displacement where precise domain-level or even base-leve kinetic pathways are engineered.^16,31^ We hypothesised that the combination of thermodynamic favourability and multiple plausible kinetic pathways would enable a straightforward temperature anneal to overcome kinetic traps, unlike previous carefully optimised systems that use temperature holds.^13–16,18,20,21,23,24,32^

To avoid unintended hairpin formation within compute strands, each 3-bit sequence has two distinct compute domains (e.g. `000` and `000`, see Supplementary Note S5.1), giving 2 × 2^3^ = 16 distinct compute domains for *ℓ* = 3. A quenched-fluorescence reporting mechanism, operating at any scaffold position was designed. By convention, quenched (low) signals report output bit 0 and unquenched (high) signals bit 1. Fluorescence- and quencher-labelled strands are 20 bases long. *N* = 4 scaffold positions, and *ℓ* = 3 bits per compute domain implies 201 computing strands: one 120-base scaffold (named scaff-120, synthetically synthesised and purified), 64 strands at each position *B, C* and *D*, and 8at anchor position *A* (unpurified). An additional 64 strands for reporting (purified), and 130 for renewable programs (unpurified), gives a pool of 395 strands for *N* ≤ 4, plus an additional 1,144 for *N* ≤ 25(Supplementary Note S5.5). We used a thermodynamic-based sequence design approach for compute and reporting domains, building on previous work^13^ with details in Supplementary Note S6. The synthetic scaffold scaff-120had domains from M13, and other longer scaffolds used biologically-sourced subsequences of M13 bacteriophage^28^ chosen to give reasonably isoenergetic position domains. (see Methods).

## SDC Programming: Addition example

Any Finite State Machine (FSM)—an important sub-class of computer programs, amenable to molecular implementation^33^—can be compiled into a 1D SDC (Fig. 3a, Supplementary Note S4.1). The *N* = 4, *ℓ* = 3 SDC implements any 2-state 8-bit input FSM, such as 4-bit Addition (Fig. 3b). In ADDITION, each compute domain stores two input bits *x*_*i*_, *y*_*i*_, and a carry *c*_*i*_ computed, via tile/strand choice, as the mod 2sum, of the previous carry and input pair. The 4 output bits were read in 4 separate experiments as shown in Figs. 2h, 3D and Supplementary Note S5.2.

**Fig. 3.**
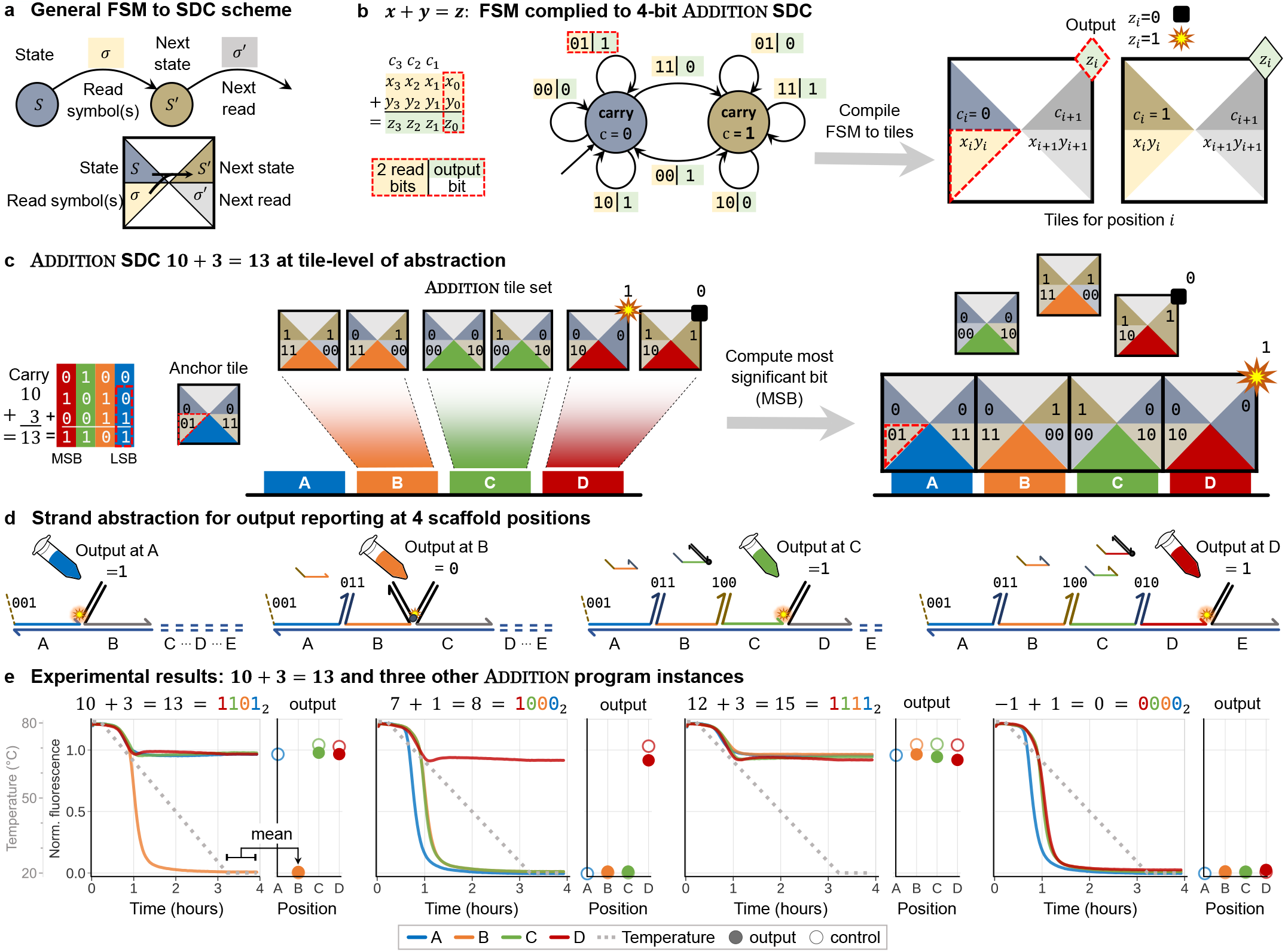
Programming and implementation of the Addition FSM. **a.** General compilation scheme from FSM into tiles, each compute domain encodes an FSM state and transition. **b.** The Addition FSM adds two binary numbers *x* + *y* = *z*: each transition reads two bits, *x*_*i*_ and *y*_*i*_, writes output bit *z*_*i*_ and enters carry state *c*_*i*_ ∈ {0, 1}. The Addition FSM is compiled to an SDC with two tiles per scaffold position. **c.** Example: 10 + 3 = 13, using colour to illustrate how bit positions map to scaffold positions. The target configuration is the most favourable as it is the only configuration with zero mismatches. **d.** Strand diagrams for reporting at each of positions A (output bit = `1`), B (= `0`), C (= `1`), and D (= `1`). Unused positions are covered with a single-domain complementary strand.**e.** Experimental result for 10 + 3 = 13, as well as three other Addition experiments. In each case the traces show signal with respect to time/temperature, and the dot plot shows completion level as mean of 22 datapoints at 20 °C.

To test the hypothesis that our thermodynamically favoured computing system could be run using simple annealing protocols, experiments were single-pot and simple, a so-called *typical-anneal* dropped from 80 °C to 20 °C in three hours, with scaffold at 100 nM, then was held for 45 minutes to obtain a signal completion level (Supplementary Note S7.1). Flat completion levels and strong signal suggested a lack of kinetic traps and signal loss (leak); Fig. 3e, analysis below.

Data was minimally processed: (1) Each raw fluorescence trace was normalised by dividing by the mean of its readings at 80 °C; (2) traces were averaged across at least 2 repeats; and (3) for scaff-120, traces were re-scaled to have the mean *completion level* of 0- and 1-reporting controls be 0.0 and 1.0, respectively; see Supplementary Notes S7.2.3 and S7.2.4. The rescaling step was not done for scaled-up systems using M13 for scaffold, where individual control completion levels are reported instead. Controls have one strand per scaffold position, and no per-position competition or computation; they should simply anneal to the ideal target configuration. The anchor position 1 is always non-competitive, and thus only a control.

We estimated SDC performance, or yield, in two ways: the ability to separate bit-0 from bit-1 and the proportion of correctly assembled target structures. Data for all typical-anneal computations for *N* = 4 are 100% linearly separable: all samples have completion levels closer to their target mean control level than off-target. We estimated yield as distance from completion levels to their target mean control level, or, for scaled-up M13 scaffold systems, their corresponding experiment-specific controls: *N* = 4 typical-anneals had mean 96.7% (SD=0.027) yield over all Addition inputs (the single worst Addition output bit gave 91.6%), with mean 95.3% (SD=0.035) over all computations, see Supplementary Note S7.2.6.

## SDC programming

We sought to test additional hypotheses about the *N* = 4 SDC system. First, we tested programmability on a further 9 programs, beginning with a simple BITCOPY system that acts as a length-4 wire (Fig. 4a). We programmed four FSMs, which were automatically compiled to tile/strand sets (Fig. 4b–e) from our stock, including the Parity program from Fig. 2. A single logical/computation step executes by an SDC choosing among competing tiles at a position. Programs tested thus far had two competing tiles per position (*B, C, D*), or *competitive complexity* 2. The MULTIPLYBY3 program multiplies by 3 in base 2, testing competitive complexity 3; average yields for competitive complexity 2 and 3 are similar: 96.2% (SD=0.033) and 95.3% (SD=0.041) respectively. 3-STATE Nondeterministic Finite Automaton tests competitive complexity 4, nondeterministically assembling up to four distinct target structures in parallel reported using distinct signal levels (Supplementary Note S7.3.5). DIVBY2 divides by 2 in base 3, requiring ternary digit reporting. We used our usual scheme for reading 0 and 1, and a novel, serendipitously-discovered, but carefully characterised, temperature-dependent spurious quenching technique for reading ternary digit 2 (Supplementary Note S5.2.2). Fig. 4f–i tests programs with competitive complexity up to 8, without significant signal loss: average yield is 95.1% (SD=0.019), see Supplementary Note S7.3 for analysis. Finally, a 20-hour postanneal temperature hold showed neither leak nor slow completion, Supplementary Fig. S27.

**Fig. 4.**
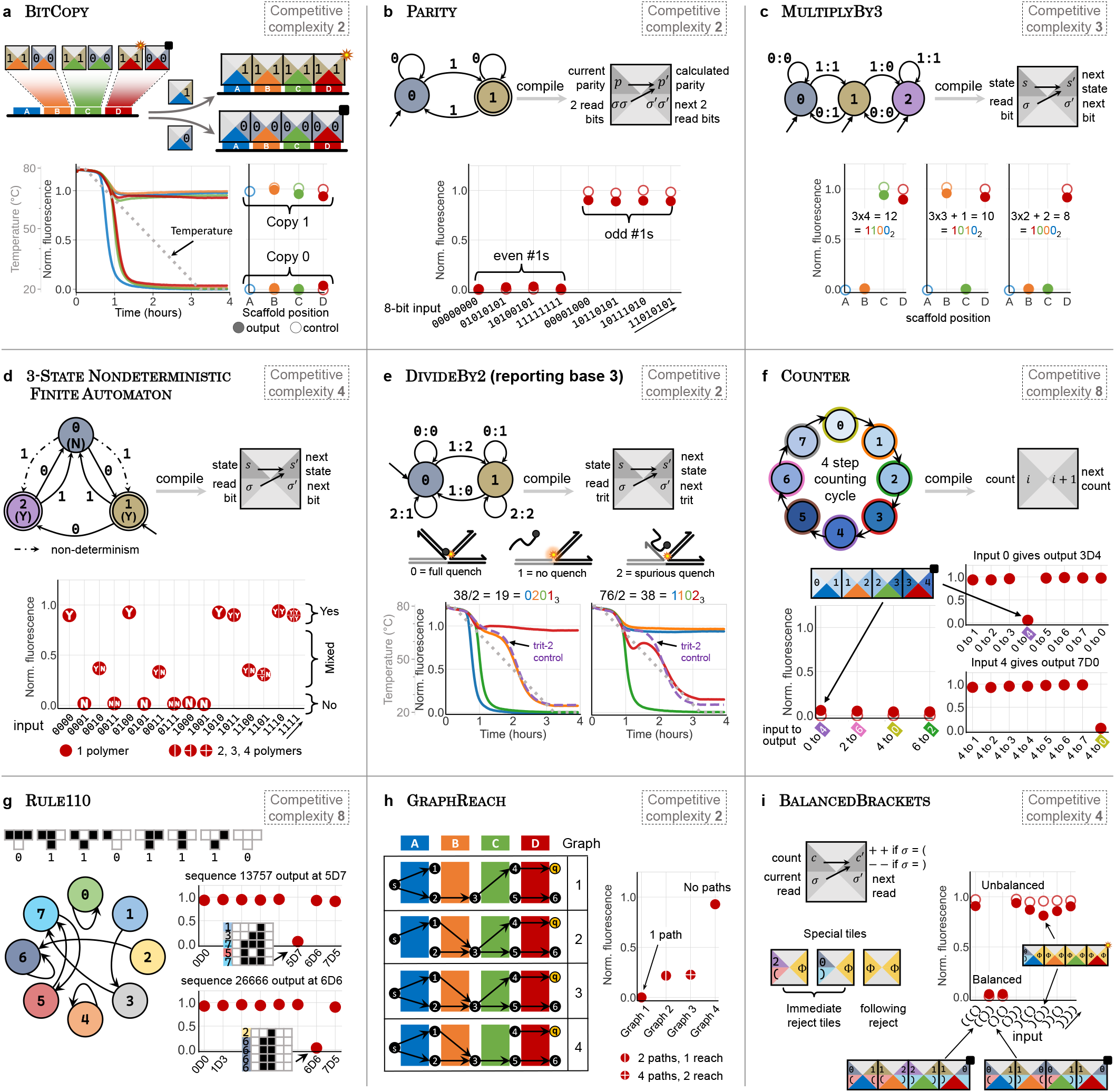
Programming 10 SDC programs of increasing complexity. **a.** Simple BITCOPY program that acts as a distance-4 wire; 8 traces show copying of input `0`or `1`, to positions *A, B, C* or *D*. **b.** Parity program: 8-bit binary input (2 input bits per compute domain), outputs `1`if the number of `1`s is odd and outputs `0`otherwise. **c.** MultiplyBy3, multiplies a base 2 number by 3. **d.** 3-STATE Nondeterministic FiniteAutomaton: each 4-bit input causes some fraction of Yes/No outputs (giving different ratios of target structures). **e.** DIVBY2 divides a 4-digit base-3 number by 2, giving output in base 3 (ternary) using a novel temperature-dependent spurious-quenching system. Ternary output ‘`2`’ should roughly track the *trit-2 control* in dashed-purple (i.e. orange curve in 38/2 = 19 (left), red in 76/2 = 38 (right). **f.** Given a value *x* ∈ 0, 1 *…*, 7 in binary, Counter counts *x* + 4 steps (e.g. input 2 at *A* yields output 6 at *D*). **g.** Rule110 simulates 4 steps of a 3-bit instance of the cellular automaton Rule 110 (with boundary condition `0`on the sides). **h.** GRAPHREACH: finding if there is a path from start vertex *s* to goal/quench vertex *q* by forming a target structure for each path from *s*. **i.** BalancedBrackets: accepts balanced brackets, e.g. (()) and rejects unbalanced brackets, e.g. ())): increments for an open bracket and decrements for a matching close bracket, quenching only if the count is 0 at *D*.

## Renewable SDC computations

We hypothesised that our method of programming SDC equilibria could facilitate *renewable* programs: simply switch equilibrium by adding two strands that swap old input *x* for new *x*′ (Fig. 5a,b). The number of renews is limited only by the concentration decrease per re-run, which can be made arbitrarily small in the limit of large initial volume, small renew volume, or low concentrations.

**Fig. 5.**
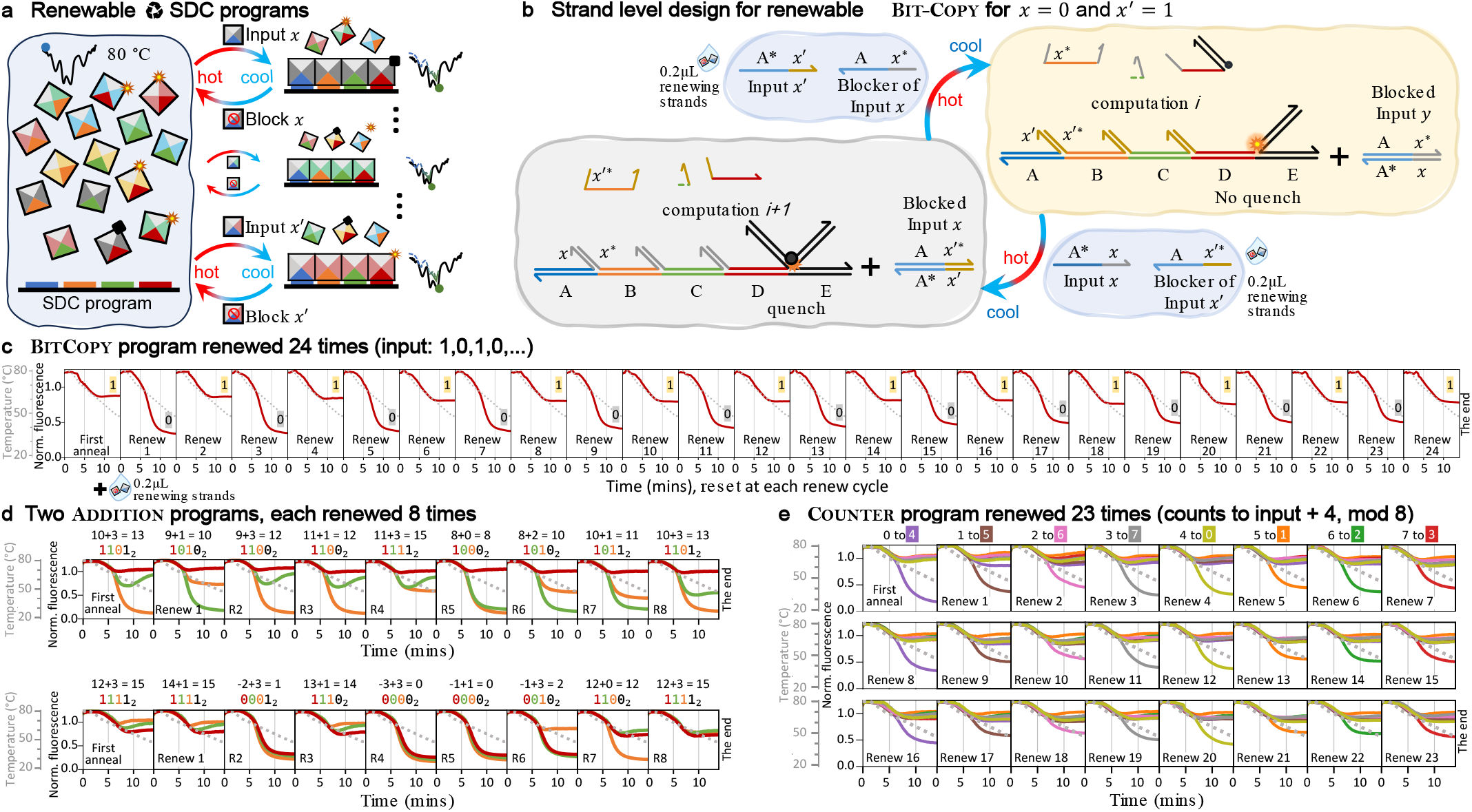
Renewable SDC programs. **a.** Principles for renewable SDC programs: At each cycle, to switch the target equilibrium we add new input *x*′ and blocker (complement) of old input *x* and reanneal to quickly reach the new target equilibrium. **b.** Strand-level renewable design: at each cycle add two strands, and reanneal. **c.** Renewable BITCOPY data: 25 runs, starting with input bit *x* = `1`, repeatedly cycling between input bit-`0`and `1`as in panel B: the blocker and new input were added at ~5.7× over scaffold (1× = 100 nM) in a 0.2 *µ*L droplet at each renewing cycle (0 mins). Sample was reannealed from 80 to 48 °C in 12 mins (holding at 80 and 48 °C for 1 min (shown), and (not shown) a 3–15 mins gap for inter-renew sample-handling). **d.** Two renewable Addition programs: to facilitate renewing a large number (8) of distinct inputs at position *A*, the programs have 8 tiles competing at *B*, see Supplementary Note S8.2). **e.** 24 runs of renewable Counter that counts from input *x* to *x* + 4. Although conceptually simple, Counter has high competitive complexity with 8 tiles competing at scaffold positions *B, C, D* for base-8 counting. Each plot shows 8 samples, each sample ran 24 times: row 1 had 8 distinct inputs *x* ∈ {0, 1, *…*, 7} in each of the 8 samples, then row 2 and 3 repeat those 8 inputs in order (192 traces total).

Fig. 5c shows BITCOPY run once and then renewed 24 times, flipping between input bit`-1` and `0`. Each renewal added 0.2*µ*L volume containing two strands using an acoustic liquid handler. For convenience, we ran fast 12 min anneals; lowering completion relative to typical 3-hour anneals, however the 0/1 signals were clearly separated. Renewals showed remarkably low signal degradation over time (we expected signal loss due to (i) dilution, (ii) buildup of strands/errors distorting the intended target equilibrium, (iii) insufficient equilibration time). We sought to renew more complex programs. Two ADDITION programs were renewed 8 times, Fig. 5d, each had 8 tiles compete at position *B* allowing 8 distinct inputs at position *A* per program (see Supplementary Note S8.2). Finally, to evaluate a high competitive complexity program on a large number of renewals, we renewed COUNTER on 24 successive inputs, Fig. 5e. Despite 8 tiles per non-anchor scaffold position, and somewhat fast anneals, all 24 renews gave a clear output, with mean signal loss of 17.3% (SD=0.067) every 8 repeats, see Supplementary Note S8.2.4.

## SDC speed and size scale-up

We hypothesised that fast computations might be possible for short scaffold lengths, *N* ≤ 4. *Super-fast* anneals, dropping from 80 °C to 55 °C in under 1 minute resulted in clear separation of bits 0 and 1 (Fig. 6a, Supplementary Note S7.2.5). This demonstrates the SDC can compute as fast as one minute, or even half a minute, the fastest in the DNA literature and almost at our lab equipment speed-limit. In terms of our yield performance metric, super-fast anneals have mean 82.4% (SD=0.116) yield at 1-minute for Addition and 81.2% (SD=0.16) over all computations, and outputs are linearly separable with a perprogram-threshold, Supplementary Note S7.2.6.

**Fig. 6.**
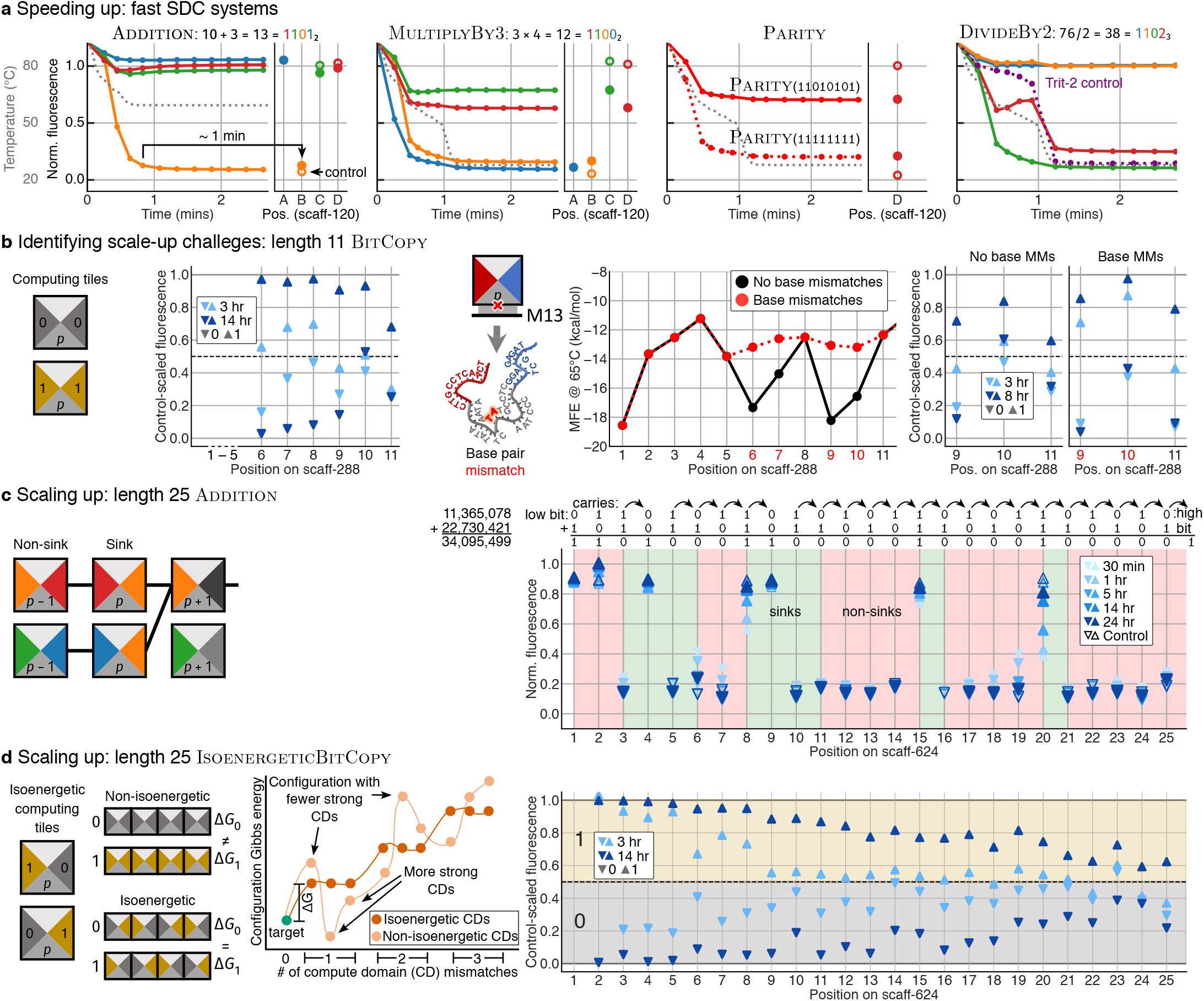
Speed-up and system scale-up. **a.** Super-fast anneal (3 mins) for 4 programs, with the dot plot showing readout at ~1 min. **b.** Scale-up of Bitcopy program to an 11-domain scaffold (scaff-288), yields are improved by longer anneal times, and DNA-base mismatches to improve the scaffold energy landscape. **c.** Scale-up of Addition program to a 25-domain scaffold (scaff-624). The program logic uses sinks/non-sinks to provide explicit error-correction by absorbing carry bits in green regions of the plot, which truncates error pathways in the energy landscape. **d.** The Isoenergeticbitcopy program was explicitly designed to have isoenergetic configurations: each scaffold position’s competing tiles have the same set of domains, but ordered differently. The 25-domain experiments used scaff-624. In (b) and (d), high and low controls at each position allowed sample readouts to be scaled per-position between their respective controls, while in (c), controls are shown directly.

It is tempting to interpret fast kinetics as evidence that scaleup to *N*> 4is possible. As Supplementary Note S3.3 shows, if the SDC actually obeys the bind/replace kinetics proposed in Fig. 1, computations would take time merely bounded by *O*(*N*^2^); the time for all compute domain mismatches to disappear by random-walk to position *N*. However, theory and naive optimism need tempering by some practical challenges.

We designed a rather ambitious and harsh scale-up test. Our choice of biologically-synthesised 7.2kb M13 meant having long and easily obtainable scaffold, but brought interesting non-idealities to the SDC model: (1) Scaffold domains have widely varying free-energies. (2) Experiments needed to run at 10 nM scaffold concentration, 10× lower than before, saving material but worsening signal-to-noise and lowering tile onrate. (3) Potential unintended interactions from thousands of superfluous single-stranded M13 DNA bases. Additionally, all except two labelled strands were unpurified, to save time/cost. We initially tested an *N* = 11 BITCOPY system, on a 288 nt M13 region called scaff-288, chosen for backwards compatibility with scaff-120. Increasing anneal time from 3 to 14 hours improved results. To combat poor M13 scaffold domain energetics, we employed DNA base mismatches,^34,35^ giving a clear yield improvement from 39% to 57% on 11-positions over an 8 hr anneal (Fig. 6b, Supplementary Notes S9.2 and S9.3).

To achieve further scale-up, and improve kinetics we used thermodynamic sequence design principles to choose a good 624-base segment (Supplementary Note S9.1). But even our best choice, scaff-624, had large variability in scaffold domain duplex Δ*G* of −20.9 to −11.1 kcal/mol at 65°C, compared to designed compute domains (−9.42 to −6.61 kcal/mol).

Computational theory suggested two more systematic approaches to programming the energy landscape. First, SDC tile programs embed logic with good error-reduction properties. A pair of Addition inputs typically has positions where a `0` or `1` carry bit is absorbed; a *sink*. Non-sinks pass carries along increasing tile competition; making non-sinks more prone to logical bit-flip errors. Let *M* denote the longest run of non-sinks for a given input pair. Addition has low *M* values: for example, two 25-bit inputs have mean *M* of 3.0 (SD 1.1), meaning that its logical structure should be conducive to error suppression. An *M* = 3 computation is computed in only an hour, Supplementary Note S9.4. A harder *M* = 5 example in Fig. 6c has good completion within 14 hours, noting that *M* ≤ 5covers over *>* 9% of pairs of 25-bit numbers, hence almost all of the *M*-hardness of ADDITION. Supplementary Note S9.4 has details and other computations.

Second, we developed a program-independent approach to isoenergtics: The ISOENERGETICBITCOPY program has the target configuration for inputs `0` and `1` being almost isoenergetic, since both reuse the same compute domains, respectively encoding `0101..01` and `1010…10`.This form of computedomain relabelling can be systematically applied to many SDC programs, enforcing configurations with *k* compute domain mismatches to sit on a roughly flat energy plateau with Δ*G* of *O*(*k*)(Fig. 6d, Supplementary Note S9.5). This gave excellent results, copying a bit across 20 positions with 71% yield, or 25 positions with 59% yield, on a 14 hr anneal. DNA base mismatches were unable to further improve yield significantly.

## Discussion

The conceptual step forward in Fig. 1 enabled robust, fast, renewable and scaled-up thermodynamically favoured molecular computing. The core concept was elucidated on 10 SDC programs on over 700 computations with binary input of up to 50-bits, as well as base-3 strings and graphs. Unlike out-of-equilibrium molecular computers, the SDC did not need explicit error correction via redundancy/scale-up^13,15,27,36–39^ nor compute-strand purification, precisely-calibrated temperature holds,^13–15,37,38^ multi-step manual protocols^16,20–24^ or early experiment termination to avoid leak. Unlike previous computing systems, the SDC is concentration-robust, inheriting the beautiful principle from DNA Origami^28^ of having staple (compute) strands in large concentration excess over the scaffold. We even repeated one experiment after 1.5 years, adding water as the 96-well plate had partially dried out, giving the data reported in Fig. 6a, MultiplyBy3 and Parity.

Intuition suggests equilibrium computation could be imprecise, since any desired output configuration competes with exponentially many off-target structures. We outline thermodynamic and kinetic arguments against this intuition for the SDC. Our thermodynamic design principles engineer the predicted target (MFE) structure to have probability approaching 1.0, driving the sum of probabilities of all other structures to 0 (Fig. 2f, Supplementary Note S3.5.1). Indeed, the correct SDC output was reported in all experiments. Notably, average completion/yield on typical-anneals, at scaffold length ≤4, was % (SD=0.035) of strict controls with fewer strands and simpler equilibria. Mean yield at scaffold positions *B* and *C* was 98.1 % (SD=0.014), dropping to 94.4 % (SD=0.034) at *D*. Increasing competitive complexity from 2 to 8 did not significantly degrade yield for typical 3-hour anneals.

For larger *N*, mere thermodynamic favourability may not be sufficient: we need a nicely-shaped and traversable energy landscape. The abstract SDC tile model gives a stepped landscape, efficiently explorable by random walks that ratchet forward via compute domain mismatch repair. Hence, in theory and practice, we took a somewhat liberal approach to kinetics. The bind/replace model in Fig. 2b and Supplementary Note S3.3 begins a computation by guessing one of a large set of 2^*O*(*N*)^ initial states, yet by mere random walk completes in only *O*(*N*^2^)time. However, experimentally, we did not enforce this particular kinetics, nor any other: we simply annealed the system, hypothesising that something akin to bind/replace is happening at higher temperatures. We used a strand design without intentional kinetic traps (Fig. 2), but did not carry out design optimisations, nor did we seek to characterise and prevent off-pathway interactions, approaches common in typical non-equilibrium molecular computing.^13,15,16,20,21,23,24,37,38^ Our approach is perhaps akin to that in DNA Origami^28^ where assembly kinetics is not precisely controlled yet works beautifully.^40–42^ Super-fast 1-minute anneals ran at the speed limit of our laboratory equipment, and beat all previous speed records for non-trivial DNA computations.^26,43–45^ Results for our super-fast, and typical, anneals suggest the SDC has fast, high-temperature, reversible assembly kinetics (Supplementary Note S3.6). Scaling-up to to scaffold length 25systems required slower anneals. The awkward energetics of cheap biologically-derived scaffold (M13) slowed computation at some positions, but in a way that seems not unlike our proposed random-walk kinetics (Supplementary Note S3.3). We conclude that the SDC is robust by design despite much remaining to be understood.

Although the earliest DNA motor was reusable,^46^ designing renewable molecular computers is a major challenge due to system complexity and out-of-equilibrium operation.^47–53^ Here, three SDC programs were successfully renewed up to 24 times at a reasonable speed of 12 mins per annealing cycle (plus time to add inputs).

Overall the principle of thermodynamic favourability provided a simple and effective method for program renewal, beating previous records in: (i) number of cycles (we show 25 runs, previous was 6 for a small system,^50^ or 3^47,48^ and 2 for larger systems^49^ or resetting the same computation^52^), and (ii) time per renewal,^48^ without need of manual steps^48,49^ or extra molecular technology beyond DNA^51,53^.

Thermodynamic computing principles are applicable to nanoscale engineering: Scaffolded DNA Origami^28^ could be endowed with computational abilities^54^ to algorithmically drive shape formation by leveraging work on folding dynamics,^41,42^ or scaffolded DNA data storage platforms, like SIMD *||*DNA^24^ could have thermodynamic biases against errors. Inherent robustness to temperature and concentration, and lack of leak/errors, could facilitate thermodynamic computing in complex biological environments, engineered in RNA or protein. Indeed, we used a biologically-sourced scaffold sequence as a proof-of-principle for under-designed DNA sequences.

Our work shows that DNA systems are amenable to ‘freeenergy landscaping’ enabling equilibrium molecular computing to move from theory to practice. More theory^1,4^ is needed to understand molecular equilibrium computation. The thermodynamic binding network model^3^ provides some principles, and entropy-driven equilibrium programs have been implemented.^25^ Also, equilibrium computing by strand commutation^26^ encodes Boolean circuit layers as a cascade of weakly bound, base-mismatching, strands. The energy landscape is challenging to control due to DNA binding model uncertainties, but theory may show how to amplify the signal.^55^ Fuel-consuming, infinite-time, out-of-equilibrium molecular dynamics^1,19,46^ are, by definition, unsuited to thermodynamically favoured computation. However, even there, energy landscapes can be sculpted to encode complex, finite-time, dynamics while driving to a ground state—a fixed-count chemical oscillator^19^ being an example. Beyond molecular programming, the theory of thermodynamics of computation^1–4,8^ seeks to determine ultimate energetic costs for computation. The SDC has several costs (number of synthesised strands, strand length, heat/annealing, etc.), but so far has not required explicit error-correction scale-up/time/energy/material costs, which seem to dominate computing architectures be they molecular,^13,15,27,36–39^ classical digital-electronic^2,4,8,9^ or quantum.^56^ With this work, we hope to stimulate dialogue^11^ towards consideration of thermodynamically-favoured, energyefficient, computing platforms.

## Methods

## Sequence design

For scaff-120, we intentionally chose **not** to use *de novo* designed DNA sequences for the scaffold since one goal is to prototype a system that is robust to sequence choice, even biologically sourced. We chose to have four scaffold position domains (*A, B, C, D*), and the additional reporting position *E*, of 24-base each giving a 120-base scaffold sequence, scaff-120, somewhat arbitrarily chosen to be two contiguous subsequences of biologically-sourced M13 (see Supplementary Note S6). For scaff-288, we moved to a subregion of M13, using full M13 in solution instead of a synthetic strand, and used the same domain sequences from scaff-120, (*B, C, D, E*), extended by eight 24-base contiguous M13 domains scaling up the computation length to 11 positions, (see Supplementary Note S9.3. For scaff-624, a 624-base subregion of M13 (still using full M13 in solution) was chosen based on criteria given in Supplementary Note S9. We used a thermodynamic sequence design approach^13^ for compute and reporting domains. For the compute domains, we used a 3-letter ATC code with at most one G as an exception, motifs CCCC, GGGG, AAAAA, TTTTT were forbidden, and isoenergetic complementary binding interactions of −11.7 ± 0.1 kcal/mol (chosen simply by our metric). −11.6 is the computed mean for a pool of 10,000 length 12 sequences. As soft constraints (i.e. enforced with potential exceptions), orthogonal interactions (unintended binding) *>* −2 kcal/mol evaluated using `binding(A,B) = pfunc(A,B)-pfunc(A)-pfunc(B)` as in reference^13^‘s SI Section S4.2.1 p. 43, with `pfunc()` given by`NUPACK4`.^57^ Secondary structure within each sequence compute domain was optimised to be *>* −0.25 kcal/mol. Reporter domains were designed using similar principles (isoenergetic binding between −21.7 and −21.95 kcal/mol at 53 °C and no G was permitted to be within 4 bases of a fluorophore.^58,59^ Software packages`nuad`^60^ and NUPACK4^57^ were used for DNA sequence design.

## DNA synthesis and fluorophore/quencher labelling

All 1,539 DNA strands were ordered in 384-well plates and 96-well plates from Integrated DNA Technologies (IDT) unpurified and normalised to 200 µM in IDTE pH 8.0 Buffer except for 9 strands which are the 120-base synthetic scaffold (Ultramer™ DNA Oligo PAGE purified and dry, see Supplementary Note S7.7), the four labelled strands (HPLC purified and normalised to 100 µM in IDTE pH 8.0 Buffer), and the four strands that the fluorophore-labelled strands bind to scaff120for reporting (PAGE purified and normalised to 100 µM in IDTE pH 8.0 Buffer). Of the four labelled strands, two had an ‘ATTO590’ fluorescent label at the 5′ prime end and the other two an ‘Iowa Black® FQ’ quencher label at the 3′ prime end.

## Sample mixing and buffer conditions

Most experimental mixes were automatedly generated using a custom code pipeline (Cosmix); some were prepared using alhambra-mixes library.^61^ SDC programs and their controls were mixed using an Echo 525 acoustic liquid handler (Beckman Coulter, supplied by Labplan Ltd. Ireland) for the data reported in Figures 3–6, and by hand for some of the data in the Supplementary information. Also, earlier versions of Figs. 3–5 were mixed by single/multi-channel hand pipettes. The liquid handler is not necessary to run SDC programs, but we found it gave more consistent results (lower signal variance) than hand-mixing. Picklists specifying the liquid transfer sequence and volumes from the 384-well source plates to the 96 destination plates were automatedly generated using Python. Mixes using synthetic scaff-120were prepared with a scaffold concentration of 100nM, while scaff-288and scaff-624directly used M13 at 10 nM concentration. Scaffold concentration in all experiments is set to the standard 1× baseline. All systems used 10× of every competing strand, 0.93 × of the fluorophore binding strand,0.79 × of the fluorophore strand and 17.86× of the quencher strand. DNA strands for an SDC program were placed in a single 0.1 mL 96-well PCR plate (or tube) to a volume of 35 *µ*Lin 12.5mM Mg++ in a tris acetate EDTA (TAE) buffer with0.01 % tween buffer.

## Fluorescence spectroscopy and temperature protocols

Bulk fluorescence data was run in a quantitative 96-well PCR machine (QuantStudio™ 5 Real-Time PCR System operated using the open-source python library `qslib`^62^) for fluorescence measurement over time. All experiments used the ATTO590 with excitation/emission at 593/622 nm. Seven plates were used to run the experiments of the ten programs shown in Figs. 3–4 including controls and repeats. Anneals essentially consist of dropping temperature from 80 °C to 20 °C over durations ranging from 30 seconds to 24 hours, then holding at 20 °C where *completion levels* are computed, see Supplementary Note S7.1. Often, temperature is dropped at a quicker rate in the 75 °C to 55 °C range than in the 55 °C to 20 °C range. All temperature protocols are made available, see “Data and code availability” below. As discussed in Supplementary Note S7.2.3, the data undergoes minimal processing. Performance metrics calculations are discussed in Supplementary Notes S7.2.3, S7.2.4 and S9.2.

## Data and code availability

Python code and libraries for sequence design, Echo mixing automation, qPCR protocols, and data analysis and processing are available as supplementary files to this submission, at doi:10.5281/zenodo.15869378.

## Acknowledgements

We thank Ahmed Shalaby for fast partition function and MFE code used for thermodynamic prediction, and Angel Cervera Roldan for his work on an SDC kinetic simulator used in sequence design. Also, Boya Wang for sharing data on tween; Chris Thachuk, David Soloveichik, David Doty and Doan Dai Nguyen for helpful conversations on theory and design, and Erik Winfree, David Soloveichik, Lulu Qian, Kim Reilly, Nathan Ronceray, Sébastien Ohleyer, for valuable feedback. Funding: Research supported by the European Union’s European Research Council (ERC, Active-DNA, no 772766); European Innovation Council (EIC, DISCO, No 101115422); and Science Foundation Ireland (SFI) under grant numbers 18/ERCS/5746 and 20/FFP-P/8843. Views and opinions expressed are however those of the author(s) only and do not necessarily reflect those of the European Union, European Research Council, European Innovation Council or Science Foundation Ireland. Neither the European Union nor the granting authority can be held responsible for them.

## Author Contributions

All authors participated in all aspects of the work: TS, AE, JA, CGE, DW designed experiments, carried out data analysis, and wrote the paper; TS, CGE and DW focused on theoretical concepts, AE, JA on experiments, TS, DW on DNA sequence design, DW, CGE on thermodynamic and kinetic analysis.

## Author Information

Correspondence should be addressed to, and. The authors declare competing financial interest: TS, AE and DW are listed as inventors on pending patents.

## Notes

https://zenodo.org/records/15869378

### Competing Interest Statement

The authors declare competing financial interests: TS, AE and DW are listed as inventors on pending patents.

