## Supplementary Information for "A Thermodynamically Favoured Molecular Computer: Robust, Fast, Renewable, Scalable"

Tristan Stérin[*,†,‡], Abeer Eshra[*,‡], Janet Adio[*,§], Constantine Glen Evans[*,‖], Damien Woods[*]

[*]Hamilton Institute and Department of Computer Science, [§]Department of Biology, Maynooth University, Co. Kildare, Ireland. [†]prgm.dev, 9 rue des colonnes, Paris, France. [‖]Evans Foundation for Molecular Medicine, Pasadena, CA, USA [‡]These authors contributed equally. [*]Corresponding authors.

### S1 Thermodynamic favourability of SDC: an overview

Section S1.1 gives a brief overview of what we mean by thermodynamic favourability in the SDC and provides justification for some assumptions in the model. Section S1.2 gives some historical context.

#### S1.1 Thermodynamic favourability of the SDC at all abstraction levels

Roughly speaking, we say that a model of computation is *thermodynamically favoured* if two rather desirable conditions are met.

- The first is that given any program and input, the intended output is energetically preferred over all other configurations, in the intended operation conditions. Thus, if we design it correctly, the computer will eventually energetically roll downhill to give the output.
- The second is that the initial configuration(s) should be easy to prepare. Specifically, the input should not need special out-of-equilibrium preparation.

These conditions assert that special out-of-equilibrium conditions are neither needed to prepare the input nor to read the output. Hence, the system should not require pre-annealing, purification, pre-computation or other kind of manual/special preparation for some special out-of-equilibrium input state. Nor does it require us to read the output at some special time before errors happen.

For example, for a DNA computer, there can be two different temperatures—the initial and target temperatures—such that the initial and target configurations are preferred, respectively. The input state might simply be all strands of the system, melted, at thermodynamic equilibrium at a high temperature. We cool the system down, and the output configuration becomes the most favoured one; assuming good kinetics, we eventually get the answer. Or, another example: maybe we care less about kinetics and prefer to hold at a single temperature, in that case, the input might be prepared by throwing all DNA strands together into a test tube at that working temperature, and we just wait to get the output.

To make these two conditions more precise, one needs a model of computation, including its physics, in mind, as detailed in this work. Also, there are a large number of other concerns that need to be met and investigated: having output with a large enough probability to be easily observable, input states that are easy to prepare, good kinetics, computational expressiveness and programmability. Here, we address all of these issues in the formalism of the SDC tile and strand models.

**Tile-level.** When we claim that the SDC tile-based model is thermodynamically favoured, we mean the model definition satisfies the two conditions above. Specifically, the input configuration has all components (tiles, scaffold) initially unbound, which means the input configuration can be prepared by simply mixing those individual components together, and it is a corollary of the model definition that the target, or output, configuration is the most favoured configuration and is eventually reached in any computation. The latter claim is mathematically proven in Lemma S3.1 in Section S3 for the kinds of programs studied in this work. The tile-based model is intentionally oversimplified by having unit-strength binding, no notion of temperature, an rigid scaffolded structure, and ignoring entropy. In Section S3 the model is defined mathematically and some theory of computation is developed, including details on programming. Hence, to believe the claim carries through to implementation, we need to provide believable justifications at lower abstraction level(s). We do this by stepping down the abstraction hierarchy and showing that each level implements, in a believable way, the assumptions of the level above. The next level down is the SDC strand level model.

**Strand level.** Below the SDC tile model, we have the SDC strand model (mathematically defined in Section S3.4 and with some implementation-specific design details in Section S5). **We give our main theoretical and simulation-based arguments for thermodynamic favourability in the language of the SDC strand-level model**, also Section S1.1.1 gives some physical justification for our thermodynamic claims. At the strand-level, we define an energy model that includes entropy of binding (i.e. a strand association/binding initialisation penalty $\Delta G_{\text{assoc}}$ and a strand concentration penalty for taking a strand out of solution), real-valued binding energies $\Delta G(\text{domain})$, and each configuration (strands bound to a scaffold) has an associated free-energy $\Delta G(\text{configuration})$ with configurations weighted by the Boltzmann distribution over their free energies, where more negative free-energies are more favourable. In our theoretical and computer simulation results, for simplicity we assume that all compute domains have equal binding free-energy $\Delta G$(compute domain), as do all scaffold position binding domains $\Delta G$(scaffold position), and that $\Delta G$(scaffold position) $\leq \Delta G$(compute domain). Assuming low enough temperature, and strong (negative) enough scaffold position domain binding energies to overcome a strand association penalty (i.e. $\Delta G$(scaffold position) $< \Delta G_{\text{assoc}}$ plus a concentration penalty, Theorem S3.3 shows that scaffold and compute domain strength scales nicely, only logarithmically in scaffold length. To better illustrate this, we simulated the strand-level model [30] over a wide temperature range (see Figure S3 below, or Figure 2f in the main text). This simulation used domain free-energies estimated from our DNA sequences via thermodynamics-based nucleic acid software NUPACK4 [57]. The simulation shows, for a variety of scaffold lengths, that at low enough temperature the target configuration has probability approaching 1, outweighing the sum of the probabilities of all other configurations despite their being exponentially many of them in scaffold length $N$ (see computer simulation in Section S3.5.1 and mathematical proof in Section S3.5.2).

**DNA sequence level.** At the DNA sequence level, we sought to implement the constraints of the strand-level model so as to preserve its thermodynamic favourability. This included avoiding self-binding within compute strands, avoiding unintended binding between pairs of compute domains/strands, etc. See Section S6 for details.

**DNA implementation.** We set experimental conditions (concentration excesses, temperature anneal, etc.) so that the strand model is faithfully approximated, in a classical Gibbs free-energy model, with enthalpy trading off against entropy in the usual way that is defined by the Boltzmann distribution (see also Section S7.1). As discussed in more detail in the main text, the successful demonstration of the SDC gives direct evidence of the favourability of the target (fairly high completion levels, no sign of 'leak' or other errors, no sign of completion not being reached on the one hour timescale of typical experiments, fairly good results with a minute on super-fast anneals, the fact that there are no slow kinetic traps even despite not explicitly designing against that too strongly). In addition, to compare to other systems: (1) Section S3.6 gives equilibrium-based and kinetic-based arguments as to why the so-called toehold-mediated strand displacement (TMSD) mechanism, that is common in DNA strand displacement systems, is likely not the main driver of SDC computations (despite compute domains "kinda looking like" toeholds). (2) Also, the fact that a simple, and even super-fast anneal, can be used to drive computations forward is strong evidence that the SDC is quite unlike the other popular form of DNA computing: algorithmic strand displacement. (3) Point (1) is also an argument that the form of SDC computation is quite different from systems that run TMSD on top of a DNA origami surface. See Section S2 for a more detailed comparison with other models.

##### S1.1.1 Justification of some physical assumptions of the SDC strand-level model

First, **we assume that strands (or tiles) and domains are indestructible**. On large timescales, such as the lifetime of the universe, and even on much smaller timescales where covalent bonds break down, no experimental SDC implementation would maintain its strand/domain structure and thus the claim of enthalpic favourability evaporates, perhaps literally—this is an unavoidable implication of the second law of thermodynamics. However, we assert **our assumption is justified in the usual, implicit, sense that underlies all work to date in molecular computing and DNA nanotechnology**: the experimental system is only used on time-scales we are interested in (minutes, hours, days, weeks, or even months) where enough domains/strands remain intact (we say "enough" since due to compute strands being in 10x over scaffold, the SDC system can handle quite a lot of strand degradation; also degrading of scaffolds simply lowers the yield in a predictable way). Indeed, these assumptions are analogous to those used when considering the thermodynamics of information processing (see Section S1.2), i.e., it is assumed that the computer and its fundamental components remain intact. Secondly, by insisting on the principle of significant excess of compute domains over scaffold (usually 10x in our experiments), the system is in fact somewhat robust to slow domain/strand degradation, and likely significantly more robust than other DNA and molecular computing paradigms (see Section S2).

Our second assumption is that we can set experimental conditions so that the SDC tile model, and **the SDC strand model are faithfully approximated**, in a classical Gibbs free-energy model, with enthalpy trading off against entropy in the usual way that is defined by the Boltzman distribution. It is a straightforward matter to choose scaffold position domain DNA sequence lengths, compute domain DNA sequence lengths, temperature, and salt concentration so that our DNA sequence designer enforces that the various constraints listed above on domain binding strengths hold. In fact, although we carefully designed the compute domain DNA sequences, we intentionally chose non-designed DNA sequences (from the M13 scaffold) as a proof-of-principle to use longer scaffolds that would need to be biologically sourced due to present-day limitations on DNA synthesis techniques (sequence design details in Section S6).

#### S1.2 Context with previous work on the thermodynamics of computing

One of the primary goals of the field of thermodynamics of computing is to understand the energy requirements for computing, a topic that is perhaps coming back in vogue due to the overwhelming burden compute power places on our energy infrastructure [9, 63]. Work in this field provides inspiration and background for our work, but it is important to note the distinctions between the goals in that work and our own, and not to confuse the two.

**The field of thermodynamics of computing** The field of thermodynamics of computing takes ideas from classical thermodynamics, statistical mechanics (including chemical reaction theory) and computer science to (a) characterise and elucidate the elusive and necessary energy requirements for computation, and (b) within that framework to give techniques that minimise energy usage during various kinds of computation. Highlights from this scientific history include Landauer [8] asking about thermodynamic costs of computing and considering the thermodynamic energy cost of reducing entropy, focusing on bit erasure since erasing a bit from memory (i.e. setting the value from an unknown `0` or `1` state, to a known/certain `0` state) is a way that programs reduce the entropy of a computer's memory. This was followed by Bennett's inspirational work using clever algorithms to show that arbitrary computations can be made reversible (i.e. forward *and* backward deterministic, no need for bit erasure nor any other form of irreversible operation) [4, 64, 65], albeit with increases in time/space usage. Bennett noted that with a small energetic cost per step, one could bias the forward direction so as to put a constant probability on seeing the completed computation (output) at equilibrium, although it was later observed that one can do the same without the bias (but perhaps with an additional, relatively small, cost to switch input) [66]. Other algorithmic work on reversible computing continues to this day [10, 67].

In molecular computing, researchers have been inspired by the work of Bennett and others, and there have been various molecular designs and implementations for reversible computing devices in theory [1, 68–70] and practice [23, 46, 50, 71] (although most systems do have at least some forward, albeit small, bias, or require manual resetting, which has implied energetic costs, etc.). More recent theoretical work on the thermodynamics of computation attempts to analyse computations using non-equilibrium statistical physics, leading to the definition of other thermodynamic costs beyond bit-erasure [1, 72]. Work is ongoing in this field, and will likely continue given humanity's concerns around energy wastage in computing [63, 73, 74]. A more detailed comparison between the SDC and molecular programming systems is given in Section S2.

**Energy, and other resource, costs** Although it would be interesting to do so, here we do not attempt to precisely estimate the necessary and sufficient energetic costs of SDC computing, nor to try to optimise (minimise) such costs in our DNA-based implementation, although it is worth noting that costs might include, at the very least, the length of our DNA sequences (synthesis costs), the number of strands, heating and temperature annealing control of the sample, shining light on samples for readout. However, we do note that as a corollary of being thermodynamically favoured, our system dispenses with the need for some energetically expensive error-correction costs or other costs involved in holding typical computers in an out-of-equilibrium, or far-from-equilibrium, state. Previous DNA computing systems handled error correction/leak using incredibly clever techniques, but techniques that cost extra strands [13–15, 36–38] (proofreading), and/or longer strands [27, 32, 39] (leakless designs). We had no need for such costs here.

The SDC presents a new paradigm of computing that leaves much room to analyse the trade-offs between various resources (time, robustness, strand excess, domain length (in DNA bases), number of scaffold domains, number of compute domains, energy use per experiment) with respect to other forms of DNA/molecular computing, including algorithmic self-assembly, strand displacement systems and surface-mediated computing (see Section S2).

**Ising models** Finally, it should be noted that, at the higher abstraction levels, the SDC can be thought of as a kind of Ising model [75, 76]. Indeed, many of the concepts of the SDC model, and many of its properties that we analyse, can all be expressed in the language of Ising systems, including locality, configuration/state space, partition function, minimum free energy, kinetic pathways and annealing protocols. It should be noted that the SDC would be a somewhat unusual Ising system in that it has information-encoding and scaffold-position encoding bonds that may have varying bond strength (at the SDC tile and SDC strand levels of abstraction), there are spurious interactions between orthogonal bonds at the SDC sequence level, there are a variety of concentration-related experimental constraints at the implementation level. Hence, if one were to think about the SDC model in the Ising formalism, it may be most relevant at the higher levels of abstraction and may prove most useful in dimensions higher than one. We note that there has been a significant amount of work on the computational complexity of Ising systems [76], and there may be fruitful connections between such results and the SDC model to be made.

**Algorithms inspired by thermodynamic energy-based principles** The long and intertwined history between computer science and physics is exemplified by the large number of algorithmic ideas inspired by thermodynamics and ubiquitous energy-based models [77]. A simple example with a decades-long history is simulated annealing [6] takes a straightforward approach, where candidate solutions to a search problem are assigned an energy, with better solutions having lower energy. A randomised algorithm navigates the solution landscape, with a time-varying temperature parameter controlling stochasticity: typically starting at high temperature to widely traverse the search space while avoiding getting trapped in local minima, then lowering the temperature to optimise within a minimum. As with *all* algorithmic ideas, success is a function of the match between algorithm and problem, with computational complexity theory being the ultimate arbiter on what is efficiently solvable [76]. That is just one example, there has been a huge amount of research on Energy-based models, see for example [77], with the cannon including simple models like Hopfield neural networks and Boltzmann machines, as well as more complex deep-learning and generative models.

There is also an inspiring history of using computer science ideas to show the hardness of problems in physics [78], by showing that those problems, whether we like it or not, embed algorithms. Since computation represents some of the hardest, yet reasonable, dynamics we can imagine, this endows those physics problems with unpredictability [76].

### S2 Comparison between SDC and other DNA computers/nanostructures

This section details (a) how previous thermodynamically favoured DNA nanostructures do not compute, and (b) why almost all previous forms of implemented DNA computation are not thermodynamically favoured. We begin with the former.

#### S2.1 Thermodynamically favoured, but does not compute!

##### S2.1.1 DNA origami

Rothemund's DNA origami technique [28] has achieved a solid reputation as a low-error assembly technique for engineering 2D and 3D structures out of DNA [29]. That reputation is built on the inherent thermodynamic favourability of the target structures provided by the beautiful design principles of having a low-concentration scaffold that seeds/touches exactly one copy of each high-concentration staple species. Likewise, some designs that predate DNA origami also have an enthalpically favoured target structure [79, 80], while other work [81] utilises the scaffolding principle (but scaffolds complex multi-stranded tiles that complicate thermodynamic arguments).

However, from the molecular computing point of view, DNA origami (i.e. scaffold plus staple strands) simply **does not compute**![1] In particular, each pixel in a target shape can be uniquely assigned a short DNA sequence that specifies that pixel and nothing else—there is no form of function computation nor any computationally complex notion of input-to-output mapping other than uniquely addressed "hardcoding". This type of hardcoding stands in contrast to known molecular computing methods such as algorithmic self-assembly where each pixel in a target structure could have one of several possible associated sequences/strands, the choice of which is determined by (a) a short information-encoding input (e.g. a short bit sequence) and (b) a sequence of information-processing steps (e.g. via Boolean logic gates) leading to the assembly of that pixel [13].[2]

Important caveat, to avoid confusion in the literature: Although published DNA origami designs do not perform computation, origami structures have been used extensively as a nanoscale structural support in various studies on DNA computation, examples include:

1. DNA origami as an information-encoding structural-support *seed* for algorithmic self-assembly systems; including systems where the origami seed encodes inputs with 2 to 6 bits [13, 15, 37, 82]. Here, **DNA origami seed does not compute anything**, it simply provides a place to start the computation from, and the attaching DNA tiles perform the computation.
2. DNA origami being used as a (flat) substrate upon which Boolean DNA circuits are assembled [20, 21, 83–85], and (somewhat relatedly) upon which DNA walkers trek or signals move [22, 23, 86–91]. Here, the **DNA origami structure does not compute anything**, it simply provides a structural support to place computing strands, or walkers, the computation/walking runs on top of the origami.

Our proposal stands in contrast to that prior work since in our SDC system, computation occurs in the process of forming the scaffolded structure.

#### S2.2 Computes, but not thermodynamically favoured!

Almost all experimental work in DNA computing falls under this category, whether it be algorithmic self-assembly, DNA strand displacement circuits or DNA circuits on a surface. Two notable exceptions are a recent exciting manuscript [25] on implementing the thermodynamic binding network model [3] and a recent

[1] Many papers in DNA/RNA/protein nanotechnology and engineering use the word "programming" to refer to the *design* capabilities enabled by the use of information-based polymers. However, in this work, when we talk about computing with DNA, we mean something stronger: the ability to encode an *algorithm* in the DNA molecules. We do not wish to debate what is or is not an algorithm, but for here it suffices to say that when claiming something as computing we believe the claim should be backed up by reference or characterisation in terms of either a well-defined model of computation, such as finite state machines, Boolean circuits, Turing machines, register machines, or some other mathematically-defined, theoretically justified, model of computation that captures what it means to compute a function/process, or some such formal object (whether already existing in the literature or perhaps completely novel).

[2] For brevity, we omit a more formal and nuanced analysis of what is and is not computing in this context.

equilibrium-based system that uses non-perfect complementary binding [26]. We elaborate in the following subsections.

##### S2.2.1 Algorithmic self-assembly

From its inception in the PhD work of Winfree [92], algorithmic self-assembly has been a driving force in the field of DNA computing. Experimental algorithmic self-assembly systems [93, 94] have seen significant progress showcasing self-replicators [15], binary counters [37] and a system with twenty-one 6-bit programs on over 100 distinct computations [13]. A related system that reuses tile types in a similar fashion to algorithmic systems, but also beautifully exploits nucleation, is capable of recognising patterns (vectors) of tile concentrations [14]. Work to date has crucially relied on the establishment of several key principles [95] that took almost two decades to develop [36, 38, 92, 96, 97] but are now, arguably, reaching fruition. The topic has benefited from significant theoretical underpinnings elucidating the computational and expressive power of such systems [98–100], with distinct flavours of complexity, from careful studies on the self-assembly complexity of building shapes [101, 102], to various novel ideas linking computation and self-assembly [103–105], to a complexity theory called intrinsic universality that has been used to tease apart the power of self-assembly models [106–109]. Perhaps we are getting to a time where imaginative ideas from the minds of theorists and experimental practice are starting to meet. There remains much to do in this exciting research direction.

**Inspiration** Our tile-based SDC formalism inherits many ideas from algorithmic tile assembly, perhaps primarily the idea of the tile sides encoding information, computational universality of tile structures, sequence design principles, etc. Some of the challenges of algorithmic self-assembly have provided major inspirations to the SDC, including wanting to avoid spurious nucleation and the need for precise operating conditions. However, there are some key differences that the SDC takes to molecular computation, and by doing so avoids some of the challenges for algorithmic tile systems, as we now detail.

**Spurious nucleation** Spurious nucleation of off-scaffold structures is a fascinating major challenge and open research theme for tile-based algorithmic self-assembly [13, 82, 96, 97]. A technical issue that is specific to algorithmic self-assembly, as implemented to date, is that after a correct computation has grown—from the seed all the way to completing its computation—there are necessarily large excesses of tiles in solution, which are available for the error mode of spurious nucleation. In correct computations, these tiles are of a special type that should not be bound into any structure whatsoever; they must be left free-floating. Otherwise, the system is in an error state. For example, running a bit-copying nanotube from ref. [13] on input bit-string `000000` would mean that the intended final system state has (i) collection of seeds copying the string `000000` ("the correct answer"), and (ii) a huge number of excess free-floating `1`-tiles in solution that should not be bound to anything ("the unused tiles, only useful for other inputs"). Unfortunately, those `1`-tiles can be considered an out-of-equilibrium system by themselves, sitting, waiting to eventually spuriously nucleate, at the growth temperature (because the system is being held in growth-favourable conditions, temperature, tile concentration, etc.). This kind of error mode is avoided by the SDC, since (a) SDC compute tiles only bind weakly to each other (hence after all scaffolds are coated with tiles there is nothing for them to bind strongly to) and (b) we would never drop to a temperature where such compute strands aggregate off-scaffold.[3] In fact, we can be much more liberal with temperature in the SDC, as the next paragraph explains.

**Fine tuning of conditions** Just like regular DNA origami, by exploiting a low concentration scaffold to tether a completed computation together, it should be impossible for large unwanted structures to spuriously form in the SDC (assuming the system is held at a temperature above that of the stable compute domain binding temperature), even for large SDC systems that would span a full M13 scaffold (7.2 kb). However, one could argue that in algorithmic self-assembly, one could super-precisely tune seed concentration, tile concentration, and the speed (i.e. slowness) of the annealing protocol so that for the subset of tiles used in the computation, the rate of spurious nucleation is extremely low, perhaps almost zero. The main point here is that such tuning is not required for SDC (we ran the SDC system at 10x excess of tiles over scaffold,

[3] Of course, one could try to mitigate this error mode, by (cleverly) designing a tile set that simultaneously uses up the intended tiles, and then has some gadget/trick to also consume the other tiles (perhaps along the lines of some theory work [103]).

and we did not finely tune temperature protocols in the way one needs to do for algorithmic self-assembly). Small SDC systems should be robust to large, e.g. factor 2, discrepancies in concentration, but very long scaffold lengths would present interesting challenges. Presumably, compute/scaffold domain strengths would need to be increased to handle concentration variation.

###### S2.2.2 DNA strand displacement circuits

TMSD circuits are another popular form of DNA computing [16, 19, 39, 46, 110–120], or see an early survey for principles [31]. Most implementations do not have a thermodynamically favoured target (output) configuration, which is one of the primary reasons for error ('leak') pathways in such systems. Recent work on enforcing kinetic [32, 39, 114, 117] and thermodynamic [27, 39] barriers to leak pathways hold great promise (see also Section S2.3.3). Here, with the SDC, our proposal is to aim for a purely thermodynamically favoured output; hence, by definition, the SDC system does not have a leak (all pathways lead to the desired output/target).

Despite SDC compute domains "looking like toeholds" (Section S5.4), in Section S3.6 we give arguments as to why we are likely not seeing toehold-mediated strand displacement as the main driver of SDC computation, serving to emphasise key differences between SDC computation and classical TMSD systems.

We should emphasise, that for some TMSD systems, e.g. [19, 113], the main intention is to build an out-of-equilibrium system, comparisons between those and SDC are less relevant than for systems where the aim is to do a typical, one-shot computation.

**Thermodynamic analysis of a TMSD seesaw AND gate** A simple seesaw AND-gate motif from Figure 2 of [16] can be used to illustrate the distinction between kinetic and thermodynamic computation. In Fig. S1 we compare the intended kinetic pathway against equilibrium. The top of Fig. S1 shows the intended kinetic pathway, and circuit output of $\text{AND}(\texttt{0},\texttt{0}) = \texttt{0}$, is as described in [16]. In contrast, we used NUPACK [57] to compute the system equilibrium: inputting the AND-gate and two input bit $(\texttt{0},\texttt{0})$ strands, with appropriate concentrations, temperature and buffer conditions. As expected, and can be checked by hand for this simple example, the NUPACK-predicted equilibrium conditions show the gate produces some output, (but the intended kinetic pathway for inputs $(\texttt{0},\texttt{0})$ does not). Hence, correct seesaw gate execution relies on irreversible, fuel-driven steps and precise kinetic control. Even with good kinetic control, we expect the gate to eventually leak and settle into a favoured, but incorrect, final state—a key trick with seesaw circuits is to read the output before that happens. In fact, it is quite a feat to design large systems that manage to stave off thermodynamic equilibrium for long enough to carry out sophisticated computations.[16, 118]

**SIMD||DNA** The SIMD||DNA model, introduced by Wang, Chalk and Soloveichik [121], is a scaffold-based system that uses TMSD for computation. The design endows DNA storage systems with DNA computation, and has been implemented experimentally [24], showcasing a binary counter and cellular automaton Rule 110 computations. SIMD||DNA nicely brings two fields together: DNA storage and DNA computing, it is suited to computing on DNA storage systems that store data using the 'nick pattern' of DNA strands bound to a long scaffold [122]. Although scaffolded, SIMD||DNA does not share the SDC's property of thermodynamic favourability of the target, and requires some manual experimental steps. Nonetheless, from the data storage point of view, SIMD||DNA, and earlier nick-pattern DNA storage systems [122], provide significant motivation and inspiration for the SDC style of computing on a scaffold.

###### S2.2.3 Other applications of DNA origami to non-thermodynamically favoured DNA computation

It is important to make a clear distinction between our proposed SDC implementation and the decade-long history of DNA computation and DNA walkers that use DNA origami as a support (citations in Section S2.1.1). Such systems still have assembly and computational-error problems as they are out-of-equilibrium systems: the use of DNA origami as a supporting structure only provides good thermodynamic benefits *for the support.* The actual computing structure / walker system built on top of the origami typically

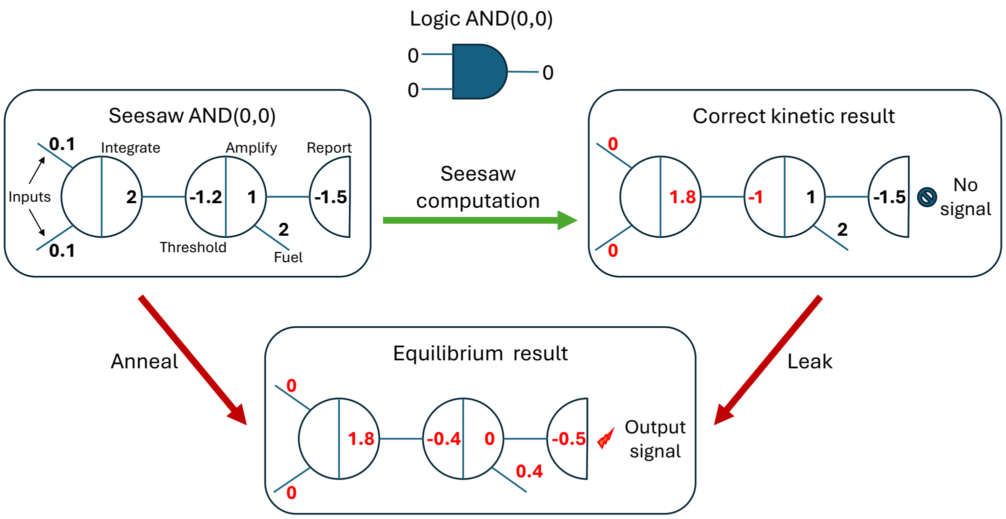

Figure S1: Illustrating a distinction between kinetic and thermodynamic DNA computation, using the example of a seesaw AND gate [16] with inputs (0, 0) encoded as two strands at concentration of 0.1× relative to a standard 1× concentration. The intended kinetic execution of the seesaw circuit (green path on top) gives the intended bit-0 output, encoded as a lack of reporter signal. However, the NUPACK-predicted [57] equilibrium is in an unintended, but thermodynamically-favoured, error configuration with incorrect output (the 'Anneal' arrow should produce this equilibrium, up to NUPACK modelling inaccuracies). The bottom right arrow shows leakage paths, where the system, if left long enough, drifts toward the equilibrium error state.

has implementation challenges requiring complex preparation steps in order to prepare the system in a high-energy state and/or prevent pre-triggering, for example [23].

#### S2.3 Thermodynamically favoured systems that compute

We said that almost all forms of experimentally implemented DNA computation are not thermodynamically favoured, here we mention a few recent exceptions that are paving the way for thermodynamically favoured molecular computing.

##### S2.3.1 Thermodynamic Binding Networks: a theoretical model of thermodynamically-favoured computation

Thermodynamic Binding Networks (TBNs) are a simple, and inspirational, theoretical model of thermodynamically favoured computation [3] that is gaining interest [123–125]. For simplicity of modelling, the model [3] intentionally ignores certain features: enthalpy, or domain binding strength, is given infinite weight over the entropic cost of bringing molecules together, and there is no physical geometry.[4] Such simplifications can be useful for (a) programming and (b) analysis of computational power by allowing one to intentionally ignore complications from kinetics, and (c) giving a mathematically simple setting to prove negative "no-go", results. In summary, the TBN model helps clarify to what extent a simple discrete model of thermodynamics can be used to drive computation.

The TBN model provides intellectual inspiration for our SDC model, but there are key differences, partially due to scaffolded DNA origami being a more direct SDC inspiration. First, the SDC tile-level and strand-level models include geometry; TBNs do not. Second, the SDC strand energy model has an inherent notion of temperature, and as a corollary, implicitly has the notion of weak glue/domain binding, which validates assumptions in the tile model, such as no off-scaffold interactions. But in TBNs, all domains are infinitely strong, meaning we can think of the system as operating in the limit of low temperatures or strong bonds.

[4]There is no geometry in the original and most-studied TBN model, although a variant with geometry has been studied [123].

Related to the second point, the SDC tile and strand models have a geometric embedding in 2D that disallows various forms of binding that would be allowed in TBNs. In particular, we allow no more than $N$ compute strands bound to the length-$N$ scaffold—one per scaffold position—thus, scaffold-bound compute strands are not permitted to bind to other compute strands in solution to form "hairy"/"branched" structures or "bridges of compute strands" between two scaffold-bound strands–features we would certainly see in favoured TBN configurations. In the SDC, we simply assume such TBN-like interactions are prevented by having low scaffold excess, and working at a temperature above the stable binding temperature of compute domains stably. Although we omit details, it can be argued that in our experimental working conditions, such interactions are unlikely as they have additional free energy costs, and we stay above the temperature where those costs are negligible.

Nevertheless, one can imagine taking any SDC and modifying it by adding enough distinct domain types to translate/compile it into some sort of TBN system, although without extra model assumptions, that TBN system would still have off-scaffold interactions and potential geometric non-idealities. Even a more geometric TBN model [123] would suffer from unwanted bindings, that we believe are not an issue in our experimental conditions. Thus, it remains as interesting future work to potentially develop simulation techniques for the SDC to implement some interesting subclasses of TBNs, or explore ways to use SDC principles to implement certain classes of TBNs in the wet-lab.

**Experimental implementation.** In a recent paper, Wang et al [25] implemented several TBN programs in DNA including two Boolean circuit designs and a polymer-based system that computes the lowest common multiple of two numbers. One of the circuit architectures has the property that TBN polymers are small ($O(1)$ size, independent of the input), giving a kind of "well-mixed" equilibrium circuit implementation, including the ability to do fan-in and fan-out. The other circuit design is seeded and grows a polymer whose size is linear in the size of the circuit. The main focus of the experimental work is to implement systems that exploit the entropy of mixing as the main driving force, under the assumption that the system has already maximised domain bindings. In contrast, our work here on the SDC is primarily driven by enthalpy through the binding of scaffold and compute domains. However, at higher temperatures, there are thermodynamic and kinetic issues at play: a multitude of similar-energy states, many with similar numbers of errors (entropy), are reachable in a reasonable time (kinetics), at reasonable temperatures. Lower temperatures both put a worse probability on bad states but also slow down the kinetics. In any case, this plurality of approaches in the TBN and SDC models, as well as the strand commutation approach (below), suggests there are, in fact, many ways to think about ways of thermodynamic computing with molecular systems: where and when each model should be used demands more study.

##### S2.3.2 Strand commutation using non-complementary binding networks

Nikitin [26] recently proposed a system called 'strand commutation' that uses partially-complementary strands: the idea is to have a set of strands that weakly bind to each other, and let that binding network go to equilibrium. Introduction of an input strand that binds strongly to many other strands will switch that equilibrium, which can be exploited to build logic gates. The system is novel, expressive and fast on small examples. Like the SDC system, the use of thermodynamics as the main driving force and an apparent lack of unintended kinetic traps in the examples implemented leads to rapid processing of information.

Although both strand commutation and the SDC utilise the idea of letting the system go to equilibrium, they have a number of key differences, in particular information-encoding is quite distinct between them, and the two systems program different kinds of equilibria: for SDC the system output/readout is designed to assess only scaffolded structures, and we have levers (compute domain binding strength, tile concentration excess) to drive, or program the intended/correct structure to be extremely favoured. In the strand commutation system, there is a more 'mixed' equilibrium where intended and unintended results may appear together in a mixed population, but with the intended winning the vote. With strand commutation, there could be room to design clever gadgets to suppress error, between circuit layers, possibly taking inspiration from kinetic proofreading for algorithmic self-assembly [36], or leveraging ideas from the Thermodynamic Binding Network model [3].

Also, as noted in [26], thermodynamic DNA sequence prediction algorithms, and their parameter sets, are likely more accurate for perfect duplexes than for partially complementary strands, or even for systems that use fully complementary partial-strand domains with nicks, overhangs, geometrically constrained loops, etc. There seems to be scope to improve that situation, with clever design/abstraction improvements such as those by Berleant [55, 126], or perhaps by using modern high-throughput laboratory techniques to find better DNA sequence parameter estimates. We should add the major caveat that both strand commutation, and SDC, can be modified/extended in so many ways that future systems could crossover to utilise features of both simultaneously. There remains space for a full theoretical characterization of the abilities and limitations of the approach.

##### S2.3.3 Leakless TMSD systems, with a thermodynamic barrier to leak

In Section S2.2.2, TMSD systems were briefly discussed, including that almost all such systems are out of equilibrium, and for one-shot computing (e.g. running a Boolean circuit), systems show some leak/error. Specifically, after we read the output, but then leave a TMSD system sitting for more time, the correct answer may get 'overwritten' by an incorrect answer. One way to deal with this is to specifically design TMSD systems with kinetic barriers to leak. Early work on leakless systems used redundancy to 'design-in' such giant kinetic barriers to leak [32], with some entropic bias [39], but more recent work [27] uses stronger thermodynamic barriers. So far, this principle has been shown on signal-transduction cascades, where an input signal is pushed through a linear sequence of translator gates. Future realisation of a computing platform, such as Boolean circuits, using these principles would present a beautiful method to use thermodynamics as the main driving force for computation while avoiding errors. Indeed, part of the beauty of such leakless designs is that they can be integrated into the 'compile chain' for existing systems (from abstract program specification to final leakless TMSD system), and that, from a theoretical prospective they can be used to give provable guarantees on error reduction, since leakless designs have a rigorous mathematical basis [27].

### S3 SDC tile model, SDC strand model and thermodynamics theory

This section makes a number of theoretical observations about SDCs.

#### S3.1 SDC tile model: definition

The SDC tile-level model is described in the main paper, and Figure 2 of the main text, and, in a bit more mathematical detail, here. Tiles are assumed to be unit-sized squares with colours on their left, bottom and right sides. The left and right sides of a tile are called *compute sides*, and the bottom side is called a *scaffold position side.* Let $\mathbb{N} = \{0, 1, 2, \dots\}$ be the set of natural numbers.

**Definition S3.1** (SDC tile program/instance)**.** A *SDC at the tile level of abstraction*, illustrated in Figure 2a–c, consists of a rigid 1D scaffold with $N \in \mathbb{N}$ distinct binding domains $d_1, d_2, \dots, d_N$ called *scaffold positions*, and a collection of $N$ distinct *computing tile* sets $T_1, T_2, \dots, T_N$, where the bottom *position domain* of each square tile in $T_i$ binds to domain $d_i$ on the scaffold. Tiles' left and right sides are called *compute domains* and two bind to each other if their colours match. Tiles' top sides are not used.

A *configuration* is $N$-tuple from $T_1 \cup \emptyset \times T_2 \cup \emptyset \times \dots \times T_N \cup \emptyset$, i.e. a list of tiles, with at most one for each scaffold position and the symbol $\emptyset$ is there is no tile at a scaffold position.

**Definition S3.2** (step, computation)**.** Starting from an empty scaffold (called the *initial configuration*), a *step* is where either (a) a tile binds at a matching empty scaffold position, or (b) a tile replaces another tile at a matching scaffold position but where replacement may only occur if the displacing tile binds with equal or more matching colours than the replaced tile. A *computation* is a sequence of steps, each one taking one configuration to the next. A *terminal configuration* is one to which no step applies.

A *program* and its *input* are encoded in tiles; details are given in the main paper Figures 2–6, as well in this SI Section. After specifying a program and input, the intended output configuration(s) is/are called the *target* or *output* configuration(s). Generally, a programmer would desire the target configuration(s) to be terminal.

An SDC is *deterministic* if it has a unique terminal configuration,[5] and *nondeterministic* otherwise (it reaches several terminal configurations and/or a cycle of configurations). If all terminal configuration(s) of a system have the same number of mismatches, and that number is the minimum over all configurations of the system, the system is said to be *thermodynamically favoured.* In other words, the system has no permanent kinetic traps on any path from the initial to the target configuration(s). Moreover, if all target configuration(s) of a system have zero mismatching domains, the system is said to be both thermodynamically favoured and *fully bound.*

**Design challenges implied by the tile model** As with other abstract models of molecular computing, e.g. [3, 54, 76, 92, 98, 99, 127–130], the tile-level of abstraction is convenient for programming and mathematical analysis of the computational capabilities of SDCs, but ignores quite a number of implementation challenges, including some of those specified in Figure 2b: viable molecular implementation of bind and replace operations, no off-scaffold tile-tile binding, no multi-scaffold structures. We leave these implementation challenges to lower abstraction levels.

#### S3.2 Thermodynamic favourability of the SDC tile model

The following lemma shows one sense in which the tile model favours the desired output, for the kinds of programs studied in this work. The lemma uses two hypotheses. The first is of "fairness", commonly used in asynchronous, distributed systems to avoid low-probability infinite, useless, loops and defined as follows: A *fair* computation is one where if there is a step (binding or replacement event) that can happen, then eventually will it happen; more precisely it can not be ignored for infinitely many computation steps. The second hypothesis is that the tile set has tiles that match each other on the sides (without this one can design systems with mismatching colours in any target state, which is not of interest in this paper). Simply stated,

[5] This notion of determinism is called directed in the abstract tile assembly model [102].

the lemma proves that target configurations are terminal and fully bound, meaning enthalpically favoured, i.e. what we want to happen. The question of whether that *will* happen in a more physically-realistic thermodynamic model is left to later sections.

**Lemma S3.1.** *Let $S$ be an SDC where, for $i < N$, for every tile $t \in T_i$, there is a tile $t' \in T_{i+1}$ where the right side of $t$ colour-matches the left side of $t'$, and where for all $j \in \{1, 2, \ldots, N\}$, $T_j$ is nonempty. Then, every fair computation in $S$ ends in a thermodynamically favoured fully-bound configuration.*

*Proof.* Since for all $j \in \{1, 2, \ldots, N\}$, $T_j$ is nonempty, by fairness, every computation $c$ reaches a configuration $C$ where each scaffold position has a tile. Let $0 \geq k \geq N$ be the number of mismatches in $C$. Consider the rightmost compute comain mismatch along the scaffold, and let $u$ and $v$ at scaffold positions $j, j+1$ be those mismatching tiles. Since, by a hypothesis of the lemma statement, each tile has a matching tile to it's right, there is a sequence of $N - j$ replacement steps to lower the number of mismatches from $k$ to $k - 1$: i.e. simply replace $v$ with the tile $v'$ that matches $u$, and so on to the right end of the scaffold. This argument can be repeated to give a (longer) sequence of steps that lowers the number of mismatches to $k - 2$, then $k - 3$, and so on, all the way down to 0, yielding a computation that ends in a fully bound configuration. Thus from any reachable configuration, there is *at least one path* to a fully bound configuration.

It remains to show that no other path gets trapped in a mismatching 'dead end' configuration, forever, for which we'll use the hypothesis of fairness. Suppose, for the sake of contradiction, that there is a computation $c'$ that never lowers its number $k' > 0$ of mismatches. We've already shown that $c'$ can not be of finite length (a 'dead end'), since we've shown that there is always an available, finite, sequence of steps to lower $k'$ to $k' - 1$ mismatches via replacement (which can be applied to get to 0 mismatches in finite steps). Therefore, it must be the case that $c'$ is infinite. Let $l$ be the largest index such that there are tiles at position $l, l + 1$ that mismatch in a configuration of $c'$. However, since the number of configurations on a finite length scaffold is itself finite, and since $c'$ has infinitely many steps, this implies that there is a replacement step that can happen at $l + 1$ that is being ignored infinitely often, hence the computation is not fair, contradicting the hypothesis. □

#### S3.3 Bind/replace kinetics for the SDC tile model

In this section, we propose a putative kinetic model, building on the kinetics in Figure 2b of the main text. Our goal is not to quantitatively explain the data, a task far beyond the scope of this paper, but rather to give a simple model that matches our intuition for how a perfectly-designed SDC might work and qualitatively shows agreement with our experiments. We leave it as future work to precisely parameterise this model or find a better one.[6]

Consider the tile-based model described in Figure 2a–c of the main text, and more precisely defined in Definitions S3.1 and S3.2. The initial state consists of a free scaffold, with tiles in solution (in excess), and where bind and replace operations happen. Lemma S3.1 says the system eventually reaches a target state, next we consider the time to get there. We augment Definition S3.2 with rates:

**Definition S3.3** (Bind/replace tile kinetics)**.** To Definition S3.2 we add rates for each step: The time for a bind or replace step is an exponential random variable with mean 1.

The resulting kinetic model is a continuous time Markov chain (CTMC), with the rate of the next event being the sum of the rates of all applicable events at that time instant.

Lemma S3.1 showed that any deterministic system (defined in Section S3.1) eventually goes to its target structure, the following lemma shows the expected time for that to happen under bind/replace kinetics. The energetic landscape is illustrated in Figure S2(a), and the artificially slow kinetic model used in the proof is simulated in Figure S2(b).

**Lemma S3.2** (Completion time)**.** *For a deterministic SDC that obeys the hypothesis of Lemma S3.1, with 1 anchor tile that binds to the scaffold position 1, and $O(1)$ tiles for each other scaffold position, the expected time to reach the target is $O(N^2)$.*

[6] In fact we believe that at different temperatures, different forms of kinetics are possible, we'd conjecture for one important fast kinetics comes from a model like what is suggested here, and perhaps some of the slowest, and most irrelevant, coming from low-temperature toehold mediated strand displacement (as discussed in Section S3.6, with data in Figure S4).

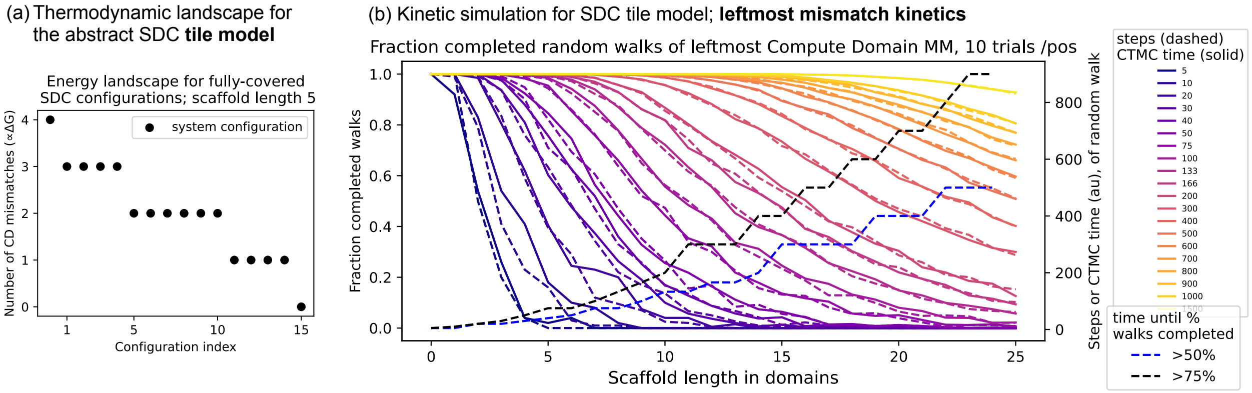

Figure S2: Two plots that illustrate the thermodynamics and kinetics of idealised SDC tile programs, such as BitCopy, assuming all domains are isoenergetic. These illustrate the proof of Lemma S3.2. (a) Simplified energy landscape for the SDC tile model, assuming a tile on each scaffold position. Each additional mismatch gives an enthalpic penalty, assuming compute domains are strong enough and perfectly isoenergetic (so several other enthalpic and entropic complications can be ignored). (b) Kinetic simulations for a model that is at least as slow as, and likely slower than, the bind/replace kinetic model. Brightly coloured curves show the fraction of completed random walks, of a mismatch starting at the leftmost position 0, on a length $N$ scaffold, within a certain number of steps, or time. The random walk models the migration of the leftmost compute domain mismatch, assuming that it never meets another mismatch. This simplified model ignores speedups such as parallel cancelling of compute domain mismatches, hence this is a kind of worst-case kinetics for the SDC bind/replace, nevertheless finishes in expected time quadratic in scaffold length $N$. The black and blue dashed curves show, for a given scaffold position, how much time to wait until that % of walks have completed.

*Proof.* We prove the $O(N^2)$ time bound for a simpler artificially slow process, one that is necessarily at likely slower than the modelled process (but certainly no faster). First, we allow only bind steps, by (artificially) preventing any replace steps, until the scaffold is filled with tiles. For a scaffold of length $N \in \mathbb{N}$, the expected time to cover the scaffold with tiles is $O(\log N)$, since each tile binding event happens independently and in parallel. Only after covering the scaffold do we allow replacements to occur.

In the exponentially (in $N$) unlikely event we get a scaffolded structure with $N$ tiles with all adjacent compute domains matching, we are done, but since that is unlikely we ignore that case and consider instead that there is at least one compute domain mismatch. Consider the leftmost such compute domain mismatch. We claim that the leftmost mismatch, over time, undergoes a random walk, that completes in $O(N^2)$ expected time, the proof is straightforward:

1. First, a compute domain mismatch between two tiles at positions 1 and 2 can not move to the left since there is only 1 anchor tile, which thus can not be removed by a replace step, hence we say that position 1 is a reflecting barrier.
2. Second, if the leftmost mismatch is between the tiles at positions $N-1$ and $N$, then there is always a replace step that replaces the tile at $N$ with a matching tile which removes that mismatch completely; in other words position $N$ is an absorbing position.
3. Third, if the leftmost mismatch is at some other position $1 < i < N$, it is repairable by a tile being replaced on the left or on the right of the mismatch: (1) if the replacement is to the left (respectively, right) and has a mismatch on its left at position $i-1$ (respectively, right at position $i+1$), we say the leftmost mismatch has taken a step to the left or, respectively right; else if (2) the new replacement tile (either at $i-1$ or $i+1$) has no mismatches on either of its compute domains then the leftmost mismatch of the system has either jumped to a new position $j > i$ (i.e. what was the second to left mismatch becomes the new leftmost mismatch), or we had started out with one mismatch and now there is no mismatch at all. Artificially, assuming the worst case where there are no jumps, the leftmost mismatch undergoes an equiprobable step to the left or right.

Hence, in our artificially slowed-down model, the leftmost mismatch undergoes a random walk until absorbed at position $N$, which is well-known to take $O(N^2)$ expected time. Hence the expected completion time of the full system, without artificial suppression of steps, is also $O(N^2)$. □

#### S3.4 SDC strand model: definition and energy model

The definitions and energy model in this section are simplified versions of those in a companion theoretical paper[7] [30], which also gives the fast partition function and MFE algorithms that we make use of in our analysis below.

##### S3.4.1 SDC strand model definition

The following definitions are strand-based concretisations of the more abstract tile model that was described at a high level in Figure 2a–c, and in detail in Section S3.1, with a more precise and detailed energy model. An $N$-position *scaffold strand* consists of $N$ concatenated *position domains*, or simply *positions*, numbered 1 to $N$, or labelled using letters $A, B, C, \ldots$. A tile with left and right compute domains $l$ and $r^*$, and position domain $p^*$, is implemented at the strand level as an abstract *compute strand* $s$ with three domains $s = d^{\mathrm{L}}(s)d^{\mathrm{M}}(s)d^{\mathrm{R}}(s)$ (i.e. left, middle, right domains of $s$), defined to run in a $5'$ to $3'$ order. The left and right domains are called *compute domains*, and the middle domain is called a *position domain.* Domains may bind if they are complementary, formally: domain $d$ binds to domain $e^*$ iff $d = e$.

**Definition S3.4.** An *$N$-position $\ell$-bit SDC at the strand level of abstraction*, illustrated in Figure 2d,e of the main text, consists of a scaffold strand with $N \in \{1, 2, 3, \ldots\}$ unique positions, and $N$ distinct sets of compute strands $S_1, S_2, \ldots, S_N$, where the position domain of each compute strand in $S_i$ has domain $i^*$.

In the strand-level model, the notions of *step, configuration, initial configuration, computation, terminal configuration, program, input* and *target/output* are the obvious and direct implementation of their tile-based definitions given in Section S3.1.

We highlight that this means that one major assumption in our strand model, and energy model below, is that **we ignore all off-scaffold interactions**; in particular each configuration consists of a scaffold with 0 or 1 compute strands at each position, nothing else. This simplifying assumption can be enforced at lower levels of abstraction our implementation, i.e. strand design level (Section S5), DNA sequence design (Section S6), and wet-lab implementation (Section S7). Also, we assume compute strands don't self-bind in any way. As shown in Section S5.1 there is a simple redundancy trick to avoid unwanted hairpins in compute strands, and good DNA sequence design can be used to prevent other non-intended interactions within a strand. Likewise we assume no self-binding of the scaffold.

##### S3.4.2 Strand-level energy model

Let $[s] \in \mathbb{R}, [s] \geq 0$ denote the concentration of a strand $s$. A domain $d$ has an associated, typically negative, real-valued free energy $\Delta G(d) \in \mathbb{R}$. Intuitively, in our simple strand-level model, the free energy $\Delta G(X)$ of a configuration $X$ is a sum of terms: for each bound domain pair $(d, d^*)$ there is a free energy of binding ($\Delta G(d) \leq 0$), while there is a per-strand concentration penalty $(-k_{\mathrm{B}}T \ln([s]u_0^{-1}))$, where $u_0 = 1$ M is a standard concentration constant, to account for the entropic cost of taking a strand from solution. Formally, the free energy $\Delta G(X)$ of a configuration $X$, where strands$(X)$ denotes the compute strands attached to the scaffold, is:

$$\Delta G(X) = \sum_{s \in \mathrm{strands}(X)} \left( \Delta G(d^{\mathrm{M}}(s)) - k_{\mathrm{B}}T \ln([s]u_0^{-1}) \right) + \sum_{s_i, s_{i+1} \in \mathrm{strands}(X)} \Delta G(d^{\mathrm{R}}(s_i), d^{\mathrm{L}}(s_{i+1})) \qquad (1)$$

where the latter summation sums the compute domain free energies of *scaffold-adjacent* strand pairs.

Since we assume a very low concentration for the scaffold, we assume that the concentrations of compute strands are constant, that is, they are not depleted by binding to the scaffold, and so the concentration of scaffolds, or the overall free energy of many complexes in a volume, does not need to be considered [131]. This is consistent with work on the kTAM where we assume a limit of low concentration for the seed (scaffold) [95]. All $\Delta G$ values are temperature dependent. Unlike some works that include a per-strand helix initiation term $\Delta G^{\mathrm{init}}$ or $\Delta G^{\mathrm{assoc}}$ [57, 131, 132], since we do not specifically model the junction between domains, and each bond forms a separate helix, we include the same helix initiation term in the $\Delta G(d)$ of each domain. Letting

[7] Many thanks to Ahmed Shalaby for helpful discussion, ideas and theorems that we use here.

$\Omega$ denote the set of all configurations of our SDC system, the strand-level minimum free energy (MFE) and partition function (PF) are the, respectively, (typically) negative and (typically) positive real-valued quantities:

$$M = \min_{X \in \Omega} \{\Delta G(X)\} \tag{2}$$

$$Q = \sum_{X \in \Omega} e^{-\Delta G(X)/k_{\mathrm{B}}T} \tag{3}$$

$Q$, the PF, gives a scaling factor that allows us to assign a probability to any configuration $X$, at some temperature $T$. For example, the probability of an MFE configuration is simply $M/Q$, which we sometimes write as MFE/PF.

#### S3.5 Thermodynamic favourability of the SDC strand model: computer simulation and theorem

This subsection has two components that justify the principles behind thermodynamically-driven SDC computation: (a) Computer simulation-based arguments showing that the target SDC structure is favoured over all configurations (indeed, has probability approaching 1.0) in our strand-level model, with strand binding energetics computed from out designed DNA sequences, and (b) a theorem proving that short compute domains suffice to have the target outweigh al other configurations.

##### S3.5.1 Strand-level thermodynamic analysis for scaling up 1: The target is favoured over all configurations by computer simulation

We wish to compare, at equilibrium, the probability of the target configuration (output) of an SDC system versus the sum of the probabilities of all other configurations, at the strand level. Leveraging the theory from Section S3, and Equations (2) and (3), those two probabilities are the quotient MFE/PF and 1−MFE/PF.

The set of all SDC configurations, $\Omega$, is typically of size exponential in the number of strands in the system.[8] For example, an $N$-position BitCopy system has $2 \cdot 3^{N-1}$ configurations since there are 2 possibilities for the anchor scaffold position $A$ (anchor strand is bound or not bound), and $3^{N-1}$ possibilities for the $N-1$ non-input scaffold positions which are each in 3 possible states (no strand bound, bit-`0` strand bound, bit-`1` strand bound). Hence a naïve and explicit evaluation of Equations (2) and (3) would take exponential time, which is infeasible for even modest scaffold lengths. Instead, we make use of the fast (polynomial time) and exact, dynamic programming algorithm developed especially for SDC systems [30][9] to generate Figure 2f in the main text, and Figure S3 below, as well as for some analysis in this subsection.

Figure 2f in the main text plots the quotient MFE/PF (Equations (2) and (3)), in the strand-level model with BitCopy SDC program, at resolution 0.5 °C for various scaffold lengths. Energy parameters for the strand-level model were as follows. $\Delta G_T$ for compute domains was computed, at every temperature $T$ in 0.5 °C increments, as the mean $\Delta G_T$ over all compute domain pairs, where the $\Delta G_T$ of an 8-base compute domain is the nearest-neighbour summation of the sum of it's 8 base pair stacks, using the nearest neighbour model and parameters from Santa-Lucia and Hicks [132], including a $\Delta G^{\mathrm{assoc}}$ penalty (Section S3.4.2). DNA sequences are in Section S10, and we used the `nuad`[10] function `nuad.np.wcenergy()` plus $\Delta G^{\mathrm{assoc}}$. Also, two corrections were applied since only doing what was described results in unreasonably high melting temperature of the target configuration (and for many other configurations), since it does not account for off-target interaction between scaffold-position-adjacent mismatching compute domains (which could favour off-target assemblies), nor a likely-large penalty for formation of the 3-way junction between a pair of

[8] Indeed, more generally beyond SDC, for systems with multiple strands where the number of strands is 'large' (part of the input), MFE determination is NP-complete and even hard to approximate [133]. Ours is a special case of that which is tractable [30].

[9] Java source code for the 1D SDC MFE and partition function algorithms [30] is available at https://github.com/ASShalaby/1D-SDC-SYSTEM-TMSD. These plots were generated using a Python implementation of the same algorithms, but with the addition of a logarithmic strand concentration penalty as defined in Eq. (1). For code see main text Methods section: *Data and code availability*.

[10] `nuad` is a DNA sequence designer developed by David Doty and others that allows specifying thermodynamic DNA sequence design principles such as those used in [13] and this work: https://github.com/UC-Davis-molecular-computing/nuad

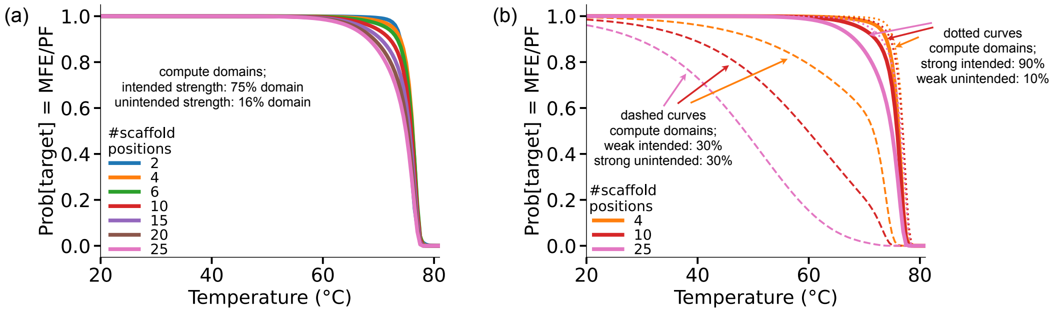

Figure S3: SDC strand-level thermodynamic analysis of a BitCopy system. Data points are plotted every 0.5 °C from 20 to 80 °C. Each data point shows the probability of the target, i.e. MFE divided by the partition function (PF), using the energy model in Section S3.4.2, for varying temperatures and numbers of scaffold positions, as well as varying parameters for matching (intended) and mismatching (unintended) compute domain interaction. (a) Plot in Figure 2f in the main text, but with wider x-axis. Domain strengths were computed from our designed DNA sequences (as described in Section S3.5.1) and then modified to account for 3-way junction penalty (75% of free energy of perfectly bound compute domains), and unintended/mismatching compute domain interactions (mismatches are assumed to bind with 16% of mismatch energy). (b) Exploring other values for these two corrections. Dotted curves have stronger intended binding and weaker unintended binding. Dashed curves have weaker intended binding and stronger unintended binding.

matching compute domains and two scaffold domains. Hence, as described in Figure S3(a), we applied two binding parameters to account for these unintended, but somewhat unavoidable[11], interactions. Figure S3(b) explores two extremes for the parameters. See main text Methods section *Data and code availability*, for Python code to generate the figures.

##### S3.5.2 Strand-level thermodynamic analysis for scaling up 2: A mathematical proof that compute domains need only be logarithmic in scaffold length

Here we provide mathematical justification for the previous computational analysis. In particular, the result proven in Theorem S3.3 gives a mathematical justification for the claimed thermodynamic favourability of SDC and for the nice scaling properties seen in Fig S3: i.e. that even with long scaffold lengths, the target configuration has probability approaching 1 at low temperature. As notational convience in the statement of Theorem S3.3, we asume the duplex scaffold domain free energy $\Delta G(\mathrm{PD})$ include an entropic strand concentration penalty, i.e. for a strand $s$ we let $\Delta G(\mathrm{PD}) = \Delta G(d^{\mathrm{M}}(s)) - k_B T \ln([s]u_0^{-1})$ and we define compute domain duplex free energy $\Delta G(\mathrm{CD})$ to not have that penalty: $\Delta G(\mathrm{CD}) = G(d^{\mathrm{R}}(s_i), d^{\mathrm{L}}(s_{i+1})$, using the notation from Eq. (1). The theorem assumes low temperature conditions such that $\Delta G(\mathrm{CD}) \leq 0$, $\Delta G(\mathrm{PD}) \leq 0$ and every scaffold position has a bound compute strand.

Intuitively, Theorem S3.3 says that as the number of scaffold positions $N$ grows, the strength (or DNA sequence length) of compute domains needs only grow logarithmically in $N$.[12] Since we are assuming low temperature conditions with a strand at each scaffold position, the also implies scaffold domains are no weaker than compute domains, so scaffold domain strength should also scale at least logarithmically in $N$.

**Theorem S3.3.** *Let $S$ be an $N$-position SDC with at least one compute strand per scaffold position. We assume all compute domains have the same duplex free energy, denoted $\Delta G(\mathrm{CD}) \leq 0$. Similarly, we assume all $N$ scaffold position domains have duplex free-energy $\Delta G(\mathrm{PD}) \leq 0$. We ignore configurations that have $< N$ strands on the scaffold. For notational convenience let $c = -\Delta G(\mathrm{CD})/k_{\mathrm{B}}T$ and $p = -\Delta G(\mathrm{PD})/k_{\mathrm{B}}T$. If $c > 0.37 + \ln n$ then $\Pr[\mathrm{target}] \geq 0.5$.*

*Proof.* Let $\Omega'$ denote the set of all configurations of the SDC $S$ that have $N$ compute strands bound to the

[11] Mismatches, or an unbound poly-T loop, around the 3-way junction could avoid the penalty, however one could see that as simply a more controlled method to achieve the same thing as the penalty we applied.

[12] Thanks to Ahmed Shalaby, Doan Dai Nuygen, and https://math.stackexchange.com/a/2652273/1318259 for invaluable assistance with the proof.

scaffold (i.e. one per position). The partition function (PF) for that class of structures is given in Eq. (3) as

$$Q = \sum_{X \in \Omega'} \mathrm{e}^{-\Delta G(X)/k_\mathrm{B}T} \tag{4}$$

Let $n = N - 1$. Observing that each configuration has some number $k \in \{0, 1, \ldots, n\}$ of mismatching compute domains, and that there are $\binom{n}{k}$ configurations with $k$ mismatching compute domains, allows us to rewrite $Q$ in terms of number of mismatches:

$$Q = \sum_{k=0}^{n} \binom{n}{k} \mathrm{e}^{c(n-k)+p(n+1)}$$

The probability of the target (no mismatches) being at least the probability of all other configurations (i.e. $\geq 0.5$) can be written as

$$\frac{1}{Q}\mathrm{e}^{cn+p(n+1)} \geq \frac{1}{Q}\sum_{k=1}^{n} \binom{n}{k} \mathrm{e}^{c(n-k)+p(n+1)}$$

Simplifying and adding $\mathrm{e}^{cn+p(n+1)}$ to both sides:

$$2\mathrm{e}^{cn+p(n+1)} \geq \sum_{k=0}^{n} \binom{n}{k} \mathrm{e}^{c(n-k)+p(n+1)}$$

$$2\mathrm{e}^{cn} \geq \sum_{k=0}^{n} \binom{n}{k} \mathrm{e}^{c(n-k)}$$

$$2 \geq \sum_{k=0}^{n} \binom{n}{k} (\mathrm{e}^{-c})^k = (\mathrm{e}^{-c} + 1)^n$$

with the last step using the binomial theorem. Simplifying, gives:

$$c \geq -\ln(2^{1/n} - 1) \tag{5}$$

In order to compute the right hand side of Equation (5), we begin by noting that $n \in \{1, 2, 3, \ldots\}$ and we will use $\lim_{n\to\infty} \frac{1}{n} = 0$, by first deriving:

$$\lim_{n\to\infty} n(2^{1/n} - 1) = \lim_{n\to\infty} \frac{2^{1/n} - 1}{1/n} = \lim_{x\to 0} \frac{2^x - 1}{x} = \lim_{x\to 0} \frac{2^x - 2^0}{x - 0} = \left[\frac{d}{dx}2^x\right]^{x=0} = \ln 2$$

Hence

$$\lim_{n\to\infty} \frac{1}{n(2^{1/n} - 1)} = \frac{1}{\ln 2} = 1.442\ldots < 1.45 \tag{6}$$

with the inequality holding for all $n \in \{1, 2, 3, \ldots\}$ since the limit in Equation (6) is approached from below (because $n(2^{1/n} - 1)$ increases monotonically and the limit exists: $\forall n, 0 < (n+1)(2^{1/(n+1)} - 1)) - (n(2^{1/n} - 1)) < 1$). By the inequality in Equation (6):

$$1.45n > \frac{1}{2^{1/n} - 1}$$

$$\Rightarrow \ln(1.45n) > -\ln(2^{1/n} - 1)$$

Combining that with Equation (5), we get the required logarithmic lower bound on $c$:

$$c > \ln(1.45n) = 0.37 + \ln n$$

□

#### S3.6 Strand-level kinetic analysis of experiments

##### S3.6.1 Introductory remarks and context

It is important to note that in this work we do not specifically prescribe, nor show that our system follows, particular kinetic pathways beyond the simple bind/replace shown pathways in Figure 2. The reader familiar with the beautiful concept [46, 110, 134] of DNA toehold mediated strand displacement systems (TMSDs) will likely have noticed that the compute domains in our strand-level model look like toeholds[13], and might immediately jump to the conclusion that we intend for the system to follow the typical TMSD pathway: **but we do not!** Existing TMSD systems are carefully designed so that such pathways are indeed the most plausible pathway for computation, and other pathways can lead to errors, called *leak* [16, 19, 20, 23, 31, 32, 39, 48, 51, 110, 135, 136]. Enforcing that a system computes by only, or mainly, using TMSD pathways builds on years of expertise and involves somewhat careful experimental and system control (including, typically, constant temperature, kinetically-optimised sequence design, optimisation of precise toehold design, etc.). Here, we intentionally do not do that, instead since the desired SDC target is thermodynamically favoured we don't really care which pathways the system uses to get to the target, so long as the system gets there. Although inconsistent with work on DNA computing, our approach is entirely consistent with the almost two-decade history of DNA origami [28, 29, 137]: heat it up and cool it down, or even run the system isothermally [42], without too much need for prescribing pathways (isothermal DNA origami folding/refolding can be quite slow though, and intentional use of toeholds can speed it up). This is not an unreasonable approach: researchers are still uncovering the folding pathways in DNA origami formation [40–42, 137]; but DNA origami has worked rather well despite those pathways being under-prescribed by the design, and still not fully-understood in practice! Our experimental setup is straightforward: we run a simple (non-optimised) temperature anneal from hot to cold, where we let the annealing process find a variety of, likely different and likely temperature-dependent, kinetic pathways to reach the thermodynamically preferred state.

Thus far in this subsection, we are describing what we want to happen–by system design– but in the remainder of this section, we give some data-backed evidence that our system is likely acting in such a manner, in particular *not* following the typical TMSD-style pathways. We acknowledge there remains much experimental and theoretical work to fully characterise this form of thermodynamic computing.

##### S3.6.2 Decisions are made above the compute domain melting temperature, and thus above typical TMSD operating temperatures

A key feature of TMSD systems is that they usually operate at a fixed temperature, somewhere around the toehold melting temperature [110]. That way, toeholds bind reversibly, which is a *key defining feature* of such systems.[14] Here, using a simple temperature-based argument, we hypothesise that compute domains do not act to drive a typical TMSD pathway as the sole kinetic pathway for SDC completion. The reasoning goes as follows.

The melting temperature for compute domains is around 48 °C, as estimated by NUPACK [57] (online version, accessed 7 Feb, 2024) with input: compute domain $d$ and its complement $d^*$ at 1 $\mu$M concentration each (the concentration used in our experiments), in 12.5 mM $Mg^{++}$ (and 50 mM NaCl, even though there was no NaCl in our experiments, but because NUPACK requires that as a minimum), complex size 2. At 48 °C we get 0.5$\mu$M of the $d$-$d^*$ complex (i.e. 50 %). Indeed, at 56 °C NUPACK estimates 0.08 $\mu$M of the $d$-$d^*$ complex (8 %), and at 63.5 °C only 0.02 $\mu$M (2%).

However, our standard anneals (an hour) show `0`/`1` decisions already beginning to happen above 65 °C in many cases, and in all cases the decision is 100% predictive of the final output by the time the anneal has dropped to a rather hot 60 °C. Intuitively, before the system gets down to 60 °C, more scaffolded polymers have voted one way (e.g. `0`) than the other (e.g. `1`).[15]

Control data shows melting mean temperatures of 63.6 °C for an entire scaffold with 2, 3 or 4 positions i.e. read at scaffold position B, C or D (see Figure S9 for an explanation and ss of good friendshiptrand diagram for controls, and see see Section S7.2.4 for the results), Interestingly, that melting temperature of

[13]See Section S5.4 for comparisons to systems that use toeholds, with design that look similar to our compute domains.

[14]It's important for toeholds to be able to bind in the case the displacement domain is correct, and be able to unbind in cases the displacement domain is incorrect [16, 110, 138]. We note that there is some flexibility in the choice of temperature [138].

[15]That intuition is complicated by the difficulties in matching a bulk fluorescence signal with a precise concentration of polymers in solution, a complication we leave to future experimental and modelling work.

a scaffolded polymer complex is around the temperature where we see 0/1 decisions happening during an anneal.

We conclude that the SDC system is making its 0/1 decision at temperatures closer to the melting temperature of an entire scaffolded polymer complex than to the much lower compute domain (or putative "toehold") melting temperature. This in turn suggest that the precise mechanism of TMSD is not the driver of those decisions, but the equilibrium distribution of polymers of those higher temperatures is playing a role (and whatever kinetic mechanisms are reasonable at such a hot temperature, but certainly some form of binding and replacement as prescribed in the abstract SDC tile and strand-level models in Figure 2 of the main text).

##### S3.6.3 Decisions are made too fast for TMSD to be the main driver

Our super-fast anneals show 0/1 decisions happening in under a minute, and typically in less than 30 seconds and above 60 °C (e.g. Figure 3f in the main text, or the collection of super-fast results shown in Section S7.3).

The rate for TMSD ranges over six orders of magnitude, capping out at $10^6$ $M^{-1}$ $s^{-1}$, which at our compute strand concentration of 1 $\mu$M gives 1 $s^{-1}$. Although TMSD can be fast, those fast rates are for a somewhat standard, and kinetically well-designed, three-way branch migration [31, 110], or other nicely designed systems [113]. Intuitively more complicated systems, such as ours, with remote toeholds should be slower, in particular SDC systems require melting off of long compute domains which need to happen at higher temperatures (which is fine, but a constraint), but more importantly initiation of displacement will require closing of a large loop while waiting for such a disassociation to happen.[16] Also, complicated circuits with several downstream reactions (which SDC has), take hours to reach a good completion level.[17] We have not attempted a careful kinetic characterisation of the obvious TMSD-style SDC pathways (that can be derived from Figure 2 in the main text), however, data (Figure S4) suggests a timescale of hours, perhaps up to half a day, for a cascade to copy information from an anchor strand at scaffold position A, all the way to position D. Such a slow reaction is perhaps not surprising, since the SDC compute domains, compute strands and scaffold are intentionally not optimised for TMSD and should have some kinetic issues, as mentioned above, when forced to operate under conditions (low temperature) that may require TMSD-style pathways. That data alone is perhaps not sufficient to completely rule out TMSD as the only kinetic mechanism behind our system, but taken together with the previous argument about decision making happening well-above the compute domain ('toehold') melting temperature, we hypothesise that it is highly unlikely that TMSD is the main driver of annealed SDC computations.

Theoretically, and experimentally, our system does not suffer from leak (slow changing of signal to an off-target/error – common in TMSD systems, which are out-of-equilibrium systems [39]). Further evidence that the SDC target/output really is the equilibrium.

Finally, it should be mentioned that, after our slow anneals (over the course of an hour or so), we hold at a fixed temperature and we see 'slow completion' to target/intended output (even holding up to 20 hours, see Figure S27). If we did see such a slow finishing-off of the signal at constant temperature, that could be interpreted as evidence that the system has not reached equilibrium during annealing, and required waiting on some kinetic mechanism to achieve the target. However, this is not the case in our data.

**Future work: kinetics.** Of course, like in all programmed nucleic acid systems (whether thermodynamically favoured or not) our implementations of the SDC could have unintended kinetic traps, due to DNA sequence impurities, or inadequate DNA sequence design, or our incomplete biophysical, energetic and mechanical understanding of DNA in the complex setting of DNA origami (helices, crossovers, adversarial scaffold sequence). These present interesting and novel challenges for this approach to DNA computing that we leave for future work. In particular, there remains much to understand, and likely design improvements to make [44], if we care about isothermal SDC computations, which could be useful for certain applications.

[16]We acknowledge Chris Thachuk for pointing some of this out, and Ahmed Shalaby for recent kinetic data backing it up.

[17]TMSD rates can be increased by increasing concentration [43], but leak rates also go up. Leakless systems are specifically designed to have large kinetic, or thermodynamic, barriers to leak, and thus have the potential to be quite fast at high concentrations without any noticeable leak [27, 32, 39].

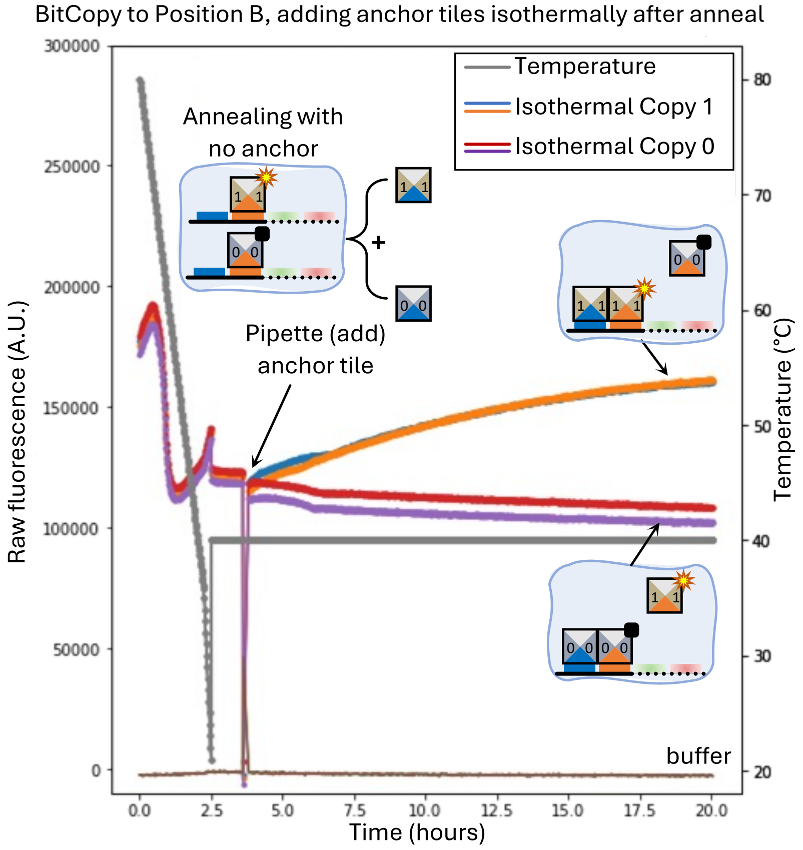

Figure S4: BitCopy to scaffold position $B$, adding input isothermally (at fixed temperature). Four samples are shown. At time zero, samples contained scaffold and two tiles for position $B$, but anchor (input) tiles were not present initially. All 4 samples were annealed down to 20 °C, then raised to 40 °C. After some time holding at 40 °C, at about 3.5 hours, an anchor/input strand was added to each sample (two samples received input `0`, and the other two received input `1`), isothermally at 40 °C. Holding at that temperature, slow completion is observed over the course of 20 hours. The down-spike ~3.5 hours is due to the qPCR drawer being opened at that time to pipette the anchor/input tile into each sample—an accidental data point was taken during the opening of the drawer and hence the sudden drop in signal at that time. Samples were pipetted by hand.

### S4 SDC tile model: programming and computational theory

#### S4.1 Simulating finite state machines (FSMs) with SDCs

Finite State Machines (FSMs) are common objects of study in computer science as they represent an important sub-class of computer programs and all the necessary background information can be found in standard textbooks [76, 139, 140]. FSMs have been a model of choice for molecular implementation [12, 33, 141–143] arguably because it is a *local* model in the sense that it does not rely on an unbounded randomly addressable memory, which also makes it a weaker model than Turing machines.

Nonetheless, we reintroduce the main concepts that we need to show that SDCs simulate FSMs.

In this work, we mainly implemented deterministic Finite State Machines with SDC with the exception of one nondeterministic Finite State Machine. However, for the sake of theory, we focus on nondeterministic FSMs as they are more general than deterministic FSMs, i.e. the set of nondeterministic FSMs includes deterministic FSMs.

**Definition S4.1** (Nondeterministic FSM)**.** A nondeterministic Finite State Machine (or Automaton) is given by:

1. an input alphabet $\mathcal{A}$

2. a set of states $Q$
3. a set of *initial* states $I$
4. a set of *final* states $F$
5. a transition function $\delta : Q \times \mathcal{A} \to \mathcal{P}(Q)$ where $\mathcal{P}(Q)$ is the set of subsets of $Q$

A *deterministic* FSM is a nondeterministic FSM such that $I$ and $\delta(q,a)$ contain only one state for all $(q,a) \in Q \times \mathcal{A}$.

**Definition S4.2** (Nondeterministic FSM computation)**.** Let $M = (\mathcal{A}, Q, I, F, \delta)$ be a nondeterministic FSM. Computation is performed on a finite word $w = w_0 \dots w_{n-1} \in \mathcal{A}^n$ of length $n \in \mathbb{N}$. The computation is performed in $n$ steps and at step, $0 \leq i < n$, machine $M$ is *in set of states* $S_i \subseteq \mathcal{P}(Q)$ reading character $w_i$ such that:

1. $S_0 = I$
2. $S_{i+1} = \{\delta(q, w_i) \mid q \in S_i\}$

$S_n$ is called the *terminal set of states* of the computation. Machine $M$ is said to *accept* word $w$ iff $S_n \cap F \neq \emptyset$. $M$ *rejects* word $w$ iff it does not accept it. The *Directed Acyclic Graph of $M$ reading $w$* is defined by nodes $\{(q,i)\}$ for $0 \leq i \leq n$, $q \in S_i$, and there is an edge between $(q,i)$ and $(q', i+1)$ iff $q' \in \delta(q, w_i)$ for $0 \leq i < n$.

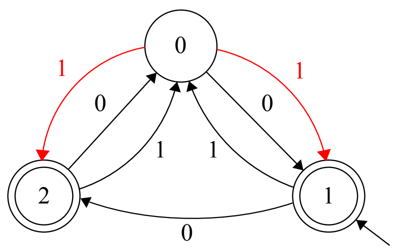

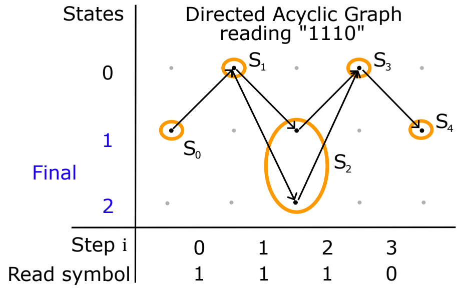

**Example S4.1** (3-STATE NONDETERMINISTIC FINITE AUTOMATON)**.** *Above is the 3-STATE NONDETERMINISTIC FINITE AUTOMATON implemented in our work, see S7.3.5. Following Definition S4.1, it is defined as:* $\mathcal{A} = \{0,1\}$, $Q = \{q_0, q_1, q_2\}$, $I = \{q_1\}$, $F = \{q_1, q_2\}$ *and* $\delta(q_0, 0) = \{q_1\}$, $\delta(q_0, 1) = \{q_1, q_2\}$, $\delta(q_1, 0) = \{q_2\}$, $\delta(q_1, 1) = \{q_0\}$, $\delta(q_2, 0) = \{q_0\}$, $\delta(q_2, 1) = \{q_0\}$. *The nondeterminism is highlighted in red in the above diagram: reading a* 1 *in state* $q_0$ *leads to* 2 *different states. For the word* 1110 *we have* $S_0 = I = \{q_1\}$, $S_1 = \{q_0\}$, $S_2 = \{q_1, q_2\}$, $S_3 = \{q_0\}$ *and* $S_4 = \{q_1\}$*; the word is accepted since* $F \subset S_4$*; its directed acyclic graph (Definition S4.2) is given above (right). The following words are also accepted:* 000, 11, 110 *with terminal set of states* $\{q_1\}$, $\{q_1, q_2\}$, $\{q_0, q_2\}$ *respectively. The following words are rejected:* 1, 111, 01 *with terminal set of states* $\{q_0\}$ *for all of them.*

**Remark S4.2** (Nondeterminism uses less states)**.** *A practical advantage of using nondeterminism in Finite State Machines is the ability, for some problems, to use less states compared to an equivalent deterministic machine. This is the case for 3-STATE NONDETERMINISTIC FINITE AUTOMATON where the smallest deterministic machine that accepts exactly the same set of words has 7 states compared to only 3 nondeterministically, see SI Section S7.3.5. This means that we would not have been able to implement this computation deterministically with the 3-bit SDC built in this work (indeed, 4 bits are needed to encode 8 states and 1 input bit).*

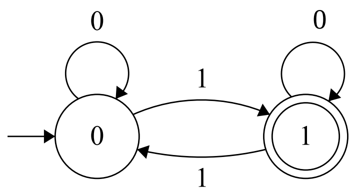

**Example S4.3** (Deterministic FSM: PARITY)**.** *Above is the PARITY 2-state deterministic Finiste State Machine, see S7.3.3. Following Definition S4.1, it is defined as:* $\mathcal{A} = \{0,1\}$,$Q = \{q_0, q_1\}$, $I = \{q_0\}$, $F = \{q_1\}$ *and* $\delta(q_0, 0) = \{q_0\}$, $\delta(q_0, 1) = \{q_1\}$, $\delta(q_1, 0) = \{q_1\}$, $\delta(q_1, 1) = \{q_0\}$. *This machine only accepts words that have an odd number of 1s, such as* 001, 1011 *or* 10111010.

**Remark S4.4** (Finite State *Transducers*). *Machines* MULTIPLYBY3 *and* DIVBY2 *implemented in this work (see S7.3) are fundamentally similar to FSMs as per Definition S4.1 with the exception that there is no notion of final state but that instead, output characters are produced (from the same alphabet $\mathcal{A}$ as for input characters) at each computation step.*

The following theorem is the main result of this section and gives us a simple method to program a wide range of SDCs. The theorem says that any FSM, and input word, can be translated into an SDC system. More precisely, each terminal configuration of the SDC corresponds to a path in the directed acyclic graph of the FSM $M$ reading input $w$ (Definition S4.2). It should be noted that some of the problems we solve with SDCs in the main paper have longer input length than scaffold length–for example, ADDITION of two 4-bit numbers (8 bits of input), and PARITY applied to 8 input bits. In those cases we use a simple trick of encoding 2 bits of input into a single base-4 symbol, and then we think of that base-4 string as the 'input', and we define an FSM with that base-4 input. See for example the ADDITION FSM in Figure 3a, which reads two bits of input at a time, i.e. base 4.

**Theorem S4.5** (SDCs simulate FSMs). *Let $M = (\mathcal{A}, Q, I, F, \delta)$ be a given nondeterministic FSM. Let $w = w_0 \dots w_{n-1} \in \mathcal{A}^n$ be a length $n$ input word. We define $\mathcal{S}$, a SDC with $n$ scaffold positions that will simulate the computation of $M$ on $w$. Let $c(q, i)$ be a tile side's colour uniquely identified by $q \in Q$ and $0 \leq i < n$. The tile set $T_i$ at position $0 < i < n$ is given by tiles with left colour $c(q, i)$ and right colour $c(q', i + 1)$ for all $q \in Q$ and $q' \in \delta(q, w_i)$. The set $T_0$ is defined similarly but where $q$ is from the initial states $I$ instead of $Q$. Then:*

1. *$\mathcal{S}$ has as many terminal configurations as paths from nodes $(q, 0)$ to nodes $(q', n)$ with $q \in S_0$ and $q' \in S_n$ in the directed acyclic graph of $M$ reading $w$ (Definition S4.2)*
2. *If $(q_0, 0), (q_1, 1), \dots, (q_n, n)$ is a path in the directed acyclic graph of $M$ reading $w$ then the configuration where, for each $i$, position $0 \leq i < n$ of the scaffold has tile with left colour $c(q_i, i)$ and right colour $c(q_{i+1}, i + 1)$ is a terminal configuration of $\mathcal{S}$*

*Proof.* There is a 1-1 correspondence between terminal configurations of $\mathcal{S}$ and paths from nodes of the form $(q, 0)$ to nodes of the form $(q', n)$ in the in the directed acyclic graph of $M$ reading $w$, with $n$ the length of $w$ (Definition S4.2). In short, tiles of $T_i$ that are used in terminal configurations of $\mathcal{S}$ exactly correspond to edges in the graph. In detail:

- A path $(q_0, 0), \dots, (q_n, n)$ in the graph corresponds to the SDC configuration where the tile at position $0 \leq i < n$ has left colour $c(q_i, i)$ and right colour $c(q_{i+1}, i+1)$, i.e. the edge from $q_i$ to $q_{i+1}$ corresponds to choosing the appropriate tile in $T_i$. The configuration is terminal since the binding is maximal because each tile has matching colours on shared edges; hence no SDC step can increase the number of matching colours (Definition S3.2)
- A terminal configuration of $\mathcal{S}$ corresponds to a path $(q_0, 0), \dots, (q_n, n)$ in the graph. Indeed, (a) all positions of a terminal configuration have a tile (no $T_i$ is empty and if there is no tile at a position a step can always be performed to add any random one since that won't decrease the number of matching colours) and (b) there can be no mismatch. Indeed, take $i$ the position of the first mismatch, the tiles that come before the first mismatch must match and the colours on their side are $c(q_0, 0), c(q_1, 1), \dots, c(q_i, i)$ with $q_0 \in I$ and $q_{j+1} \in \delta(q_j, w_j)$ $j \leq i$, meaning that they correspond to successive edges in the graph and if there was a mismatch at position $i$, we can replace the mismatch with the tile that corresponds to the next edge in the graph (always defined because $\delta$ is total) and this step is valid because it will either increase or conserve the number of matches. Repeating this operation at each mismatch shows that the terminal configuration has no mismatch and corresponds to a path of the form $(q_0, 0), \dots, (q_n, n)$ in the graph.

From this we get 1. and 2. □

**Remark S4.6** (A note on transducers). *Although Theorem S4.5 was stated for FSMs we note that it is easily generalised to Finite state transducers: with the minor change that for transducers the output is reported not by a final state, but at each step of the computation (examples in this work include* ADDITION*,* DIVBY2*,* MULTIPLYBY3*, and* RULE110*).*

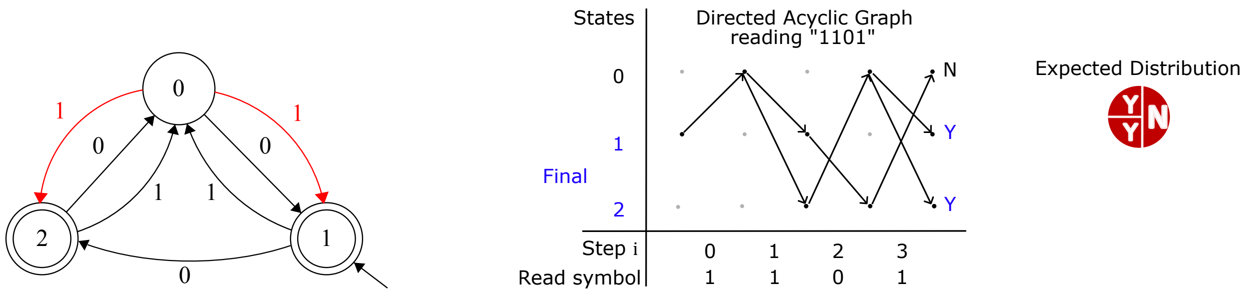

**Remark S4.7** (Concentration of terminal configurations)**.** *As seen in Theorem S4.5, when simulating a FSM $M$ reading $w$, a SDC will end up in as many terminal configurations as there are paths in the directed acyclic graph of $M$ reading $w$. For instance, see the above directed acyclic graph: the nondeterministic FSM implemented in this paper (Example S4.1) gives rise to 3 distinct paths when reading the word "1101" meaning that there will be 3 terminal configurations of the SDC. Two of them reach a final state (labelled* ***Y*** *like "Yes") and one of them is rejected (labelled* ***N*** *like "No"). It is natural to ask: for a supply of $K >> t$ scaffolds with $K, t \in \mathbb{N}$ and $t$ the number of terminal configurations, what is the proportion of each terminal configuration after annealing? (In this work with scaffold at 100 nM in 35$\mu$L we have $K \simeq 2000$ billion $>> 3$.) The answer is read on the directed acyclic graph: each path (i.e. terminal configuration) starts with proportion $1/n_0$ with $n_0 = |I|$ the number of initial states and this proportion is divided by the out-degree of each successive node of the graph. For instance, here this reasoning gives that $1/2$ of the population will be for the terminal configuration reporting* ***N*** *and $1/2$ of the population for two different ways of reporting* ***Y****, with each* ***Y*** *being $1/4$ proportion. This expected distribution over* ***N*** *and* ***Y*** *is illustrated to the right of the directed acyclic graph in red. The same style is used in, for example, Figure 4g of the main text to allow the reader to easily verify the experimental result.*

##### S4.1.1 Computational sinks

In the context of Finite State Machines (SI Section S4.1), a *computational sink* happens when, for a given read symbol state $a$, reading $a$ in any state brings to a single state $q'$. Mathematically: $\exists a \in \mathcal{A}$, $\exists q' \in Q$, $\forall q \in Q$, $\delta(q, a) = \{q'\}$.

Given the way SDCs simulate FSMs (Theorem S4.5), a computational sink in a FSM means that there is a scaffold position of the SDC at which all competing tiles have the same output computation domain. This has the effect of *resetting* the SDC at that scaffold position: computation happening after this scaffold position becomes independent of computation happening before it.

In practice, for instance, this gives the surprising and unintuitive result that performing BitCopy is harder than performing Addition because there are no sinks in BitCopy while there can be some in Addition. In accordance with this observation, we experimentally measure better performance for computations with sinks than without, see SI Section S7.2.6.

#### S4.2 Simulating FSMs with separate tiles for program and input

The standard encoding of FSMs into tiles, used in the main text (Figures 2–5) and in this document, is more compressed, allowing input and program bits to be encoded in a single compute domain. Here, in Figure S5 we give a second method for SDCs to simulate FSMs, where input bits and program bits are encoded in two separate tile sets. One advantage of Figure S5 over our standard encoding is that it would allow for renewing a program by resetting an input, but without needing to re-supply program bits.

#### S4.3 Other modes of SDC computing, beyond FSMs

In the language of computer science, there is a sense in which all problems solved by SDCs in this paper, and all problems solved on modern digital silicon-electronic computers are *regular*. This is simply because all such devices have finite memory, thus computers can only solve problems with finite-length inputs, and they run for only a finite time. However, theoretical computer science, and algorithms and computational complexity in particular, considers this an uninteresting observation and prefers to characterise the difficulty

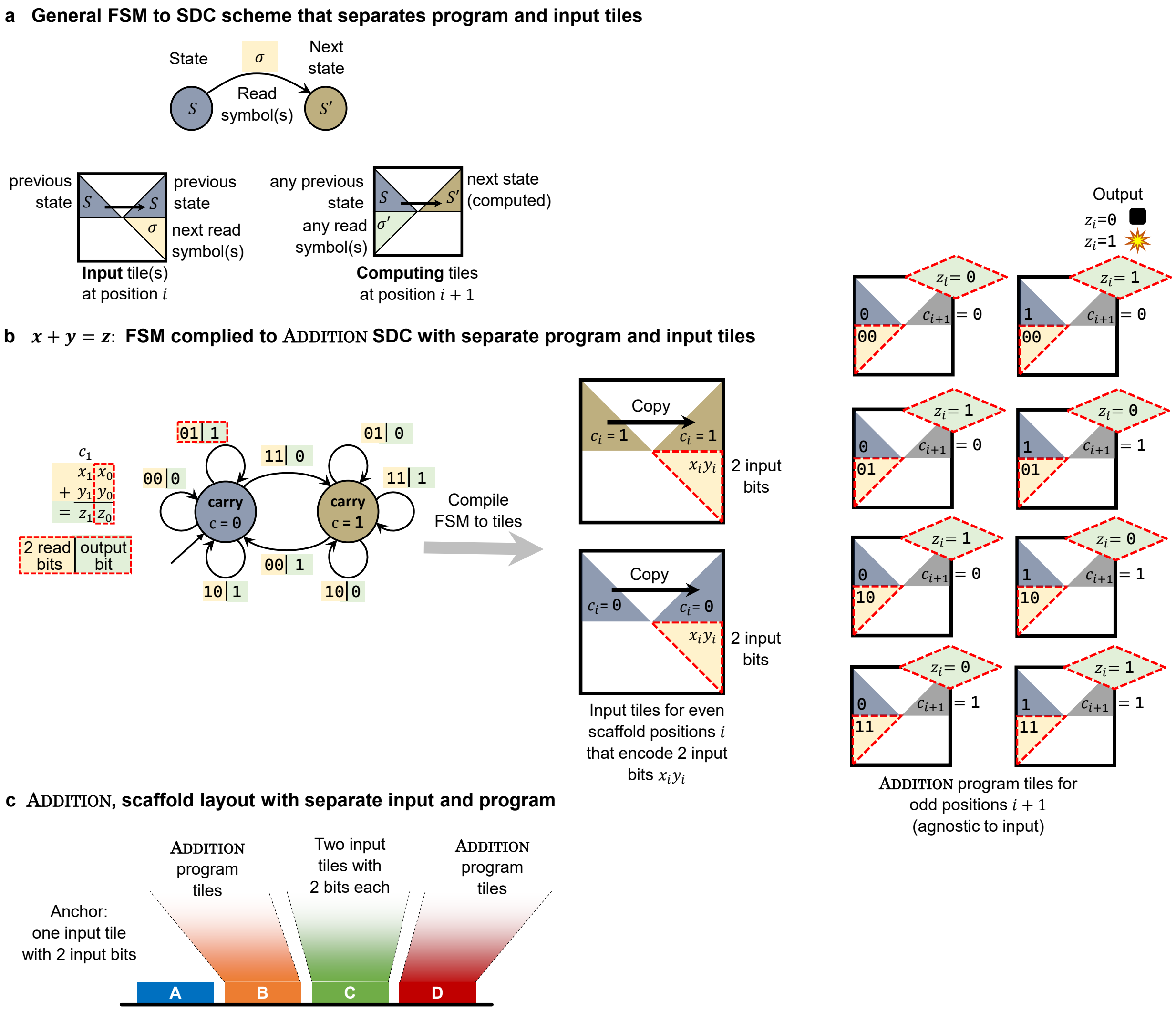

Figure S5: Scheme for simulating FSMs using SDCs that separately encode input (at odd scaffold positions) and program on one tile set (at even scaffold positions). a. Mapping from FSM transition to tile schematic. b. Example: the ADDITION FSM, that adds two $n$-bit numbers, is encoded as $n$ tile sets. Two tiles that bind to even scaffold positions ($i = \{0, 2, 4, \ldots\}$) encode a pair of input bits $x_i, y_i$, and copy-through a carry bit from left-to-right. Eight tiles that bind to odd scaffold positions ($i = \{1, 2, 3, \ldots\}$) encode the addition logic, and may (or may not) give output bits. c. Schematic of layout on a length 4 scaffold.

of problems as the input length tends to infinity, as that seems to better capture our intuition of 'problem complexity' as well as actual resource usage on computers.

The following SDC systems, from Figures 2–4 solve problems solvable by FSMs or finite state transducers: ADDITION, BITCOPY, PARITY, MULTIPLYBY3, 3-STATE NONDETERMINISTIC FINITE AUTOMATON, DIVBY2, COUNTER. The proof being trivial: in each case we defined a simple FSM, and converted that into the relevant SDC system using Theorem S4.5 (or similar reasoning). We note that 3-STATE NONDETERMINISTIC FINITE AUTOMATON solves a problem solvable by a non-deterministic FSM, which noted in Remark S4.2 solve the same problems as deterministic FSMs, just more efficiently in terms of number of states.

In Figure 4 various other SDC systems are given that were not converted directly from FSMs: RULE110, GRAPHREACH, BALANCEDBRACKETS. Indeed the problems solved by these machines, when (canonically) scaled up to arbitrary input size, are not regular, meaning such problems provably do not have an associated FSM [76, 139, 140]. We next discuss each of them.

**BalancedBrackets** The arbitrary-length generalisation of the problem solved by BalancedBrackets is not a regular language, but is context-free [76]. The canonical model of computation for languages of matching brackets is a pushdown automaton, which runs by pushing opening brackets to a stack and popping them off as the correct closing brackets are met. One can instead use a few counters. For SDC systems this means that for long bracketed expressions, with deeply nested brackets, beside having a long scaffold, many compute domain types would be needed to count up the opening brackets and then count down to match them off.

**GraphReach** is NL-complete [76], meaning that it is solved by nondeterministic Turing Machines that use logarithmic space, and is as hard as any problem in that class (hence it is not regular, as regular sets are decided by finite state machines (a restriction of Turing Machines) with only constant memory. For large graphs, with many nodes at each layer, an instance of a GraphReach SDC would require many compute domains.

**Rule110** The generalisation of the rule 110 cellular automaton to arbitrary input lengths is computationally universal [144, 145] in the following sense: given a Turing machine and its input, there is an instance of Rule110 that simulates it. Our Rule110 SDC (Figure 4g) simulates small (length 3), instances of rule 110 for finite time (only 4 steps). This construction puts almost all of the complexity of the simulation into the compute domains, making it essentially unscalable: indeed we claim that **it would be utter nonsense to claim it Turing universal**. Specifically: First, if the Turing machine doesn't halt this means that the rule 110 simulation uses wider and wider configurations, meaning in turn there is no finite length compute domain that can handle those configurations. Even if we restrict to halting Turing machines, on configurations of some size $s$, we would need compute domains that encode something like $O(s)$ bits [145]. Perhaps even worse, we would need exponentially many (in $s$) of those huge domains (and thus that many tiles) to be able to compute on any such configuration. Nevertheless, we felt it was fun to show the (small!) Rule110 SDC system in Figure 4g.

**Yet other styles of SDC computation** In a deterministic SDC we have a single target configuration. Even more restrictive in this work we have a single tile at position A, and then the choice of one out of two or more tiles at each other position. This leads to left-to-right computation. One could also compute right-to-left (by having only one tile at the final/rightmost scaffold position), or one could have the best of both worlds by having systems that compute inside-to-out (single deterministic position in the centre).

We leave it to future work to explore other models of SDC computing.

#### S4.4 Our tile set encoding does not cheat

Our simulation of finite state machines (FSMs) with the SDC, Figure 3a, has the curious property that both FSM program bits and input bits are shared among tiles: i.e. each compute domain has a few bits of the program and a few bits of input. A reasonable concern is, since program and input appear mixed together, could the programmer cheat somehow? Specifically, given an FSM and its input, could the programmer run the computation themselves on pen and paper, and then merely choose a tile set that essentially does nothing of interest, and outputs the correct answer? Actually, yes. In such a scenario, the programmer did the hard work, and the DNA in the test tube is merely giving a hardcoded answer. The purpose of this subsection is to show that our program-input encoding does not cheat in this way.

Some of our SDC programs and inputs are encoded using the technique in the proof of Theorem S4.5, and all 10 SDC programs in this paper use similar ideas. Thus we focus on the terminology in the proof of Theorem S4.5: We fix an FSM (or finite state transducer) and a length $n$ input word $w = w_0 \dots w_{n-1} \in \mathcal{A}^n$. From that we get a compiled tile set $T$. In particular at some position $i$ of the scaffold, the proof gives a compiled set of tiles $T_i$ that compete for scaffold position $i$. Now consider running the same procedure but on a new, different, input $w'$ that has the same values as $w$ at indexes $i, i+1$ in $w$ but is different at some other indices, more precisely we have: $w \neq w'$ (different inputs), but $|w| = |w'|$ (same length), and $w_i w_{i+1} = w'_i w'_{i+1}$ (same values near $i$). From $w'$, applying the theorem, we get a new tile set $T'$. However, at scaffold position $i$, the tile sets are identical, i.e. $T_i = T'_i$. Hence, the programmer has no opportunity to cheat in the way described, since if the tiles near the output scaffold position $n$ somehow encode "special" non-local information about bits throughout $w$, an adversary can come along and switch out $w$ for $w'$, which does not change the tiles near the output region, but *crucially will change the output itself*, thus "fooling" the programmer who thought they could cheat. It is possible to formalise the claim into a theorem with a full proof, but we omit the details.

Indeed a stronger statement can be formalised and proven that leverages ideas from uniform Boolean circuit complexity theory [76, 146]. We give a brief summary of the argument: Using the formalism of Boolean circuits, one can show that our encoding functions are in the special, weak, complexity class $\mathrm{FAC}^0$, which gives a formal sense to the intuitive meaning that these encoding functions are 'simple'.[18] Since we know that SDCs solve the parity problem ("is the number of `1`s in the input odd?"), and since it is known that parity is not in $\mathrm{AC}^0$ (and also not in $\mathrm{FAC}^0$), and since our encoding is in $\mathrm{FAC}^0$, we get that the SDC system is itself solving the parity problem, and the encoding function is provably not sufficiently expressive to solve parity. Thus 'cheating' is mathematically impossible with the kinds of encoding functions we use when solving parity with an SDC.

**An alternative encoding** Section S4.2 gives an encoding for SDC to simulate FSMs, but where input and program bits are not mixed. That shows that mixing the bits on the same compute domain is not necessary for FSM simulation.

[18] $\mathrm{FAC}^0$ is the class of functions that map bit string to bit strings, and are computable by polynomial sized, constant depth, Boolean circuits. To establish this one leverages a suitably simple encoding of tiles as short bitstrings.

### S5 SDC strand model: system design details

This section gives design details in the strand model that are needed to understand our experiments. It builds upon the strand-level SDC model given in Section S3.4, by fixing SDC model parameters $N$ and $\ell$, adding strands for fluorescence reporting, cover strands to block unintended interactions, and domains/strands for renewable programs.

In the experimental system reported in the main paper, we set $N = 4, \ell = 3$ (meaning 4 scaffold positions for computing and $\leq 3$ bits per compute domain). As shown in Figure 2h and Figure 3d, for $N = 4$ the strand-level design requires $N + 1 = 5$ unique binding domains ($A$, $B$, $C$, $D$, and $E$). Each scaffold position domain is 24 bases long. In the programs we implemented, position $A$ always binds the *anchor* strand (Figure 2a and Figure 3c). Domains $B, C, D$ were involved in computation and reporting of programs of scaffold length $\leq 3$, and domain $E$ is for reporting programs of scaffold length 4.

#### S5.1 Compute strand design: avoiding unwanted hairpins

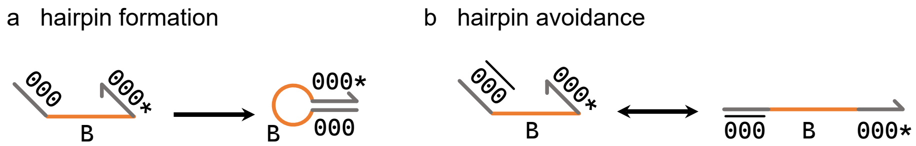

Figure S6: a. An undesired hairpin forms when compute domains are reverse complements, i.e. $\texttt{000}^*$ is the reverse complement of $\texttt{000}$. b. To avoid unwanted hairpins we make two versions of each logical domain: Compute strand $\overline{\texttt{000}}B\texttt{000}^*$ remains single-stranded due to the two compute domains $\overline{\texttt{000}}$ and $\texttt{000}$ being distinct. i.e. $\texttt{000}^*$ is not the reverse complement of $\overline{\texttt{000}}$.

As shown in Figure 2d, and described in the main text, a compute strand is a 3-domain strand that represents the left, bottom and right domains of a tile. In this work, we chose to have an $\ell = 3$-bit SDC, meaning that each compute domain represents 3 bits to encode programs and data. That gives 8 *logical* compute domains, numbered in binary: `000`, `001`, `010`, `011`, `100`, `101`, `110` and `111`. The programmer is then free to choose a compute strand that has any pair of compute domains, for example a strand at scaffold position $B$ with `000` on its left and `111` on its right which we would write as $\texttt{000}B\texttt{111}^*$, (giving $8 \times 8 = 64$ compute strands per scaffold position; and where $^*$ denotes reverse complement of DNA). This flexibility of choice enables reprogramming of the SDC system by simply choosing a set of tiles/strands. However, unlike square tiles, single-stranded strands are considered to be floppy which creates an issue with this design: if the programmer were to choose a strand that binds to scaffold position $B$ with two logically identical compute domains, e.g. $\texttt{000}B\texttt{000}^*$, that strand would form a hairpin, Figure S6(a). Hence as described in Figure S6(b) we designed two versions of each logical compute domain, $\overline{\texttt{000}}$ and `000` – the former appears between odd-even scaffold positions, and the latter at even-odd scaffold positions, thus avoiding unwanted hairpins within compute strands.

#### S5.2 Reporter design: Sequence independent reporting mechanism (SIRM)

##### S5.2.1 Reporting binary outputs: 0/1

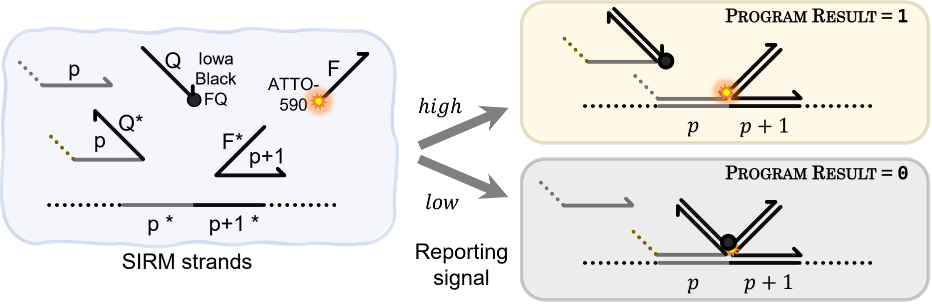

Figure S7: Sequence independent reporting mechanism (SIRM). Left: strands for reporting at some position $p$. Top right: reporting a high signal when the output of the computation is the strand at position $p$ with no quenching domain. The strand with the quencher domain is not bound. Bottom right: reporting low signal when the output of the computation is the strand at position $p$ with the quenching domain.

The reporting mechanism used to read results of the 1D SDC is depicted in the paper main text Figure 2h, and here in Figure S7. This design is used for reporting results of ADDITION in Figure 3e,f, as well as BITCOPY, PARITY, MULTIPLYBY3, and 3-STATE NONDETERMINISTIC FINITE AUTOMATON, in Figure 4a–d,f. In those programs, we report output bit-0 with a low signal and output bit-1 as a high signal. The design includes an 'Iowa Black FQ' quencher (from supplier IDT) attached to 20-base domain $Q$ and 'ATTO-590' fluorophore attached to 20-base domain $F$.

To report, for example at position $p = B$ (see Figure S7), the programmer does the following: (a) Include the strand $F^*C$ in the system (here $p + 1 = C$), (b) the compute strands reporting bit-0 at position $B$ should be of the form $cBQ^*$ (where $c$ is some compute domain), **attachment of these strands causes quenching**, (c) the compute strands reporting bit-1 at position $B$ should be of the form $cB$, i.e. they have no right-hand-side ($3'$ end) domain at all, **attachment of these strands implies no quenching**.

In a typical SDC program several compute strands (e.g. 2 for BITCOPY) compete at the reporting position, some with domain $Q^*$ and some without, depending on the output those strands encode. In this design, a single identical fluorophore and quencher strand allow for reporting at any position, a feature which allows for reduced reporting cost and increased experimental consistency. Although not pursued here, we note that a multiplexed design with a distinct fluorophore per position would allow for reading 4 positions at once albeit sacrificing signal quality due to spectral overlap.

##### S5.2.2 Reporting ternary outputs: 0/1/2

Here, we discuss our novel ternary (base 3) reporting mechanism, for reporting one of three trits denoted 0, 1 or 2. The design is illustrated in Figure S8(a). Trit-0 and trit-1 are reported using precisely the mechanism described for bit-0 and bit-1 in the previous section (binding, or not, to a quencher-labelled strand $Q$ by presence/absence of the domain $Q^*$). Trit-2 is reported using a domain that intentionally, weakly binds to the quencher-labelled strand. To give an example: for reporting trit-2 at position $B$ there is a strand $Bc^*$ where the domain $c^*$ binds weakly to the quencher-labeled strand $Q$ (i.e. $c^*$ and Q are not full reverse complements but do share some partial reverse complementarity).

So, what sequence did we choose for $T^*$? Somewhat serendipitously, we discovered that one of our compute domains happened to have sufficient complementarity with $Q$ to give the required signal for trit-2. In particular we used compute domain $\overline{\texttt{CD5}}^* = \overline{\texttt{101}}^*$, with sequence GGACTGGTAGTG, for the orange domain called T* in Figure S8 As shown in Figure S8(b)(bottom), T* has a significant secondary structure (7 bases of complementarity) with the quencher-labelled strand $Q$, with MFE of -12.10 kcal/mol at 30 °C, albeit less than between the perfect complements $Q$ and $Q^*$. This means that the quencher-labelled strand $Q$ will spuriously bind to compute domain $\overline{\texttt{CD5}}^*$ (i.e. T* in the figure above), but only at temperatures lower than $Q^*$-$Q$ binding. Since our experiments follow a temperature annealing protocol from high to low

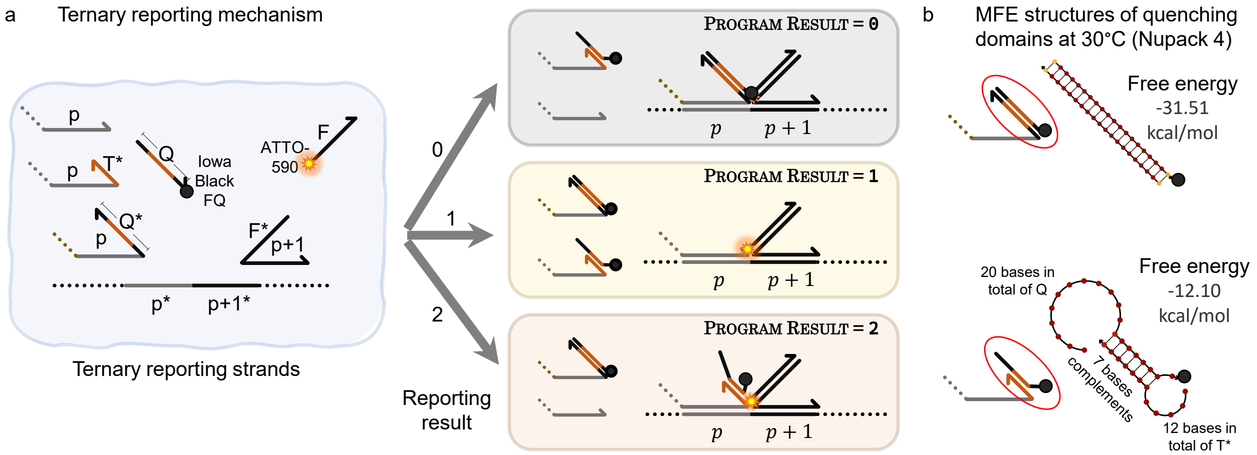

Figure S8: a. Mechanism of reporting ternary 0/1/2 used in Figure 4e in the main text. b. MFE structures if $Q$ domain exists in the reported result at 30 °C (NUPACK4 [57]). Top shows when reporting trit 0 and bottom shows when reporting trit 2 with 12 bases compute domain 5 (101) and 20 bases quencher strand.

temperature, during an experiment, we see quenching that reports trit-2 occurring later (lower temperature) than quenching that reports trit-0. More precisely, we tend to see $Q^*$-$Q$ quenching in the 65–75 °C range (green curves in Figure 4e in the main text) whereas $T^*$-$Q$ occurs in the 30–50 °C range: purple curves in main text Figure 4e show a control for the mechanism where there is no competition between strands at any scaffold position in the control—i.e. the control shown in purple has exactly one scaffold-binding strand per scaffold position.

##### S5.2.3 Reporting outputs for high competitive complexity programs

Previously, we gave designs for reporting 2 (binary), or 3 (ternary), different output signals. Here, we give another method for reporting any number of output signals, with the downside that it requires more samples to be run. Specifically, in order to assess high competitive complexity SDC programs, we separately report each of $k \in \{2, 3, 4, \ldots\}$ potential outputs by having the correct output cause quenching, but also running $k - 1$ parallel experiments where we test each of the other (wrong!) outputs to asses if they quench: if the correct output quenches and the other $k - 1$ do not, we have a successful experiment.

We use this reporting method for the SDC programs Counter, Rule110, GraphReach, Balanced-Brackets (main text Figure 4f,g,h, and e). In those SDC programs, there are $> 2$ tiles at the last computing position $D$ that we wish to report. We illustrate with the Counter program, which takes one of 8 inputs $x \in \{0, 1, 2, \ldots, 7\}$ and reports the output $x + 4 \in \{0, 1, 2, \ldots, 7\}$. Let's suppose we have input $x = 0$ (as shown in Figure 4f in the main text). In a "positive sample", at the reporting position $D$, we choose the correct answer (compute domain 4, or CD4, in this case) to be the quenching domain $Q^*$ while the other 7 strands competing at position $D$ do not. We then run 7 other samples, sample number $i \in \{0, 1, 2, 5, 6, 7\}$ having the quenching domain $Q^*$ at position $D$—this should report the fraction of incorrect polymers. As expected, all 7 show a high signal in the relevant plot in Figure 4f, suggesting (at the very least) that we are seeing a very low fraction of errors. A concrete Counter mix, showcasing this reporting technique, is shown in SI Section S7.8, SI Figure S35.

One might ask: Why did we flip our reporting scheme from 'correct answer = no quenching', to 'correct answer = quench'? If we had reported the correct output with a high signal, then only the correct output would have the no-quenching domain, and all other $\leq 7$ reporting domains would have an attached $Q$ domain. But then, this might require a high excess of quencher-labeled strands, a large waste of a labeled strand (labelled strands take longer for shipping and are more expensive than unlabelled strands).

Strand excesses (concentrations) are discussed in detail in Section S7.5.

#### S5.3 Compute controls

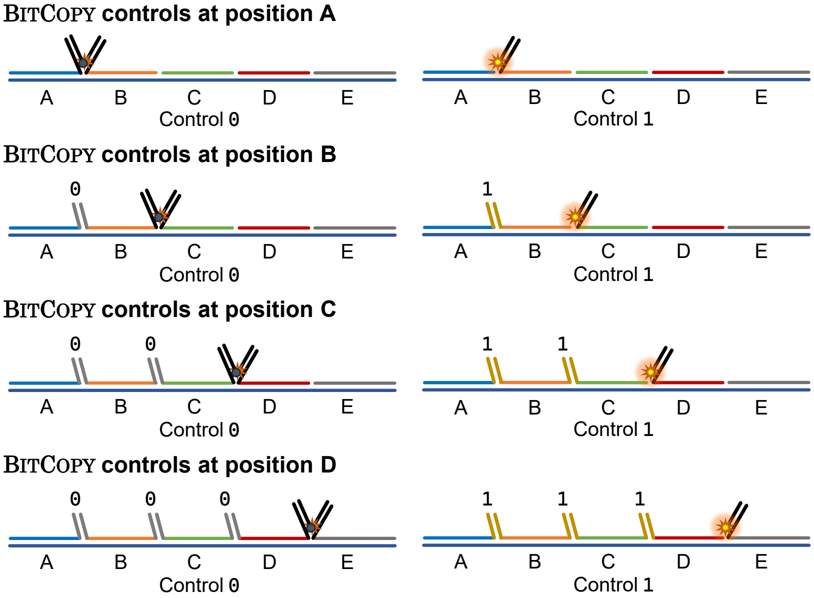

Figure S9: Strand diagram illustrating controls for bit-0 and bit-1 at each of position $A$, $B$, $C$, and $D$. A *control* is an SDC sample where there is exactly one strand per scaffold position, i.e. there is no competition for a scaffold position. For each compute domain there are 16 possibilities, for the BitCopy data, reported in Figure 4 in the main text, bit-0 is encoded by compute domain CD0 and bit-1 is encoded by compute domain CD2. Unused positions are covered by a single-domain scaffold complement, or *cover*. Each compute domain has its own set of control experiments.

Throughout this paper, we use the term 'control' to mean 'compute control', more precisely, a control is an experiment that (a) does not compute anything, and (b) is designed to have at most one compute strand per scaffold position. Strand diagrams for (computing) controls are given in Figure S9. The idea is to have a scaffolded structure assemble, without any competition between strands, giving a clear bit-0 (quenched/low), or bit-1 (unquenched/high), signal. Completion levels, for computing samples, are measured against these controls.

#### S5.4 Comparison of SDC with other toehold-based designs

Although we intentionally do not prescribe a toehold-mediated strand displacement (TMSD) pathway for the SDC replace primitive, we note that the SDC configurations, with exposed (mismatching) compute domains, have superficial similarities with a variety of toehold-based designs. Indeed we leave as future work to explore, characterise and optimise isothermal, TMSD style, computation with the SDC that would treat compute domains as toeholds. In anticipation, it is worth giving some context by discussing related work on various toehold designs.

At first glance, perhaps the closest in spirit is the associative toehold design of Chen [114] where the toehold is not on the same strand as the migrating domain. The invading strand attaches to the toehold then displaces a strand that is not attached to the strand with the toehold. For contrast, in more common and standard TMSD [46, 110] design the invading strand displaces the strand that is binding to the strand with the toehold. Another strand displacement design by Cabello-Garcia et al [113] has some similarities with the SDC design but they use a second toehold called a *handhold* beside the toehold to accelerate the displacement rate. Other designs of strand displacement that are different from the SDC are the allosteric toehold design [116] and the remote toehold [115]. The allosteric toehold design controls strand displacement by making it on-demand: meaning that the toehold is sequestered as double-stranded, and displacement can only be activated if an auxiliary strand exists in solution to displace the toehold and make it single-stranded. The remote toehold design gives extra control on hybridization kinetics by placing a spacer domain between the toehold and the migrating domain.

#### S5.5 Strand count

A total of 1,539 strands were designed and used in the reported experiments, these strands were as follows. Strand design names/details are given earlier in this section, except where stated below. See Section S10 for DNA sequences.

**395 strands** were used for the **scaff-**120 experiments, listed as follows:

- With $N = 4$ scaffold positions, and $\ell = 3$ bits per compute domain, tiles and scaffold are encoded as 201 strands $=1 + (2^\ell)^2 \times (N-1) + 2^\ell$:
  - 1 scaffold strand that contained five scaffold positions named $A$–$E$ (where $A$–$D$ were used for computation, and $B$–$E$ for reporting and the $5'$ to $3'$ position domain order is $E, D, C, B, A$).
  - At each of positions $B$, $C$, and $D$ there were 64 compute strands. $(+3 \times 64)$
  - At position $A$ there were 8 compute strands: each had a 3-bit compute domain on the right ($5'$ end), and since there was no need for domains on the left ($3'$ end) we did not include those compute domains. (+8)
- 64 strands for reporting (see Section S5.2 for details):
  - 5 strands each consisting of a single scaffold position-complement domain, i.e $A^*$, $B^*$, $C^*$, $D^*$, and $E^*$; for use in computing at any position ($A$–$D$), reporting at any position ($B$–$E$), and with any depth (1–4, see Section S7.6). (+5)
  - at each of positions $B$, $C$, and $D$, there were 8 strands that had a quencher domain (3 bits on left/$3'$ domain, and the quencher domain on the right/$5'$), as well as 8 strands with no domain on the right. $(+3 \times 2 \times 8)$
  - 4 strands at each position that report a quenching signal $AQ^*$, $BQ^*$, $CQ^*$, and $DQ^*$ with a quencher domain on the right, and no computing domain on the left. (+4)
  - 4 strands ATTO$^*B$, ATTO$^*C$, ATTO$^*D$, and ATTO$^*E$ that have domain $F^*$ that binds to the fluorophore-labelled strand $F$. (+4)
  - one quencher-labelled strand $Q$ ('Iowa Black® FQ') and one fluorophore-labelled strand ('ATTO 590') $F$. (+2).
  - one strand that is complementary to the quencher-labelled strand $Q$ and the fluorophore-labelled strand $F$ where the 'Iowa Black® FQ' and the 'ATTO 590' are perfectly beside each other for maximum quenching. (+1).
- 130 strands for renewable SDC programs (see Section S8.2.1 for details):
  - blocker strands of the anchor tile at position $A$: 8 blocker strands that are perfect complements to the 8 compute strands that bind at $A$. 1 blocker strand that is perfect complements to the strand $AQ^*$. 1 blocker strand that is a perfect complement to the strand with no compute domain at position $A$ (+10)
  - 64 blockers at position $D$ that are perfect complements to the 64 compute strands that bind at position $D$. (+64)
  - 16 blockers for output strands at $D$ (8 with quenching domain on the right and 8 with no domain on the right). (+16)
  - 20 strands with two different left domain at position $A$ with all possible 10 right domains (8 compute domains, one quencher domain, and no right domain) at its right. $(+2 \times 10)$
  - 20 blocker strands that are perfect complements to the strands with left domains at position $A$. (+20)

This gives:

$$395 = 1 + (3 \times 64) + 8 + 5 + (3 \times 2 \times 8) + 4 + 4 + 2 + 1 + 10 + 64 + 16 + (+2 \times 10) + 20$$

For the scale-up programs, a number of 1,144 strands were ordered and used. A more concise breakdown is as follows:

- Both scaff-288 and scaff-624 used M13 as the scaffold in experiments, but were listed as separate entries in the sequence list to indicate the specific M13 fragment used in each scale-up length. (+2)
- Scaled-up scaff-288BitCopy used 215 strands in total, including anchors, computing strands, reporting strands, and strands with mismatches. (+215)
- 25-bit IsoenergeticBitCopy used 387strands including anchors, computing strands, reporting strands, and strands with mismatches. (+387)

- 25-bit Addition used 540 strands including anchors, computing strands, and reporting strands. (+540)

This gives:

$$1,144 = 2 + 215 + 387 + 540$$

Both length-4 and scaled-up experiments number of strands give a total of:

$$1,539 = 395 + 1,144$$

### S6 DNA sequence design for SDC

In this section, we describe how DNA sequences were designed to implement the strand-level model. See the main text (Data and code availability), for the DNA sequence design code. All DNA interactions computed in this section used temperature parameter 53 °C and salt parameters [Na] = 1 M and $[\mathrm{Mg}^{++}] = 0$.

#### S6.1 Scaffold sequence for $N = 4$ SDC (scaff-120)

The 120-base scaffold (called scaff-120) consists of five contiguous 24-base domains called positions: $A$, $B$, $C$, $D$ and $E$. Computations ending at position $A$, $B$, $C$, and $D$ are, respectively, reported at positions $B$, $C$, $D$, and $E$.

To create scaff-120, we joined two unrelated subsequences from M13 (type p7249 using terminology from the company `tilibit nanosystems`[19]): one of length 96 and the other of length 24. Using M13 was an important design choice as we desired the system to be robust to biologically-sourced sequences, i.e. sequences that are not designed *de novo* by us for future applicability.

The use of two subsequences of M13 was more accidental than essential. First, we chose the sequence for four domains to be somewhat isoenergetic, specifically: we chose the length-96 subsequence of M13 that minimises the standard deviation of reverse-complement binding strength amongst four contiguous domains (using the typical straightforward summation of stack-loop energies in the nearest-neighbour model with parameters taken from SantaLucia and Hicks [60, 132]). Then, the use of a second (length-24) subsequence of M13 for a fifth domain was later added on after we realised the need for an extra domain for reporting purposes—we decided to use an M13-extracted sequence from earlier prototypes [20].

The scaffold sequence is given in SI Section S10.1.1 and was sourced from IDT as a PAGE-purified ultramer.

#### S6.2 DNA sequences for compute domains

As noted in Section S5.1, there are 16 compute domain types in the system:

- 8 "logical" compute domain types (denoted in base 2 as `000`, `001`, ..., `111` or base 10 as `CD0`, `CD1`, ..., `CD7` or sometimes simply `0`, `1`, ..., `7`), Figure S10(a)
- 8 domains that are logically identical to the first 8 (in terms of encoded data/computation) but have distinct sequences to avoid hairpin formation within certain compute strands, denoted (denoted in base 2 as $\overline{\texttt{000}}, \overline{\texttt{001}}, \ldots, \overline{\texttt{111}}$ or base 10 as $\overline{\texttt{CD0}}, \overline{\texttt{CD1}}, \ldots, \overline{\texttt{CD7}}$ or sometimes simply $\overline{\texttt{0}}, \overline{\texttt{1}}, \ldots, \overline{\texttt{7}}$), Figure S10(a)

Compute domains were designed with only a few criteria in mind, Figure S10(c). The main constraint that we designed for was orthogonality between compute domains: interactions between two distinct compute domains (in all possible 4 ways in terms of reverse complementation, i.e. for two non-matching domains $d_i, d_j$ we check the four pairs $(d_i, d_j), (d_i, d_j^*), (d_i^*, d_j), (d_i^*, d_j^*)$) should be unfavorable ($> -2$ kcal/mol) and interaction between complementary compute domains' sequences should be favorable ($< -11.6$ kcal/mol, that is slightly below the average binding energy computed on 10,000 random length-12 strands using the nearest-neighbor model with parameters from Santa Lucia and Hicks [132]) with interactions measured at 53 °C, [Na] = 1 M and $[\mathrm{Mg}^{++}] = 0$ using `binding(A,B) = pfunc(A,B)-pfunc(A)-pfunc(B)` as in [13], SI Section S4.2.1 p. 43, with `pfunc()` given by `NUPACK4` [57]. We designed the sequences using `nuad`[21], a Python library that allows for easy specification of constraints, and runs a stochastic local search for domains using the sequence design principles developed by Woods, Doty et al. [13]. The orthogonality constraint mentioned above was *softly* enforced to favour the convergence of the search algorithm meaning that it was not necessarily achieved by all sequences. In practice orthogonal binding energies ranged from $-3.31$ kcal/mol to $-0.19$ kcal/mol with an average of $-2.00$ kcal/mol.

Note that we intentionally (as a design principle) designed the compute domains independently of the length 120-base scaffold sequence. This choice was motivated as follows:

[19] `https://www.tilibit.com/`

[20] In an earlier experimental iteration that is not reported in this paper, the scaffold sequence order was $A^*B^*C^*D^*E^*$ where the first M13 96-base subsequence was $A^*B^*C^*D^*$ and domain $E^*$ was the second M13 24-base subsequence. The current scaffold order is $E^*D^*C^*B^*A^*$ as given in SI Section S10.1.1

[21] `https://github.com/UC-Davis-molecular-computing/nuad`, project led by David Doty, UC Davis.

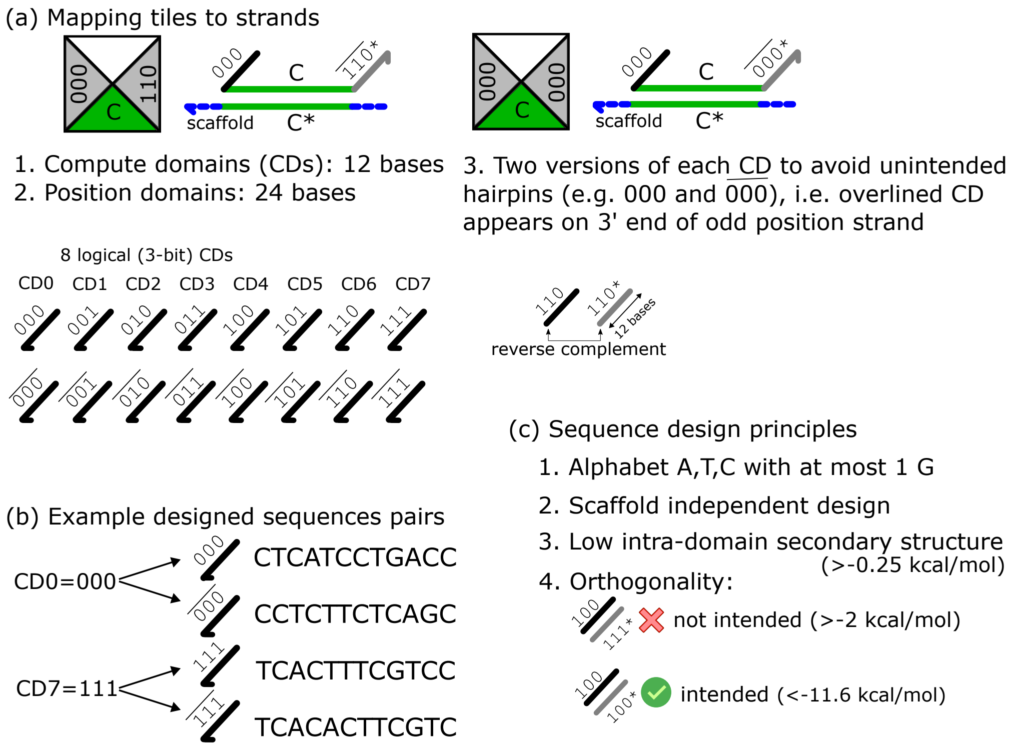

Figure S10: Sequence design principles for compute domains. (a) Mapping from a tile to a strand. Square tiles are rigid, but strands are not: to avoid unwanted hairpin formation—which could happen in any strand with two logically identical compute domains, such as the example one shown on the right—in Section S5.1 we described how each of 8 logical 3-bit strings are represented by two CDs, one with an overline (overlined version is used if a compute domain is at an *odd* scaffold position $(A, C)$ and $3'$ end of the compute strand, or at an *even* scaffold position $(B, D)$ and $5'$ end of the compute strand). Light grey represents the reverse Watson-Crick complement of a compute domain. (b) The two pairs of designed sequences for compute domains CD0=`000` and CD7=`111`. (c) Summary of main principles for compute domains sequence design. Energies were computed using `NUPACK4`'s `pfunc()` function [57] at 53 °C, [Na] = 1 M and $[Mg^{++}]$=0.

1. Designing against spurious interactions with long strands (e.g. 7,249 bases for M13 scaffold) is a challenging problem. One issue is that it tends to over-constrain the sequence design task because of the huge number of possible interactions.
2. With long compute domains, and high formation temperature (likely well above 50 °C), even long compute domains of 12 bases would be likely not to have too many interactions with the scaffold.
3. Having a *universal* set of compute domains that could be used (in theory) with almost any scaffold sequence would allow the use of the domains designed in this work in many other contexts in the future. However, we leave to future work to experimentally test the hypothesis that our domains would indeed work with common, non-adversarially chosen, scaffold sequences.

Despite the above Point 3, with a risk-adverse mindset, we conducted significant post-design analysis on ten sets of designed compute domains before choosing which one of those ten performed best over a set of criteria that included examining some of the many possible unintended interactions between them and our previously-chosen scaffold sequence.

Computation strands were formed by prepending all possible compute domains before, and appending their reverse-complement after, each scaffold-position complementary domain $A^*, B^*, C^*$, and $D^*$, giving 64 computation strands at each scaffold position (apart from position $A$ where only 8 computation strands were needed as we did not put domains on the left side (same orientation as main text Figure 2h) since they are not required at the beginning of the scaffold).

The sequences for computation strands are given in SI Section S10.1.3.

#### S6.3 DNA sequences for reporter strands: Sequence independent reporting mechanism (SIRM)

**Sequence design.** Similar principles were used to design the two domains $F$ and $Q$, respectively labelled with a fluorophore and quencher and used as part of the reporting mechanism (the domains $F$ and $Q$ in Figure 2h of the main text have strand names `3RQ` and `5RF` in Section S10).

- Hard constraints (i.e. non-compliant sequences are rejected):
  1. 20 bases.
  2. Alphabet `A,T,C` and at most 1 `G`.
  3. No patterns `CCCC, GGGG, AAAAA, TTTTT`.
  4. Reverse-complement nearest-neighbour binding free energy (according to a summation of stack-loop free energies in the nearest-neighbour model [132]) is between $-21.7$ and $-21.95$ kcal/mol ($-21.7$ is the average perfect reverse-complement binding energy computed over 10,000 20-base strands) computed at 53 °C using `nuad`'s nearest-neighbour implementation (see Section S6.2).
  5. The 4 bases directly adjacent to the fluorophore on the same strand should not be G [59].
- Soft constraints (i.e. sequences may not comply but are optimised with respect to these criteria):
  1. Intra-domain secondary structure $\geq -0.1$ kcal/mol
  2. Binding energy[22] between `3RQ` and `5RF` $\geq -3$ kcal/mol (in all possible 4 ways in terms of whether or not these two strands are reverse complemented).

##### S6.3.1 Summary and purification

In total, the SDC system tested in this paper consisted of 1,539 strands, including reporting strands. See SI Section S10 for the designed DNA sequences. All strands were ordered unpurified in pH 8.0 buffer from IDT except for nine strands: (a) the 120 base scaffold was a PAGE-purified "IDT ultramer", dry, (b) the four strands with the quencher and fluorophore labels (`3RQ`, `5RF`) were HPLC purified, and (c) four strands that bind to the flurophlor-labelled strand in experiments of scaff-120, namely: `ATTO`$^*B$, `ATTO`$^*C$, `ATTO`$^*D$, `ATTO`$^*E$ (the latter four strands were inconsistently ordered PAGE purified or unpurified at times, without any measured impact on results).

[22] We use the same notion of binding energy as in [13], SI Section S4.2.1 p. 43: `binding(A,B) = pfunc(A,B)-pfunc(A)-pfunc(B)` with `pfunc` given by `NUPACK4` [57].

### S7 Implementation: 10 SDC programs; fast programs; data analysis – experiments

#### S7.1 Experimental parameters and temperature protocols

As noted in the main paper, and Section S1, in our experiments we seek to set experimental conditions (concentration excesses, rate of temperature anneal to explore configuration space, etc.) so that the SDC strand model, in a classical Gibbs free-energy sense, is faithfully approximated with enthalpy trading off against entropy as defined by the Boltzmann distribution. Hence, we use the excesses, sample mixing and buffer conditions described in the Methods section. In particular (a) having compute strands in high excess (10×) over the scaffold (1×) ensures that there is a thermodynamic concentration-driven bias towards each scaffold position having a strand (a requirement of the tile model). (b) Using a temperature anneal, from hot (melted) to cold (assembled scaffolded structures), allows the system to explore a large number of scaffolded configurations, without needing to enforce any precise kinetic pathways.

A 3-hour annealing protocol, used for all 'typical anneals' in the main paper is shown below both in Python source-code (left) used to control the qPCR machine, and graphically (right). We also run faster ("super-fast", 1-min) anneals which are described in Section S7.2.5, as well as 12-minute and other fast anneals for renewable programs described in Section S8).

Compute strand and anchor strands were at 10× over the scaffold for data reported in Figure 3 and Figure 4 of the main text, and throughout the work (except where otherwise noted), but at 5.7× for those renewable programs reported in Figure 5 and Section S8.2.

**Pipetting: acoustic liquid handler versus manual pipetting** All samples providing data reported in the main paper were dispensed using an acoustic liquid handler (Echo 525, Beckman Coulter), for speed and consistency. However, earlier versions of Figures 3–4 were done by hand with good yield, but wider variance as would be expected, hence a liquid handler is not necessary to repeat our results. All samples giving data reported in the Supplementarity Information were also dispensed using an acoustic liquid handler, except where otherwise noted (notably some renewing experiments were tested by hand pipetting, results shown in Section S8.3 and some early tests of the system in Sections S7.5, S7.6 and S7.7).

```
from qslib.common import Protocol, Stage
protocol = Protocol(
    filters = ['x4-m4'],
    stages = [
        # 10 minutes at 80C, 1 datapoint per minute
        Stage.hold_at(temperature=80,
                      total_time="10 minutes",
                      step_time="1 minute",
                      collect=True),

        # 80C -> 20C in 3h, 1 datapoint per 2 minutes
        Stage.stepped_ramp(from_temperature=80,
                           to_temperature=20,
                           total_time="3 hours",
                           n_steps=90,
                           collect=True,
                           start_increment=True),

        # Hold at 20C for 45 minutes, 1 datapoint per 2 mins
        Stage.hold_at(temperature=20,
                      total_time="45 minutes",
                      step_time="2 minutes",
                      collect=True),
    ],
    # volume in units of uL
    volume = 35
)
```

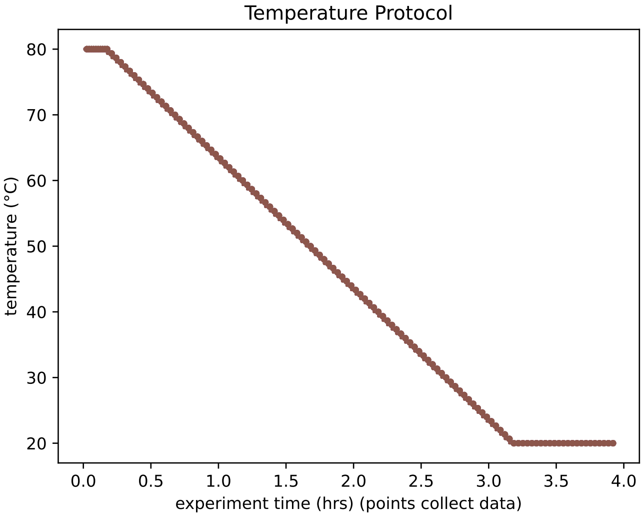

Figure S11: Left: Specification of *typical-anneal* temperature protocol in python, using qslib library [62]. Right: Plot of temperature protocol (the qPCR machine has six temperature zones, which here are all being held at the same temperature hence there are 6 identical overlapping curves).

#### S7.2 Treatment of dataset

##### S7.2.1 Dataset description

The data presented in the main paper corresponds to 506 imaged individual qPCR wells, corresponding to 194 distinct mixes (each mix was repeated at least twice) of which:

- 104 mixes for distinct SDC computations, see SI Section S7.3
- 80 mixes for distinct SDC controls, see SI Section S7.2.4
- 8 mixes for distinct fluorescence controls with no compute domains and 1 for dye-only control (i.e. no scaffold), see SI Section S5.2
- 1 mix for only buffer and tween

**Coverage: usage of designed compute domains** All 16 compute domains' sequences that we designed (SI Section S6) were used in some experiment. The least used compute domain sequence was used 228 times, and the most used compute domain sequence was used 524 times. These counts are obtained by summing all appearances of a compute domain's sequence in each mix (including repeated mixes) – a compute domain can appear several times within the same mix.

##### S7.2.2 Explaining the early drop in signal at the beginning of annealing and the motive behind rescaling the y-axis

Fluorescence data showed a consistent drop in signal at the beginning of each anneal, from 80 °C to roughly 67 °C. To understand the phenomenon, we ran two sets of control experiments on the dye:

1. **SIRM Control experiments.** In these experiments we tested the SIRM on scaffold positions $B, C, D$ and $E$, with the caveat that non-reporting positions were covered with a single-domain complementary strand: For example, if reporting position $C$, positions $C$ and $D$ will be involved in signal reporting with strands ATTO$^*D$, 5RF, 3RQ, $C$–reporting high signal and $CQ^*$–reporting low signal. Positions $A, B$, and $E$ are not involved in the signal reporting and will be covered with compliments. Figure S12(left) shows normalised data of 40 SIRM experiments at all possible scaffold positions that were run as controls to different computation experiments. We hypothesised that this consistent drop in signal happens due to the composition of the reporting mechanism and the effect of having the fluorophore labeled strand becoming double-stranded. Therefore, we sought to test the hypothesis with the next control experiment, described below in bullet point 2. We also ran other control experiments in relation to each computation that are explained in S7.2.4. In the control experiments, it is observed that, although no computation is happening, the high signals drop to a mean value of 0.832.
2. **ATTO-590 control experiments.** To test the hypothesis that the initial signal drop in the experiment is due to the ATTO-590 labeled strand becoming double-stranded (as described in bullet point 1), we ran experiments to compare the single-stranded ATTO-590-labeled strand vs a double-stranded one. Figure S12(right) shows that the double-strand ATTO-590-labeled strand has an initial signal drop similar to what is observed with the reported computing experiments and controls.

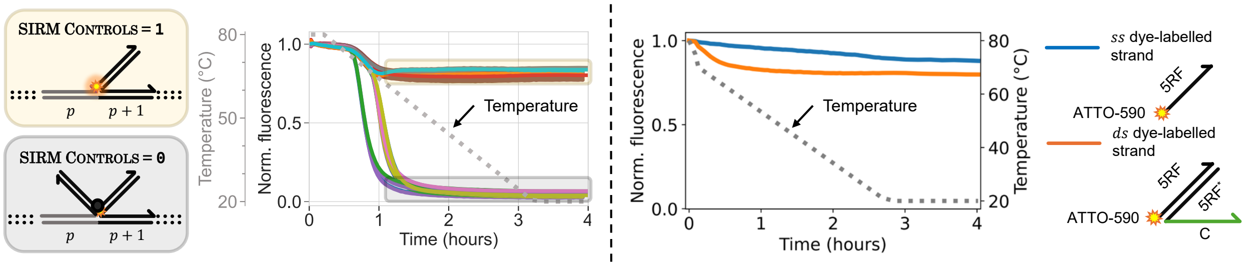

Figure S12: Control experiments showing the initial signal drop as temperature drops from 80 °C to roughly 67 °C. Left: Normalised SIRM control experiments, a signal drop is seen from 80 °C to 67 °C. Right: blue curve shows fluorescence of a single-stranded ATTO-590 labeled strand. The fast initial drop is not seen. The orange curve shows a rapid drop from 80 °C to 67 °C, thus showing the effect of having a strand that is a complement to the ATTO-590 labeled strand. (The blue curves show a slow drop as temperature lowers from 80 °C all the way down to 20 °C, we believe this is due to a temperature-dependence of the due when single-stranded.)

##### S7.2.3 Data treatment (normalisation)

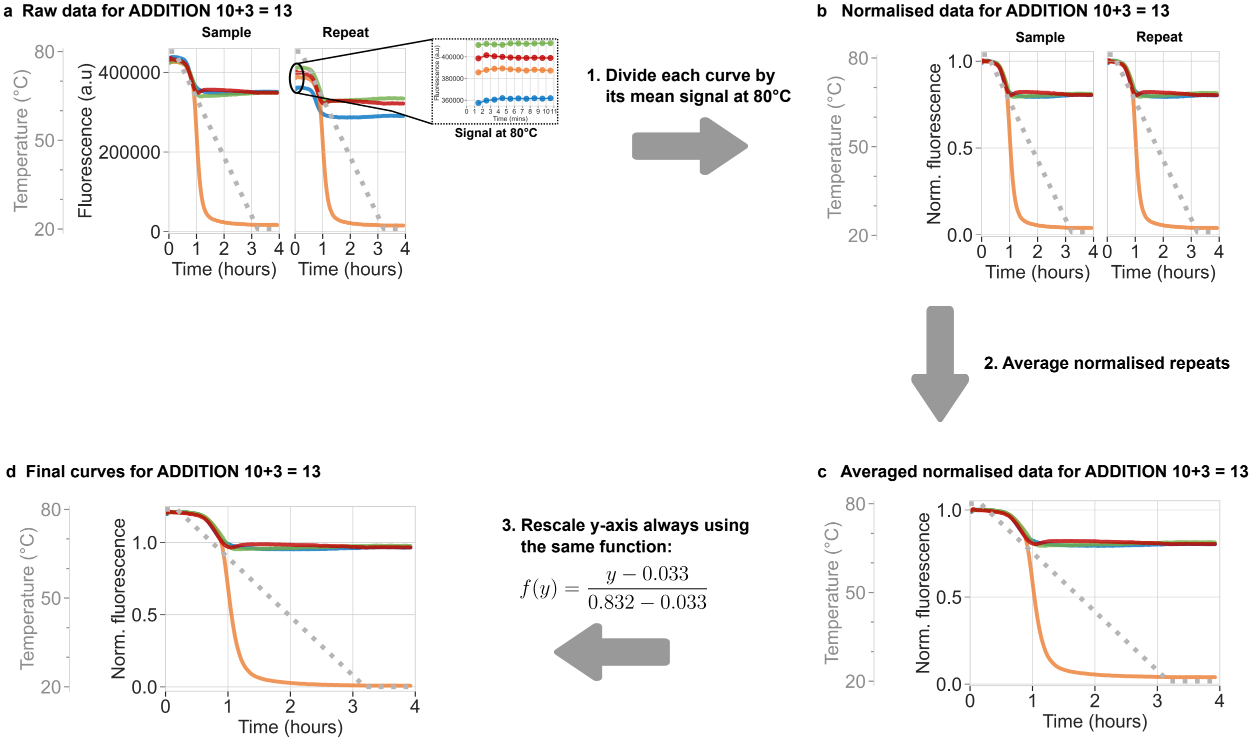

Figure S13: Fluorescence data is processed using a simple 3-step process. **a.** Raw fluorescence data for the Addition 10+3=13 experiment (see Figure 3e in main text) acquired using a quantitative PCR machine (QuantStudio$^{tm}$ 5 Real-Time PCR System controlled using the `qslib`[62] open source python library). Each mix is repeated at least twice, and the raw sample and repeat are shown side-to-side. Zoom-in on the signal at 80 °C is shown. Step 1 of data processing is that each curve is divided by its mean signal at 80 °C (mean of 10 datapoints for that signal). **b.** Normalised data for Addition 10+3=13 experiment. Step 2 of data-processing consists in averaging each sample's curve with its repeat(s) (e.g. here, the two same-colour curves on the left and right side plots are averaged, point by point). **c.** Averaged normalised data for Addition 10+3=13 experiment. Step 3 of data-processing rescales the y-axis by applying the linear transform $f(y) = \frac{y-0.033}{0.832-0.033}$. This same function is applied to all averaged signals at this stage, it maps value 0.832 to 1.0 and 0.033 to 0.0, these values are based on control data analysis, see SI Section S7.2.4. **d.** Fully processed data for Addition 10+3=13 experiment. The data processing steps depicted in this Figure are systematically applied to all of the data shown in the main text. Samples were pipetted using Echo 525 Acoustic Liquid Handler.

We strive to minimally treat data. All of the fluorescence data presented in the main text is processed using only the following three steps, described graphically in Figure S13:

1. **Divide by mean signal at 80°C.** Raw fluorescence curves, Figure S13a, are divided by the mean of their fluorescence data points at 80°C, giving normalised fluorescence data with values in the range $[0, 1]$, Figure S13b.
2. **Average normalised repeats.** Each sample is mixed at least twice giving a set of repeats for the same experiment. Data obtained after Step 1 is averaged amongst repeats, Figure S13c.
3. **Rescale y-axis linearly.** Normalised averaged repeats obtained after Step 2 are rescaled linearly so that y-axis value 0.832 becomes 1.0 and 0.033 becomes 0.0, i.e. the linear function $f(y) = \frac{y-0.033}{0.832-0.033}$ is applied to the data, Figure S13d. Values 0.033 and 0.832 were fitted on 0-reporting and 1-reporting control data respectively; see SI Section S7.2.4. However, for the scaled-up systems in SI Section S9, owing to the number of positions, controls, and scaffolds involved, this rescaling step is omitted, and values are reported directly after step 2. Instead, for these systems, controls are plotted alongside samples.

Data given in this SI document is processed similarly, unless stated otherwise (mainly, step 3 is sometimes not applied in this SI).

**Experimental variance** Experimental variance between repeats was small: between 0.1% and 6.7%, with a mean of 0.6%, of average normalised signal difference between repeats. This small variance was achieved through the use of an acoustic liquid handler (Echo 525, Beckman Coulter).

##### S7.2.4 Control data

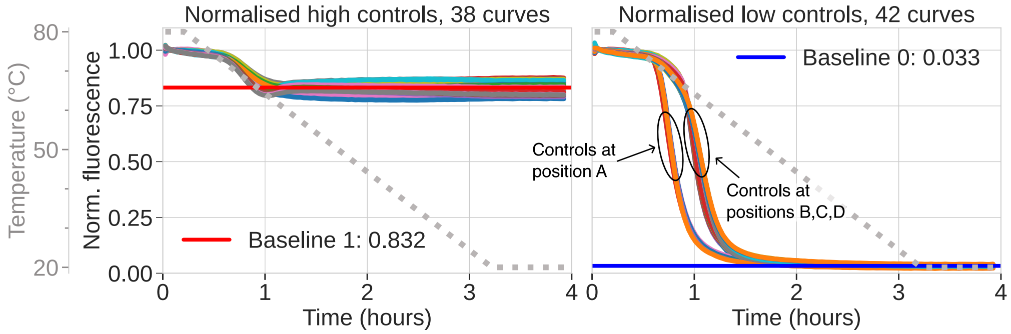

Figure S14: All 80 SDC controls (processed using Steps 1 and 2, SI Section S7.2.3). Left: high controls (reporting 1). Baseline 1, equal to 0.832, is the average of high controls' completion levels (i.e. mean signal at 20°C). Left: low controls (reporting 0). Baseline 0, equal to 0.033, is the average of low controls' completion levels (i.e. mean signal at 20°C). Notably, low controls at position $A$ have a higher melting temperature than controls at positions $B, C, D$: 68.5°C and 63.6°C respectively in average. Samples were pipetted using Echo 525 Acoustic Liquid Handler.

We define *control data* as being SDC samples where computation domains are used but there is no competition: only the correct strand at each scaffold position $\{A, B, C, D\}$ is put in the mix. Within control data, 52% of controls report at position $D$ and the remainder report at positions $A$, $B$ and $C$ evenly (similarly to all experiments, when reporting before position $D$, unused domains are covered using complementary domains).

In contrast, *computation data*, that executes algorithms (such as depicted in main text Figures 3, 4, 5), comes from samples where several strands compete for scaffold positions. Almost all of our computation data has control data that is *directly* associated to it: that is, control data where only the correct combination of strands (corresponding to the algorithm's output) is annealed. Some samples do not have associated control data, mainly in the case where several polymers are formed after anneal (this is the case in 3-STATE NONDETERMINISTIC FINITE AUTOMATON and GRAPHREACH computations, see SI Sections S7.3.5 and S7.3.9).

Control data is shown in Figure S14 and is processed using Steps 1 and 2 of SI Section S7.2.3. Baseline 1 (resp., baseline 0) is computed by taking the average completion level (i.e. mean signal at 20 °C) of all high (resp., low) controls. These baselines are equal to 0.832 and 0.033 respectively and are used in Step 3 of SI Section S7.2.3 to rescale the y-axis of our data.

The following features of our control data can be noted:

1. **Higher standard deviation for high controls.** From Figure S14, one can note that the standard deviation for baseline 1 is higher than for baseline 0: standard deviations are 0.026 and 0.004 respectively. One likely explanation for the higher variance is how we chose our reporting strand excess: details are in Section S7.5, the key point is that for the high signal (and not for the low signal) close to zero quencher-labelled strands should be bound to the scaffold (quenching the fluorophore), hence any large enough sample-to-sample variation in fluorophore concentration will be picked up by the qPCR machine. For low-signal control samples we should not see variation since the excess of the fluorophore strands is $< 1\times$ of scaffold concentration, with the intention being that all fluorophore strands are quenched in low signal samples. This is just one hypothesis, that we did not further investigate. Another hypothesis could be that in high samples, nearby single-stranded bases are affecting the fluorophore in a sequence-dependent manner – something that should not happen for low signal since there are only duplexes in the locality of the dye.

2. **Higher melting temperature of controls at position $A$.** The set of low controls, depicted in Figure S14 right, can clearly be separated in two sets of curves: one set where the melting point[23] is reached before the 1-hour mark, at, 68.5°C, and the other set after the 1-hour mark at, 63.6°C. All controls with the higher 68.5°C melting temperature are controls at position $A$. Strands at position $A$ have two particularities: (a) they are at the 3'-extremity of the scaffold and (b) they do not have any compute domain at their 5'-extremity (otherwise it would just be hanging outside the scaffold). We hypothesise that only (a) explains the difference in melting temperature, giving a thermodynamic (enthalpic) advantage to these strands. This is because we tested (b) with controls at positions $B, C, D$ without seeing a change in their melting temperature.

###### S7.2.5 Super-fast programs

We ran SDC programs as fast as we could, given the equipment we have at hand. We use qPCR machines (QuantStudio$^{tm}$ 5) which have significantly better temperature control properties than other readily available devices common to labs in our field. In particular, qPCR machines have at least an order-of-magnitude better temperature precision and accuracy than platereaders, order-of-magnitude better ramp-speed than plate readers and water-heater/chiller controlled fluorescence spectrophotometers, and can go to higher temperatures than platereaders. In the following protocols we ran our qPCR machines at their fastest speed limit.[24] Our super-fast protocol (Figure S15(a)) asks the machine to change temperature every second (by large decrements of, respectively: $-25, -5, -5, -5$ °C), starting at 80 °C and ending at 55 °C—something that is impossible for the machine to do.[25]

**It is important to understand that in our super-fast protocols, we instruct the qPCR machine to change temperature much faster than it is capable of doing, with the goal of forcing the machine go as fast as it can**, thus our python temperature protocols (Figure S15(a,b)) look rather different than the final experimental temperatures (Figure S15(c,d,e)). Throughout this work, to avoid any confusion on this matter, when we report data we report the qPCR machine 'sample' temperature at data acquisition time. Figure S15 plots time of temperature changes and data acquisition, showing that the qPCR machine only changes temperature after it has acquired data at the specified setpoint. All super-fast data plots in the main paper (Figure 3) and Sections S7.3.1– S7.3.10 are of the style of plot in Figure S15(c), i.e. showing temperature changes and data acquisition times.

[23]Melting temperatures are computed as the average temperature amongst repeats where raw qPCR data (see Figure S13a) reaches the value (max+min)/2.

[24]It would be useful, especially to inform and parameterise theoretical models along the lines suggested in Section S3.5, to push SDC systems to their absolute fastest speed limit. We note that it is possible to build, or purchase, custom, purpose-designed fluorescence-readout systems with faster response times.

[25]For the DivBy2 program we used a slightly different super-fast protocol optimised for ternary readout.

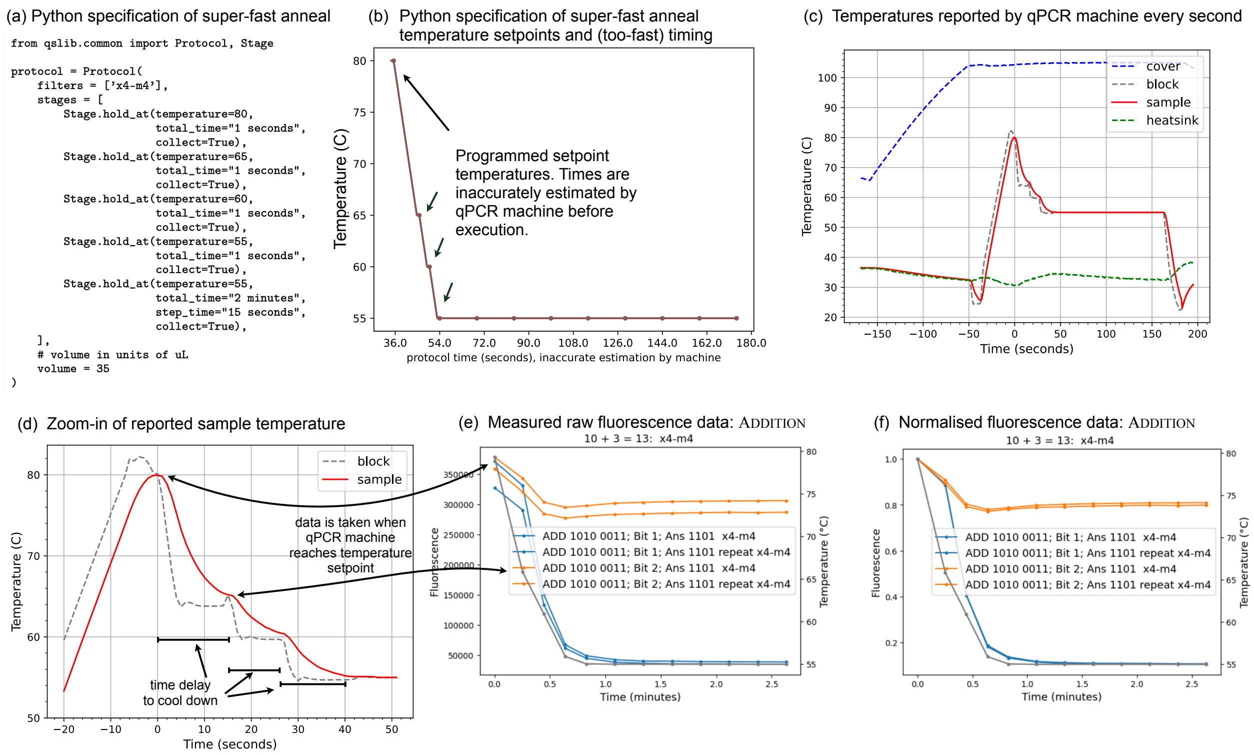

Figure S15: Programming, and assessing, super-fast anneals on a QuantStudio$^{tm}$ 5 qPCR machine. (a) Specification of *super-fast anneal* temperature protocol in python, using the `qslib` [62] library. **It is important to note that the qPCR machine will move at a slower speed than specified in the program.** (b) Plot of temperature protocol, before running it, using `qslib`'s function `Experiment.plot_protocol()` with axis changed to seconds. **It is important to note that this plot has an inaccurate x-axis: the qPCR machine's software is simply estimating how long it will take to run the protocol, likely based on some previous internal calibration outside of our control.** (c) Four temperature values are reported by the qPCR machine, every second: (1) cover (headed lid to prevent condensation inside sample cover/lid), (2) block (contains samples), (3) sample (machine reported sample temperature), (4) heatsink (ancillary device). (d) Zoom-in of sample and block temperature, with explanations of how the machine is responding to our (too fast) commands. (e) Experimental data plotted with respect to recorded temperature at data-acquisition timepoints. (f) Experimental data after normalisation (Section S7.2.3), i.e. as it is reported throughout the main text and SI. Samples of data in (e) and (f) were dispensed using an Echo 525 Acoustic Liquid Handler.

##### S7.2.6 Measuring program performance

SDC program performance was measured using two definitions: (1) the ability to separate bit-`0` from bit-`1` and, (2) the proportion of correctly assembled polymers after anneal. Definition (1) is handled through **binary classification** and definition (2) by introducing two performance metrics (#1 and and #2 below) for **estimating polymer concentration**. Both methods were applied separately on typical and super-fast anneal data.

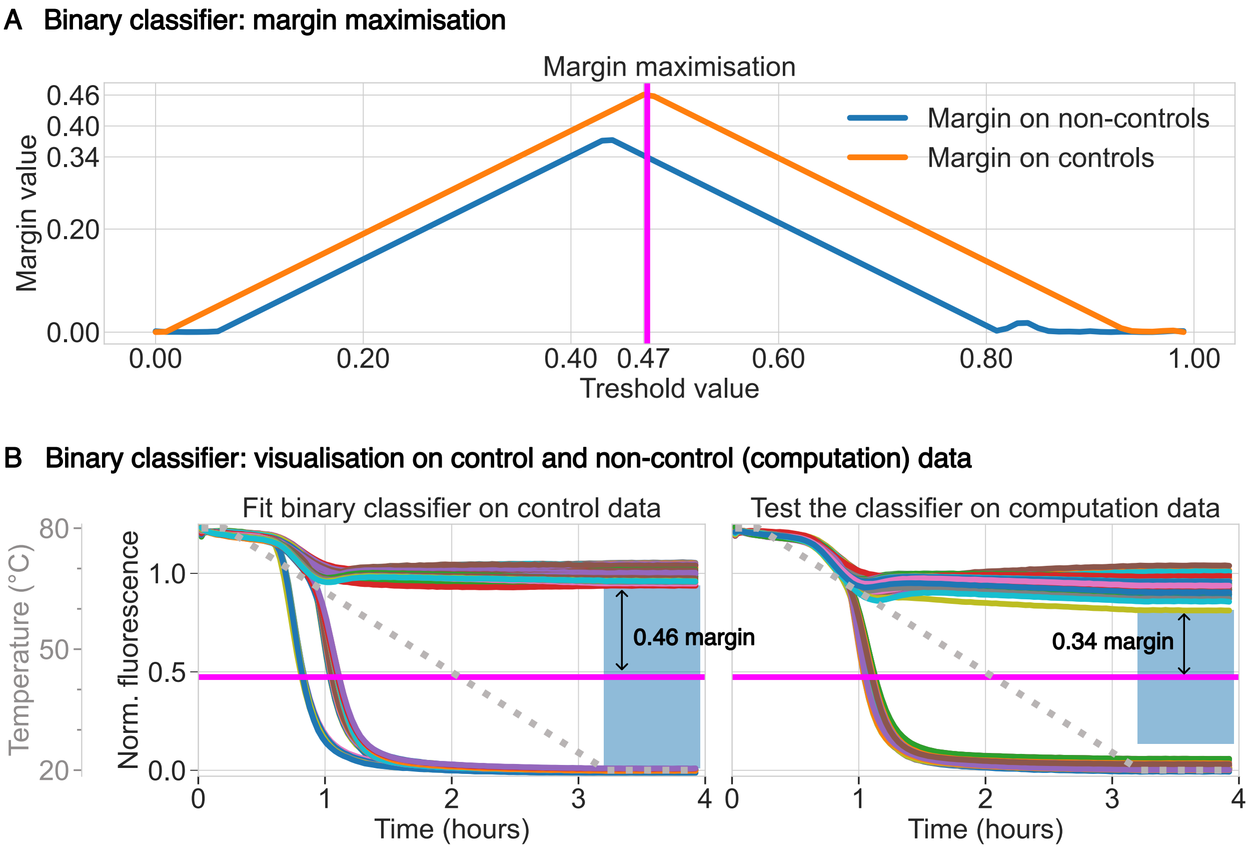

Figure S16: Linear binary classification of typical-anneal completion levels using margin maximisation on control data. Samples were pipetted using Echo 525 Acoustic Liquid Handler.

**Non-quantitative binary classification.** Binary classification, i.e. the ability to distinguish between a high signal for bit-`1` and a low signal for bit-`0`, is the first considered approach for measuring the performance of SDC computations. In practice, a threshold value is fitted on control data (SI Section S7.2.4) below which a normalised (SI Section S7.2.3) typical-anneal signal's completion level (mean value at 20 °C) is considered to be bit-`0` and over which is considered to be bit-`1`. The technique used to fit this threshold is *margin maximisation*, i.e. the threshold maximises the distance to the closest control signal, Figure S16. As seen on Figure S16(A) fitted threshold (magenta curve) for typical-anneals is 0.47 with 0.46 margin (blue area in Figure S16(B,left) meaning that the distance from the threshold to the closest control curve is 0.46. This margin drops to 0.34 on the computation (non-controls) typical-anneal data but the data remains linearly classifiable with 100 % accuracy: all bit-`1` are above the threshold and all bit-`0` are below. Concerning super-fast data, no threshold separates all of our 1-minute data with significant margin, however, for each program, a threshold can be fitted to linearly separate the computation's super-fast 1-minute data points with 100% accuracy.

**Two quantitative metrics: Estimating structure concentration.** We challenge the results of our SDC computations by defining two performance metrics and measuring how our computations perform for each of them:

- **Performance metric #1: "Distance to 0/1 (mean of controls)".** Completion levels (i.e. mean of signal at 20 °C) of typical-anneals and 1-minute data points of super-fast anneals are computed on the processed data (SI Section S7.2.3) and compared to the expected 0 or 1 signal (mean of controls, Section S7.2.4). For $c$ the completion level of an experiment, performance, expressed as percentage, is given by $1 - \mathrm{abs}(1 - c) \in [0, 1]$ if the expected output is 1 and $1 - \mathrm{abs}(c) \in [0, 1]$ if the expected output is 0.

- **Performance metric #2: "Distance to associated control".** When an experiment has an associated control (SI Section S7.2.4), we compute the distance between completion levels of typical-anneals and 1-minute data points of super-fast anneals on the processed data (SI Section S7.2.3) of the experiment and the control. Performance is given in percentage by $1 - \text{abs}(c_{\text{experiment}} - c_{\text{control}})$ with $c_{\text{experiment}}, c_{\text{control}} \in [0, 1]$ the completion levels of the experiment and control respectively.

Metric #2 is not computed on 3-state Nondeterministic Finite Automaton and GraphReach because they do not have controls (because there are several target polymers, see SI Section S7.2.4), excluding them from the scope of metric #2. Also, metric #1 and #2 are not computed on experiments reporting ternary 2 in DivBy2 or intermediary levels in 3-state Nondeterministic Finite Automaton and GraphReach.

**Standard Deviation (SD).** When reporting the mean of a metric (or any other data), we systematically report the Standard Deviation (**SD**) associated to the sample on which the mean was computed, in the format "SD=...". Standard deviation of a sample $\{x_0, \ldots, x_{N-1}\}$ of $N$ elements is defined as $\text{SD} = \sqrt{\sum_{0 \leq i < N} \frac{(x_i - \mu)^2}{N}}$ with $\mu$ the mean of the sample.

| Program | Mean performance metric #1 | Mean performance metric #2 |
|---|---|---|
| Addition | 96.7% (SD=0.027) | 95.0% (SD=0.044) |
| BitCopy | 96.5% (SD=0.028) | 96.2% (SD=0.025) |
| Parity | 93.9% (SD=0.047) | 94.5% (SD=0.039) |
| MultiplyBy3 | 95.6% (SD=0.041) | 95.4% (SD=0.039) |
| DivBy2 | 97.7% (SD=0.025) | 97.6% (SD=0.021) |
| 3-state Nondeterministic Finite Automaton | 94.9% (SD=0.038) | N/A: no controls |
| Counter | 95.8% (SD=0.018) | 95.2% (SD=0.008) |
| GraphReach | 96.4% (SD=0.05) | N/A: no controls |
| Rule110 | 94.2% (SD=0.017) | 95.7% (SD=0.005) |
| BalancedBrackets | 90.4% (SD=0.055) | 93.4% (SD=0.039) |
| Mean over all computations | 95.3% (SD=0.035) | 95.4% (SD=0.035) |

Table 1: Typical-anneal average performance results for metrics 1 and 2 on all computations, see SI Section S7.2.6.

Table 1 reports the mean performance for typical-anneals (3 hours) on each metric for each computation. Both metrics give similar results, with a global mean performance of 95.3% for metric #1 and 95.4% for metric #2.

| Program | Mean performance metric #1 | Mean performance metric #2 |
|---|---|---|
| Addition | 82.4% (SD=0.116) | 84.7% (SD=0.108) |
| BitCopy | 78.3% (SD=0.126) | 76.8% (SD=0.139) |
| Parity | 76.8% (SD=0.124) | 73.6% (SD=0.096) |
| MultiplyBy3 | 85.9% (SD=0.02) | 85.5% (SD=0.099) |
| DivBy2 | 91.9% (SD=0.059) | 93.6% (SD=0.053) |
| 3-state Nondeterministic Finite Automaton | 82.5% (SD=0.109) | N/A: no controls |
| Counter | 81.6% (SD=0.225) | 57.0% (SD=0.096) |
| GraphReach | 93.6% (SD=0.068) | N/A: no controls |
| Rule110 | 79.5% (SD=0.227) | 52.0% (SD=0.056) |
| BalancedBrackets | 78.5% (SD=0.173) | 79.6% (SD=0.116) |
| Mean over all computations | 81.2% (SD=0.16) | 78.8% (SD=0.144) |

Table 2: Super-fast anneal average performance results for metrics 1 and 2 on all computations, see SI Section S7.2.6.

Table 2 reports the mean performance for super-fast anneals (∼1 minute datapoint) on each metric for each computation. We get a global mean performance of 81.2% for metric #1 and 78.8% for metric #2. Discrepancies between metric #1 and #2 on Counter and Rule110 are explained by the fact that, in one minute, non-quenching samples (which are in majority) reach signal level of 1, hence giving a good score on metric#1, while quenching samples (which were the only ones to have associated controls) are still quite far from reaching signal level 0, already reached by their associated control, see SI S7.3.7 and SI S7.3.8. Interestingly, the performance of super-fast anneals is not considerably lower than the performance of typical-anneals.

#### S7.3 10 SDC programs: design and results

This section presents each of the 10 implemented SDC programs and their results in details.

##### S7.3.1 Addition

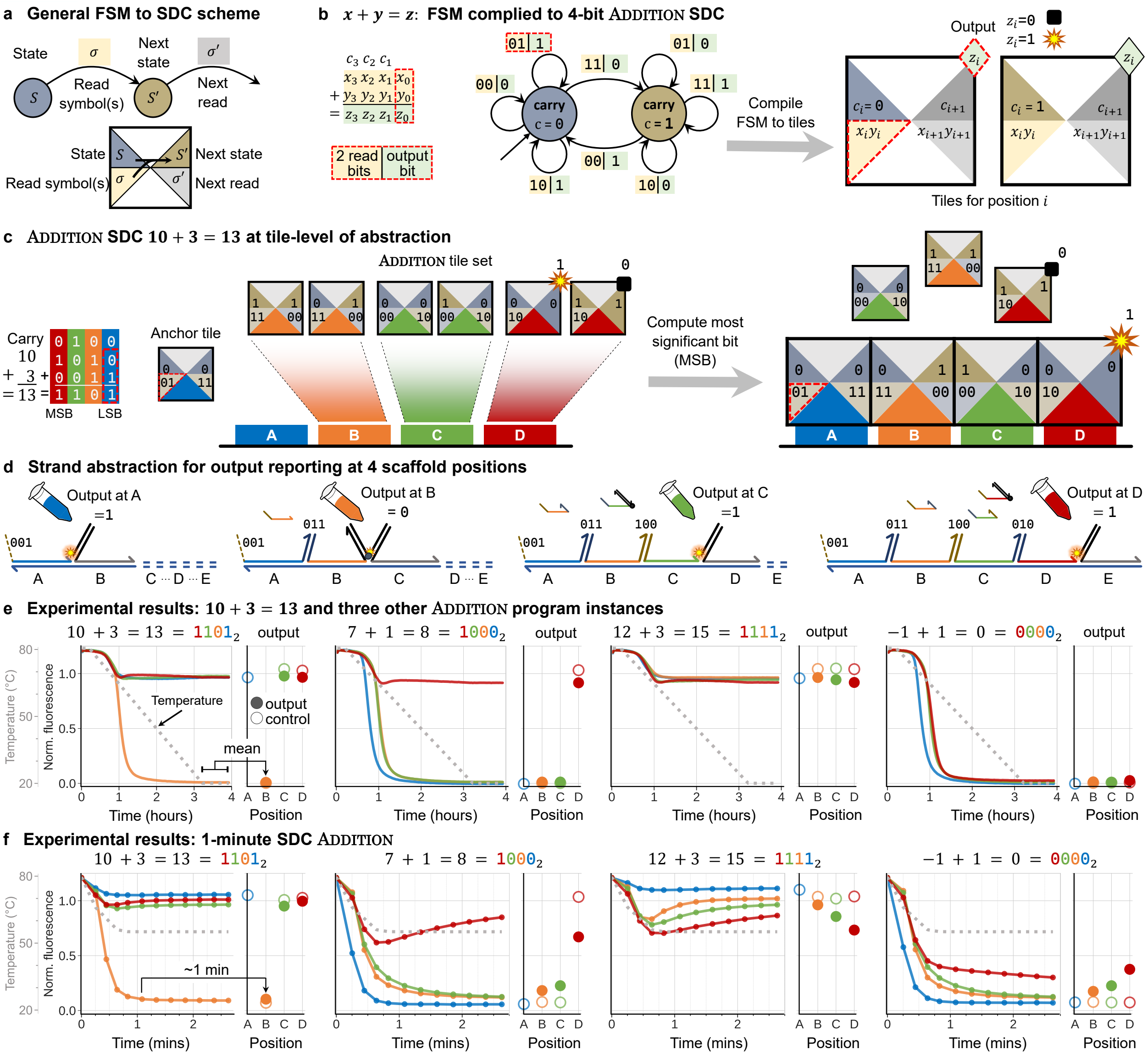

Figure S17: Programming and implementation of the Addition FSM. a. General compilation scheme from FSM into tiles, each compute domain encodes an FSM state and transition. b. The Addition FSM adds two binary numbers $x + y = z$: each transition reads two bits, $x_i$ and $y_i$, writes output bit $z_i$ and enters carry state $c_i \in \{0, 1\}$. The Addition FSM is compiled to an SDC with two tiles per scaffold position. c. Example: $10 + 3 = 13$, using colour to illustrate how bit positions map to scaffold positions. The target configuration is the most favourable as it is the only configuration with zero mismatches. d. Strand diagrams for reporting at each of positions A (output bit = 1), B (= 0), C (= 1), and D (= 1). e. Experimental result for $10 + 3 = 13$, as well as three other Addition experiments. In each case the traces show signal with respect to time/temperature, and the dot plot shows completion level as mean of 22 datapoints at 20 °C. f. Super-fast 3 min anneal for Addition, with the dot plot showing readout at 1 min. Samples were pipetted using Echo 525 Acoustic Liquid Handler.

We ran the following four additions with SDCs:

- $10 + 3 = 13$ corresponding to $1010 + 0011 = 1101$ in binary.
- $7 + 1 = 8$ corresponding to $0111 + 0001 = 1000$ in binary.
- $12 + 3 = 15$ corresponding to $1100 + 0011 = 1111$ in binary.
- -1 + 1 = 0. This addition uses the overflow over four bits: $1111 + 0001 = (1)0000$, only four bits of the output are used.

The results for all these additions are given in main text Figure 3e (typical-anneal protocol) and Figure 3f (super-fast protocol).

Addition contains a computational sink (see SI Section S4.1.1) as soon as two binary 0s (resp. two 1s) are added together: whether the current carry is 0 or 1, the next carry will be 0 (resp. 1). This situation occurs in $10 + 3 = 13$ and $7 + 1 = 8$. Performance for computations with sinks is better than without, see SI Section S7.2.6.

#### S7.3.2 BitCopy

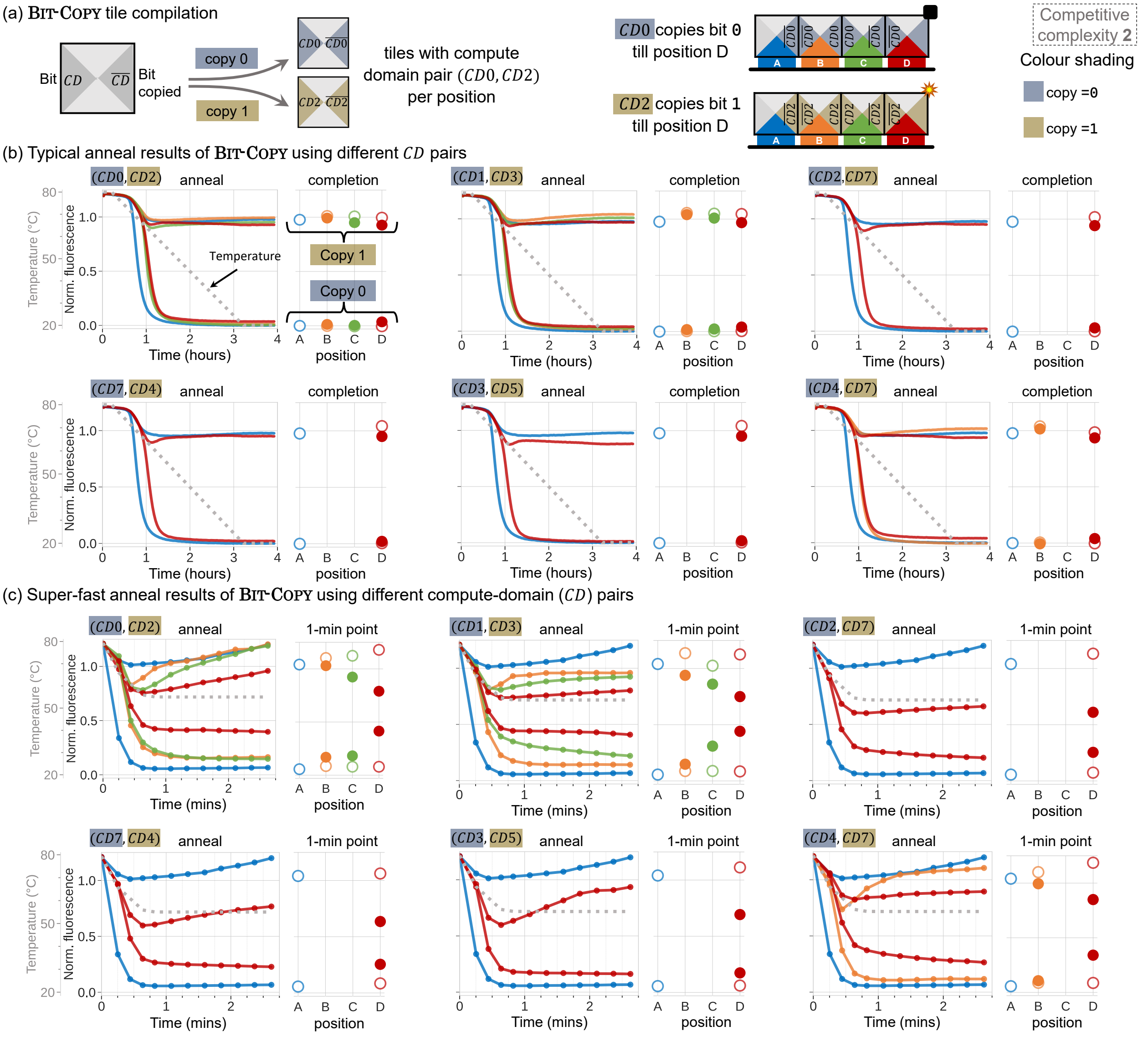

Figure S18: Details of BitCopy SDC program. (a) Tile compilation into compute-domains. Example of compute-domains pair (CD0,CD2) means the former in the pair copies 0 and propagates low signal (i.e. quench) while the latter copies 1 and propagates high signal (i.e. no-quench). (b) Typical anneal results of BitCopy for all tested compute domain pairs and reporting positions (i.e. distance to anchor anchor position A). Completion levels as mean of the last 22 points in the annealing curve are given on the right of each plot. (c) Super-fast anneal results for all tested compute-domain pairs with the 1-min data point. Samples were pipetted using Echo 525 Acoustic Liquid Handler.

Several pairs of computation domains were tested for BitCopy. Pair (CD0,CD2) means that computation domains CD0 represents signal 0 (quenching) and CD2 represents signal 1 (non-quenching). We test the ability of the pairs to propagate their signal at various distances from anchor position A, results are depicted in Figure S18:

- Pair (CD0,CD2) at positions B, C, D.
- Pair (CD1, CD3) at positions B, C, D.
- Pair (CD4, CD7) at positions B and D.
- Pair (CD7, CD4) at position D.

- Pair (CD3, CD5) at position D.
- Pair (CD2, CD7) at position D.

The fact that results for pairs (CD4, CD7) and (CD7, CD4), which only swaps which computation domain quenches, are similar in quality gives support to the claim that a non-quenching signal, which could in theory corresponds to any other reason than having the correct output, is indeed meaningful and corresponds to the SDC reporting a 1 as intended.

Although BitCopy is arguably a *simple* computation from the algorithmic point of view, it does not have any computational sink (see SI Section S4.1.1), which makes it a computation of interest from the SDC point of view.

##### S7.3.3 Parity

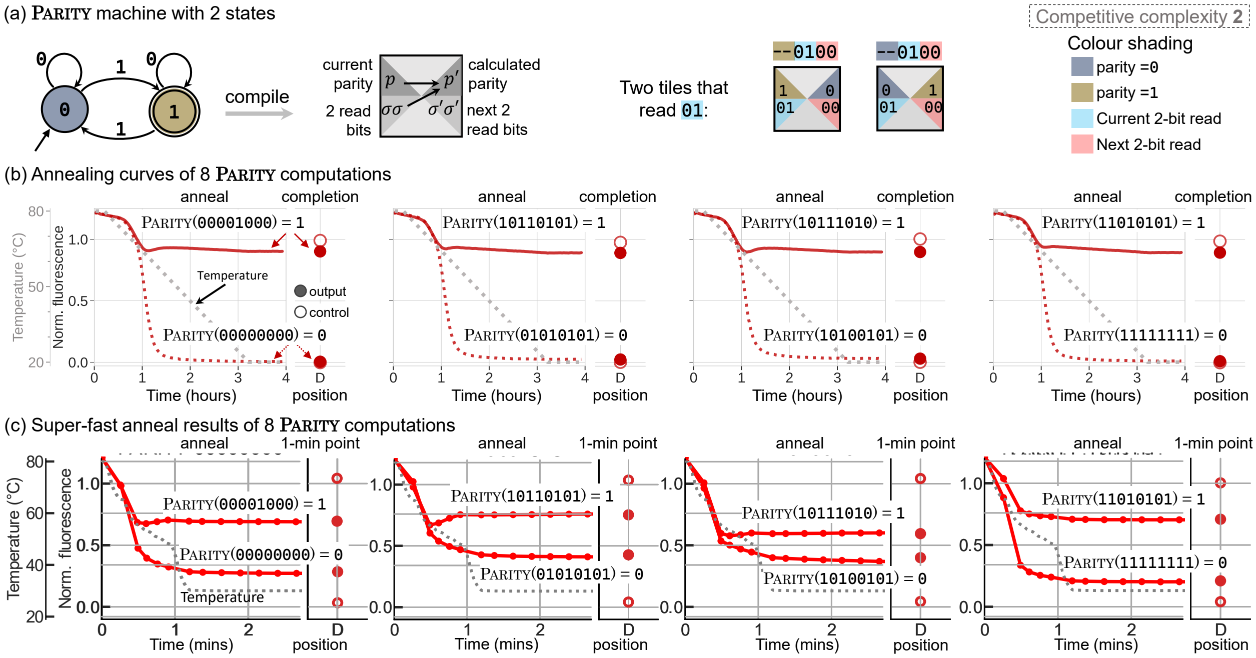

Figure S19: Details of the SDC Parity program. (a) The Parity FSM computes the parity of 1 in a binary input ending at state 1 if the number of 1s in the input is odd and at state 0 if the number of 1s is even. The FSM is compiled to SDC tiles by reading two bits and parity of previous read input sequence at the left side and the next two read bits with the calculated parity of the current read on the right side. Each scaffold position gates two competing tiles, one assumes the parity of previous input sequence is odd (1) and the other assumes it is even (0). (b) Experimental results of 8 Parity computations, two per plot. Solid curves shows inputs with Parity 1 and dotted curves show inputs with Parity 0. Completion levels as mean of the last 22 points in the annealing curve are given on the right of each plot. (c) Super-fast results for Parity: 0/1 results are distinguishable at the 1-minute mark. Samples were pipetted using Echo 525 Acoustic Liquid Handler.

Parity is a program that outputs 1 if and only if the binary input contains an odd number of 1s, the corresponding Finite State Machine is given in Figure S19 (a). This program is of interest in theoretical computer science because it is known to separate the complexity class $AC^0$ (see definition[26]) from P: Parity is in P but not in $AC^0 \subset P$[76]. On our 3-bit SDC, one bit is used to encode the parity state (even or odd) and two bits are used to encode the input, which allows for 8-bit input on our 4-position scaffold, Figure S19 (a). Both typical-anneal and super-fast anneals produce the correct output on all our tested inputs, Figure S19 (b) and (c).

[26] $AC^0$ is the class of problems solved by uniform Boolean circuits of unbounded-fanin AND, OR gates, and fanin-1 NOT gates, of constant depth and polynomial size in input length n.

##### S7.3.4 MultiplyBy3

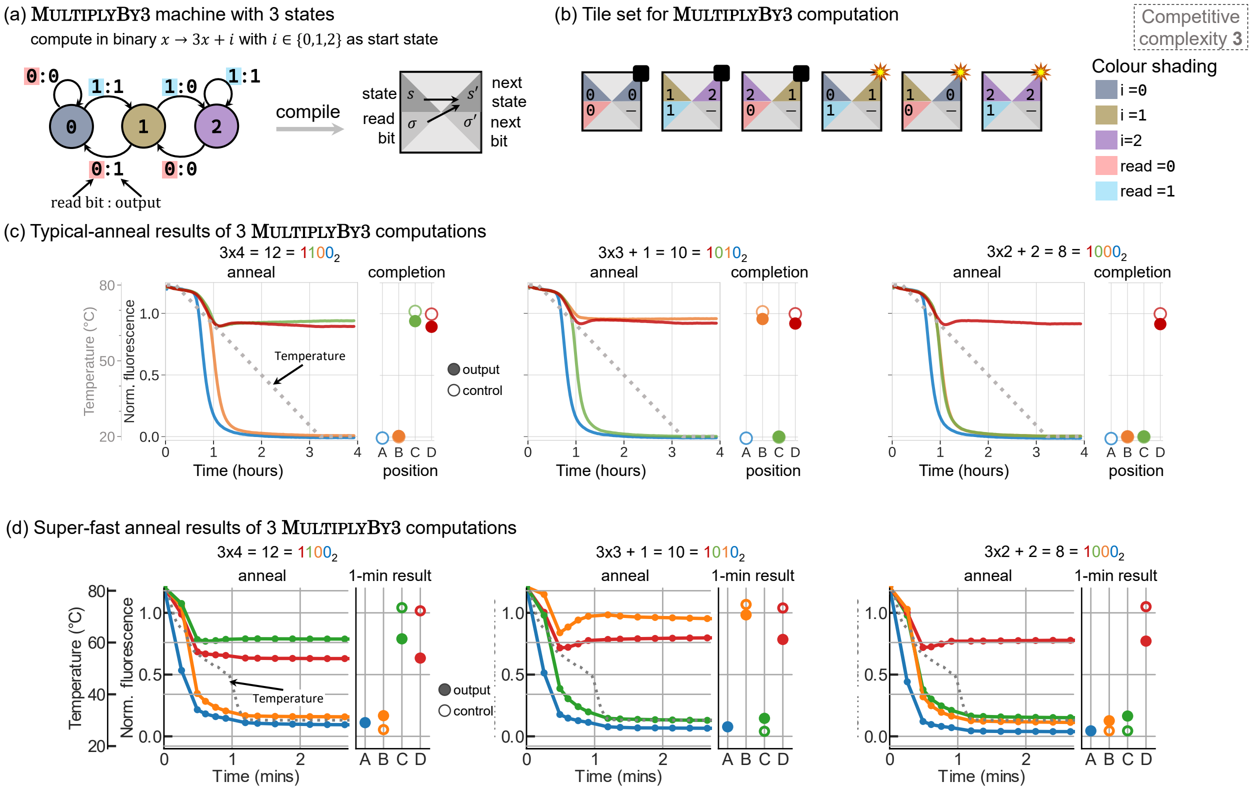

Figure S20: Details of the SDC MultiplyBy3 program. (a) Finite state machine that compute $x \to x + i$ where the start state in a computation is the value of $i$ with $i \in \{0, 1, 2\}$ and its compilation to SDC tiles. (b) The full set of tiles of the SDC MultiplyBy3 program. Three tiles of the set compete at every position in a MultiplyBy3 computation giving competitive complexity of 3. (c) Typical-anneal results of three different MultiplyBy3 computations. On the right of every anneal plot is the completion level plot as mean of the last 22 points in every annealing curve. (d) Super-fast anneal results for all tested inputs with the 1-min data points. Samples were pipetted using Echo 525 Acoustic Liquid Handler.

MultiplyBy3 computes multiplication by 3 in binary: when processing a binary input (reading from Least Significant Bit) starting in state $i \in \{0, 1, 2\}$, the FSM depicted in Figure S20 (a) outputs the binary representation of $3x + i$. Competitive complexity (i.e. number of tiles/strands in competition at scaffold positions B, C, D) is equal to 3 as there are 3 states in the FSM. Figure S20 (c) and (d) give the results of computations $3 \times 4 = 12$, $3 \times 3 + 1 = 10$ and $3 \times 2 + 2 = 8$ both using typical-anneal and super-fast anneal.

##### S7.3.5 3-STATE NONDETERMINISTIC FINITE AUTOMATON

3-STATE NONDETERMINISTIC FINITE AUTOMATON is a non-deterministic computation (see SI Section S4.1) because reading a bit `1` in state 0 leads to both states 1 and 2, see Figure S21 (a). In practice, this means that the execution of the machine on an input bit string yields a Directed Acyclic Graph (Definition S4.2) instead of a linear succession of states. We label the states 1, 2 of 3-STATE NONDETERMINISTIC FINITE AUTOMATON with $Y$ for "yes" and state 0 with $N$ for "no". The machine *accepts* an input if and only if at least one "yes" is reached, Figure S21 (d). See an example computation on input `11101` in Figure S21 (e).

In terms of SDC, nondeterminism implies that several target structures (*polymers*) can be annealed depending on the input instead of having only one. The relative proportions of each target polymer can be deduced, in theory, from the Directed Acyclic Graph of the computation (Figure S21 (e)), see Remark S4.7.

Figure S21 (f) and (g) give the experimental SDC results of 3-STATE NONDETERMINISTIC FINITE AUTOMATON for both typical and super-fast anneals. For each experiment, the expected final proportion of target structures with output "yes"/"no" are given schematically inside red dots: we expect either 1, 2, 3 or 4 target structures depending on the input. Concerning the four inputs where half "yes" / half "no" population is expected (such as for input `1101`), we measured an average normalised fluorescence signal of 0.34 (SD=0.022) which is less than 0.5, Figure S21 (f). Given that these four results are rather tightly located around their mean, we hypothesise that there is a small thermodynamic bias responsible for lowering the signal below 0.5.

Finally, it is worth mentioning that the equivalent minimal deterministic machine, Figure S21 (c), has 7 states instead of 3, which could not have been implemented with our 3-bit SDC (we would have needed 3 bits for encoding the state and 1 bit for the input, 4 bit total), which demonstrates one of the utility of running nondeterministic machines in a constrained setup. Another advantage of nondeterminism is that it takes advantage of the parallel potential of molecular computing: with trillions of scaffolds in solution we can afford to split the final population of target structures.

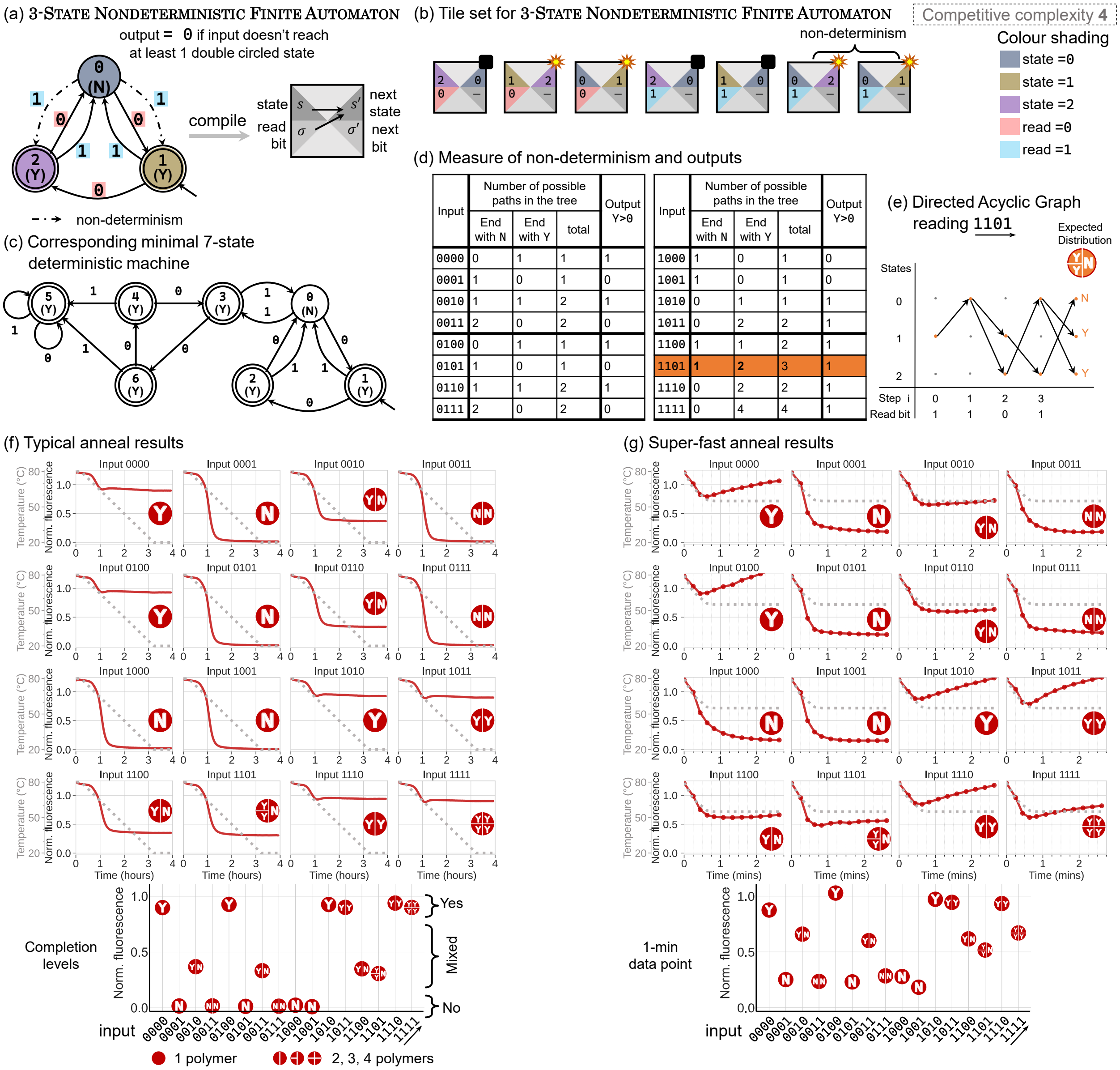

| Input | Number of possible paths in the tree: End with N | End with Y | total | Output Y>0 |
|---|---|---|---|---|
| 0000 | 0 | 1 | 1 | 1 |
| 0001 | 1 | 0 | 1 | 0 |
| 0010 | 1 | 1 | 2 | 1 |
| 0011 | 2 | 0 | 2 | 0 |
| 0100 | 0 | 1 | 1 | 1 |
| 0101 | 1 | 0 | 1 | 0 |
| 0110 | 1 | 1 | 2 | 1 |
| 0111 | 2 | 0 | 2 | 0 |
| 1000 | 1 | 0 | 1 | 0 |
| 1001 | 1 | 0 | 1 | 0 |
| 1010 | 0 | 1 | 1 | 1 |
| 1011 | 0 | 2 | 2 | 1 |
| 1100 | 1 | 1 | 2 | 1 |
| 1101 | 1 | 2 | 3 | 1 |
| 1110 | 0 | 2 | 2 | 1 |
| 1111 | 0 | 4 | 4 | 1 |

Figure S21: Details of the SDC 3-state Nondeterministic Finite Automaton program. (a) 3-state Nondeterministic Finite Automaton of 3 states and its compilation to an SDC tile. Dotted-connectors show non-determinism where a read bit value can lead to either of two states. Accept states are of answer Y meaning Yes (no-quench) and reject state gives answer of N which means No (quench). (b) The full tile set of the SDC 3-state Nondeterministic Finite Automaton program. Non-determinism is shown by having two tiles reporting high signal (i.e. Y) while ending at either state 2 or state 1. (c) Simulation of the 3-state Nondeterministic Finite Automaton by its deterministic, minimal, equivalent DFA with a cost of more states. Here it is 7-states instead of 3: it would not have been possible to run the deterministic version on our 3-bit SDC. (d) The machine output and number of Y and N answers of 16 possible 4-bit inputs. (e) Example execution of input 1101: the computation is an acyclic graph, 2 Y and 1 N are reached hence this output is 1 (at least 1 Y). (f) Typical-anneal results of computations with 16 possible 4-bit inputs to the 3-state Nondeterministic Finite Automaton. Below the curves is the completion level plot as mean of the last 22 points in every annealing curve. (g) Super-fast anneal results of computations with 16 possible 4-bit inputs to the 3-state Nondeterministic Finite Automaton. Below the curves is the 1-min data point plot as mean of the exact value of the data point taken at 1 min in every annealing curve. Samples were pipetted using Echo 525 Acoustic Liquid Handler.

##### S7.3.6 DivBy2

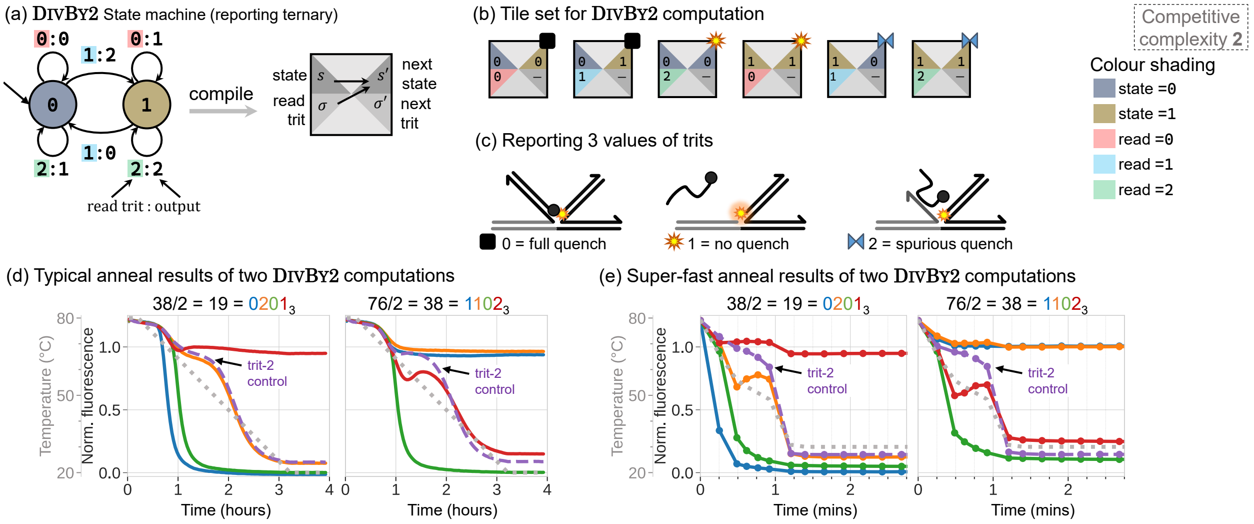

Figure S22: Details of the SDC DivBy2 program where computation is in ternary, i.e. base 3 of three digits $0, 1, 2$. (a) Finite state machine of DivBy2 program showing two states where each state reads a trit and output a trit. FSM is followed by its compilation to SDC tiles. (b) The full set of DivBy2 tiles. Two tiles compete at every position in a DivBy2 computation. One of three symbols on top right of every tile indicating the three output trits. (c) Strand diagram of the reporting mechanism showing the three trits. Details of ternary reporting is in section S5.2.2. (d) Typical-anneal results of two DivBy2 computations. (e) Super-fast anneal results of two DivBy2 results. Samples were pipetted using Echo 525 Acoustic Liquid Handler.

DivBy2 computes division by 2 in ternary (base 3): when processing a ternary input (reading from Most Significant Trit) starting in state $i \in \{0, 1\}$, the FSM depicted in Figure S22 (a) outputs the ternary representation of $(x - i)/2$ if $x - i$ is divisible by two[27]. The machine is structurally similar to the Parity FSM, Figure S19 (a), and that is no coincidence: a base 3 number is divisible by 2 if and only if its number of `1`s is even.

In order to report base-3 digits (trits) we employ a serendipitously-discovered trick, that we call "spurious quenching", depicted in Figure S22 (c) and explained in Section S5.2.2.

Typical-anneal and fast-anneal results for DivBy2 are given in Figure S22 (d) and (e).

[27] If $x - i$ is not divisible by two, running the machine infinitely by appending to the input $0^\infty$ will output the so-called 3-adic representation of $(x - i)/2$. For instance, the 3-adic representation of $5/2$ is ...121012102.

##### S7.3.7 Counter

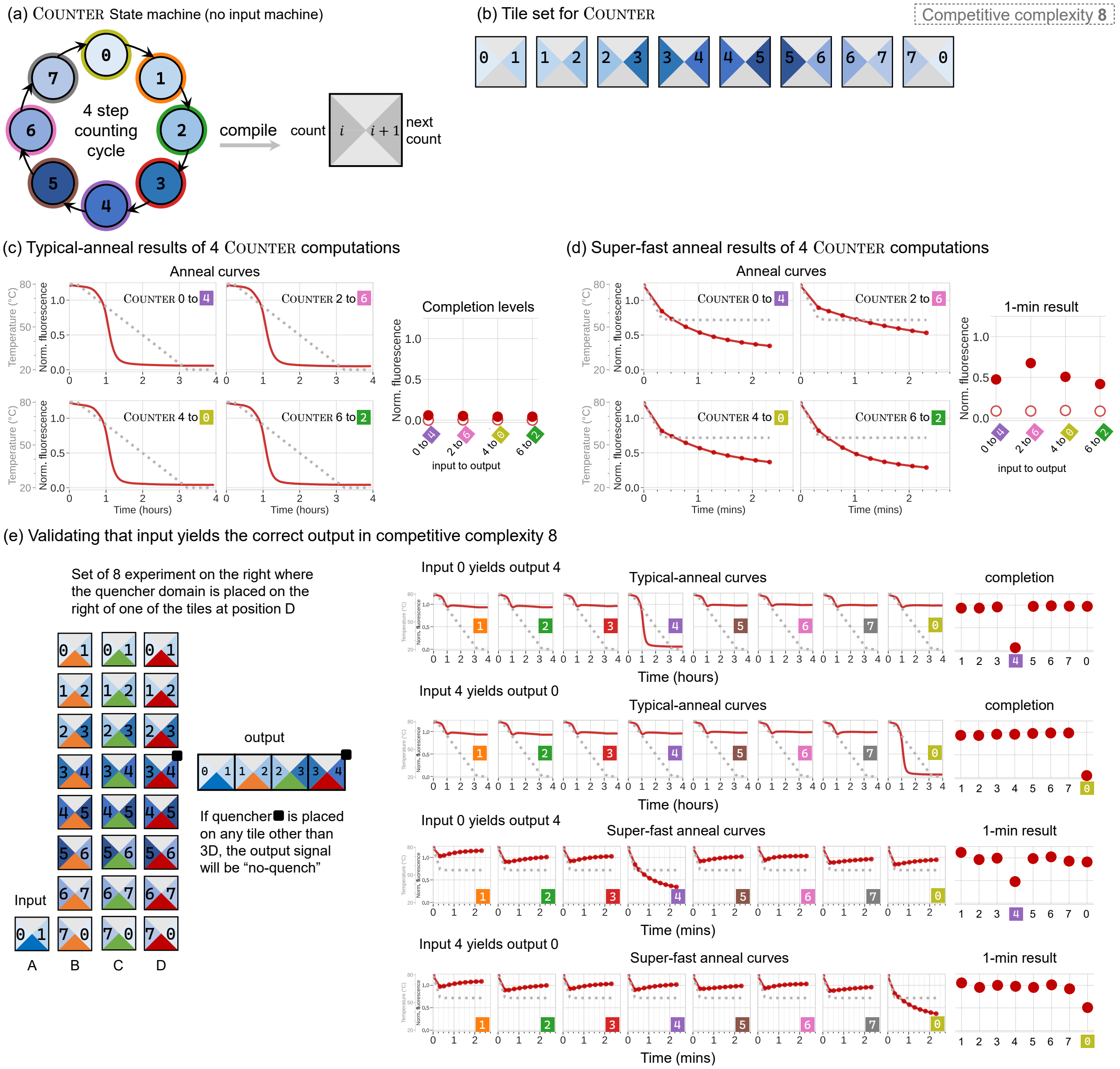

Figure S23: Details of the SDC Counter program. (a) An 8-state Counter finite state machine with its compilation to an SDC tile. The machine can start at any state and do a following 4 counts. For example if it starts at 1, it ends at $1 + 4 = 5$ and the count is $1 \rightarrow 2 \rightarrow 3 \rightarrow 4 \rightarrow 5$. (b) The SDC Counter tile set. (c) Typical-anneal results of 4 Counter computations with their completion level plot as mean of the last 22 points in every annealing curve. The correct target structure reports with quench signal. (d) Super-fast anneal results of 4 Counter computations with their 1-min result plot. (e) Validating input yielding to the correct output in a program of a competitive complexity 8. At each computing domain $B, C,$ and $D$, 8 tiles are competing. The anchor tile at $A$ leads the computation and selects the following tiles. The quencher domain is only present at one tile at position $D$ in an experiment. If the quencher domain is placed on any tile other than the correct answer, the signal will be a no-quench, see Section S5.2.3. 8 experiment results are showing that quenching only happens when the quenching domain is at the correct tile at the end of the counting cycle. For example, when input is 0, it yields an output of 4. Both typical-anneal and super-fast anneal results are given for two sets of experiments. Samples were pipetted using Echo 525 Acoustic Liquid Handler.

##### S7.3.8 RULE110

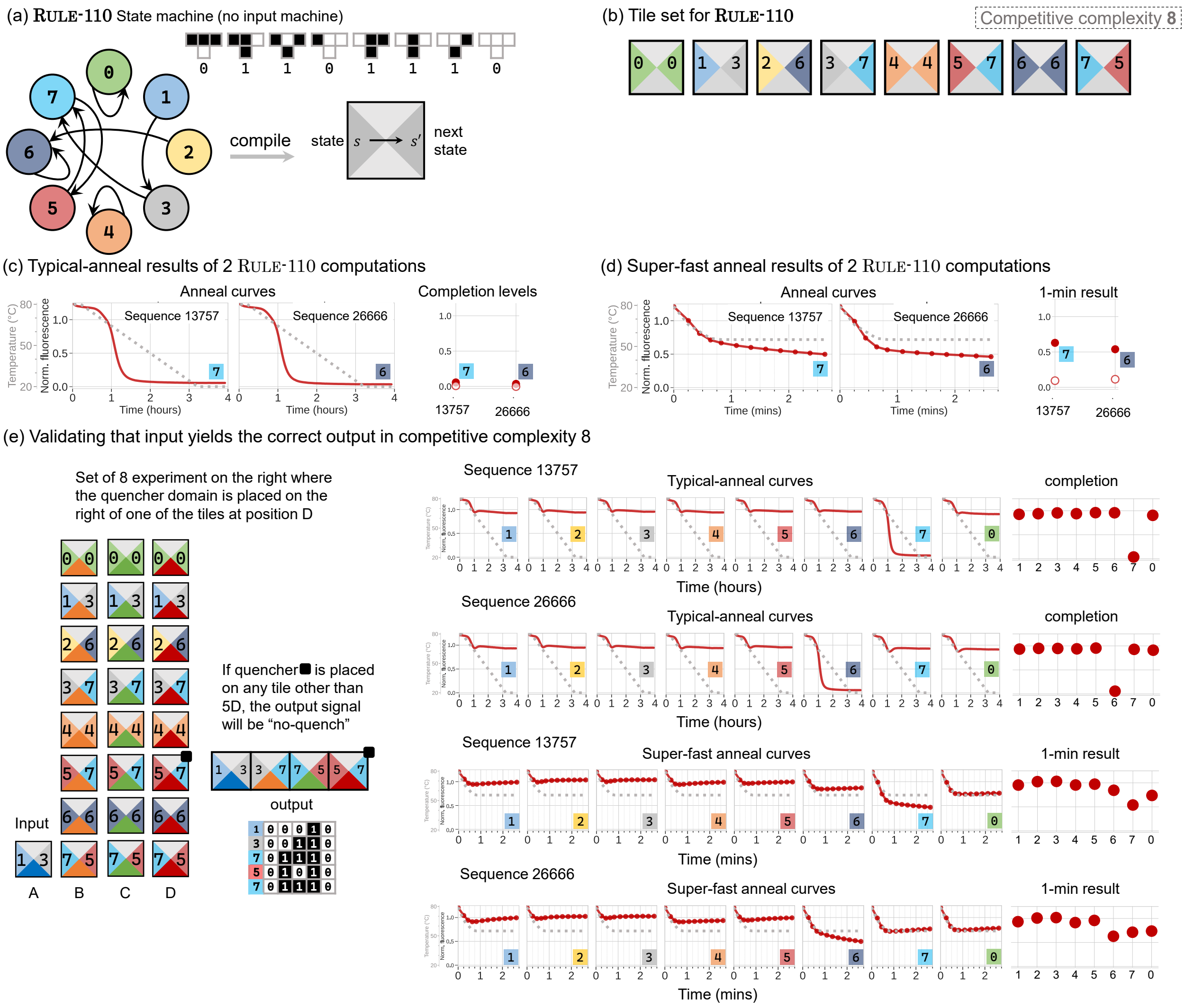

Figure S24: Details of the SDC RULE110 program. (a) An 8-state RULE110 finite state machine with its compilation to an SDC tile. The machine can start at any state run a sequence of 4 digits. For example if it starts at 1, the sequence is $1 \to 3 \to 7 \to 5 \to 7$. (b) The SDC RULE110 tile set. (c) Typical-anneal results of 2 RULE110 computations with their completion level plot as mean of the last 22 points in every annealing curve. The correct target structure reports with quench signal. (d) Super-fast anneal results of 2 RULE110 computations with their 1-min result plot. (e) Validating input yielding to the correct output in a program of a competitive complexity 8. At each computing domain $B, C$, and $D$, 8 tiles are competing. The anchor tile at $A$ leads the computation and selects the following tiles. The quencher domain is only present at one tile at position $D$ in an experiment. If the quencher domain is placed on any tile other than the correct answer, the signal will be a no-quench, see Section S5.2.3. 8 experiment results are showing that quenching only happens when the quenching domain is at the correct tile at the end of the counting cycle. For example, when input is 1, it yields an output of 7. Both typical-anneal and super-fast anneal results are given for two sets of experiments. Samples were pipetted using Echo 525 Acoustic Liquid Handler.

RULE110 is an important program in theoretical computer science because it is the only (up to symmetry) Elementary Cellular Automaton that is known to be Turing Complete[144], and efficiently so [145]. We encode 5-bit RULE110 configurations with our SDC, Figure S24 (e) shows such encoding. We use the same reporting system as for COUNTER: we perform 8 experiments per input, only varying the position of the quencher extension on the tile at position D, see Section S5.2.3; only one of these 8 experiments should quench and this is what we observe in both Typical and fast anneals.

##### S7.3.9 GraphReach

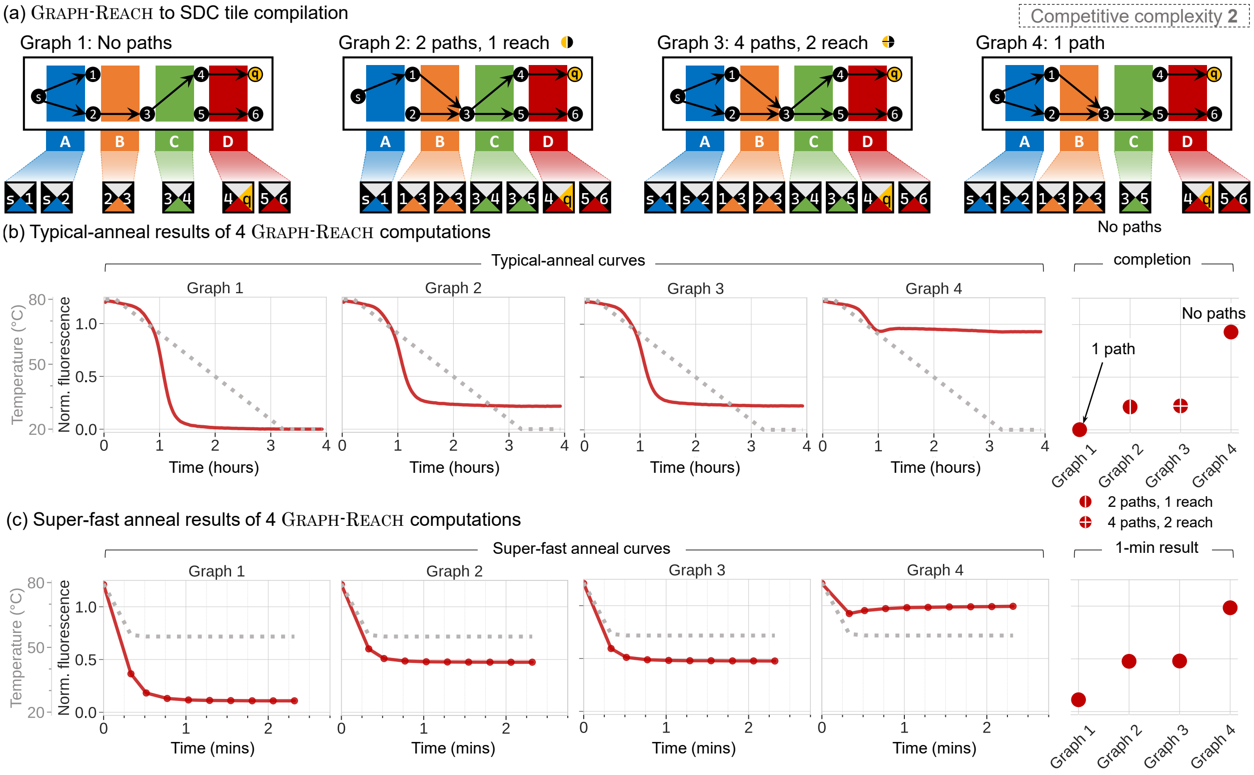

Figure S25: Details of the SDC GraphReach program. (a) 4 graphs each has different reach/no-reach paths. Below each graph is its compilation to SDC GraphReach tiles. (b)Typical-anneal results of the 4 graph computations with their completion level plot as mean of the last 22 points in every annealing curve. The correct target structure reports with quench signal. Where there is only correct path/s, a full quench happens as seen in results of Graph 1. When no path reaches the destination (q), no-quench signal is reported as in graph 4. When some paths reach destination while others don't, we get different reporting structures mixed in solution that some of them report no-quench and some report quench signal. Hence, the signal of Graph2 and Graph3 is above the full-quench level. (c) Super-fast anneal results of the 4 GraphReach computations in (b) with their 1-min result plot. Samples were pipetted using Echo 525 Acoustic Liquid Handler.

GraphReach is a *natural* computation to run with SDC: edges at different layers of a Directed Acyclic Graph are encoded as tiles at different scaffold positions, Figure S25. A node with out-degree greater than 1 (such as node 3 in graph 2) yields several target structures. In the case of graphs 2 and 3, we expect half of the target structures to quench and the other half not to quench; similarly to 3-state Nondeterministic Finite Automaton, SI Section S7.3.5, we witness a signal that is below expected 0.5, yet distinguishably above 0.

##### S7.3.10 BalancedBrackets

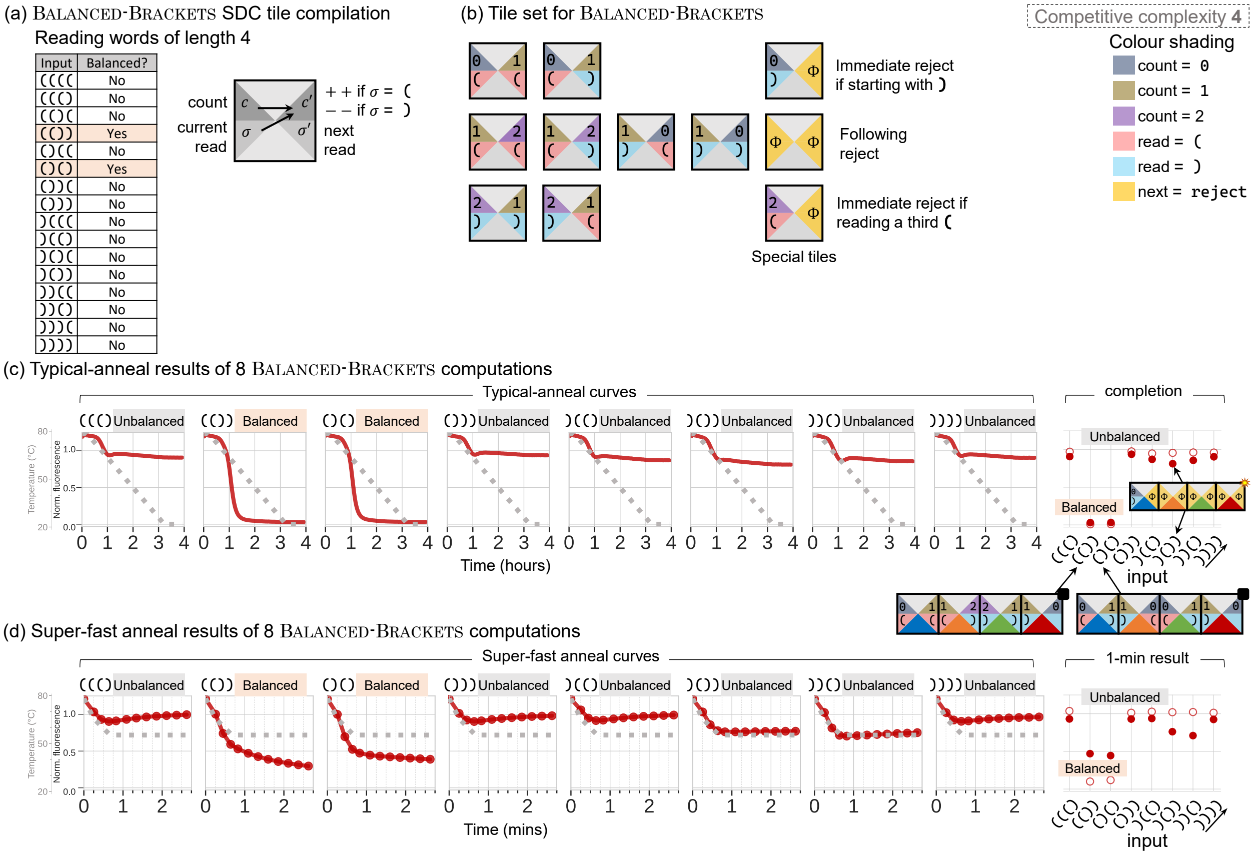

| Input | Balanced? |
|---|---|
| (((( | No |
| ((() | No |
| (()( | No |
| (()) | Yes |
| ()(( | No |
| ()() | Yes |
| ())( | No |
| ())) | No |
| )((( | No |
| )(() | No |
| )()( | No |
| )()) | No |
| ))(( | No |
| ))() | No |
| )))( | No |
| )))) | No |

Figure S26: Details of the SDC BalancedBrackets program. (a) The SDC BalancedBrackets reads words of 4 characters which can either be ( or ). Compilation to tiles shows that the left side encode the current count of brackets and reads the new one, while the left side compute the count after the new read by either incrementing if ( or decrementing if ). (b) The SDC BalancedBrackets tile set. The set includes 3 special tiles for special reject cases like a word starting with a ). (c) Typical anneal results of 8 BalancedBrackets computations where only if balanced a quench takes place. On the right is their completion level plot as mean of the last 22 points in every annealing curve. (d) Super-fast anneal results of the same 8 BalancedBrackets computations shown in (c). Samples were pipetted using Echo 525 Acoustic Liquid Handler.

BalancedBrackets is program that is used by most coding editor to warn the user in case of unbalanced sequence of parentheses/brackets, such as ())). This program is also of theoretical interest because it cannot be implemented by FSM in general (i.e. arbitrary input size). In its 4-character restriction, BalancedBrackets can be implemented using a FSM, hence SDC, Figure S26 (a) and (b). Notably, implementation is set to early reject invalid strings, such as the ones starting with ). We ran BalancedBrackets on 8 inputs and report balanced strings using quenching, see SI Section S5.2.3. Both typical and fast anneals are able to correctly recognise the only two balanced strings (()) and ()(), Figure S26 (c) and (d).

##### S7.4 Robustness over time: long temperature holds show no yield decrease

Figure S27 shows a 20-hour long temperature hold for program BalancedBrackets. We notice no degradation of either high or low signals over this period of time (photobleaching accounts for 1.33% loss of high signal in 20 hours): SDC-style computation is robust over time and does not suffer from leak [27].

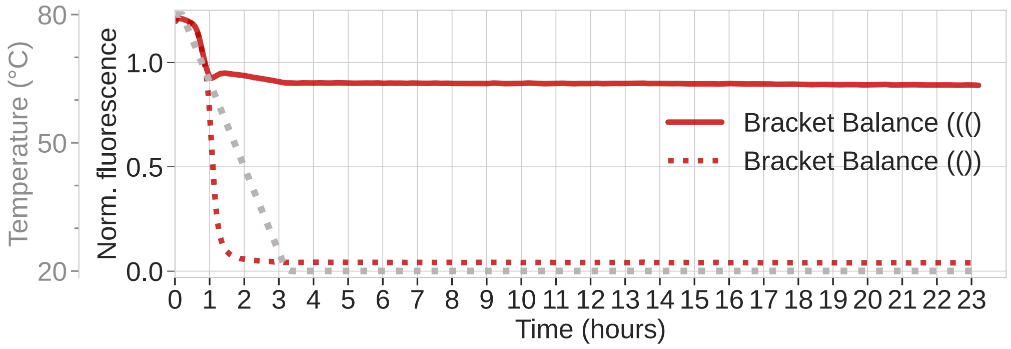

Figure S27: Long temperature hold (20 hours) after typical-anneal for BalancedBrackets program. No yield decrease is measured over time for either high (unbalanced input) or low (balanced inputs) signals. Samples were pipetted using Echo 525 Acoustic Liquid Handler.

##### S7.5 Strand concentration

###### S7.5.1 Scaffold concentration

The SDC-style computations should, in principle, work at a wide concentration range. We chose a scaffold concentration of 100 nM (termed "1×" concentration throughout) as this gave smooth, noise-free, signals with our qPCR-machine based assay, which ran on a QuantStudio$^{tm}$ 5 Real-Time PCR System. At 10 nM, high and low signals were still distinguishable but signal-to-noise ratio was worse (Figure S28 bottom right). We believe that further qPCR machine calibration or optimization or the use of different equipment, such as a platereader, would be sufficient to obtain smooth results at 10 nM or even 1 nM (the typical scaffold concentration for DNA Origami [28])—however it should be noted that typical platereaders have significantly poorer temperature control abilities than qPCR machines, so likely a net loss.

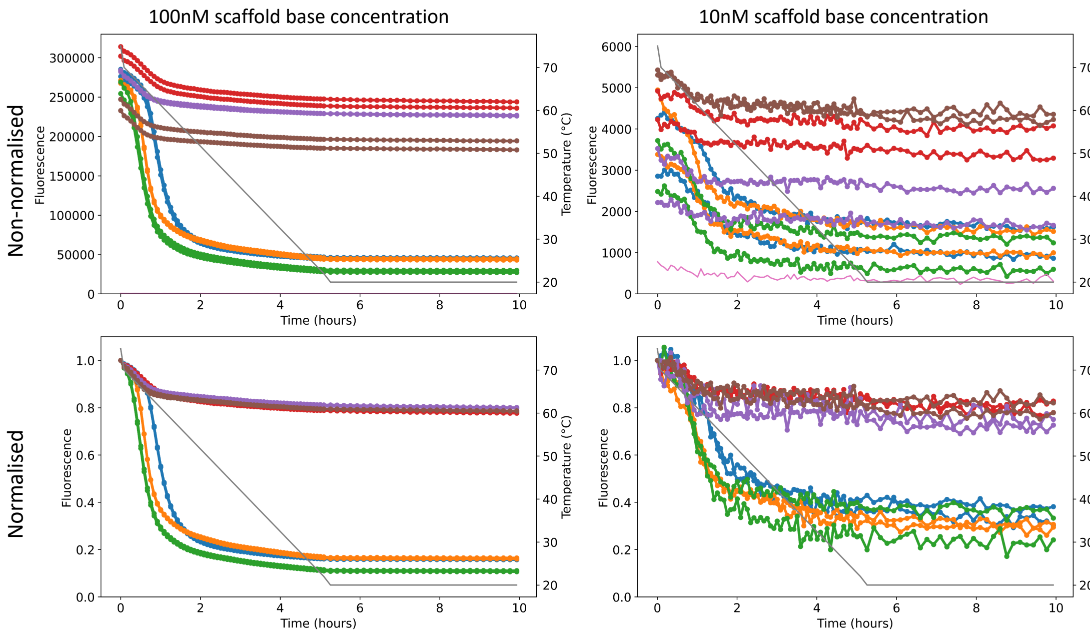

Figure S28: Quench and non-quench signals of the SIRM reporting binary (described in SI section S5.2.1) with scaffold concentrations of 100 nM (left) and 10 nM (right). Top: raw data. Bottom: data normalised by dividing by the initial raw data value at 80 °C. Samples were pipetted by hand.

##### S7.5.2 Strand concentration for anchor, compute and reporting strands

Excesses (assume scaffold excess of $1\times = 100$nM, chosen as discussed in the previous subsection):

- $10\times$ of every compute strand, the goal is that each[28] scaffold position gets a strand
- $10\times$ of anchor strand at position $A$, same goal as above

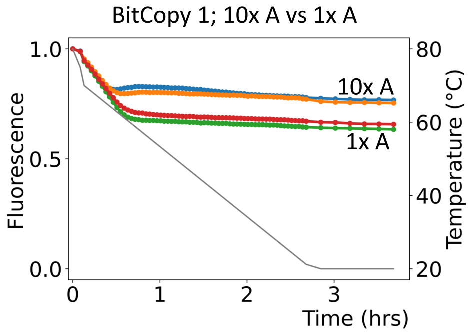

Figure S29: Data justifying our choice of anchor at position $A$ excess of $10\times$, versus $1\times$, shown for BitCopy of distance 4 copying bit value 1. Here $1\times$ = scaffold concentration at 100nM. Samples were pipetted by hand.

- $17\times$ of quencher-labeled strand (3RQ): the goal is to ensure that every[28] reporting-low strand (with a quencher-complement domain), which is at $10\times$, gets a quencher.
- $0.8\times$ of fluorophore-labeled strand (5RF): the goal is to ensure that every[28] fluorophore is bound to a structure (since scaffold is at $1\times$). We tested $1.2\times$ and $0.8\times$ with some controls, and the results were: less variance on fluorescence signal with $0.8\times$ versus $1.2\times$, see Figure S30.

[28]Up to some equilibrium-determined value, at the very leats with our chosen excess this value should be high.

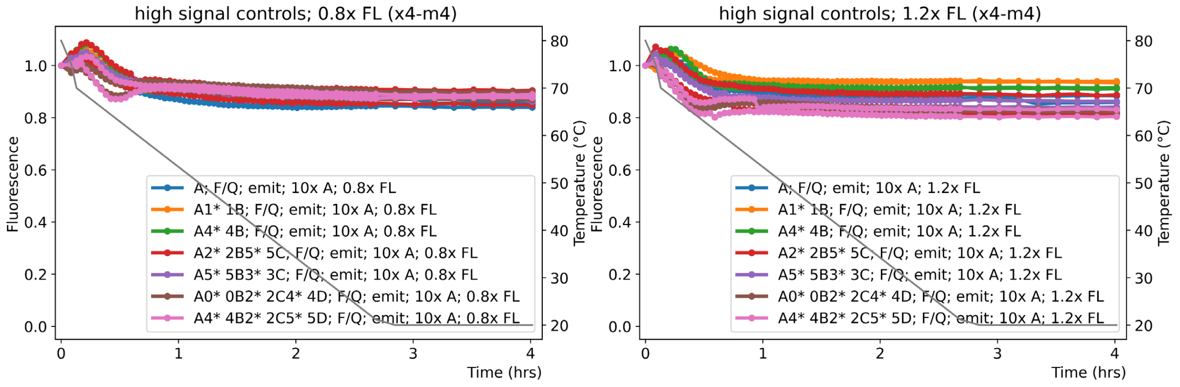

Figure S30: Effect of fluorophore-labeled strand excess on bit-1 reporting control data (controls are defined in SI Section S7.2.4): 0.8× fluorophore-labeled strand excess (left), 1.2× fluorophore-labeled strand excess (right). Excesses are relative to scaffold concentration. Samples were pipetted by hand.

- 0.9x of sequence-independent reporting strand: the goal is to ensure that every[28] reported structure has a fluorophore complex on it. A fluorophore complex consists of the strand labeled with the dye (5RF) and the sequence-independent reporting strand (ATTO*_).

#### S7.6 Starting a computation at various scaffold positions

Here, we show that computations can be started at any position $A, B, C$, or $D$, and ended at any position $A, B, C$, or $D$. Computations starting specifically at $A$ and ending at any of $A, B, C, D$, were shown in the main text as various results of the 10 programs reported there. Figure S31 shows designs, and Figure S32 shows data.

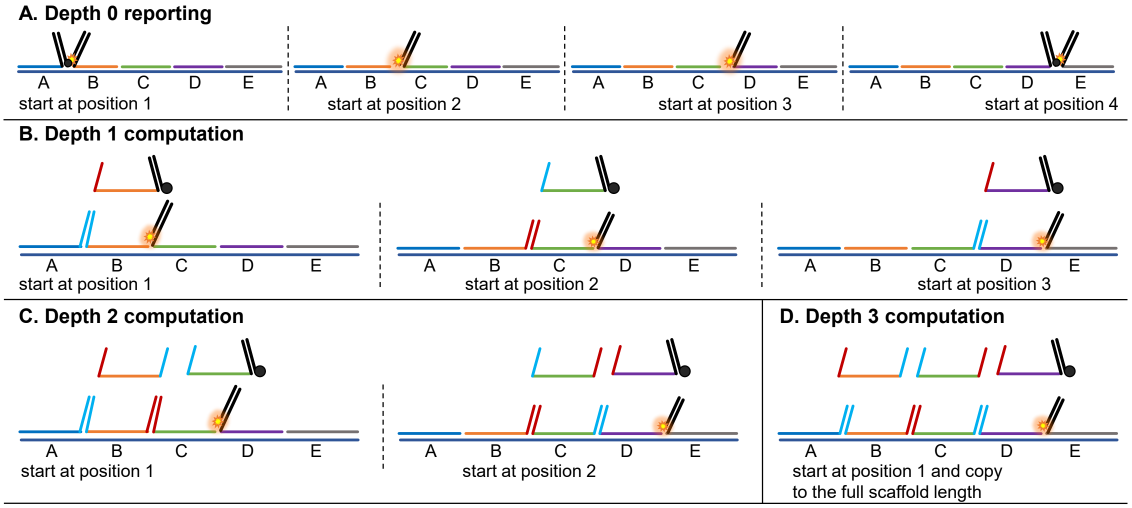

Figure S31: Computations can start and end at any scaffold position. On a scaffold with 5 position domains, we may: (A) Report a non-computing signal (scaffold distance 1) either high or low with no computation on 4 positions of the scaffold. (B) Compute with scaffold distance 2 at 3 starting scaffold positions. (C) Compute at distance 3 starting at 2 scaffold positions. (D) Compute at distance 4 at starting at only 1 scaffold positions.

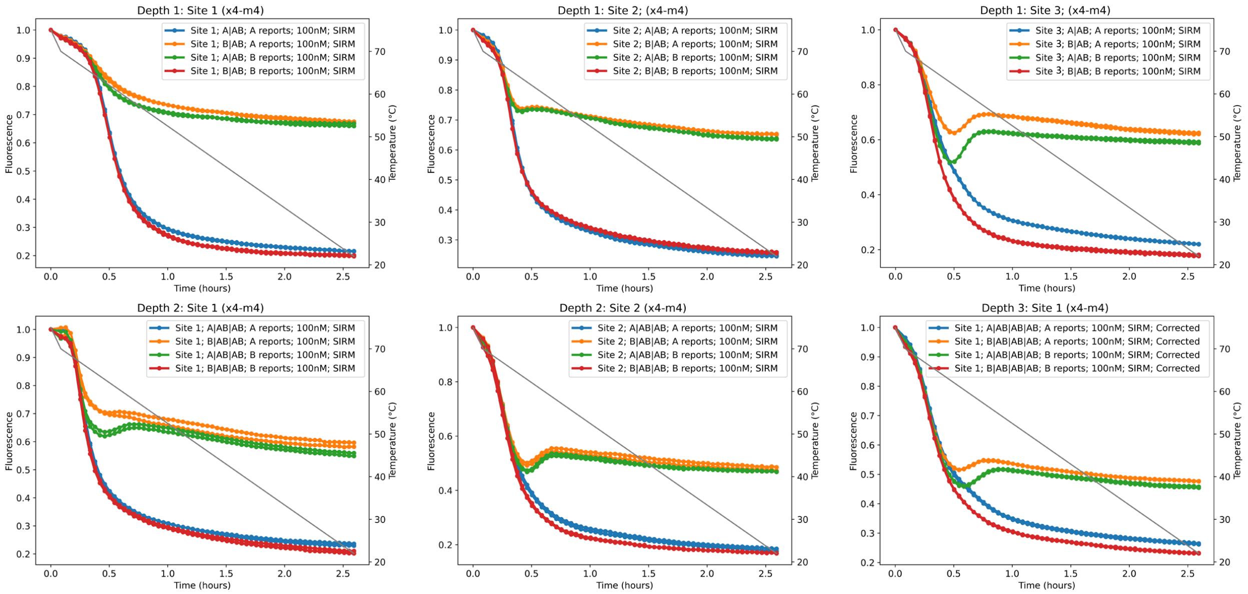

Figure S32: Data from early stages of the project (using manual pipetting and a somewhat slower temperature protocol than reported elsewhere) showing computations at various different scaffold positions, reporting at various positions and with computations of various scaffold distances. Depth in the figure means scaffold distance (measured in units of scaffold positions). Samples were pipetted by hand.

#### S7.7 Justification for scaffold purification

The 120 nt scaffold was initially tested unpurified, Figure S33 left. The curve shows a signal of BitCopy with input bit-1 (orange and green curves) and BitCopy with input bit-0 (red and blue curves). The high and low signal levels are distinguishable. IDT reports that synthesizing strands of lengths above 60 nt results in lower coupling efficiency, decreasing the full-length product yield. Therefore, we switched to using a PAGE-purified scaffold from IDT with the product named IDT Ultramer DNA Oligos. Figure S33(right) shows an improved signal when reporting 'high' with the PAGE-purified scaffold.

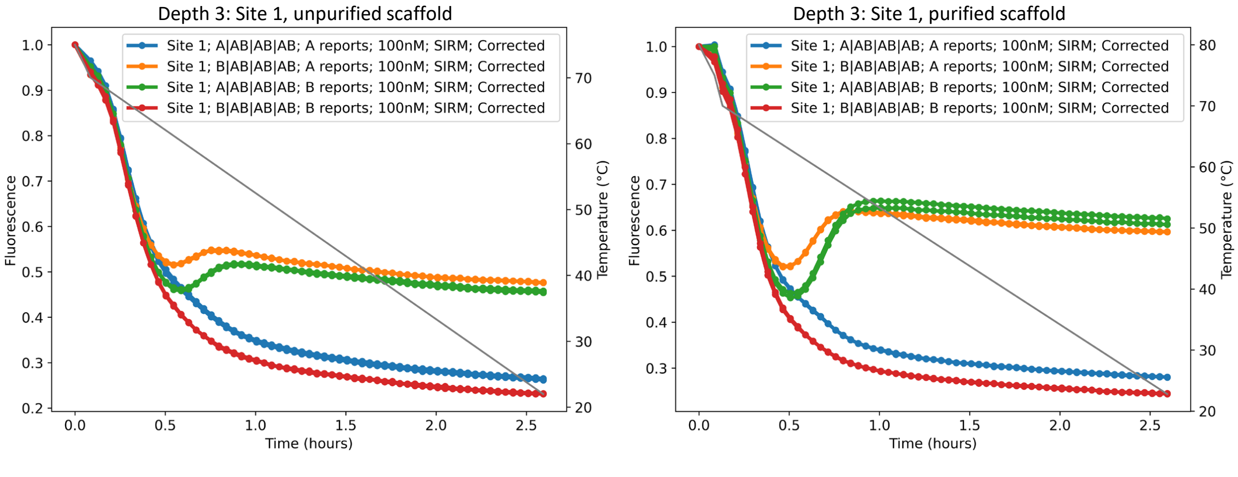

Figure S33: Data for justification of why we chose purified versus unpurified scaffold: specifically the green orange curves on right are higher than left, while red/blue are similar for both. Data from early stages of the project (using manual pipetting and a somewhat slower temperature protocol than reported elsewhere) shows scaffold purification's effect on SDC computation. Left: results from four BitCopy samples using the unpurified scaffold. Right: same, but using the purified scaffold. Samples were pipetted by hand.

#### S7.8 Example SDC mixes

| BITCOPY; (T0, T2) ; Position D ; Input T2 | | | | | |
|---|---|---|---|---|---|
| Species | Stock conc (µM) | Target conc (µM) | Volume to move (nL) | Exact num droplets | Excess |
| scaffold 10uM | 10 | 0.1 | 350 | 14 | 1.00x |
| A2* | 200 | 1 | 175 | 7 | 10.00x |
| 0B0* | 200 | 1 | 175 | 7 | 10.00x |
| 2B2* | 200 | 1 | 175 | 7 | 10.00x |
| 0C0* | 200 | 1 | 175 | 7 | 10.00x |
| 2C2* | 200 | 1 | 175 | 7 | 10.00x |
| 0DQ* | 200 | 1 | 175 | 7 | 10.00x |
| 2D | 200 | 1 | 175 | 7 | 10.00x |
| ATTO*E 10uM | 10 | 0.09285714286 | 325 | 13 | 0.93x |
| 5RF 10uM | 10 | 0.07857142857 | 275 | 11 | 0.79x |
| 3RQ | 100 | 1.785714286 | 625 | 25 | 17.86x |
| 10x Mg++ | N/A | N/A | 3500 | 140 | |
| 0.1x tween | N/A | N/A | 3500 | 140 | |
| 1x TAE | N/A | N/A | 25200 | 1007 | |
| Total | | | 35000 | | |

Figure S34: Example 35$\mu$L mix (Echo 525 compatible) for BitCopy, Copy `1` at Position D.

Figure S34 shows the mix content for BitCopy at scaffold position D with input bit `1`, implemented using compute domains 0 and 2, see SI Section S7.3.2. The experimental results of this mix are given in main text Figure 4a (position D, Copy `1`). Competitive complexity 2 is apparent: there are 2 strands in competition at each scaffold position B,C,D. In this mix format, the use of even/odd compute domain depending on scaffold position is implicit, SI Section S5.1. At position D, the strand `0DQ*` has the extension `Q*` on which the quencher strand can bind, see SI Section S5.2.1. Strands `5RF` and `3RQ` are respectively the fluorophore-modified and quencher-modified strands. Details about buffer, tween and magnesium are given in main text Methods section. A liquid handler (Echo 525, Beckman Coulter) was use to prepare the mix, the column `Exact num droplets` indicates the number of 25nL droplets to use in order to meet the `Volume to move` target. The column `Excess` shows the relative excess of each strand with respect to the scaffold, see SI Section S7.5.2 for excess justification. See SI Section S10 for all DNA sequences.

Figure S35 shows the mix content for Counter "0 to 4". The experimental results of this mix are given in main text Figure 4f. General comments made above about the BitCopy mix apply. Competitive complexity 8 is apparent: there are 8 strands in competition at each scaffold position B,C,D. The way of reporting results described in SI Section S5.2.3 is apparent in this mix: this mix is part of a set of 8 quasi-identical mixes where the only difference is which strands bares the `Q*` domain at position D. Only one of these 8 mixes will quench, which we experimentally demonstrate, Figure 4f.

Mixes like the one depicted in Figure S34 were generated automatically using code, based on their formal description (e.g. `BITCOPY; (T0, T2) ; Position D ; Input T2`). Then, these mixes were converted automatically into Echo 525 *picklists* (i.e. list of atomic mixing instructions) for mixing.

| COUNTER; A1*; 3DQ*; should quench | | | | | |
|---|---|---|---|---|---|
| Species | Stock conc (µM) | Target conc (µM) | Volume to move (nL) | Exact num droplets | Excess |
| scaffold 5uM | 5 | 0.1 | 700 | 28 | 1.00x |
| A1* | 200 | 1 | 175 | 7 | 10.00x |
| 0B1* | 200 | 1 | 175 | 7 | 10.00x |
| 1B2* | 200 | 1 | 175 | 7 | 10.00x |
| 2B3* | 200 | 1 | 175 | 7 | 10.00x |
| 3B4* | 200 | 1 | 175 | 7 | 10.00x |
| 4B5* | 200 | 1 | 175 | 7 | 10.00x |
| 5B6* | 200 | 1 | 175 | 7 | 10.00x |
| 6B7* | 200 | 1 | 175 | 7 | 10.00x |
| 7B0* | 200 | 1 | 175 | 7 | 10.00x |
| 0C1* | 200 | 1 | 175 | 7 | 10.00x |
| 1C2* | 200 | 1 | 175 | 7 | 10.00x |
| 2C3* | 200 | 1 | 175 | 7 | 10.00x |
| 3C4* | 200 | 1 | 175 | 7 | 10.00x |
| 4C5* | 200 | 1 | 175 | 7 | 10.00x |
| 5C6* | 200 | 1 | 175 | 7 | 10.00x |
| 6C7* | 200 | 1 | 175 | 7 | 10.00x |
| 7C0* | 200 | 1 | 175 | 7 | 10.00x |
| 0D | 200 | 1 | 175 | 7 | 10.00x |
| 1D | 200 | 1 | 175 | 7 | 10.00x |
| 2D | 200 | 1 | 175 | 7 | 10.00x |
| 3DQ* | 200 | 1 | 175 | 7 | 10.00x |
| 4D | 200 | 1 | 175 | 7 | 10.00x |
| 5D | 200 | 1 | 175 | 7 | 10.00x |
| 6D | 200 | 1 | 175 | 7 | 10.00x |
| 7D | 200 | 1 | 175 | 7 | 10.00x |
| ATTO*E 10uM | 10 | 0.09285714286 | 325 | 13 | 0.93x |
| 5RF 10uM | 10 | 0.07857142857 | 275 | 11 | 0.79x |
| 3RQ | 100 | 1.785714286 | 625 | 25 | 17.86x |
| 10x Mg++ | N/A | N/A | 3500 | 140 | |
| 0.1x tween | N/A | N/A | 3500 | 140 | |
| 1x TAE | N/A | N/A | 21700 | 868 | |
| Total | | | 35000 | | |

Figure S35: Example 35$\mu$L mix (Echo 525 compatible) for COUNTER, "0 to 4" instance.

### S8 Implementation: Renewable SDC programs

#### S8.1 Prior work

DNA computers are typically one-time use, a significant drawback. Usually, after all the effort in designing and running the computation, the material in the test tube becomes waste. To run a new computation, one must carry out additional experimental steps to remix a new DNA program that costs time, material, and labour.

However, there are exciting alternatives! A number of reusable DNA computing platforms have been developed using approaches such as forcing dehybridization by activation of photo-regulated molecules [51, 147, 148], controlling pH [149, 150], using magnetic beads [151, 152], using enzymes [153–155], switching temperature [52], or adding extra DNA strands [47–50, 156].

**Renewing with the same computation/input as before.** Many of the approaches that achieve high yields renew/reset the program but run the exact same previous computation (input) [52, 149, 150, 156]. Two of those approaches [52, 156] are pure nucleic acid systems (meaning: no DNA modifications, no enzymes, no magnetic beads, no pH change) hence can be compared to SDC renewable programs. For example, the thermal cycling approach [52] demonstrated how to include different motifs (3) in one solution and uniquely activate one at a specific temperature. The authors design logic gates that can be reset by cycling the temperature, thus rerunning the program with the same set of inputs. Output signals require 8 hours to reach completion—somewhat slower than the SDC results presented here.

**Renewing a program with a different input than before.** Some designs [47–50] run new computations by disabling previous inputs and introducing new inputs (which is what we do with SDC renewable programs). However, they tend to accumulate waste in solution. The more sophisticated the circuit is [48, 49], the more complex and constrained renewal becomes, giving a poorer yield of the renewed cycles. The waste from the previous computation severely interferes with the new/renewed computation and decreases the yield of the new output; hence, it progressively degrades over time. The fastest record for each computing cycle (renewing) in such circuits is 15 minutes [48, 156]. The technique called clip-strand [47] also allows for changing of inputs and was used to build a full-adder logic circuit. Each use and reset cycle needs about 12 hours to see the signal flat line, although signal levels show a leak over time.

#### S8.2 Renewable SDC programs: dispensed by acoustic liquid handler

A renewable SDC program is an SDC program such that by adding a new input, and possibly a “small” number of other strands to reset the system, we get a new output for the program. A “small number” could mean a constant, $O(1)$, independent of the number of scaffold positions $N$, for example. For the renewable SDC programs here, we merely add 2 strands for BitCopy, Counter, and Addition.

##### S8.2.1 Renewable Counter, Addition and BitCopy

In the main text (Figure 5), we showed data for three renewed programs using an acoustic liquid handler: BitCopy, Addition and Counter. We used 5.7× excess for the anchor and competing strands, i.e. a bit less than the 10× excess used for Figure 3 and Figure 4, mainly to save on material. The method of renewing used depends on the number of tiles/strands competing at the reporting scaffold position $D$:

- BitCopy: reporting one of 2 results. Input is at the anchor position. In this case, renewing and reporting is straightforward—simply add 2 strands: 1 new input, and 1 old-input-blocker strand – complementary to the previous input. Both strands were added at a concentration ratio 1:1 to the old input; i.e 5.7× relative to the scaffold, see Figure 5b, main text. See SI Section S8.2.2 for details.
- Addition: reporting one of 2 results. (This version of the Addition program has competitive complexity 8 as there are 8 strands competing at $B$, their left compute domains have all $2^3$ 3-bit strings, which in turn allows for programming of 8 different inputs at $A$). In this case, renewing and reporting is the same as BitCopy—simply add 2 strands: 1 new input, and 1 old-input-blocker strand –

complementary to the previous input, see Figure 5b, main text. See SI Section S8.2.2 for details.[29]

- COUNTER: reporting one of 8 results (at $D$). In this case, renewing and reporting is similar to BITCOPY—simply add 2 strands: 1 new input, and 1 old-input-blocker strand – complementary to the previous input, see Figure 5b, main text. But since we wish to report one of 8 possible output values, reporting is a little more complex. There are a few methods to do this. The data for renewed-COUNTER reported in the main text was renewed by adding 2 strands: 1 new input, 1 old-input-blocker strand, and by running 8 samples, each reporting one count out of 8, with a quencher. In Figure S40 we give another method (for experiments done by hand-pipetting).

###### S8.2.2 Design details for renewable BITCOPY and other programs that report one of two results

**Summary/reminder of the BITCOPY program.** The BITCOPY program uses the usual binary reporting method (Section S5.2.1) where the output is either bit-0 or bit–1. Output bit-`0` happens when the input at the anchor position $A$ copies bit-`0` compute domains until the last position ($D$) that ends with a quenching domain complementary to the quenching strand. This brings the quencher besides the fluorophore and starts the quenching process and produces a low signal. The opposite situation applies when copying bit-`1`. When input at position $A$ copies bit-`1` through the compute domains, it reaches the last position that doesn't have a domain complementary to the quenching strand. In this case, the fluorophore remains free of quenching and reports a high signal representing bit-`1`.

**Details of the renewed BITCOPY program.** BITCOPY has identical sets of two strands at each scaffold position $B, C$, and $D$. In each set, one of the strands copies bit-`0` and the other copies bit-`1`. In the reported result in Figure 5d in the main text, we used the compute domains pair (CD1, CD3) (names CD$i$ defined in Section S6.2, DNA sequences in Section S10), where CD1 copies signal bit-`0` and CD3 copies signal bit-`1`. Figure S36(a) illustrates the strand sets used in the experiment.

###### S8.2.3 Experiment design for renewable COUNTER: competitive complexity 8 programs using multiple samples

Results of programs with competitive complexity 8 are reported by quenching as detailed in Section S5.2.3—i.e. with only two signals it is not possible to report 7 structures out of 8 in the same sample (test tube). We decided to run 8 samples where each of them would quench at 1 of the 8 possible results. We trigger a renew in each sample as usual with two strands, 1 new input, and 1 old-input-blocker strand – complementary to the previous input in the sample – all are added with a concentration ratio 1:1 to the old input; i.e 5.7× relative concentration to the scaffold. From 8 resulting curves in a renew cycle, only the one sample, that is targeted by the added input, shows quenching while the other 7 curves show no quenching.

###### S8.2.4 Renewable COUNTER: analysis

Figure S37 shows the final levels of each of the 8 renewed COUNTER instances shown in the main text Figure 5e. Each instance is renewed a total of three times (each of the three times is separated by 7 other instances renewals in between), hence three points per plot. We see that the quenching signal gets linearly higher, i.e. it degrades. We compute an average of 17.3% (SD=0.067) signal degradation every 8 renewals over these 8 COUNTER instances: this metric is computed by averaging $\frac{\text{final\_level\_1}-\text{final\_level\_0}}{\text{final\_level\_0}}$ and $\frac{\text{final\_level\_2}-\text{final\_level\_1}}{\text{final\_level\_1}}$ over the 8 COUNTER instances.

##### S8.3 Renewable SDC programs: dispensed by hand-pipetting

The work reported thus earlier in Section S8.2, and in Figure 5 of the main text, used a (rather expensive) acoustic liquid handler (Echo 525, Beckman Coulter, supplied by Labplan Ltd., Ireland). In this section, we

[29]We note that, although not pursued here, there is another way to do ADDITION (see Section S4.2) that encodes input and program bits on two separate tile sets, which in turn means that any input can be renewed without renewing any program bits. We did not demonstrate this experimentally, but it is straightforward to implement with our existing tile set, since they encode all 3-bit compute domains.

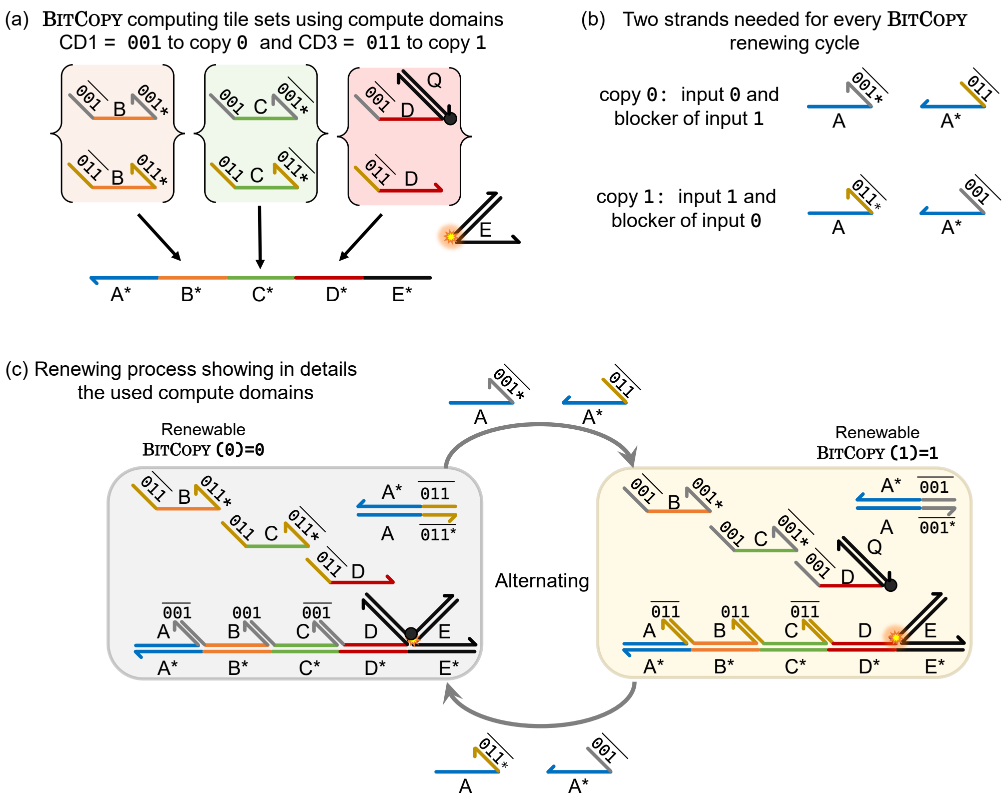

Figure S36: Renewable BitCopy strand design. (a) Strands for standard the BitCopy program, as used in Figure 5c of the main text, showing two strands at each of scaffold positions $B, C, D$, and emphasising which two domains we chose to represent bit-0 and bit-1. (b) Two strands are sufficient to renew the BitCopy program, one to block the old input and the other to encode the new input. (c) Detailed strand diagram of the renewing process, alternating between copying bit-0 and bit-1

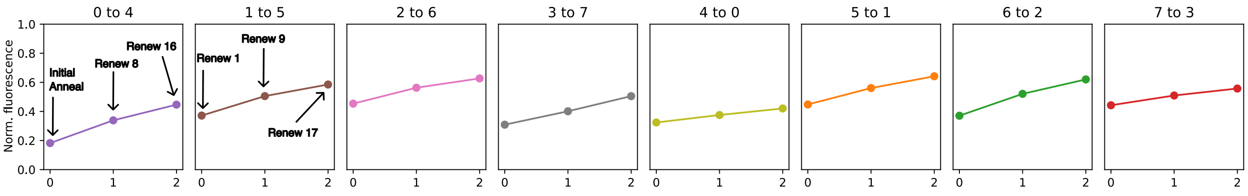

Figure S37: Final normalised levels of each of the 8 renewed Counter instances showed in main text Figure 5e.

show that SDC programs can also be renewed when pipetting by hand, i.e. no need for expensive equipment.

Our main conclusion will be that this (merely) comes at a cost of extra dilution over time, since our hand pipettors require larger minimum volumes for consistency (e.g. we hand-pipette 2 $\mu$L) than our acoustic liquid handler (e.g. we dispensed 0.2 $\mu$L per renew, but could in theory go as low as 25 nl on the device).

##### S8.3.1 Pipetting by hand: Dilution lowers signal for renewable programs

Figure S39 shows the effect of repeatedly adding a small volume (2 $\mu$L) of solution on the fluorescence signal.

**Observations and conclusions from Figure S39:**

- The most striking observation is that although the Copy 1 experiment signal (orange curve) reduces over time, a significant fraction of that reduction is merely due to dilution, as seen in the blue curve (control).

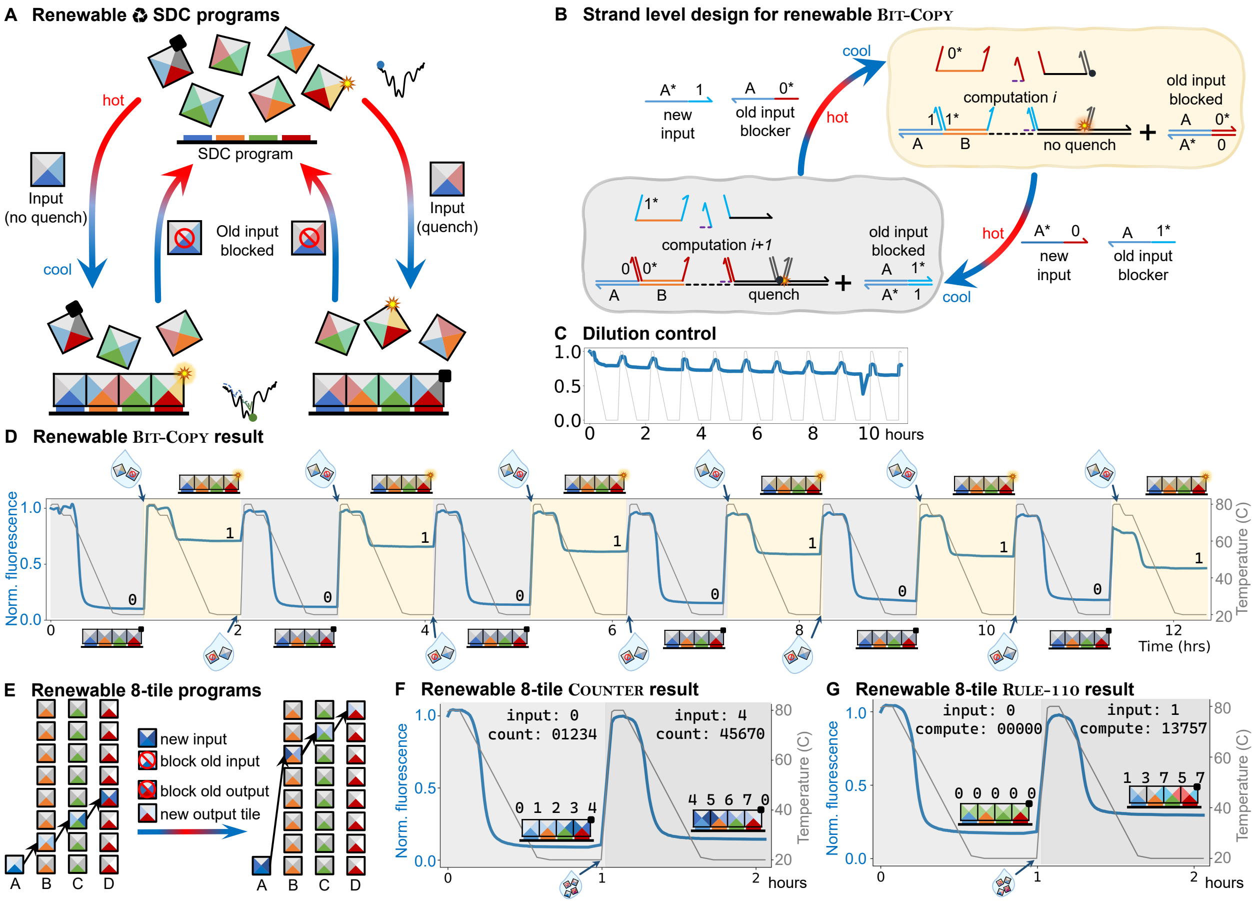

Figure S38: **Renewing SDC programs by hand-pipetting (no automated liquid handler).** **(A)** Principles for renewable SDC programs: at each renew cycle we add a new input, and a blocker for the old input. **(B)** Strand-level diagram of renewable BitCopy. At the beginning of each temperature cycle, the old input gets deactivated by its complement and the new input is available to copy the desired bit. **(C)** Effect of serial dilution on SDC signal in the absence of computation. **(D)** Renewable BitCopy: twelve runs cycling between input bits 0 and 1. Part of the decrease of the high output signal is due to serial dilution shown in **C**. **(E)** Renewing of more complex 8-tile programs, an example of a random program shows deactivation of old input and old output tiles and introduction of new input and output tiles. Otherwise, the rest of the tile set remains the same. The correct computation is reported with quenching. **(F)** Renewable 8-tile Counter experiment: two runs of the counter: Hour 0: Started with counter value 0 (counting $0 \to 1 \to 2 \to 3 \to 4$), Hour 1: renew with input 4 (counting $4 \to 5 \to 6 \to 7 \to 0$). **(G)** Renewable Rule110: Hour 0: input 0 (cycle $0 \to 0 \to 0 \to 0 \to 0$), Hour 1: renew withn input 1, (cycle $1 \to 3 \to 7 \to 5 \to 7$). Samples were pipetted by hand.

- In Copy 1, the drop in signal (from its level at 80 °C to that at 20 °C) is approximately equal for the renewing cycle at hour 2, as it is for hour 11, suggesting the system is reasonably close to the intended behaviour. However, there is some variance in the Copy 1 signal at 20 °C, which is not explained (could be qPCR machine variance, or variance due to manual handling as the plate was spun just before going to 80 °C between each cycle — to try to reduce such variance! Note that the plate was not centrifuged in the data reported in Figure S38).
- Another observation is that the Copy 0 signal increases slightly over time. It is not clear why, perhaps a changing of equilibrium, at 20 °C with the addition of unpurified blocker and 0 and 1 input strands over time.
- Besides respective downward and upward trends, there is some unintended variance in copy 1 and 0, respectively.
  - There is an obvious spike around hour 9 which can be ignored as it was an experimental error:

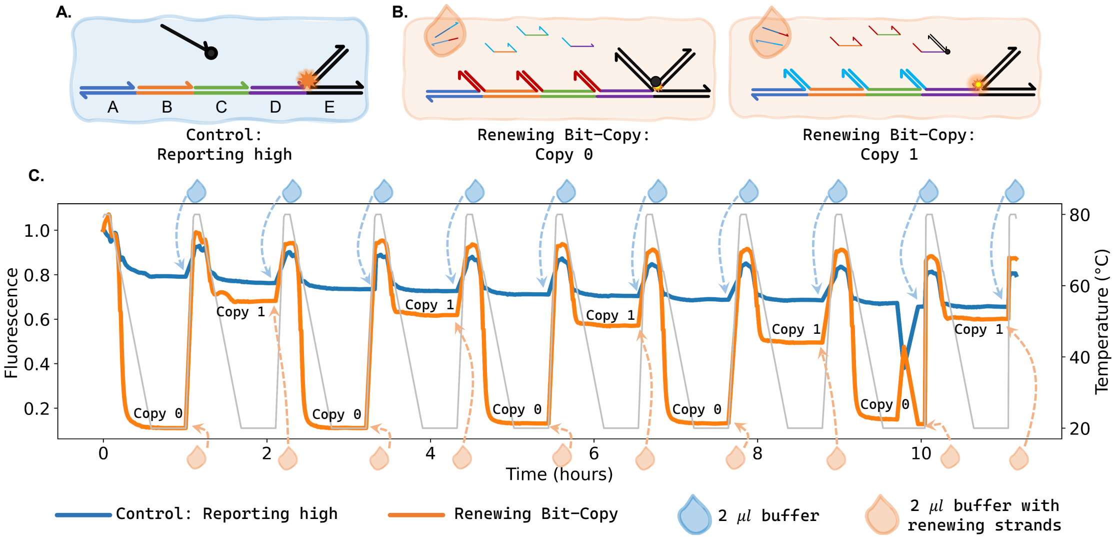

Figure S39: Data showing that dilution by hand-pipetting, by 2 $\mu$L at each renewing cycle, significantly lowers the signal in renewing experiments. (A) Dilution control: strands form a complex that reports high signal, reported by the blue curve in C (i.e. no strand competition at scaffold positions). (B) Renewable BitCopy experiment: strands, depicted as target configurations, for the Copy 0 and Copy 1 experiments reported in the orange curve in the plot in C. (C) Fluorescence trace in orange shows BitCopy with addition of 2 $\mu$L of solution (with new input and old input blocker) at each step. Fluorescence trace in blue shows that adding 2 $\mu$L of buffer lowers the signal at each renewing cycle. The data is normalised as follows: the first datapoint on a curve is divided into every other datapoint along the curve (data is not rescaled to non-renewing controls as was done elsewhere). Data taken at the following rate: Hold at 80 °C for 5 mins taking one datapoint per min. Then during the temperature drop of 1 °C per 0.5 min from 80 to 20 °C take 1 datapoint per 0.5 min, and finally one datapoint per 5 min during the temperature hold at 20 °C. The plate was centrifuged between each cycle in an effort to reduce signal variance due to unintended bubbles/droplets after adding strands (such centrifugation between cycles was *not* done in the experiment shown in Figure S38). Samples were pipetted by hand.

the qPCR machine door was opened too early, causing the machine to take a datapoint with no sample present.

– There is a systematic reduction of the signal at 80 °C, presumably due to dilution as it happens in both control and experiment. Although being systematic suggests it is reasonable to renormalise the data each time we reset to 80 °C, we decided to leave the data relatively unprocessed.

##### S8.3.2 Experiment design for renewable Counter, Rule110, and any competitive complexity $> 2$ programs using one sample and 4 strands

The programs Counter and Rule110 have the property that at each position, the same set of tiles compete (except that the position domains are unique to the position). Also, they both have competitive complexity 8: each non-anchor scaffold position has 8 competing tiles. Both are deterministic: depending on the unique input tile at the anchor position $A$, a single output number (Counter) or bit (Rule110) is reported at position $D$.

To report the correct output signal, we used the method described in S5.2.3 where we place the quenching domain only on the desired output strand at the last position, $D$. The other 7 strands at position $D$ have no 3′-end reporting/compute domain (the 3′-end of a compute strand is on the right-hand side in our drawings).

Figure S40(a) gives the computing tile set of Counter to run two cycles: the first receives input value 0 and counts to 4, counting the sequence $0 \to 1 \to 2 \to 3 \to 4$, and is then renewed receiving input 4 and counts to 0, counting the sequence $4 \to 5 \to 6 \to 7 \to 0$.

To initiate a renew cycle, we add 4 strands (all strands added at the usual compute strand excess of 10×):One strand that blocks (binds to) the previous input at position $A$, as well as a new input strand. Since we are reporting one of many outputs, we use a strategy of adding two further strands to block the

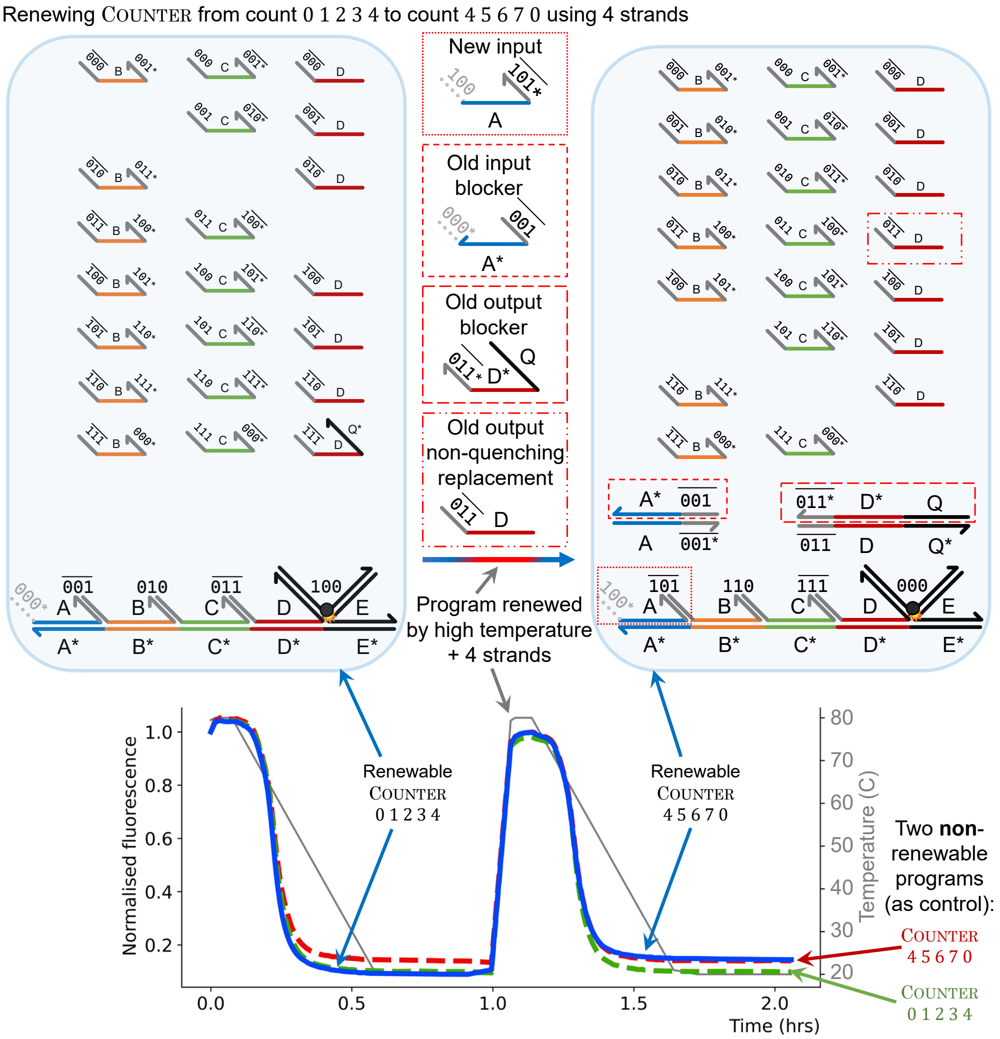

Figure S40: Renewing programs with competitive complexity $> 2$, example of COUNTER. Top shows the result of renewable COUNTER to run two cycles, on the left $0 \to 1 \to 2 \to 3 \to 4$ and on the right $4 \to 5 \to 6 \to 7 \to 0$. In between the two strand diagrams are the 4 needed strands for the renewing process. Bottom: result of three experiments. Blue curve: renewable COUNTER showing two cycles. Red curve (non-renewing COUNTER-program control): COUNTER running $0 \to 1 \to 2 \to 3 \to 4$, designed to produce the same target structure as the blue curve on its first cycle. Green curve (non-renewing COUNTER-program control): COUNTER running $4 \to 5 \to 6 \to 7 \to 0$, designed to produce the same target structure as the blue curve on its second cycle. Samples were pipetted by hand.

previous output and report the new output.[30] Figure S40 shows a COUNTER instance with input 0 ending with count 4, then 4 strands are added to renew it to start with input 4 and end with count 0.

[30] For renewing the output, ideally, we could use 4 strands: 1 strand to block the old reporting (quenching) output strand $x$, 1 strand that is the same as $x$ but no quenching domain, 1 new reporting output $y$ with quenching domain, 1 strand to block the old tile with no quenching domain that $y$ replaces. However, previous results showed that even with larger competitive complexity (8 in this case), the input only reports its correct output (see main text COUNTER and RULE110 results in Figure 4f,g). Therefore, we decided that for simplicity and to reduce the number of strands, to run two renewing cycles and make the quenching strands of both cycles available from the beginning. We ran the two computations independently to act as controls that prove that the reporting strand of the second cycle doesn't interfere with the reporting of the first cycle. Therefore, at the beginning of cycle 2, while adding the renewing strands, we didn't need to add the new cycle quenching strand and block the previous cycle non-quenching strand of the new output.

### S9 Implementation: Scaling-up SDC

#### S9.1 DNA sequence design for scale-up

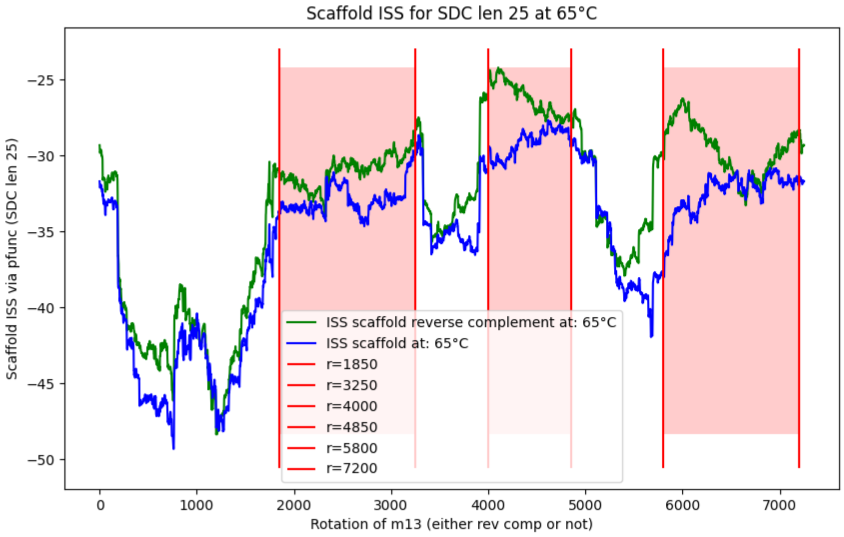

Figure S41: Internal Secondary Structure (ISS) of the first 624 bases (and reverse complement) of each of the 7249 rotations of the M13 bacteriophage. ISS is computed using NUPACK4's pfunc function at 53 °C [57].

SDC scale-up to scaffold length $> 4$ (or $> 5$, including reporters) for experiments in Figure 6 of the main text use subregions of 7249-base M13 bacteriophage: in the experiments the full 7.2kb M13 strand is present in solution (at 10 nM), but only a subregion is targeted, serving as scaffold for a scaled-up SDC. This contrasts with our use of a *synthetic 120-base scaffold* (at 100 nM) in the 5-domain setup, see Section S10.1.1.

The main design challenge for SDC scale-ups was to choose which subregion of M13 to use. Two regions were chosen: (i) a 288-base subregion (for length 11 BitCopy, discussed in Section S9.3) which is essentially the continuation of our 120-base synthetic scaffold (uses two an M13 subsequences, see Section S10.1.1) that we used for backward-compatibility, and (ii) a 624-base subregion, for programs using SDC length 20 to 25 (Sections S9.5 and S9.4), which we decided to choose from scratch.

Choosing this 624-base, 26-domain M13 subregion was challenging because M13 is known to have regions with internal secondary structure (ISS). Figure S41 gives a metric of ISS, computed using NUPACK4's pfunc function, of the first 624 bases of each of the 7,249 rotations of M13 (in blue), and of its reverse complement (in green). We see that ISS varies greatly, in a -25 to -50 kcal/mol range. The criteria for choosing the region were, from most to least important:

1. **Low ISS.** In order to avoid scaffold self-binding, candidate M13 subregions were restrained to three sections of M13 exhibiting low ISS, highlighted in red in Figure S41.
2. **Domain isoenergeticness.** In order to have a uniform energetic landscape across 24-base scaffold domains, candidate subregions were ranked by lowest standard deviation amongst domains' Watson-Crick binding, as computed using the nearest-neighbour model with parameters taken from SantaLucia and Hicks [60, 132]).
3. **Low scaffold/compute domain interaction.** In order to avoid cross-talk between scaffold and compute domains, we sought to minimise their binding as computed using NUPACK4's pfunc[57].

Other criteria such as the ability to tweak the subregions' energy landscape with compute strand mismatches, or simulated performance on length-25 IsoenergeticBitCopy using a sequence-dependent kinetic simulator [157], were taken into consideration. Finally, we hand-picked a 624-base subregion of M13 which performed relatively well on all these metrics.

See main text Methods section *Data and code availability* for accessing the code that computed these

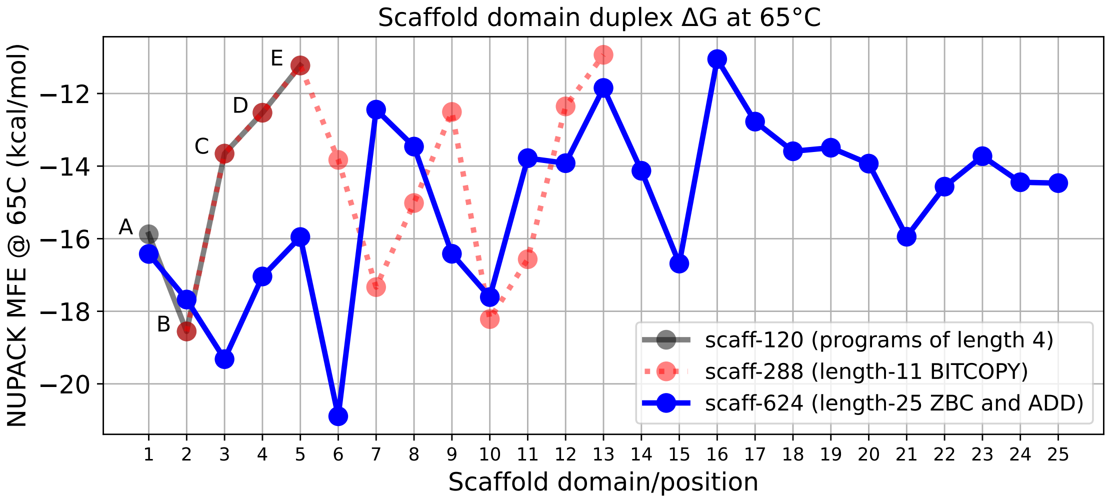

Figure S42: Blue: Domain energetics of our chosen 624-base M13 scaffold sequence scaff-624. In experiments we used the full 7.2 kb M13, of which scaff-624 is a segment. Black: short scaffold (scaff-120, synthetically synthesised 120 base strand). Red: scaff-624, a segment sitting on the full 7.2 kb M13 that includes 4 domains from scaff-120. Domains are of length 24 bases each; the plot shows the `NUPACK4` MFE [57] of the duplex of a 24-base domain and its perfect reverse complement at 65°C. The final reporting position (26) is not shown.

metrics. DNA sequences are in Sections S10.2 to **??**.

#### S9.2 Yield analysis of scaled-up programs

We define *yield* for a sample in this context to be the fractional position of the ending fluorescence of a sample between the ending fluorescence of the correct and incorrect controls for the sample's position, that is, given $p = (e_{\text{sample}} - e_{\text{low}})/(e_{\text{high}} - e_{\text{low}})$, the yield for a sample is $y = p$ if the sample's target output is high, and $y = 1 - p$ if the sample's target output is low. Intuitively, if the ending fluorescence of the high/low controls is taken to be the fluorescence at the position when no/all scaffolds have a quencher strand attached, this yield can be roughly interpreted as the percentage of scaffold strands in the sample that have the correct output. With this definition, for either high or low target states, a 100% yield would represent every scaffold being correct, a yield far higher than necessary to distinguish the output of the computation, while a yield of 50%, for a binary output, would represent a complete failure, with both outputs equally likely. When replicates are present, for calculation of yield, the sample ending fluorescence is averaged across replicates, while the minimum fluorescence (maximum reporter binding and quenching) of (two) control replicates is used.

#### S9.3 Scaling-up: length 11 BitCopy

For an initial scale-up step, we used scaffold scaff-288, which is a segment of 7,249-base M13 that extends the sequence of scaff-120 from 120 bases to 288 bases. We first ran an 11-position BitCopy program on scaff-288. 3-hour anneals showed limited success, but better results were seen on 24-hour anneals, see Figure S43.

Major changes relative to shorter 4-domain systems (or 5, including reporters) included:

- There were new 'random biological' scaffold domains, i.e. the new domains could make the system even worse because they are not chosen/designed to have good binding energetics
- Length-4 systems used a synthetic 120 base scaffold, however scaff-288 is a segment of full-length single-stranded M13, meaning that there are 6,961 single-stranded DNA bases adjacent to scaff-288 in solution.
- Experiments were carried out at 10-fold lower concentrations (100-fold decrease was also tested).
- Reporter-binding strands were unpurified.

However, this work helped us discover new principles applied in later sections below.

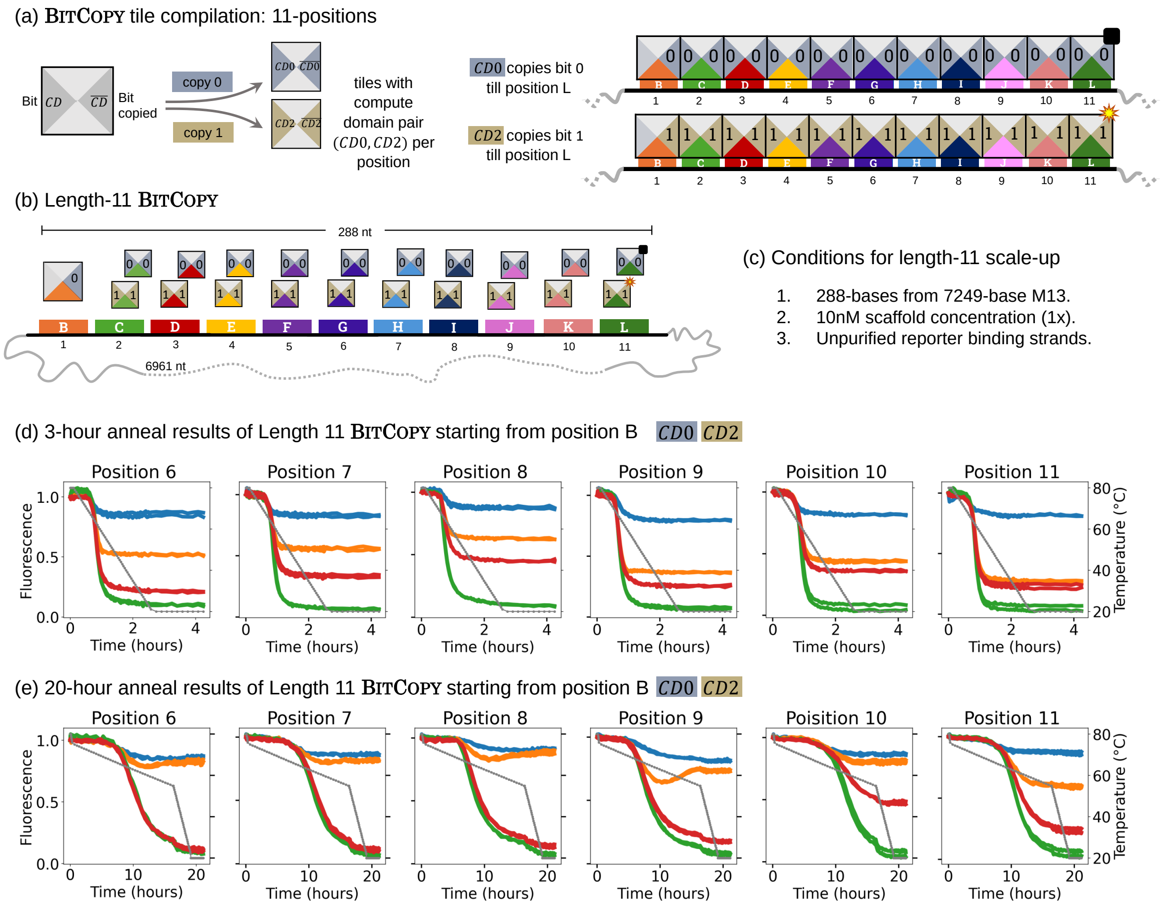

Figure S43: (a) BitCopy tiles, each containing a compute domain pair (CD0 or CD2) to propagate bit 0 or bit 1. (b) The BitCopy program is scaled up to an 11-position scaffold termed scaff-288 that incorporates positions B, C, D and E from scaff-120 and extends 24-bases to the right. Scaffold positions on the scaff-288 are labelled from B to L (Reporting position 12 is not shown in the tile abstraction). (c) Experimental conditions for length-11 scale-up: 288-base subregion of the 7249-nt M13 scaffold, 10 nM scaffold concentration (1×), and unpurified reporter-binding strands. (d) Typical 3-hour anneal normalised results for length-11 BitCopy for positions 6 to 11. (e) 24-hour anneal results for length-11 BitCopy for positions 6 to 11.

##### S9.3.1 DNA base mismatches

Using a biological M13 scaffold limits the bounds of scaffold manipulation: scaffold position sequences, and hence, their sequence-complement binding energetics, cannot be changed. Yet we hypothesise that uneven scaffold domain binding energies, with some positions that are significantly stronger, may interfere with computation by allowing uncontrolled attachments at higher temperatures in positions with more favourable scaffold domains. Fortunately, the inclusion of base-pair mismatches between scaffold and strand sequences could allow for an increase in $\Delta G$ (a weaker bond) for strand-scaffold binding without changing the M13 scaffold sequence [132]. By incorporating mismatches at specific positions, the energy landscape can be “engineered”, weakening overly strong scaffold positions and improving computation, as detailed in Figure S44. The use of mismatches in scaff-288 provided a means to improve computation when working on extended scaffold and was an inspiration for its use in the 25-position system in S9.5. Mismatched bases were chosen using `NUPACK4`’s MFE. In Figure S44, experiments with base-pair mismatches at positions 5, 6, 7, and 8 showed improved computation using the standard anneal protocol, with dependence on the mismatch variation.

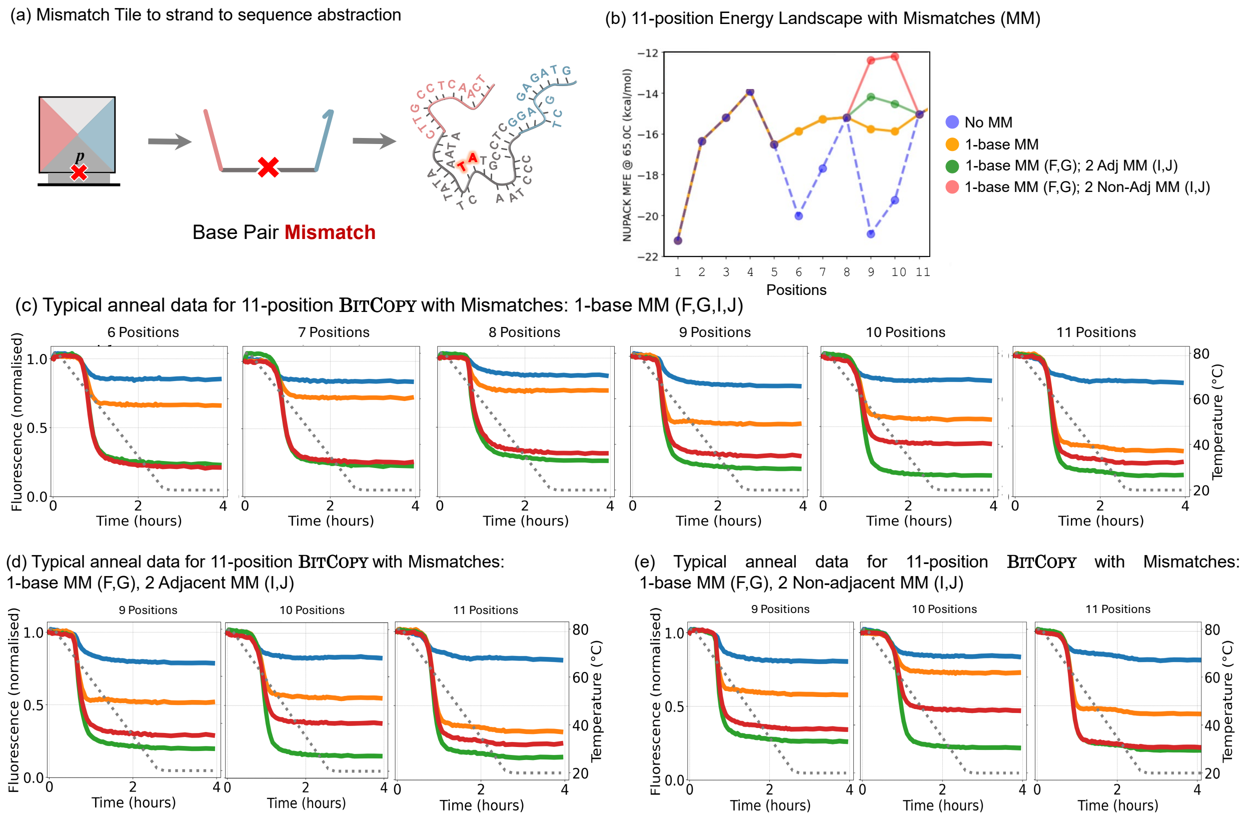

Figure S44: (a) Tile, strand and sequence abstraction of a Mismatch tile involved in BitCopy. Mismatch tiles used in scale-up have single or double mismatches. (b) Domain energetics of the 288-base M13 scaff-288 (blue) and projected domain energetics with mismatches implemented on positions F, G, I and J.(c) 11-position BitCopy normalised anneal data at typical anneal length with three mismatch variations. One base mismatches (MM) on positions F, G, I and J. (d) One base mismatch on positions F and G and two base adjacent mismatches on positions I and J. (e) One base mismatch on positions F and G and two base non-adjacent mismatches on positions I and J.

#### S9.4 Scaling-up: length 25 Addition

The Addition program has a logical property with both thermodynamic and kinetic implications: carry-propagation. Whether a carry bit must move through a given position (*non-sink*) or gets absorbed (*sink*) determines the amount of tile competition that occurs at that position. To compare: in a BitCopy system, two tiles compete at each position. In an Addition sink position, however, there are two tiles that can bind, and independent of whether or not a carry passes through, only one of those two has a matching neighbouring tile. Hence logical errors can not propagate through a sink. This in turn gives a measure of how hard the computation is: fewer sinks means a harder computation. Figure S45(a) shows the Addition computing tiles at position $i$. Figure S45(b) shows a sink and non-sink positions. At a sink position $i$ where two identical bits are added, either `00` or `11`, the carry output is the same regardless of the incoming carry. At a non-sink position $i$, where two different bits are added, either `01` or `10`, the carry output depends on the incoming carry value.

We tested 25-bit ADDITION using a range of regions of maximum non-sink run-lengths, $M$, to assess computation hardness. Higher $M$ values correspond to harder problems due to extended carry propagation.

All tests used the scaff-624 region of the M13 scaffold. Thermodynamic properties of this region are shown in Figure S46(a), and its sequence choice is detailed in Section S9.1.

Figure S46(b) shows the frequency distribution of $M$ values computed across one million 25-bit input pairs, randomly selected from the uniform distribution on inputs. Most inputs (over 74%) have $M \leq 3$, 99% have $M \leq 6$. This informed our experimental input selection: we chose representative input pairs for common ($M = 3$ to $M = 6$) and edge cases ($M = 0$, $M = 25$) to span the full difficulty range. Extreme

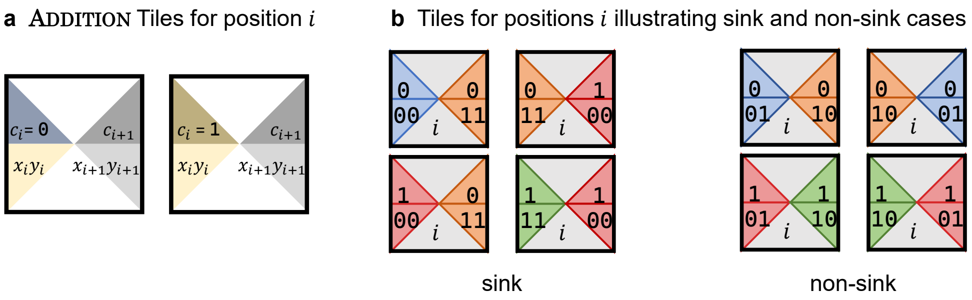

Figure S45: Explanation of sink, no-sink positions in SDC showcasing the ADDITION program. (a) ADDITION tiles at position $i$ encoding the computation. (b) sink positions happen when adding two similar bits, the resulting carry is the same despite the value of the incoming carry. Oppositely, no-sink positions happen when adding two different bits, the resulting carry depends on the incoming carry.

cases like $M = 25$ are rare. This confirms that, in practice, most 25-bit additions do not require long carry chains, which in turn helps with SDC kinetics. The full statistical breakdown is shown in Table 3.

| Max run ($M$) | Count | % | Cumulative % |
|---|---|---|---|
| 0 | 0 | 0.0000 | 0.0000 |
| 1 | 22,228 | 2.2228 | 2.2228 |
| 2 | 350,299 | 35.0299 | 37.2527 |
| 3 | 373,231 | 37.3231 | 74.5758 |
| 4 | 167,793 | 16.7793 | 91.3551 |
| 5 | 58,395 | 5.8395 | 97.1946 |
| 6 | 19,083 | 1.9083 | 99.1029 |
| 7 | 6,145 | 0.6145 | 99.7174 |
| 8 | 1,949 | 0.1949 | 99.9123 |
| 9 | 600 | 0.0600 | 99.9723 |
| 10 | 196 | 0.0196 | 99.9919 |
| 11 | 60 | 0.0060 | 99.9979 |
| 12 | 18 | 0.0018 | 99.9997 |
| 13 | 2 | 0.0002 | 99.9999 |

Table 3: Distribution of maximum run-lengths of non-sinks ($M$) across 1 million random 25-bit input pairs.

We monitored the computations' progression across different annealing protocols with lengths of 30 minutes, 1 hour, 5 hours, 14 hours, and 24 hours. The goal was to observe how $M$ impacts the kinetics and final yield. In Figure S46, panels c-j show the ADDITION results of different randomly selected 25-bit input pairs with $M = 0, 3, 4, 5, 6,$ and 25. In Figure S46:

- (c) shows the most ideal case is shown where $M = 0$ and all positions are sinks. Computation completes rapidly. Even the 30-minute anneal shows near-complete output. This confirms that in the absence of carry propagation, computation can be fast.
- (d) shows $M = 3$, where short non-sink regions are introduced due to the nature of the computation. Annealing 1 to 5 hours yields nearly complete results judging by the distance from the control.
- (e) shows $M = 4$, which introduced longer carry propagation that increased the hardness of the computation in the non-sink regions. Errors started to appear near the end of the longest non-sink stretch between positions $E$ and $I$. However, longer anneals from 5 to 14 hours helped to bring the output to near completion.
- (f) shows $M = 5$, where the competition between tiles in non-sink regions becomes more visible. Partial computation is visible even with the 5-hour anneal. However, the 14 and 24 hours improve the output completion level.
- (g) and (h) show $M = 6$ examples. The regions of 6 consecutive non-sink positions are later on the scaffold: in (g) between positions 12 and 19, and in (h) between positions 16 and 22. In these experiments, error states persist throughout the non-sink run. Long anneals of 14 and 24 hours are

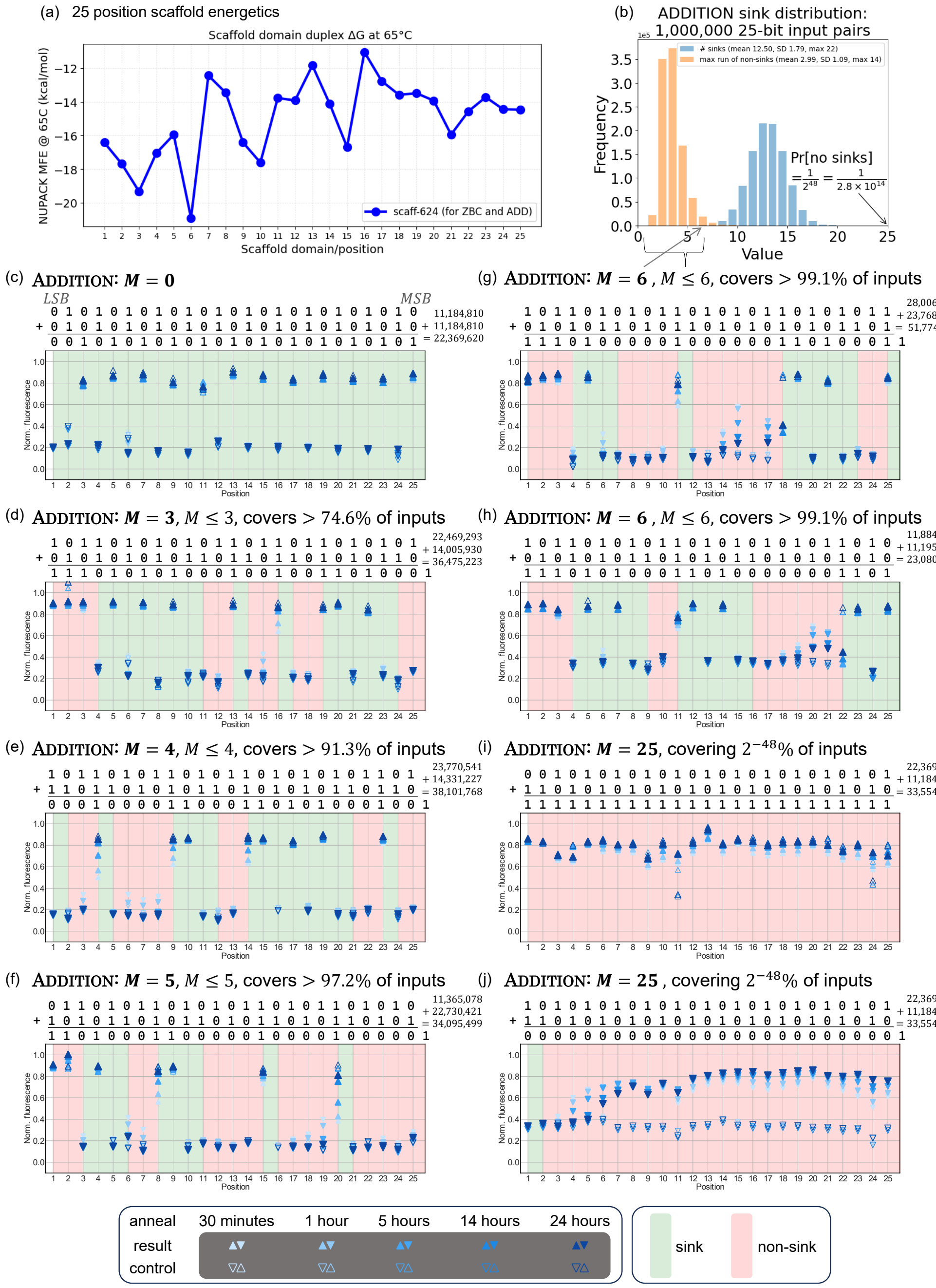

Figure S46: Scaling-up to Addition to $2 \times 25$-bit input, with 25 output bits and $\leq 25$ carries (by some definitions, 100-bit computation). (a) Energetics of the scaff-624 position domains showing some very strong domains like position 6 followed by position 3, and position 15. (b) Frequency distribution of maximum length runs $M$ over 1 million 25-bit input pairs. (c) Results of $11,184,810 + 11,184,810 = 22,369,620$ that has $M = 0$. The most optimum result where all positions reach completion even at fast anneals like 30 minutes. (d-h) different Addition computations with $M$ starting from $M = 3$ to $M = 6$ showing that the longer the no-sink region, the slower the system is to reach completion. (i-j) $M = 25$ Addition computation resulting in all `1`'s in (i) and all `0`'s in (j) using the same tile set and only changing the anchor at position 1 to propagate a carry of `0`'s in (i) and of `1`'s in (j).

required for almost full propagation of the correct output. However, in the last position of each $M = 6$ region in the two computations (position 19 in (g) and 22 in (h)), while longer anneals improve performance, completion levels remain far from target controls.

- (i) and (j) show two $M = 25$ examples. In these computations, the system was pushed to its maximum limit, with no sinks anywhere on any position. These are the worst-case inputs: the computation resembles that of BitCopy, and similar behaviour can be expected. In (i), the output pattern may appear to report the correct result of all `1`'s. However, this apparent completion is not genuine. In (j), we reprogrammed the computation by using the same tile set and changing the anchor tile, resulting in a computation of similar difficulty that instead should generate all `0`'s, as seen for the controls. Yet the result in (j) shows that the computation fails after 6 positions with long anneals. Position 6 is the strongest position on the scaffold, as seen in Figure S46(a). So, in this configuration, once an incorrect `1`-tile attaches at a given position, especially with no sinks to reset or constrain the configuration, it becomes difficult to replace. The error then propagates through the rest of the scaffold, giving the false impression of successful computation in panel (i). This contrast in the behaviour between (i) and (j) while using the same computing tiles set, suggests that the `1`-reporting tiles are thermodynamically favoured in this configuration.

The data in Figure S46(c-j) show that sink positions function as convergence checkpoints that stabilize the intermediate configurations and accelerate progression. As the maximum non-sink run length $M$ increases, the system becomes slower with more frequent errors and reduced final yields. Most practical input pairs, as indicated in Figure S46(b) and Table 3, fall within $M \leq 4$, a system where computation proceeds reliably within reasonable annealing times. Oppositely, inputs with $M \geq 6$ experience kinetic delays and need extended annealing to achieve near-completion levels. These observations support the use of $M$ as a reliable predictor of Addition program difficulty and runtime. To address the limitation imposed by high $M$, we developed the IsoenergeticBitCopy design described in section S9.5. The design strategy aims to equalize the thermodynamic unavoidability of different configurations, reducing the likelihood that incorrect tiles gain kinetic or energetic advantage. By making intended and error states isoenergetic, the system minimizes bias and improves robustness, specially in computations with extended non-sink segments.

#### S9.5 Scaling-up: length 25 IsoenergeticBitCopy

The 25-position 75-bit computer operates on 26 continuous 24-base scaffold domains, denoted as A to Z, using a carefully chosen 624-base rotation out of the 7,249 nt M13 DNA sequence, scaff-624, as detailed in S9 and Figure S42 for full system reporting. In IsoenergeticBitCopy, the anchor strand binds at position A, and positions B to Z are involved in computation and reporting programs of scaffold length less than or equal to 25. IsoenergeticBitCopy is a model program, similar to BitCopy, involving bit-flipping and the implementation of NOT gates at each scaffold position. This bit-flip behaviour optimises system energetics by choosing two CDs (CD1 and CD0) with isoenergetic binding behaviour and minimises any energetic bias between BitCopy-specific COPY 0 and 1 polymers. Using CD1 and CD0, the pool of compute strands for IsoenergeticBitCopy when input = 0 can be denoted as A0, 0B1, 1C0, etc., when input = 1, as A1, 1B0, 0C1, etc.. Since the number of times each compute domain is used is equal regardless of computation, the correct configurations are isoenergetic (indeed, erroneous configurations also inherit an isoenergetic property: for fixed $k$, all $k$-mismatch configurations are roughly isoenergetic). Figure S47(a) details a strand and tile abstraction of the IsoenergeticBitCopy program, and in Figure S47(b) an example of IsoenergeticBitCopy program is detailed.

In the 3-hour anneal shown in Figure S47(c), kinetically trapped structures are hypothesised to contribute to higher computational error rates, as indicated by the distance of output signals to control levels. Standard annealing durations in IsoenergeticBitCopy have difficulty achieving the target configuration at longer length scaffolds. This suggests that many compute strands may be incorrectly or partially bound to the scaffold, disrupting correct strand alignment and signal output. In contrast, the 8-hour and 14-hour annealing protocols exhibit greater proximity between output and control signals, indicating that extended annealing at elevated temperatures facilitates error correction and enhances the likelihood of reaching the correct configuration. Based on analysis from SDC simulation software, CD0 and CD1 were chosen for IsoenergeticBitCopy as they had isoenergetic hybridisation to their complements that would provide optimal results for a large-scale SDC system.

Throughout the IsoenergeticBitCopy scale-up process, SDC simulations sought to optimise experimental results using mismatches, as we did with 11-position BitCopy (Figure S48).

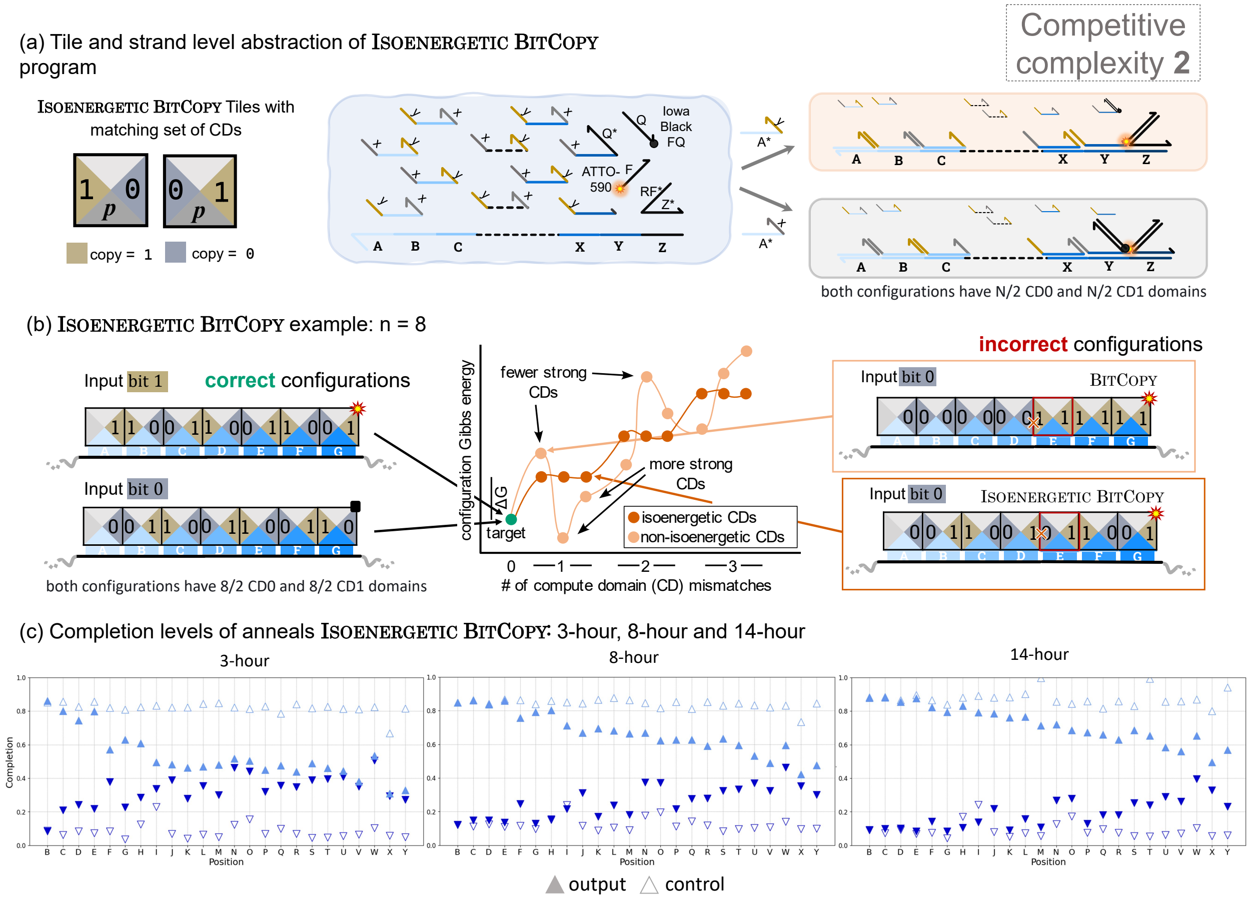

Figure S47: (a) Tile and strand-level abstraction of the IsoenergeticBitCopy program. Correct configurations have a balanced distribution of compute domains (CD0 and CD1), ensuring minimal energetic bias for bit propagation. (b) Example of an IsoenergeticBitCopy system with n=8. Both correct configurations contain equal numbers of CD0 and CD1 domains. Incorrect configurations for normal BitCopy and IsoenergeticBitCopy are illustrated based on input bit states. Alternation of CD0 and CD1 ensures all configurations with $k$ compute domain mismatches to sit on a roughly flat, staircase-like energy plateau with $\Delta G$ of $O(k)$. (c) Completion levels of IsoenergeticBitCopy anneals at 3-hour, 8-hour, and 14-hour anneals.

Data from 11-position scaff-288 indicates that implementing base-pair mismatches increases the $\Delta G$ associated with strand-scaffold hybridisation [132], which led to improved computation (Section S9.3.1). Using an in-house produced python library **(sculptor)**, mismatch base-pairs were chosen to create an elevated energy profile (black plot in Figure S48(b)) with increasing $\Delta G$ to facilitate error correction at downstream positions. We hypothesise that compute strands would bind initially at earlier (and stronger) positions, propagating a correct signal across the scaffold. As the strands bind at later positions, it becomes easier to displace less stable, higher-energy strands.

We measured the efficiency using *performance ratio* (PR), which measured system efficiency between the computations ($COPY$) and controls ($CTRL$) using the completion fluorescence values obtained from individual experiments and the formula; $(COPY_1 - COPY_0)/(CTRL_1 - CTRL_0)$. Calculated from the average of two repeats, PR values closer to 1 indicated wider separation between sample outputs, while PR values closer to 0 (or below 0) indicate less separation. Figure S48(b) shows the PR data testing mismatches effect on IsoenergeticBitCopy.

Principal findings from debugging the IsoenergeticBitCopy experiment demonstrate that modulating compute strand binding energies with mismatches can help the system when reporting on a short length SDC and at shorter anneal times. However, simply dropping the temperature by slower rates in anneals also

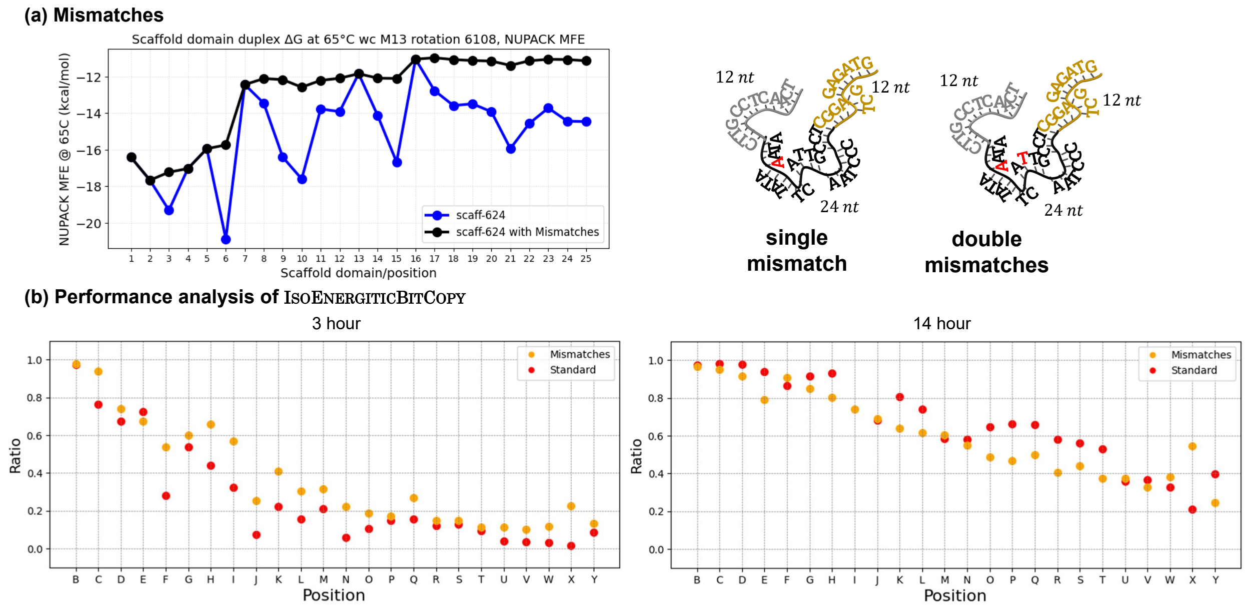

Figure S48: (a) Graph generated using the sculptor python library, illustrating the staircase-like energetic landscape at 65°C (black) compared to the standard scaffold energetic landscape (blue). Strand and sequence abstraction of potential mismatches on a IsoenergeticBitCopy compute strand. (b) Performance analysis of all four sets of experiments for IsoenergeticBitCopy for both 3-hour and 14-hour protocols; Standard (red circle), Mismatches (orange circle)

increases computational performance, which is hypothesised to allow the system to recover from hybridisation errors and kinetically trapped, off-target configurations.

157. Cervera Roldan, A., Woods, D. & Evans, C. Kinetic Simulation of a Scaffolded DNA Computer. (paper in preparation) (cit. on p. 72).

### S10 DNA sequences

This section lists all DNA sequences used in the experiments reported. We use the notation `d*` to denote the reverse complement of sequence/domain/strand `d`. The format is as follows:
`[strand_number]. [list_of_domains]: [DNA_sequence_with_space-separated_domains]`

A technical note on the naming convention used below: As described in Sections S5.1 and S6.2, there are two versions of each logical compute domain—the main text and this document uses an *overline* notation to make that distinction (e.g. `3` versus $\overline{tt3}$, when writing compute domains in base 10). This in turn explains why, for example, in the strand named $\overline{3}B3^*$ (below) the domains $\overline{3}$ = `TCAATCCTTGCC` and $3^*$ = `GACAAGGGTTGT` are not reverse complements. Indeed if they were, that strand would form a perfect hairpin, the very thing we wish to avoid.

Compute domain names of the form CD$i$ were defined in Section S6.2, but in the strand names below we simply call domains by their number $i$, i.e. we use $\overline{0}B4^*$: instead of the more verbose $\overline{\mathrm{CD0}}\,B\,\mathrm{CD4}^*$:.

The domains $F$ and $Q$ in Figure 2h of the main text have strand names `3RQ` and `5RF` below (where 3 and 5 refer to $3'$ and $5'$ ends, respectively—i.e. not compute domains!).

**Strand synthesis and purification** All strands were synthesised by IDT except for the M13 used in the scale-up experiments. Strands from IDT were ordered unpurified except for 7 strands which are the scaffold (`scaff`: Ultramer™ DNA Oligo PAGE purified and dry), the four labelled strands (`3RQ`, `5RF`, `3RQd`, `5RFd`: HPLC purified and normalised to 100 µM in IDTE pH 8.0 Buffer), and the four strands that the fluorophore-labelled strands bind to (ATTO$^*B$, ATTO$^*C$, ATTO$^*D$, ATTO$^*E$: PAGE purified and normalised to 100 µM in IDTE pH 8.0 Buffer). Of the four labelled strands, two had an 'ATTO590' fluorescent label at the $5'$ prime end and the others had an 'Iowa Black® FQ' quencher label at the $3'$ prime end. The $M13$ used in the scale-up experiments was purchased from tilibit nanosystems. Product: `Scaffold ssDNA 7249bp`, normalised to 400 nM (896 µg/ml) concentration in buffer containing 10 mM TRIS-BASE, 1 mM EDTA.

#### S10.1 DNA Sequences: 4-bit programs

##### S10.1.1 Synthetic 120nt Scaffold DNA sequence

1. scaff-120: $E^*$ $D^*$ $C^*$ $B^*$ $A^*$: ATTTTTGATTTATGGTCATTCTCG TTTTCTGAACTGTTTAAAGCATTT GAGGGGGATTCAATGAATATTTAT GACGATTCCGCAGTATTGGACGCT CTGGCAAATTAGGCTCTGGAAAGA

##### S10.1.2 Reporter complex DNA sequences

**Fluorophore-/quencher-labelled strands**

2. 3RQ: CCCACCTCTCCACACTACCC/3IABkFQ/
3. 5RF: /5ATTO590N/ACCATCCCTTCGCATCCCAA

**Strands that bind to fluorophore-labelled strands**

4. ATTO$^*B$: TTGGGATGCGAAGGGATGGT AGCGTCCAATACTGCGGAATCGTC
5. ATTO$^*C$: TTGGGATGCGAAGGGATGGT ATAAATATTCATTGAATCCCCCTC
6. ATTO$^*D$: TTGGGATGCGAAGGGATGGT AAATGCTTTAAACAGTTCAGAAAA
7. ATTO$^*E$: TTGGGATGCGAAGGGATGGT CGAGAATGACCATAAATCAAAAAT

**Strands that bind to quencher-labelled strands**

8. $AQ^*$: TCTTTCCAGAGCCTAATTTGCCAG GGGTAGTGTGGAGAGGTGGG
9. $BQ^*$: AGCGTCCAATACTGCGGAATCGTC GGGTAGTGTGGAGAGGTGGG
10. $CQ^*$: ATAAATATTCATTGAATCCCCCTC GGGTAGTGTGGAGAGGTGGG
11. $DQ^*$: AAATGCTTTAAACAGTTCAGAAAA GGGTAGTGTGGAGAGGTGGG
12. $\overline{0}BQ^*$: CCTCTTCTCAGC AGCGTCCAATACTGCGGAATCGTC GGGTAGTGTGGAGAGGTGGG
13. $\overline{1}BQ^*$: CATCTCCGATCC AGCGTCCAATACTGCGGAATCGTC GGGTAGTGTGGAGAGGTGGG
14. $\overline{2}BQ^*$: TCTTTCCAAGCC AGCGTCCAATACTGCGGAATCGTC GGGTAGTGTGGAGAGGTGGG
15. $\overline{3}BQ^*$: TCAATCCTTGCC AGCGTCCAATACTGCGGAATCGTC GGGTAGTGTGGAGAGGTGGG

16. $\overline{4}BQ^*$: CACATCCCTGTT AGCGTCCAATACTGCGGAATCGTC GGGTAGTGTGGAGAGGTGGG
17. $\overline{5}BQ^*$: CCATGTCCCATT AGCGTCCAATACTGCGGAATCGTC GGGTAGTGTGGAGAGGTGGG
18. $\overline{6}BQ^*$: CAACCAACGTTC AGCGTCCAATACTGCGGAATCGTC GGGTAGTGTGGAGAGGTGGG
19. $\overline{7}BQ^*$: TCACACTTCGTC AGCGTCCAATACTGCGGAATCGTC GGGTAGTGTGGAGAGGTGGG
20. $0CQ^*$: CTCATCCTGACC ATAAATATTCATTGAATCCCCCTC GGGTAGTGTGGAGAGGTGGG
21. $1CQ^*$: TCAACTCCGTTC ATAAATATTCATTGAATCCCCCTC GGGTAGTGTGGAGAGGTGGG
22. $2CQ^*$: AATGCCACCATT ATAAATATTCATTGAATCCCCCTC GGGTAGTGTGGAGAGGTGGG
23. $3CQ^*$: ACAACCCTTGTC ATAAATATTCATTGAATCCCCCTC GGGTAGTGTGGAGAGGTGGG
24. $4CQ^*$: CTGTTCCCAACA ATAAATATTCATTGAATCCCCCTC GGGTAGTGTGGAGAGGTGGG
25. $5CQ^*$: CACTACCAGTCC ATAAATATTCATTGAATCCCCCTC GGGTAGTGTGGAGAGGTGGG
26. $6CQ^*$: ACACACACTGTC ATAAATATTCATTGAATCCCCCTC GGGTAGTGTGGAGAGGTGGG
27. $7CQ^*$: TCACTTTCGTCC ATAAATATTCATTGAATCCCCCTC GGGTAGTGTGGAGAGGTGGG
28. $\overline{0}DQ^*$: CCTCTTCTCAGC AAATGCTTTAAACAGTTCAGAAAA GGGTAGTGTGGAGAGGTGGG
29. $\overline{1}DQ^*$: CATCTCCGATCC AAATGCTTTAAACAGTTCAGAAAA GGGTAGTGTGGAGAGGTGGG
30. $\overline{2}DQ^*$: TCTTTCCAAGCC AAATGCTTTAAACAGTTCAGAAAA GGGTAGTGTGGAGAGGTGGG
31. $\overline{3}DQ^*$: TCAATCCTTGCC AAATGCTTTAAACAGTTCAGAAAA GGGTAGTGTGGAGAGGTGGG
32. $\overline{4}DQ^*$: CACATCCCTGTT AAATGCTTTAAACAGTTCAGAAAA GGGTAGTGTGGAGAGGTGGG
33. $\overline{5}DQ^*$: CCATGTCCCATT AAATGCTTTAAACAGTTCAGAAAA GGGTAGTGTGGAGAGGTGGG
34. $\overline{6}DQ^*$: CAACCAACGTTC AAATGCTTTAAACAGTTCAGAAAA GGGTAGTGTGGAGAGGTGGG
35. $\overline{7}DQ^*$: TCACACTTCGTC AAATGCTTTAAACAGTTCAGAAAA GGGTAGTGTGGAGAGGTGGG
36. ATTO$^*Q^*$: TTGGGATGCGAAGGGATGGT GGGTAGTGTGGAGAGGTGGG

**Strands with no compute domain on the right-hand side (3′ end)**

37. $A$: TCTTTCCAGAGCCTAATTTGCCAG
38. $B$: AGCGTCCAATACTGCGGAATCGTC
39. $C$: ATAAATATTCATTGAATCCCCCTC
40. $D$: AAATGCTTTAAACAGTTCAGAAAA
41. $E$: CGAGAATGACCATAAATCAAAAAT
42. $\overline{0}B$: CCTCTTCTCAGC AGCGTCCAATACTGCGGAATCGTC
43. $\overline{1}B$: CATCTCCGATCC AGCGTCCAATACTGCGGAATCGTC
44. $\overline{2}B$: TCTTTCCAAGCC AGCGTCCAATACTGCGGAATCGTC
45. $\overline{3}B$: TCAATCCTTGCC AGCGTCCAATACTGCGGAATCGTC
46. $\overline{4}B$: CACATCCCTGTT AGCGTCCAATACTGCGGAATCGTC
47. $\overline{5}B$: CCATGTCCCATT AGCGTCCAATACTGCGGAATCGTC
48. $\overline{6}B$: CAACCAACGTTC AGCGTCCAATACTGCGGAATCGTC
49. $\overline{7}B$: TCACACTTCGTC AGCGTCCAATACTGCGGAATCGTC
50. $0C$: CTCATCCTGACC ATAAATATTCATTGAATCCCCCTC
51. $1C$: TCAACTCCGTTC ATAAATATTCATTGAATCCCCCTC
52. $2C$: AATGCCACCATT ATAAATATTCATTGAATCCCCCTC
53. $3C$: ACAACCCTTGTC ATAAATATTCATTGAATCCCCCTC
54. $4C$: CTGTTCCCAACA ATAAATATTCATTGAATCCCCCTC
55. $5C$: CACTACCAGTCC ATAAATATTCATTGAATCCCCCTC
56. $6C$: ACACACACTGTC ATAAATATTCATTGAATCCCCCTC
57. $7C$: TCACTTTCGTCC ATAAATATTCATTGAATCCCCCTC
58. $\overline{0}D$: CCTCTTCTCAGC AAATGCTTTAAACAGTTCAGAAAA
59. $\overline{1}D$: CATCTCCGATCC AAATGCTTTAAACAGTTCAGAAAA
60. $\overline{2}D$: TCTTTCCAAGCC AAATGCTTTAAACAGTTCAGAAAA
61. $\overline{3}D$: TCAATCCTTGCC AAATGCTTTAAACAGTTCAGAAAA
62. $\overline{4}D$: CACATCCCTGTT AAATGCTTTAAACAGTTCAGAAAA
63. $\overline{5}D$: CCATGTCCCATT AAATGCTTTAAACAGTTCAGAAAA
64. $\overline{6}D$: CAACCAACGTTC AAATGCTTTAAACAGTTCAGAAAA
65. $\overline{7}D$: TCACACTTCGTC AAATGCTTTAAACAGTTCAGAAAA

##### S10.1.3 DNA sequences for compute strands

**Computing strands for position $A$**

66. $A\overline{0}^*$: TCTTTCCAGAGCCTAATTTGCCAG GCTGAGAAGAGG
67. $A\overline{1}^*$: TCTTTCCAGAGCCTAATTTGCCAG GGATCGGAGATG
68. $A\overline{2}^*$: TCTTTCCAGAGCCTAATTTGCCAG GGCTTGGAAAGA
69. $A\overline{3}^*$: TCTTTCCAGAGCCTAATTTGCCAG GGCAAGGATTGA
70. $A\overline{4}^*$: TCTTTCCAGAGCCTAATTTGCCAG AACAGGGATGTG
71. $A\overline{5}^*$: TCTTTCCAGAGCCTAATTTGCCAG AATGGGACATGG
72. $A\overline{6}^*$: TCTTTCCAGAGCCTAATTTGCCAG GAACGTTGGTTG
73. $A\overline{7}^*$: TCTTTCCAGAGCCTAATTTGCCAG GACGAAGTGTGA

**Computing strands for position $B$**

74. $\overline{0}B0^*$: CCTCTTCTCAGC AGCGTCCAATACTGCGGAATCGTC GGTCAGGATGAG
75. $\overline{0}B1^*$: CCTCTTCTCAGC AGCGTCCAATACTGCGGAATCGTC GAACGGAGTTGA
76. $\overline{0}B2^*$: CCTCTTCTCAGC AGCGTCCAATACTGCGGAATCGTC AATGGTGGCATT
77. $\overline{0}B3^*$: CCTCTTCTCAGC AGCGTCCAATACTGCGGAATCGTC GACAAGGGTTGT
78. $\overline{0}B4^*$: CCTCTTCTCAGC AGCGTCCAATACTGCGGAATCGTC TGTTGGGAACAG
79. $\overline{0}B5^*$: CCTCTTCTCAGC AGCGTCCAATACTGCGGAATCGTC GGACTGGTAGTG
80. $\overline{0}B6^*$: CCTCTTCTCAGC AGCGTCCAATACTGCGGAATCGTC GACAGTGTGTGT
81. $\overline{0}B7^*$: CCTCTTCTCAGC AGCGTCCAATACTGCGGAATCGTC GGACGAAAGTGA
82. $\overline{1}B0^*$: CATCTCCGATCC AGCGTCCAATACTGCGGAATCGTC GGTCAGGATGAG
83. $\overline{1}B1^*$: CATCTCCGATCC AGCGTCCAATACTGCGGAATCGTC GAACGGAGTTGA
84. $\overline{1}B2^*$: CATCTCCGATCC AGCGTCCAATACTGCGGAATCGTC AATGGTGGCATT
85. $\overline{1}B3^*$: CATCTCCGATCC AGCGTCCAATACTGCGGAATCGTC GACAAGGGTTGT
86. $\overline{1}B4^*$: CATCTCCGATCC AGCGTCCAATACTGCGGAATCGTC TGTTGGGAACAG
87. $\overline{1}B5^*$: CATCTCCGATCC AGCGTCCAATACTGCGGAATCGTC GGACTGGTAGTG
88. $\overline{1}B6^*$: CATCTCCGATCC AGCGTCCAATACTGCGGAATCGTC GACAGTGTGTGT
89. $\overline{1}B7^*$: CATCTCCGATCC AGCGTCCAATACTGCGGAATCGTC GGACGAAAGTGA
90. $\overline{2}B0^*$: TCTTTCCAAGCC AGCGTCCAATACTGCGGAATCGTC GGTCAGGATGAG
91. $\overline{2}B1^*$: TCTTTCCAAGCC AGCGTCCAATACTGCGGAATCGTC GAACGGAGTTGA
92. $\overline{2}B2^*$: TCTTTCCAAGCC AGCGTCCAATACTGCGGAATCGTC AATGGTGGCATT
93. $\overline{2}B3^*$: TCTTTCCAAGCC AGCGTCCAATACTGCGGAATCGTC GACAAGGGTTGT
94. $\overline{2}B4^*$: TCTTTCCAAGCC AGCGTCCAATACTGCGGAATCGTC TGTTGGGAACAG
95. $\overline{2}B5^*$: TCTTTCCAAGCC AGCGTCCAATACTGCGGAATCGTC GGACTGGTAGTG
96. $\overline{2}B6^*$: TCTTTCCAAGCC AGCGTCCAATACTGCGGAATCGTC GACAGTGTGTGT
97. $\overline{2}B7^*$: TCTTTCCAAGCC AGCGTCCAATACTGCGGAATCGTC GGACGAAAGTGA
98. $\overline{3}B0^*$: TCAATCCTTGCC AGCGTCCAATACTGCGGAATCGTC GGTCAGGATGAG
99. $\overline{3}B1^*$: TCAATCCTTGCC AGCGTCCAATACTGCGGAATCGTC GAACGGAGTTGA
100. $\overline{3}B2^*$: TCAATCCTTGCC AGCGTCCAATACTGCGGAATCGTC AATGGTGGCATT
101. $\overline{3}B3^*$: TCAATCCTTGCC AGCGTCCAATACTGCGGAATCGTC GACAAGGGTTGT
102. $\overline{3}B4^*$: TCAATCCTTGCC AGCGTCCAATACTGCGGAATCGTC TGTTGGGAACAG
103. $\overline{3}B5^*$: TCAATCCTTGCC AGCGTCCAATACTGCGGAATCGTC GGACTGGTAGTG
104. $\overline{3}B6^*$: TCAATCCTTGCC AGCGTCCAATACTGCGGAATCGTC GACAGTGTGTGT
105. $\overline{3}B7^*$: TCAATCCTTGCC AGCGTCCAATACTGCGGAATCGTC GGACGAAAGTGA
106. $\overline{4}B0^*$: CACATCCCTGTT AGCGTCCAATACTGCGGAATCGTC GGTCAGGATGAG
107. $\overline{4}B1^*$: CACATCCCTGTT AGCGTCCAATACTGCGGAATCGTC GAACGGAGTTGA
108. $\overline{4}B2^*$: CACATCCCTGTT AGCGTCCAATACTGCGGAATCGTC AATGGTGGCATT
109. $\overline{4}B3^*$: CACATCCCTGTT AGCGTCCAATACTGCGGAATCGTC GACAAGGGTTGT
110. $\overline{4}B4^*$: CACATCCCTGTT AGCGTCCAATACTGCGGAATCGTC TGTTGGGAACAG
111. $\overline{4}B5^*$: CACATCCCTGTT AGCGTCCAATACTGCGGAATCGTC GGACTGGTAGTG
112. $\overline{4}B6^*$: CACATCCCTGTT AGCGTCCAATACTGCGGAATCGTC GACAGTGTGTGT
113. $\overline{4}B7^*$: CACATCCCTGTT AGCGTCCAATACTGCGGAATCGTC GGACGAAAGTGA
114. $\overline{5}B0^*$: CCATGTCCCATT AGCGTCCAATACTGCGGAATCGTC GGTCAGGATGAG

115. $\overline{5}B1^*$: CCATGTCCCATT AGCGTCCAATACTGCGGAATCGTC GAACGGAGTTGA
116. $\overline{5}B2^*$: CCATGTCCCATT AGCGTCCAATACTGCGGAATCGTC AATGGTGGCATT
117. $\overline{5}B3^*$: CCATGTCCCATT AGCGTCCAATACTGCGGAATCGTC GACAAGGGTTGT
118. $\overline{5}B4^*$: CCATGTCCCATT AGCGTCCAATACTGCGGAATCGTC TGTTGGGAACAG
119. $\overline{5}B5^*$: CCATGTCCCATT AGCGTCCAATACTGCGGAATCGTC GGACTGGTAGTG
120. $\overline{5}B6^*$: CCATGTCCCATT AGCGTCCAATACTGCGGAATCGTC GACAGTGTGTGT
121. $\overline{5}B7^*$: CCATGTCCCATT AGCGTCCAATACTGCGGAATCGTC GGACGAAAGTGA
122. $\overline{6}B0^*$: CAACCAACGTTC AGCGTCCAATACTGCGGAATCGTC GGTCAGGATGAG
123. $\overline{6}B1^*$: CAACCAACGTTC AGCGTCCAATACTGCGGAATCGTC GAACGGAGTTGA
124. $\overline{6}B2^*$: CAACCAACGTTC AGCGTCCAATACTGCGGAATCGTC AATGGTGGCATT
125. $\overline{6}B3^*$: CAACCAACGTTC AGCGTCCAATACTGCGGAATCGTC GACAAGGGTTGT
126. $\overline{6}B4^*$: CAACCAACGTTC AGCGTCCAATACTGCGGAATCGTC TGTTGGGAACAG
127. $\overline{6}B5^*$: CAACCAACGTTC AGCGTCCAATACTGCGGAATCGTC GGACTGGTAGTG
128. $\overline{6}B6^*$: CAACCAACGTTC AGCGTCCAATACTGCGGAATCGTC GACAGTGTGTGT
129. $\overline{6}B7^*$: CAACCAACGTTC AGCGTCCAATACTGCGGAATCGTC GGACGAAAGTGA
130. $\overline{7}B0^*$: TCACACTTCGTC AGCGTCCAATACTGCGGAATCGTC GGTCAGGATGAG
131. $\overline{7}B1^*$: TCACACTTCGTC AGCGTCCAATACTGCGGAATCGTC GAACGGAGTTGA
132. $\overline{7}B2^*$: TCACACTTCGTC AGCGTCCAATACTGCGGAATCGTC AATGGTGGCATT
133. $\overline{7}B3^*$: TCACACTTCGTC AGCGTCCAATACTGCGGAATCGTC GACAAGGGTTGT
134. $\overline{7}B4^*$: TCACACTTCGTC AGCGTCCAATACTGCGGAATCGTC TGTTGGGAACAG
135. $\overline{7}B5^*$: TCACACTTCGTC AGCGTCCAATACTGCGGAATCGTC GGACTGGTAGTG
136. $\overline{7}B6^*$: TCACACTTCGTC AGCGTCCAATACTGCGGAATCGTC GACAGTGTGTGT
137. $\overline{7}B7^*$: TCACACTTCGTC AGCGTCCAATACTGCGGAATCGTC GGACGAAAGTGA

**Computing strands for position $C$**

138. $0C\overline{0}^*$: CTCATCCTGACC ATAAATATTCATTGAATCCCCCTC GCTGAGAAGAGG
139. $0C\overline{1}^*$: CTCATCCTGACC ATAAATATTCATTGAATCCCCCTC GGATCGGAGATG
140. $0C\overline{2}^*$: CTCATCCTGACC ATAAATATTCATTGAATCCCCCTC GGCTTGGAAAGA
141. $0C\overline{3}^*$: CTCATCCTGACC ATAAATATTCATTGAATCCCCCTC GGCAAGGATTGA
142. $0C\overline{4}^*$: CTCATCCTGACC ATAAATATTCATTGAATCCCCCTC AACAGGGATGTG
143. $0C\overline{5}^*$: CTCATCCTGACC ATAAATATTCATTGAATCCCCCTC AATGGGACATGG
144. $0C\overline{6}^*$: CTCATCCTGACC ATAAATATTCATTGAATCCCCCTC GAACGTTGGTTG
145. $0C\overline{7}^*$: CTCATCCTGACC ATAAATATTCATTGAATCCCCCTC GACGAAGTGTGA
146. $1C\overline{0}^*$: TCAACTCCGTTC ATAAATATTCATTGAATCCCCCTC GCTGAGAAGAGG
147. $1C\overline{1}^*$: TCAACTCCGTTC ATAAATATTCATTGAATCCCCCTC GGATCGGAGATG
148. $1C\overline{2}^*$: TCAACTCCGTTC ATAAATATTCATTGAATCCCCCTC GGCTTGGAAAGA
149. $1C\overline{3}^*$: TCAACTCCGTTC ATAAATATTCATTGAATCCCCCTC GGCAAGGATTGA
150. $1C\overline{4}^*$: TCAACTCCGTTC ATAAATATTCATTGAATCCCCCTC AACAGGGATGTG
151. $1C\overline{5}^*$: TCAACTCCGTTC ATAAATATTCATTGAATCCCCCTC AATGGGACATGG
152. $1C\overline{6}^*$: TCAACTCCGTTC ATAAATATTCATTGAATCCCCCTC GAACGTTGGTTG
153. $1C\overline{7}^*$: TCAACTCCGTTC ATAAATATTCATTGAATCCCCCTC GACGAAGTGTGA
154. $2C\overline{0}^*$: AATGCCACCATT ATAAATATTCATTGAATCCCCCTC GCTGAGAAGAGG
155. $2C\overline{1}^*$: AATGCCACCATT ATAAATATTCATTGAATCCCCCTC GGATCGGAGATG
156. $2C\overline{2}^*$: AATGCCACCATT ATAAATATTCATTGAATCCCCCTC GGCTTGGAAAGA
157. $2C\overline{3}^*$: AATGCCACCATT ATAAATATTCATTGAATCCCCCTC GGCAAGGATTGA
158. $2C\overline{4}^*$: AATGCCACCATT ATAAATATTCATTGAATCCCCCTC AACAGGGATGTG
159. $2C\overline{5}^*$: AATGCCACCATT ATAAATATTCATTGAATCCCCCTC AATGGGACATGG
160. $2C\overline{6}^*$: AATGCCACCATT ATAAATATTCATTGAATCCCCCTC GAACGTTGGTTG
161. $2C\overline{7}^*$: AATGCCACCATT ATAAATATTCATTGAATCCCCCTC GACGAAGTGTGA
162. $3C\overline{0}^*$: ACAACCCTTGTC ATAAATATTCATTGAATCCCCCTC GCTGAGAAGAGG
163. $3C\overline{1}^*$: ACAACCCTTGTC ATAAATATTCATTGAATCCCCCTC GGATCGGAGATG
164. $3C\overline{2}^*$: ACAACCCTTGTC ATAAATATTCATTGAATCCCCCTC GGCTTGGAAAGA
165. $3C\overline{3}^*$: ACAACCCTTGTC ATAAATATTCATTGAATCCCCCTC GGCAAGGATTGA
166. $3C\overline{4}^*$: ACAACCCTTGTC ATAAATATTCATTGAATCCCCCTC AACAGGGATGTG

167. $3C\overline{5}^*$: ACAACCCTTGTC ATAAATATTCATTGAATCCCCCTC AATGGGACATGG
168. $3C\overline{6}^*$: ACAACCCTTGTC ATAAATATTCATTGAATCCCCCTC GAACGTTGGTTG
169. $3C\overline{7}^*$: ACAACCCTTGTC ATAAATATTCATTGAATCCCCCTC GACGAAGTGTGA
170. $4C\overline{0}^*$: CTGTTCCCAACA ATAAATATTCATTGAATCCCCCTC GCTGAGAAGAGG
171. $4C\overline{1}^*$: CTGTTCCCAACA ATAAATATTCATTGAATCCCCCTC GGATCGGAGATG
172. $4C\overline{2}^*$: CTGTTCCCAACA ATAAATATTCATTGAATCCCCCTC GGCTTGGAAAGA
173. $4C\overline{3}^*$: CTGTTCCCAACA ATAAATATTCATTGAATCCCCCTC GGCAAGGATTGA
174. $4C\overline{4}^*$: CTGTTCCCAACA ATAAATATTCATTGAATCCCCCTC AACAGGGATGTG
175. $4C\overline{5}^*$: CTGTTCCCAACA ATAAATATTCATTGAATCCCCCTC AATGGGACATGG
176. $4C\overline{6}^*$: CTGTTCCCAACA ATAAATATTCATTGAATCCCCCTC GAACGTTGGTTG
177. $4C\overline{7}^*$: CTGTTCCCAACA ATAAATATTCATTGAATCCCCCTC GACGAAGTGTGA
178. $5C\overline{0}^*$: CACTACCAGTCC ATAAATATTCATTGAATCCCCCTC GCTGAGAAGAGG
179. $5C\overline{1}^*$: CACTACCAGTCC ATAAATATTCATTGAATCCCCCTC GGATCGGAGATG
180. $5C\overline{2}^*$: CACTACCAGTCC ATAAATATTCATTGAATCCCCCTC GGCTTGGAAAGA
181. $5C\overline{3}^*$: CACTACCAGTCC ATAAATATTCATTGAATCCCCCTC GGCAAGGATTGA
182. $5C\overline{4}^*$: CACTACCAGTCC ATAAATATTCATTGAATCCCCCTC AACAGGGATGTG
183. $5C\overline{5}^*$: CACTACCAGTCC ATAAATATTCATTGAATCCCCCTC AATGGGACATGG
184. $5C\overline{6}^*$: CACTACCAGTCC ATAAATATTCATTGAATCCCCCTC GAACGTTGGTTG
185. $5C\overline{7}^*$: CACTACCAGTCC ATAAATATTCATTGAATCCCCCTC GACGAAGTGTGA
186. $6C\overline{0}^*$: ACACACACTGTC ATAAATATTCATTGAATCCCCCTC GCTGAGAAGAGG
187. $6C\overline{1}^*$: ACACACACTGTC ATAAATATTCATTGAATCCCCCTC GGATCGGAGATG
188. $6C\overline{2}^*$: ACACACACTGTC ATAAATATTCATTGAATCCCCCTC GGCTTGGAAAGA
189. $6C\overline{3}^*$: ACACACACTGTC ATAAATATTCATTGAATCCCCCTC GGCAAGGATTGA
190. $6C\overline{4}^*$: ACACACACTGTC ATAAATATTCATTGAATCCCCCTC AACAGGGATGTG
191. $6C\overline{5}^*$: ACACACACTGTC ATAAATATTCATTGAATCCCCCTC AATGGGACATGG
192. $6C\overline{6}^*$: ACACACACTGTC ATAAATATTCATTGAATCCCCCTC GAACGTTGGTTG
193. $6C\overline{7}^*$: ACACACACTGTC ATAAATATTCATTGAATCCCCCTC GACGAAGTGTGA
194. $7C\overline{0}^*$: TCACTTTCGTCC ATAAATATTCATTGAATCCCCCTC GCTGAGAAGAGG
195. $7C\overline{1}^*$: TCACTTTCGTCC ATAAATATTCATTGAATCCCCCTC GGATCGGAGATG
196. $7C\overline{2}^*$: TCACTTTCGTCC ATAAATATTCATTGAATCCCCCTC GGCTTGGAAAGA
197. $7C\overline{3}^*$: TCACTTTCGTCC ATAAATATTCATTGAATCCCCCTC GGCAAGGATTGA
198. $7C\overline{4}^*$: TCACTTTCGTCC ATAAATATTCATTGAATCCCCCTC AACAGGGATGTG
199. $7C\overline{5}^*$: TCACTTTCGTCC ATAAATATTCATTGAATCCCCCTC AATGGGACATGG
200. $7C\overline{6}^*$: TCACTTTCGTCC ATAAATATTCATTGAATCCCCCTC GAACGTTGGTTG
201. $7C\overline{7}^*$: TCACTTTCGTCC ATAAATATTCATTGAATCCCCCTC GACGAAGTGTGA

**Computing strands for position $D$**

202. $\overline{0}D0^*$: CCTCTTCTCAGC AAATGCTTTAAACAGTTCAGAAAA GGTCAGGATGAG
203. $\overline{0}D1^*$: CCTCTTCTCAGC AAATGCTTTAAACAGTTCAGAAAA GAACGGAGTTGA
204. $\overline{0}D2^*$: CCTCTTCTCAGC AAATGCTTTAAACAGTTCAGAAAA AATGGTGGCATT
205. $\overline{0}D3^*$: CCTCTTCTCAGC AAATGCTTTAAACAGTTCAGAAAA GACAAGGGTTGT
206. $\overline{0}D4^*$: CCTCTTCTCAGC AAATGCTTTAAACAGTTCAGAAAA TGTTGGGAACAG
207. $\overline{0}D5^*$: CCTCTTCTCAGC AAATGCTTTAAACAGTTCAGAAAA GGACTGGTAGTG
208. $\overline{0}D6^*$: CCTCTTCTCAGC AAATGCTTTAAACAGTTCAGAAAA GACAGTGTGTGT
209. $\overline{0}D7^*$: CCTCTTCTCAGC AAATGCTTTAAACAGTTCAGAAAA GGACGAAAGTGA
210. $\overline{1}D0^*$: CATCTCCGATCC AAATGCTTTAAACAGTTCAGAAAA GGTCAGGATGAG
211. $\overline{1}D1^*$: CATCTCCGATCC AAATGCTTTAAACAGTTCAGAAAA GAACGGAGTTGA
212. $\overline{1}D2^*$: CATCTCCGATCC AAATGCTTTAAACAGTTCAGAAAA AATGGTGGCATT
213. $\overline{1}D3^*$: CATCTCCGATCC AAATGCTTTAAACAGTTCAGAAAA GACAAGGGTTGT
214. $\overline{1}D4^*$: CATCTCCGATCC AAATGCTTTAAACAGTTCAGAAAA TGTTGGGAACAG
215. $\overline{1}D5^*$: CATCTCCGATCC AAATGCTTTAAACAGTTCAGAAAA GGACTGGTAGTG
216. $\overline{1}D6^*$: CATCTCCGATCC AAATGCTTTAAACAGTTCAGAAAA GACAGTGTGTGT
217. $\overline{1}D7^*$: CATCTCCGATCC AAATGCTTTAAACAGTTCAGAAAA GGACGAAAGTGA
218. $\overline{2}D0^*$: TCTTTCCAAGCC AAATGCTTTAAACAGTTCAGAAAA GGTCAGGATGAG

219. $\overline{2}D1^*$: TCTTTCCAAGCC AAATGCTTTAAACAGTTCAGAAAA GAACGGAGTTGA
220. $\overline{2}D2^*$: TCTTTCCAAGCC AAATGCTTTAAACAGTTCAGAAAA AATGGTGGCATT
221. $\overline{2}D3^*$: TCTTTCCAAGCC AAATGCTTTAAACAGTTCAGAAAA GACAAGGGTTGT
222. $\overline{2}D4^*$: TCTTTCCAAGCC AAATGCTTTAAACAGTTCAGAAAA TGTTGGGAACAG
223. $\overline{2}D5^*$: TCTTTCCAAGCC AAATGCTTTAAACAGTTCAGAAAA GGACTGGTAGTG
224. $\overline{2}D6^*$: TCTTTCCAAGCC AAATGCTTTAAACAGTTCAGAAAA GACAGTGTGTGT
225. $\overline{2}D7^*$: TCTTTCCAAGCC AAATGCTTTAAACAGTTCAGAAAA GGACGAAAGTGA
226. $\overline{3}D0^*$: TCAATCCTTGCC AAATGCTTTAAACAGTTCAGAAAA GGTCAGGATGAG
227. $\overline{3}D1^*$: TCAATCCTTGCC AAATGCTTTAAACAGTTCAGAAAA GAACGGAGTTGA
228. $\overline{3}D2^*$: TCAATCCTTGCC AAATGCTTTAAACAGTTCAGAAAA AATGGTGGCATT
229. $\overline{3}D3^*$: TCAATCCTTGCC AAATGCTTTAAACAGTTCAGAAAA GACAAGGGTTGT
230. $\overline{3}D4^*$: TCAATCCTTGCC AAATGCTTTAAACAGTTCAGAAAA TGTTGGGAACAG
231. $\overline{3}D5^*$: TCAATCCTTGCC AAATGCTTTAAACAGTTCAGAAAA GGACTGGTAGTG
232. $\overline{3}D6^*$: TCAATCCTTGCC AAATGCTTTAAACAGTTCAGAAAA GACAGTGTGTGT
233. $\overline{3}D7^*$: TCAATCCTTGCC AAATGCTTTAAACAGTTCAGAAAA GGACGAAAGTGA
234. $\overline{4}D0^*$: CACATCCCTGTT AAATGCTTTAAACAGTTCAGAAAA GGTCAGGATGAG
235. $\overline{4}D1^*$: CACATCCCTGTT AAATGCTTTAAACAGTTCAGAAAA GAACGGAGTTGA
236. $\overline{4}D2^*$: CACATCCCTGTT AAATGCTTTAAACAGTTCAGAAAA AATGGTGGCATT
237. $\overline{4}D3^*$: CACATCCCTGTT AAATGCTTTAAACAGTTCAGAAAA GACAAGGGTTGT
238. $\overline{4}D4^*$: CACATCCCTGTT AAATGCTTTAAACAGTTCAGAAAA TGTTGGGAACAG
239. $\overline{4}D5^*$: CACATCCCTGTT AAATGCTTTAAACAGTTCAGAAAA GGACTGGTAGTG
240. $\overline{4}D6^*$: CACATCCCTGTT AAATGCTTTAAACAGTTCAGAAAA GACAGTGTGTGT
241. $\overline{4}D7^*$: CACATCCCTGTT AAATGCTTTAAACAGTTCAGAAAA GGACGAAAGTGA
242. $\overline{5}D0^*$: CCATGTCCCATT AAATGCTTTAAACAGTTCAGAAAA GGTCAGGATGAG
243. $\overline{5}D1^*$: CCATGTCCCATT AAATGCTTTAAACAGTTCAGAAAA GAACGGAGTTGA
244. $\overline{5}D2^*$: CCATGTCCCATT AAATGCTTTAAACAGTTCAGAAAA AATGGTGGCATT
245. $\overline{5}D3^*$: CCATGTCCCATT AAATGCTTTAAACAGTTCAGAAAA GACAAGGGTTGT
246. $\overline{5}D4^*$: CCATGTCCCATT AAATGCTTTAAACAGTTCAGAAAA TGTTGGGAACAG
247. $\overline{5}D5^*$: CCATGTCCCATT AAATGCTTTAAACAGTTCAGAAAA GGACTGGTAGTG
248. $\overline{5}D6^*$: CCATGTCCCATT AAATGCTTTAAACAGTTCAGAAAA GACAGTGTGTGT
249. $\overline{5}D7^*$: CCATGTCCCATT AAATGCTTTAAACAGTTCAGAAAA GGACGAAAGTGA
250. $\overline{6}D0^*$: CAACCAACGTTC AAATGCTTTAAACAGTTCAGAAAA GGTCAGGATGAG
251. $\overline{6}D1^*$: CAACCAACGTTC AAATGCTTTAAACAGTTCAGAAAA GAACGGAGTTGA
252. $\overline{6}D2^*$: CAACCAACGTTC AAATGCTTTAAACAGTTCAGAAAA AATGGTGGCATT
253. $\overline{6}D3^*$: CAACCAACGTTC AAATGCTTTAAACAGTTCAGAAAA GACAAGGGTTGT
254. $\overline{6}D4^*$: CAACCAACGTTC AAATGCTTTAAACAGTTCAGAAAA TGTTGGGAACAG
255. $\overline{6}D5^*$: CAACCAACGTTC AAATGCTTTAAACAGTTCAGAAAA GGACTGGTAGTG
256. $\overline{6}D6^*$: CAACCAACGTTC AAATGCTTTAAACAGTTCAGAAAA GACAGTGTGTGT
257. $\overline{6}D7^*$: CAACCAACGTTC AAATGCTTTAAACAGTTCAGAAAA GGACGAAAGTGA
258. $\overline{7}D0^*$: TCACACTTCGTC AAATGCTTTAAACAGTTCAGAAAA GGTCAGGATGAG
259. $\overline{7}D1^*$: TCACACTTCGTC AAATGCTTTAAACAGTTCAGAAAA GAACGGAGTTGA
260. $\overline{7}D2^*$: TCACACTTCGTC AAATGCTTTAAACAGTTCAGAAAA AATGGTGGCATT
261. $\overline{7}D3^*$: TCACACTTCGTC AAATGCTTTAAACAGTTCAGAAAA GACAAGGGTTGT
262. $\overline{7}D4^*$: TCACACTTCGTC AAATGCTTTAAACAGTTCAGAAAA TGTTGGGAACAG
263. $\overline{7}D5^*$: TCACACTTCGTC AAATGCTTTAAACAGTTCAGAAAA GGACTGGTAGTG
264. $\overline{7}D6^*$: TCACACTTCGTC AAATGCTTTAAACAGTTCAGAAAA GACAGTGTGTGT
265. $\overline{7}D7^*$: TCACACTTCGTC AAATGCTTTAAACAGTTCAGAAAA GGACGAAAGTGA

##### S10.1.4 DNA sequences for renewable programs

**Renewing strands for position $A$**

266. L($AQ^*$): CCCACCTCTCCACACTACCC CTGGCAAATTAGGCTCTGGAAAGA
267. L($A\overline{0}^*$): CCTCTTCTCAGC CTGGCAAATTAGGCTCTGGAAAGA
268. L($A\overline{1}^*$): CATCTCCGATCC CTGGCAAATTAGGCTCTGGAAAGA

269. L($A\overline{2}^*$): TCTTTCCAAGCC CTGGCAAATTAGGCTCTGGAAAGA
270. L($A\overline{3}^*$): TCAATCCTTGCC CTGGCAAATTAGGCTCTGGAAAGA
271. L($A\overline{4}^*$): CACATCCCTGTT CTGGCAAATTAGGCTCTGGAAAGA
272. L($A\overline{5}^*$): CCATGTCCCATT CTGGCAAATTAGGCTCTGGAAAGA
273. L($A\overline{6}^*$): CAACCAACGTTC CTGGCAAATTAGGCTCTGGAAAGA
274. L($A\overline{7}^*$): TCACACTTCGTC CTGGCAAATTAGGCTCTGGAAAGA
275. $0AQ^*$: CTCATCCTGACC TCTTTCCAGAGCCTAATTTGCCAG GGGTAGTGTGGAGAGGTGGG
276. $0A\overline{0}^*$: CTCATCCTGACC TCTTTCCAGAGCCTAATTTGCCAG GCTGAGAAGAGG
277. $0A\overline{1}^*$: CTCATCCTGACC TCTTTCCAGAGCCTAATTTGCCAG GGATCGGAGATG
278. $0A\overline{2}^*$: CTCATCCTGACC TCTTTCCAGAGCCTAATTTGCCAG GGCTTGGAAAGA
279. $0A\overline{3}^*$: CTCATCCTGACC TCTTTCCAGAGCCTAATTTGCCAG GGCAAGGATTGA
280. $0A\overline{4}^*$: CTCATCCTGACC TCTTTCCAGAGCCTAATTTGCCAG AACAGGGATGTG
281. $0A\overline{5}^*$: CTCATCCTGACC TCTTTCCAGAGCCTAATTTGCCAG AATGGGACATGG
282. $0A\overline{6}^*$: CTCATCCTGACC TCTTTCCAGAGCCTAATTTGCCAG GAACGTTGGTTG
283. $0A\overline{7}^*$: CTCATCCTGACC TCTTTCCAGAGCCTAATTTGCCAG GACGAAGTGTGA
284. L($0AQ^*$): CCCACCTCTCCACACTACCC CTGGCAAATTAGGCTCTGGAAAGA GGTCAGGATGAG
285. L($0A\overline{0}^*$): CCTCTTCTCAGC CTGGCAAATTAGGCTCTGGAAAGA GGTCAGGATGAG
286. L($0A\overline{1}^*$): CATCTCCGATCC CTGGCAAATTAGGCTCTGGAAAGA GGTCAGGATGAG
287. L($0A\overline{2}^*$): TCTTTCCAAGCC CTGGCAAATTAGGCTCTGGAAAGA GGTCAGGATGAG
288. L($0A\overline{3}^*$): TCAATCCTTGCC CTGGCAAATTAGGCTCTGGAAAGA GGTCAGGATGAG
289. L($0A\overline{4}^*$): CACATCCCTGTT CTGGCAAATTAGGCTCTGGAAAGA GGTCAGGATGAG
290. L($0A\overline{5}^*$): CCATGTCCCATT CTGGCAAATTAGGCTCTGGAAAGA GGTCAGGATGAG
291. L($0A\overline{6}^*$): CAACCAACGTTC CTGGCAAATTAGGCTCTGGAAAGA GGTCAGGATGAG
292. L($0A\overline{7}^*$): TCACACTTCGTC CTGGCAAATTAGGCTCTGGAAAGA GGTCAGGATGAG
293. $7AQ^*$: TCACTTTCGTCC TCTTTCCAGAGCCTAATTTGCCAG GGGTAGTGTGGAGAGGTGGG
294. $7A\overline{0}^*$: TCACTTTCGTCC TCTTTCCAGAGCCTAATTTGCCAG GCTGAGAAGAGG
295. $7A\overline{1}^*$: TCACTTTCGTCC TCTTTCCAGAGCCTAATTTGCCAG GGATCGGAGATG
296. $7A\overline{2}^*$: TCACTTTCGTCC TCTTTCCAGAGCCTAATTTGCCAG GGCTTGGAAAGA
297. $7A\overline{3}^*$: TCACTTTCGTCC TCTTTCCAGAGCCTAATTTGCCAG GGCAAGGATTGA
298. $7A\overline{4}^*$: TCACTTTCGTCC TCTTTCCAGAGCCTAATTTGCCAG AACAGGGATGTG
299. $7A\overline{5}^*$: TCACTTTCGTCC TCTTTCCAGAGCCTAATTTGCCAG AATGGGACATGG
300. $7A\overline{6}^*$: TCACTTTCGTCC TCTTTCCAGAGCCTAATTTGCCAG GAACGTTGGTTG
301. $7A\overline{7}^*$: TCACTTTCGTCC TCTTTCCAGAGCCTAATTTGCCAG GACGAAGTGTGA
302. L($7AQ^*$): CCCACCTCTCCACACTACCC CTGGCAAATTAGGCTCTGGAAAGA GGACGAAAGTGA
303. L($7A\overline{0}^*$): CCTCTTCTCAGC CTGGCAAATTAGGCTCTGGAAAGA GGACGAAAGTGA
304. L($7A\overline{1}^*$): CATCTCCGATCC CTGGCAAATTAGGCTCTGGAAAGA GGACGAAAGTGA
305. L($7A\overline{2}^*$): TCTTTCCAAGCC CTGGCAAATTAGGCTCTGGAAAGA GGACGAAAGTGA
306. L($7A\overline{3}^*$): TCAATCCTTGCC CTGGCAAATTAGGCTCTGGAAAGA GGACGAAAGTGA
307. L($7A\overline{4}^*$): CACATCCCTGTT CTGGCAAATTAGGCTCTGGAAAGA GGACGAAAGTGA
308. L($7A\overline{5}^*$): CCATGTCCCATT CTGGCAAATTAGGCTCTGGAAAGA GGACGAAAGTGA
309. L($7A\overline{6}^*$): CAACCAACGTTC CTGGCAAATTAGGCTCTGGAAAGA GGACGAAAGTGA
310. L($7A\overline{7}^*$): TCACACTTCGTC CTGGCAAATTAGGCTCTGGAAAGA GGACGAAAGTGA
311. $0A$: CTCATCCTGACC TCTTTCCAGAGCCTAATTTGCCAG
312. $7A$: TCACTTTCGTCC TCTTTCCAGAGCCTAATTTGCCAG
313. L($A$): CTGGCAAATTAGGCTCTGGAAAGA
314. L($0A$): CTGGCAAATTAGGCTCTGGAA AGAGGTCAGGATGAG
315. L($7A$): CTGGCAAATTAGGCTCTGGAA AGAGGACGAAAGTGA

**Renewing strands for position $D$**

316. L($\overline{0}D0^*$): CTCATCCTGACC TTTTCTGAACTGTTTAAAGCATTT GCTGAGAAGAGG
317. L($\overline{0}D1^*$): TCAACTCCGTTC TTTTCTGAACTGTTTAAAGCATTT GCTGAGAAGAGG
318. L($\overline{0}D2^*$): AATGCCACCATT TTTTCTGAACTGTTTAAAGCATTT GCTGAGAAGAGG
319. L($\overline{0}D3^*$): ACAACCCTTGTC TTTTCTGAACTGTTTAAAGCATTT GCTGAGAAGAGG
320. L($\overline{0}D4^*$): CTGTTCCCAACA TTTTCTGAACTGTTTAAAGCATTT GCTGAGAAGAGG

321. L($\bar{0}D5^*$): CACTACCAGTCC TTTTCTGAACTGTTTAAAGCATTT GCTGAGAAGAGG
322. L($\bar{0}D6^*$): ACACACACTGTC TTTTCTGAACTGTTTAAAGCATTT GCTGAGAAGAGG
323. L($\bar{0}D7^*$): TCACTTTCGTCC TTTTCTGAACTGTTTAAAGCATTT GCTGAGAAGAGG
324. L($\bar{1}D0^*$): CTCATCCTGACC TTTTCTGAACTGTTTAAAGCATTT GGATCGGAGATG
325. L($\bar{1}D1^*$): TCAACTCCGTTC TTTTCTGAACTGTTTAAAGCATTT GGATCGGAGATG
326. L($\bar{1}D2^*$): AATGCCACCATT TTTTCTGAACTGTTTAAAGCATTT GGATCGGAGATG
327. L($\bar{1}D3^*$): ACAACCCTTGTC TTTTCTGAACTGTTTAAAGCATTT GGATCGGAGATG
328. L($\bar{1}D4^*$): CTGTTCCCAACA TTTTCTGAACTGTTTAAAGCATTT GGATCGGAGATG
329. L($\bar{1}D5^*$): CACTACCAGTCC TTTTCTGAACTGTTTAAAGCATTT GGATCGGAGATG
330. L($\bar{1}D6^*$): ACACACACTGTC TTTTCTGAACTGTTTAAAGCATTT GGATCGGAGATG
331. L($\bar{1}D7^*$): TCACTTTCGTCC TTTTCTGAACTGTTTAAAGCATTT GGATCGGAGATG
332. L($\bar{1}D0^*$): CTCATCCTGACC TTTTCTGAACTGTTTAAAGCATTT GGCTTGGAAAGA
333. L($\bar{1}D1^*$): TCAACTCCGTTC TTTTCTGAACTGTTTAAAGCATTT GGCTTGGAAAGA
334. L($\bar{1}D2^*$): AATGCCACCATT TTTTCTGAACTGTTTAAAGCATTT GGCTTGGAAAGA
335. L($\bar{1}D3^*$): ACAACCCTTGTC TTTTCTGAACTGTTTAAAGCATTT GGCTTGGAAAGA
336. L($\bar{1}D4^*$): CTGTTCCCAACA TTTTCTGAACTGTTTAAAGCATTT GGCTTGGAAAGA
337. L($\bar{1}D5^*$): CACTACCAGTCC TTTTCTGAACTGTTTAAAGCATTT GGCTTGGAAAGA
338. L($\bar{1}D6^*$): ACACACACTGTC TTTTCTGAACTGTTTAAAGCATTT GGCTTGGAAAGA
339. L($\bar{1}D7^*$): TCACTTTCGTCC TTTTCTGAACTGTTTAAAGCATTT GGCTTGGAAAGA
340. L($\bar{3}D0^*$): CTCATCCTGACC TTTTCTGAACTGTTTAAAGCATTT GGCAAGGATTGA
341. L($\bar{3}D1^*$): TCAACTCCGTTC TTTTCTGAACTGTTTAAAGCATTT GGCAAGGATTGA
342. L($\bar{3}D2^*$): AATGCCACCATT TTTTCTGAACTGTTTAAAGCATTT GGCAAGGATTGA
343. L($\bar{3}D3^*$): ACAACCCTTGTC TTTTCTGAACTGTTTAAAGCATTT GGCAAGGATTGA
344. L($\bar{3}D4^*$): CTGTTCCCAACA TTTTCTGAACTGTTTAAAGCATTT GGCAAGGATTGA
345. L($\bar{3}D5^*$): CACTACCAGTCC TTTTCTGAACTGTTTAAAGCATTT GGCAAGGATTGA
346. L($\bar{3}D6^*$): ACACACACTGTC TTTTCTGAACTGTTTAAAGCATTT GGCAAGGATTGA
347. L($\bar{3}D7^*$): TCACTTTCGTCC TTTTCTGAACTGTTTAAAGCATTT GGCAAGGATTGA
348. L($\bar{4}D0^*$): CTCATCCTGACC TTTTCTGAACTGTTTAAAGCATTT AACAGGGATGTG
349. L($\bar{4}D1^*$): TCAACTCCGTTC TTTTCTGAACTGTTTAAAGCATTT AACAGGGATGTG
350. L($\bar{4}D2^*$): AATGCCACCATT TTTTCTGAACTGTTTAAAGCATTT AACAGGGATGTG
351. L($\bar{4}D3^*$): ACAACCCTTGTC TTTTCTGAACTGTTTAAAGCATTT AACAGGGATGTG
352. L($\bar{4}D4^*$): CTGTTCCCAACA TTTTCTGAACTGTTTAAAGCATTT AACAGGGATGTG
353. L($\bar{4}D5^*$): CACTACCAGTCC TTTTCTGAACTGTTTAAAGCATTT AACAGGGATGTG
354. L($\bar{4}D6^*$): ACACACACTGTC TTTTCTGAACTGTTTAAAGCATTT AACAGGGATGTG
355. L($\bar{4}D7^*$): TCACTTTCGTCC TTTTCTGAACTGTTTAAAGCATTT AACAGGGATGTG
356. L($\bar{5}D0^*$): CTCATCCTGACC TTTTCTGAACTGTTTAAAGCATTT AATGGGACATGG
357. L($\bar{5}D1^*$): TCAACTCCGTTC TTTTCTGAACTGTTTAAAGCATTT AATGGGACATGG
358. L($\bar{5}D2^*$): AATGCCACCATT TTTTCTGAACTGTTTAAAGCATTT AATGGGACATGG
359. L($\bar{5}D3^*$): ACAACCCTTGTC TTTTCTGAACTGTTTAAAGCATTT AATGGGACATGG
360. L($\bar{5}D4^*$): CTGTTCCCAACA TTTTCTGAACTGTTTAAAGCATTT AATGGGACATGG
361. L($\bar{5}D5^*$): CACTACCAGTCC TTTTCTGAACTGTTTAAAGCATTT AATGGGACATGG
362. L($\bar{5}D6^*$): ACACACACTGTC TTTTCTGAACTGTTTAAAGCATTT AATGGGACATGG
363. L($\bar{5}D7^*$): TCACTTTCGTCC TTTTCTGAACTGTTTAAAGCATTT AATGGGACATGG
364. L($\bar{6}D0^*$): CTCATCCTGACC TTTTCTGAACTGTTTAAAGCATTT GAACGTTGGTTG
365. L($\bar{6}D1^*$): TCAACTCCGTTC TTTTCTGAACTGTTTAAAGCATTT GAACGTTGGTTG
366. L($\bar{6}D2^*$): AATGCCACCATT TTTTCTGAACTGTTTAAAGCATTT GAACGTTGGTTG
367. L($\bar{6}D3^*$): ACAACCCTTGTC TTTTCTGAACTGTTTAAAGCATTT GAACGTTGGTTG
368. L($\bar{6}D4^*$): CTGTTCCCAACA TTTTCTGAACTGTTTAAAGCATTT GAACGTTGGTTG
369. L($\bar{6}D5^*$): CACTACCAGTCC TTTTCTGAACTGTTTAAAGCATTT GAACGTTGGTTG
370. L($\bar{6}D6^*$): ACACACACTGTC TTTTCTGAACTGTTTAAAGCATTT GAACGTTGGTTG
371. L($\bar{6}D7^*$): TCACTTTCGTCC TTTTCTGAACTGTTTAAAGCATTT GAACGTTGGTTG
372. L($\bar{7}D0^*$): CTCATCCTGACC TTTTCTGAACTGTTTAAAGCATTT GACGAAGTGTGA
373. L($\bar{7}D1^*$): TCAACTCCGTTC TTTTCTGAACTGTTTAAAGCATTT GACGAAGTGTGA
374. L($\bar{7}D2^*$): AATGCCACCATT TTTTCTGAACTGTTTAAAGCATTT GACGAAGTGTGA

375. L($\overline{7}D3^*$): ACAACCCTTGTC TTTTCTGAACTGTTTAAAGCATTT GACGAAGTGTGA
376. L($\overline{7}D4^*$): CTGTTCCCAACA TTTTCTGAACTGTTTAAAGCATTT GACGAAGTGTGA
377. L($\overline{7}D5^*$): CACTACCAGTCC TTTTCTGAACTGTTTAAAGCATTT GACGAAGTGTGA
378. L($\overline{7}D6^*$): ACACACACTGTC TTTTCTGAACTGTTTAAAGCATTT GACGAAGTGTGA
379. L($\overline{7}D7^*$): TCACTTTCGTCC TTTTCTGAACTGTTTAAAGCATTT GACGAAGTGTGA
380. L($\overline{0}DQ^*$): CCCACCTCTCCACACTACCC TTTTCTGAACTGTTTAAAGCATTT GCTGAGAAGAGG
381. L($\overline{1}DQ^*$): CCCACCTCTCCACACTACCC TTTTCTGAACTGTTTAAAGCATTT GGATCGGAGATG
382. L($\overline{1}DQ^*$): CCCACCTCTCCACACTACCC TTTTCTGAACTGTTTAAAGCATTT GGCTTGGAAAGA
383. L($\overline{3}DQ^*$): CCCACCTCTCCACACTACCC TTTTCTGAACTGTTTAAAGCATTT GGCAAGGATTGA
384. L($\overline{4}DQ^*$): CCCACCTCTCCACACTACCC TTTTCTGAACTGTTTAAAGCATTT AACAGGGATGTG
385. L($\overline{5}DQ^*$): CCCACCTCTCCACACTACCC TTTTCTGAACTGTTTAAAGCATTT AATGGGACATGG
386. L($\overline{6}DQ^*$): CCCACCTCTCCACACTACCC TTTTCTGAACTGTTTAAAGCATTT GAACGTTGGTTG
387. L($\overline{7}DQ^*$): CCCACCTCTCCACACTACCC TTTTCTGAACTGTTTAAAGCATTT GACGAAGTGTGA
388. L($\overline{0}D$): TTTTCTGAACTGTTTAAAGCATTT GCTGAGAAGAGG
389. L($\overline{1}D$): TTTTCTGAACTGTTTAAAGCATTT GGATCGGAGATG
390. L($\overline{1}D$): TTTTCTGAACTGTTTAAAGCATTT GGCTTGGAAAGA
391. L($\overline{3}D$): TTTTCTGAACTGTTTAAAGCATTT GGCAAGGATTGA
392. L($\overline{4}D$): TTTTCTGAACTGTTTAAAGCATTT AACAGGGATGTG
393. L($\overline{5}D$): TTTTCTGAACTGTTTAAAGCATTT AATGGGACATGG
394. L($\overline{6}D$): TTTTCTGAACTGTTTAAAGCATTT GAACGTTGGTTG
395. L($\overline{7}D$): TTTTCTGAACTGTTTAAAGCATTT GACGAAGTGTGA

#### S10.2 DNA sequences: 11-position programs

##### S10.2.1 288nt Subsequence of the M13 Scaffold DNA sequence

396. scaff-288: $L^*$ $K^*$ $J^*$ $I^*$ $H^*$ $G^*$ $F^*$ $E^*$ $D^*$ $C^*$ $B^*$ $A^*$:
GCAAAAATGACCTCTTATCAAAAG GAGCAATTAAAGGTACTCTCTAAT CCTGACCTGTTGGAGTTTGCTTCC
GGTCTGGTTCGCTTTGAAGCTCGA ATTAAAACGCGATATTTGAAGTCT TTCGGGCTTCCTCTTAATCTTTTT
GATGCAATCCGCTTTGCTTCTGAC TATAATAGTCAGGGTAAAGACCTG ATTTTTGATTTATGGTCATTCTCG
TTTTCTGAACTGTTTAAAGCATTT GAGGGGGATTCAATGAATATTTAT GACGATTCCGCAGTATTGGACGCT

##### S10.2.2 11-position DNA sequences

397. $A$: TCTTTCCAGAGCCTAATTTGCCAG
398. $A\overline{0}^*$: TCTTTCCAGAGCCTAATTTGCCAG GCTGAGAAGAGG
399. $A\overline{2}^*$: TCTTTCCAGAGCCTAATTTGCCAG GGCTTGGAAAGA
400. $A\mathtt{Q}^*$: TCTTTCCAGAGCCTAATTTGCCAG GGGTAGTGTGGAGAGGTGGG
401. $B$: AGCGTCCAATACTGCGGAATCGTC
402. $B0^*$: AGCGTCCAATACTGCGGAATCGTC GGTCAGGATGAG
403. $B2^*$: AGCGTCCAATACTGCGGAATCGTC AATGGTGGCATT
404. $\overline{0}B0^*$: CCTCTTCTCAGC AGCGTCCAATACTGCGGAATCGTC GGTCAGGATGAG
405. $\overline{2}B2^*$: TCTTTCCAAGCC AGCGTCCAATACTGCGGAATCGTC AATGGTGGCATT
406. $\overline{0}B$: CCTCTTCTCAGC AGCGTCCAATACTGCGGAATCGTC
407. $\overline{2}B$: TCTTTCCAAGCC AGCGTCCAATACTGCGGAATCGTC
408. $\overline{0}B\mathtt{Q}^*$: CCTCTTCTCAGC AGCGTCCAATACTGCGGAATCGTC GGGTAGTGTGGAGAGGTGGG
409. $\overline{2}B\mathtt{Q}^*$: TCTTTCCAAGCC AGCGTCCAATACTGCGGAATCGTC GGGTAGTGTGGAGAGGTGGG
410. $\mathtt{RF}^*B$: TTGGGATGCGAAGGGATGGT AGCGTCCAATACTGCGGAATCGTC
411. $C$: ATAAATATTCATTGAATCCCCCTC
412. $C\overline{0}^*$: ATAAATATTCATTGAATCCCCCTC GCTGAGAAGAGG
413. $C\overline{2}^*$: ATAAATATTCATTGAATCCCCCTC GGCTTGGAAAGA
414. $0C\overline{0}^*$: CTCATCCTGACC ATAAATATTCATTGAATCCCCCTC GCTGAGAAGAGG
415. $2C\overline{2}^*$: AATGCCACCATT ATAAATATTCATTGAATCCCCCTC GGCTTGGAAAGA
416. $0C$: CTCATCCTGACC ATAAATATTCATTGAATCCCCCTC
417. $2C$: AATGCCACCATT ATAAATATTCATTGAATCCCCCTC

418. $0C\mathtt{Q}^*$: CTCATCCTGACC ATAAATATTCATTGAATCCCCCTC GGGTAGTGTGGAGAGGTGGG
419. $2C\mathtt{Q}^*$: AATGCCACCATT ATAAATATTCATTGAATCCCCCTC GGGTAGTGTGGAGAGGTGGG
420. $\mathtt{RF}^*C$: TTGGGATGCGAAGGGATGGT ATAAATATTCATTGAATCCCCCTC
421. $D$: AAATGCTTTAAACAGTTCAGAAAA
422. $D0^*$: AAATGCTTTAAACAGTTCAGAAAA GGTCAGGATGAG
423. $D2^*$: AAATGCTTTAAACAGTTCAGAAAA AATGGTGGCATT
424. $\overline{0}D0^*$: CCTCTTCTCAGC AAATGCTTTAAACAGTTCAGAAAA GGTCAGGATGAG
425. $\overline{2}D2^*$: TCTTTCCAAGCC AAATGCTTTAAACAGTTCAGAAAA AATGGTGGCATT
426. $\overline{0}D$: CCTCTTCTCAGC AAATGCTTTAAACAGTTCAGAAAA
427. $\overline{2}D$: TCTTTCCAAGCC AAATGCTTTAAACAGTTCAGAAAA
428. $\overline{0}D\mathtt{Q}^*$: CCTCTTCTCAGC AAATGCTTTAAACAGTTCAGAAAA GGGTAGTGTGGAGAGGTGGG
429. $\overline{2}D\mathtt{Q}^*$: TCTTTCCAAGCC AAATGCTTTAAACAGTTCAGAAAA GGGTAGTGTGGAGAGGTGGG
430. $\mathtt{RF}^*D$: TTGGGATGCGAAGGGATGGT AAATGCTTTAAACAGTTCAGAAAA
431. $E$: CGAGAATGACCATAAATCAAAAAT
432. $E\overline{0}^*$: CGAGAATGACCATAAATCAAAAAT GCTGAGAAGAGG
433. $E\overline{2}^*$: CGAGAATGACCATAAATCAAAAAT GGCTTGGAAAGA
434. $0E\overline{0}^*$: CTCATCCTGACC CGAGAATGACCATAAATCAAAAAT GCTGAGAAGAGG
435. $2E\overline{2}^*$: AATGCCACCATT CGAGAATGACCATAAATCAAAAAT GGCTTGGAAAGA
436. $0E$: CTCATCCTGACC CGAGAATGACCATAAATCAAAAAT
437. $2E$: AATGCCACCATT CGAGAATGACCATAAATCAAAAAT
438. $0E\mathtt{Q}^*$: CTCATCCTGACC CGAGAATGACCATAAATCAAAAAT GGGTAGTGTGGAGAGGTGGG
439. $2E\mathtt{Q}^*$: AATGCCACCATT CGAGAATGACCATAAATCAAAAAT GGGTAGTGTGGAGAGGTGGG
440. $\mathtt{RF}^*E$: TTGGGATGCGAAGGGATGGT CGAGAATGACCATAAATCAAAAAT
441. $f$: CAGGTCTTTACCCTGACTATTATA
442. $f0^*$: CAGGTCTTTACCCTGACTATTATA GGTCAGGATGAG
443. $f2^*$: CAGGTCTTTACCCTGACTATTATA AATGGTGGCATT
444. $\overline{0}f$: CCTCTTCTCAGC CAGGTCTTTACCCTGACTATTATA
445. $\overline{2}f$: TCTTTCCAAGCC CAGGTCTTTACCCTGACTATTATA
446. $\overline{0}f0^*$: CCTCTTCTCAGC CAGGTCTTTACCCTGACTATTATA GGTCAGGATGAG
447. $\overline{2}f2^*$: TCTTTCCAAGCC CAGGTCTTTACCCTGACTATTATA AATGGTGGCATT
448. $\overline{0}f2^*$: CCTCTTCTCAGC CAGGTCTTTACCCTGACTATTATA AATGGTGGCATT
449. $\overline{2}f0^*$: TCTTTCCAAGCC CAGGTCTTTACCCTGACTATTATA GGTCAGGATGAG
450. $\overline{0}f\mathtt{Q}^*$: CCTCTTCTCAGC CAGGTCTTTACCCTGACTATTATA GGGTAGTGTGGAGAGGTGGG
451. $\overline{2}f\mathtt{Q}^*$: TCTTTCCAAGCC CAGGTCTTTACCCTGACTATTATA GGGTAGTGTGGAGAGGTGGG
452. $F$: GTCAGAAGCAAAGCGGATTGCATC
453. $F\overline{0}^*$: GTCAGAAGCAAAGCGGATTGCATC GCTGAGAAGAGG
454. $F\overline{2}^*$: GTCAGAAGCAAAGCGGATTGCATC GGCTTGGAAAGA
455. $0F$: CTCATCCTGACC GTCAGAAGCAAAGCGGATTGCATC
456. $2F$: AATGCCACCATT GTCAGAAGCAAAGCGGATTGCATC
457. $0F\overline{0}^*$: CTCATCCTGACC GTCAGAAGCAAAGCGGATTGCATC GCTGAGAAGAGG
458. $2F\overline{2}^*$: AATGCCACCATT GTCAGAAGCAAAGCGGATTGCATC GGCTTGGAAAGA
459. $0F\mathtt{Q}^*$: CTCATCCTGACC GTCAGAAGCAAAGCGGATTGCATC GGGTAGTGTGGAGAGGTGGG
460. $2F\mathtt{Q}^*$: AATGCCACCATT GTCAGAAGCAAAGCGGATTGCATC GGGTAGTGTGGAGAGGTGGG
461. $\mathtt{RF}^*F$: TTGGGATGCGAAGGGATGGT GTCAGAAGCAAAGCGGATTGCATC
462. $G$: AAAAAGATTAAGAGGAAGCCCGAA
463. $G0^*$: AAAAAGATTAAGAGGAAGCCCGAA GGTCAGGATGAG
464. $G2^*$: AAAAAGATTAAGAGGAAGCCCGAA AATGGTGGCATT
465. $\overline{0}G$: CCTCTTCTCAGC AAAAAGATTAAGAGGAAGCCCGAA
466. $\overline{2}G$: TCTTTCCAAGCC AAAAAGATTAAGAGGAAGCCCGAA
467. $\overline{0}G0^*$: CCTCTTCTCAGC AAAAAGATTAAGAGGAAGCCCGAA GGTCAGGATGAG
468. $\overline{2}G2^*$: TCTTTCCAAGCC AAAAAGATTAAGAGGAAGCCCGAA AATGGTGGCATT
469. $\overline{0}G\mathtt{Q}^*$: CCTCTTCTCAGC AAAAAGATTAAGAGGAAGCCCGAA GGGTAGTGTGGAGAGGTGGG
470. $\overline{2}G\mathtt{Q}^*$: TCTTTCCAAGCC AAAAAGATTAAGAGGAAGCCCGAA GGGTAGTGTGGAGAGGTGGG
471. $\mathtt{RF}^*G$: TTGGGATGCGAAGGGATGGT AAAAAGATTAAGAGGAAGCCCGAA

472. $H$: AGACTTCAAATATCGCGTTTTAAT
473. $H\overline{0}^*$: AGACTTCAAATATCGCGTTTTAAT GCTGAGAAGAGG
474. $H\overline{2}^*$: AGACTTCAAATATCGCGTTTTAAT GGCTTGGAAAGA
475. $0H$: CTCATCCTGACC AGACTTCAAATATCGCGTTTTAAT
476. $2H$: AATGCCACCATT AGACTTCAAATATCGCGTTTTAAT
477. $0H\overline{0}^*$: CTCATCCTGACC AGACTTCAAATATCGCGTTTTAAT GCTGAGAAGAGG
478. $2H\overline{2}^*$: AATGCCACCATT AGACTTCAAATATCGCGTTTTAAT GGCTTGGAAAGA
479. $0H\mathtt{Q}^*$: CTCATCCTGACC AGACTTCAAATATCGCGTTTTAAT GGGTAGTGTGGAGAGGTGGG
480. $2H\mathtt{Q}^*$: AATGCCACCATT AGACTTCAAATATCGCGTTTTAAT GGGTAGTGTGGAGAGGTGGG
481. $\mathtt{RF}^*H$: TTGGGATGCGAAGGGATGGT AGACTTCAAATATCGCGTTTTAAT
482. $I$: TCGAGCTTCAAAGCGAACCAGACC
483. $I0^*$: TCGAGCTTCAAAGCGAACCAGACC GGTCAGGATGAG
484. $I2^*$: TCGAGCTTCAAAGCGAACCAGACC AATGGTGGCATT
485. $\overline{0}I$: CCTCTTCTCAGC TCGAGCTTCAAAGCGAACCAGACC
486. $\overline{2}I$: TCTTTCCAAGCC TCGAGCTTCAAAGCGAACCAGACC
487. $\overline{0}I0^*$: CCTCTTCTCAGC TCGAGCTTCAAAGCGAACCAGACC GGTCAGGATGAG
488. $\overline{2}I2^*$: TCTTTCCAAGCC TCGAGCTTCAAAGCGAACCAGACC AATGGTGGCATT
489. $\overline{0}I\mathtt{Q}^*$: CCTCTTCTCAGC TCGAGCTTCAAAGCGAACCAGACC GGGTAGTGTGGAGAGGTGGG
490. $\overline{2}I\mathtt{Q}^*$: TCTTTCCAAGCC TCGAGCTTCAAAGCGAACCAGACC GGGTAGTGTGGAGAGGTGGG
491. $\mathtt{RF}^*I$: TTGGGATGCGAAGGGATGGT TCGAGCTTCAAAGCGAACCAGACC
492. $J$: GGAAGCAAACTCCAACAGGTCAGG
493. $J\overline{0}^*$: GGAAGCAAACTCCAACAGGTCAGG GCTGAGAAGAGG
494. $J\overline{2}^*$: GGAAGCAAACTCCAACAGGTCAGG GGCTTGGAAAGA
495. $0J$: CTCATCCTGACC GGAAGCAAACTCCAACAGGTCAGG
496. $2J$: AATGCCACCATT GGAAGCAAACTCCAACAGGTCAGG
497. $0J\overline{0}^*$: CTCATCCTGACC GGAAGCAAACTCCAACAGGTCAGG GCTGAGAAGAGG
498. $2J\overline{2}^*$: AATGCCACCATT GGAAGCAAACTCCAACAGGTCAGG GGCTTGGAAAGA
499. $0J\mathtt{Q}^*$: CTCATCCTGACC GGAAGCAAACTCCAACAGGTCAGG GGGTAGTGTGGAGAGGTGGG
500. $2J\mathtt{Q}^*$: AATGCCACCATT GGAAGCAAACTCCAACAGGTCAGG GGGTAGTGTGGAGAGGTGGG
501. $\mathtt{RF}^*J$: TTGGGATGCGAAGGGATGGT GGAAGCAAACTCCAACAGGTCAGG
502. $K$: ATTAGAGAGTACCTTTAATTGCTC
503. $K0^*$: ATTAGAGAGTACCTTTAATTGCTC GGTCAGGATGAG
504. $K2^*$: ATTAGAGAGTACCTTTAATTGCTC AATGGTGGCATT
505. $\overline{0}K$: CCTCTTCTCAGC ATTAGAGAGTACCTTTAATTGCTC
506. $\overline{2}K$: TCTTTCCAAGCC ATTAGAGAGTACCTTTAATTGCTC
507. $\overline{0}K0^*$: CCTCTTCTCAGC ATTAGAGAGTACCTTTAATTGCTC GGTCAGGATGAG
508. $\overline{2}K2^*$: TCTTTCCAAGCC ATTAGAGAGTACCTTTAATTGCTC AATGGTGGCATT
509. $\overline{0}K\mathtt{Q}^*$: CCTCTTCTCAGC ATTAGAGAGTACCTTTAATTGCTC GGGTAGTGTGGAGAGGTGGG
510. $\overline{2}K\mathtt{Q}^*$: TCTTTCCAAGCC ATTAGAGAGTACCTTTAATTGCTC GGGTAGTGTGGAGAGGTGGG
511. $\mathtt{RF}^*K$: TTGGGATGCGAAGGGATGGT ATTAGAGAGTACCTTTAATTGCTC
512. $\mathtt{RF}^*L$: TTGGGATGCGAAGGGATGGT CTTTTGATAAGAGGTCATTTTTGC
513. $F_{\mathtt{w}}$: GTCAGAAGCAAATCGGATTGCATC
514. $F_{\mathtt{w}}\overline{0}^*$: GTCAGAAGCAAATCGGATTGCATC GCTGAGAAGAGG
515. $F_{\mathtt{w}}\overline{2}^*$: GTCAGAAGCAAATCGGATTGCATC GGCTTGGAAAGA
516. $0F_{\mathtt{w}}$: CTCATCCTGACC GTCAGAAGCAAATCGGATTGCATC
517. $2F_{\mathtt{w}}$: AATGCCACCATT GTCAGAAGCAAATCGGATTGCATC
518. $0F_{\mathtt{w}}\overline{0}^*$: CTCATCCTGACC GTCAGAAGCAAATCGGATTGCATC GCTGAGAAGAGG
519. $2F_{\mathtt{w}}\overline{2}^*$: AATGCCACCATT GTCAGAAGCAAATCGGATTGCATC GGCTTGGAAAGA
520. $0F_{\mathtt{w}}\mathtt{Q}^*$: CTCATCCTGACC GTCAGAAGCAAATCGGATTGCATC GGGTAGTGTGGAGAGGTGGG
521. $2F_{\mathtt{w}}\mathtt{Q}^*$: AATGCCACCATT GTCAGAAGCAAATCGGATTGCATC GGGTAGTGTGGAGAGGTGGG
522. $G_{\mathtt{w}}$: AAAAAGATTAAGCGGAAGCCCGAA
523. $G_{\mathtt{w}}0^*$: AAAAAGATTAAGCGGAAGCCCGAA GGTCAGGATGAG
524. $G_{\mathtt{w}}2^*$: AAAAAGATTAAGCGGAAGCCCGAA AATGGTGGCATT
525. $\overline{0}G_{\mathtt{w}}$: CCTCTTCTCAGC AAAAAGATTAAGCGGAAGCCCGAA

526. $\overline{2}G_{\mathtt{w}}$: TCTTTCCAAGCC AAAAAGATTAAGCGGAAGCCCGAA
527. $\overline{0}G_{\mathtt{w}}0^*$: CCTCTTCTCAGC AAAAAGATTAAGCGGAAGCCCGAA GGTCAGGATGAG
528. $\overline{2}G_{\mathtt{w}}2^*$: TCTTTCCAAGCC AAAAAGATTAAGCGGAAGCCCGAA AATGGTGGCATT
529. $\overline{0}G_{\mathtt{w}}\mathtt{Q}^*$: CCTCTTCTCAGC AAAAAGATTAAGCGGAAGCCCGAA GGGTAGTGTGGAGAGGTGGG
530. $\overline{2}G_{\mathtt{w}}\mathtt{Q}^*$: TCTTTCCAAGCC AAAAAGATTAAGCGGAAGCCCGAA GGGTAGTGTGGAGAGGTGGG
531. $I_{\mathtt{w}}$: TCGAGCTTCAAATCGAACCAGACC
532. $I_{\mathtt{w}}0^*$: TCGAGCTTCAAATCGAACCAGACC GGTCAGGATGAG
533. $I_{\mathtt{w}}2^*$: TCGAGCTTCAAATCGAACCAGACC AATGGTGGCATT
534. $\overline{0}I_{\mathtt{w}}$: CCTCTTCTCAGC TCGAGCTTCAAATCGAACCAGACC
535. $\overline{2}I_{\mathtt{w}}$: TCTTTCCAAGCC TCGAGCTTCAAATCGAACCAGACC
536. $\overline{0}I_{\mathtt{w}}0^*$: CCTCTTCTCAGC TCGAGCTTCAAATCGAACCAGACC GGTCAGGATGAG
537. $\overline{2}I_{\mathtt{w}}2^*$: TCTTTCCAAGCC TCGAGCTTCAAATCGAACCAGACC AATGGTGGCATT
538. $\overline{0}I_{\mathtt{w}}\mathtt{Q}^*$: CCTCTTCTCAGC TCGAGCTTCAAATCGAACCAGACC GGGTAGTGTGGAGAGGTGGG
539. $\overline{2}I_{\mathtt{w}}\mathtt{Q}^*$: TCTTTCCAAGCC TCGAGCTTCAAATCGAACCAGACC GGGTAGTGTGGAGAGGTGGG
540. $J_{\mathtt{w}}$: GGAAGCAAACTCAAACAGGTCAGG
541. $J_{\mathtt{w}}\overline{0}^*$: GGAAGCAAACTCAAACAGGTCAGG GCTGAGAAGAGG
542. $J_{\mathtt{w}}\overline{2}^*$: GGAAGCAAACTCAAACAGGTCAGG GGCTTGGAAAGA
543. $0J_{\mathtt{w}}$: CTCATCCTGACC GGAAGCAAACTCAAACAGGTCAGG
544. $2J_{\mathtt{w}}$: AATGCCACCATT GGAAGCAAACTCAAACAGGTCAGG
545. $0J_{\mathtt{w}}\overline{0}^*$: CTCATCCTGACC GGAAGCAAACTCAAACAGGTCAGG GCTGAGAAGAGG
546. $2J_{\mathtt{w}}\overline{2}^*$: AATGCCACCATT GGAAGCAAACTCAAACAGGTCAGG GGCTTGGAAAGA
547. $0J_{\mathtt{w}}\mathtt{Q}^*$: CTCATCCTGACC GGAAGCAAACTCAAACAGGTCAGG GGGTAGTGTGGAGAGGTGGG
548. $2J_{\mathtt{w}}\mathtt{Q}^*$: AATGCCACCATT GGAAGCAAACTCAAACAGGTCAGG GGGTAGTGTGGAGAGGTGGG
549. $F_{\mathtt{w14A}}$: GTCAGAAGCAAAGCAGATTGCATC
550. $F_{\mathtt{w14A}}\overline{0}^*$: GTCAGAAGCAAAGCAGATTGCATC GCTGAGAAGAGG
551. $F_{\mathtt{w14A}}\overline{2}^*$: GTCAGAAGCAAAGCAGATTGCATC GGCTTGGAAAGA
552. $0F_{\mathtt{w14A}}$: CTCATCCTGACC GTCAGAAGCAAAGCAGATTGCATC
553. $2F_{\mathtt{w14A}}$: AATGCCACCATT GTCAGAAGCAAAGCAGATTGCATC
554. $0F_{\mathtt{w14A}}\overline{0}^*$: CTCATCCTGACC GTCAGAAGCAAAGCAGATTGCATC GCTGAGAAGAGG
555. $2F_{\mathtt{w14A}}\overline{2}^*$: AATGCCACCATT GTCAGAAGCAAAGCAGATTGCATC GGCTTGGAAAGA
556. $0F_{\mathtt{w14A}}\mathtt{Q}^*$: CTCATCCTGACC GTCAGAAGCAAAGCAGATTGCATC GGGTAGTGTGGAGAGGTGGG
557. $2F_{\mathtt{w14A}}\mathtt{Q}^*$: AATGCCACCATT GTCAGAAGCAAAGCAGATTGCATC GGGTAGTGTGGAGAGGTGGG
558. $I_{\mathtt{w13T}}$: TCGAGCTTCAAAGTGAACCAGACC
559. $I_{\mathtt{w13T}}0^*$: TCGAGCTTCAAAGTGAACCAGACC GGTCAGGATGAG
560. $I_{\mathtt{w13T}}2^*$: TCGAGCTTCAAAGTGAACCAGACC AATGGTGGCATT
561. $\overline{0}I_{\mathtt{w13T}}$: CCTCTTCTCAGC TCGAGCTTCAAAGTGAACCAGACC
562. $\overline{2}I_{\mathtt{w13T}}$: TCTTTCCAAGCC TCGAGCTTCAAAGTGAACCAGACC
563. $\overline{0}I_{\mathtt{w13T}}0^*$: CCTCTTCTCAGC TCGAGCTTCAAAGTGAACCAGACC GGTCAGGATGAG
564. $\overline{2}I_{\mathtt{w13T}}2^*$: TCTTTCCAAGCC TCGAGCTTCAAAGTGAACCAGACC AATGGTGGCATT
565. $\overline{0}I_{\mathtt{w13T}}\mathtt{Q}^*$: CCTCTTCTCAGC TCGAGCTTCAAAGTGAACCAGACC GGGTAGTGTGGAGAGGTGGG
566. $\overline{2}I_{\mathtt{w13T}}\mathtt{Q}^*$: TCTTTCCAAGCC TCGAGCTTCAAAGTGAACCAGACC GGGTAGTGTGGAGAGGTGGG
567. $J_{\mathtt{w14G}}$: GGAAGCAAACTCCAGCAGGTCAGG
568. $J_{\mathtt{w14G}}\overline{0}^*$: GGAAGCAAACTCCAGCAGGTCAGG GCTGAGAAGAGG
569. $J_{\mathtt{w14G}}\overline{2}^*$: GGAAGCAAACTCCAGCAGGTCAGG GGCTTGGAAAGA
570. $0J_{\mathtt{w14G}}$: CTCATCCTGACC GGAAGCAAACTCCAGCAGGTCAGG
571. $2J_{\mathtt{w14G}}$: AATGCCACCATT GGAAGCAAACTCCAGCAGGTCAGG
572. $0J_{\mathtt{w14G}}\overline{0}^*$: CTCATCCTGACC GGAAGCAAACTCCAGCAGGTCAGG GCTGAGAAGAGG
573. $2J_{\mathtt{w14G}}\overline{2}^*$: AATGCCACCATT GGAAGCAAACTCCAGCAGGTCAGG GGCTTGGAAAGA
574. $0J_{\mathtt{w14G}}\mathtt{Q}^*$: CTCATCCTGACC GGAAGCAAACTCCAGCAGGTCAGG GGGTAGTGTGGAGAGGTGGG
575. $2J_{\mathtt{w14G}}\mathtt{Q}^*$: AATGCCACCATT GGAAGCAAACTCCAGCAGGTCAGG GGGTAGTGTGGAGAGGTGGG
576. $I_{\mathtt{w13Tw9G}}$: TCGAGCTTCGAAGTGAACCAGACC
577. $I_{\mathtt{w13Tw9G}}0^*$: TCGAGCTTCGAAGTGAACCAGACC GGTCAGGATGAG
578. $I_{\mathtt{w13Tw9G}}2^*$: TCGAGCTTCGAAGTGAACCAGACC AATGGTGGCATT
579. $\overline{0}I_{\mathtt{w13Tw9G}}$: CCTCTTCTCAGC TCGAGCTTCGAAGTGAACCAGACC

580. $\overline{2}I_{\mathtt{w13Tw9G}}$: TCTTTCCAAGCC TCGAGCTTCGAAGTGAACCAGACC
581. $\overline{0}I_{\mathtt{w13Tw9G}}0^*$: CCTCTTCTCAGC TCGAGCTTCGAAGTGAACCAGACC GGTCAGGATGAG
582. $\overline{2}I_{\mathtt{w13Tw9G}}2^*$: TCTTTCCAAGCC TCGAGCTTCGAAGTGAACCAGACC AATGGTGGCATT
583. $\overline{0}I_{\mathtt{w13Tw9G}}\mathtt{Q}^*$: CCTCTTCTCAGC TCGAGCTTCGAAGTGAACCAGACC GGGTAGTGTGGAGAGGTGGG
584. $\overline{2}I_{\mathtt{w13Tw9G}}\mathtt{Q}^*$: TCTTTCCAAGCC TCGAGCTTCGAAGTGAACCAGACC GGGTAGTGTGGAGAGGTGGG
585. $J_{\mathtt{w14Gw12T}}$: GGAAGCAAACTCTAGCAGGTCAGG
586. $J_{\mathtt{w14Gw12T}}\overline{0}^*$: GGAAGCAAACTCTAGCAGGTCAGG GCTGAGAAGAGG
587. $J_{\mathtt{w14Gw12T}}\overline{2}^*$: GGAAGCAAACTCTAGCAGGTCAGG GGCTTGGAAAGA
588. $0J_{\mathtt{w14Gw12T}}$: CTCATCCTGACC GGAAGCAAACTCTAGCAGGTCAGG
589. $2J_{\mathtt{w14Gw12T}}$: AATGCCACCATT GGAAGCAAACTCTAGCAGGTCAGG
590. $0J_{\mathtt{w14Gw12T}}\overline{0}^*$: CTCATCCTGACC GGAAGCAAACTCTAGCAGGTCAGG GCTGAGAAGAGG
591. $2J_{\mathtt{w14Gw12T}}\overline{2}^*$: AATGCCACCATT GGAAGCAAACTCTAGCAGGTCAGG GGCTTGGAAAGA
592. $0J_{\mathtt{w14Gw12T}}\mathtt{Q}^*$: CTCATCCTGACC GGAAGCAAACTCTAGCAGGTCAGG GGGTAGTGTGGAGAGGTGGG
593. $2J_{\mathtt{w14Gw12T}}\mathtt{Q}^*$: AATGCCACCATT GGAAGCAAACTCTAGCAGGTCAGG GGGTAGTGTGGAGAGGTGGG
594. $I_{\mathtt{w13Tw12C}}$: TCGAGCTTCAAACTGAACCAGACC
595. $I_{\mathtt{w13Tw12G}}0^*$: TCGAGCTTCAAACTGAACCAGACC GGTCAGGATGAG
596. $I_{\mathtt{w13Tw12G}}2^*$: TCGAGCTTCAAACTGAACCAGACC AATGGTGGCATT
597. $\overline{0}I_{\mathtt{w13Tw12G}}$: CCTCTTCTCAGC TCGAGCTTCAAACTGAACCAGACC
598. $\overline{2}I_{\mathtt{w13Tw12G}}$: TCTTTCCAAGCC TCGAGCTTCAAACTGAACCAGACC
599. $\overline{0}I_{\mathtt{w13Tw12G}}0^*$: CCTCTTCTCAGC TCGAGCTTCAAACTGAACCAGACC GGTCAGGATGAG
600. $\overline{2}I_{\mathtt{w13Tw12G}}2^*$: TCTTTCCAAGCC TCGAGCTTCAAACTGAACCAGACC AATGGTGGCATT
601. $\overline{0}I_{\mathtt{w13Tw12G}}\mathtt{Q}^*$: CCTCTTCTCAGC TCGAGCTTCAAACTGAACCAGACC GGGTAGTGTGGAGAGGTGGG
602. $\overline{2}I_{\mathtt{w13Tw12G}}\mathtt{Q}^*$: TCTTTCCAAGCC TCGAGCTTCAAACTGAACCAGACC GGGTAGTGTGGAGAGGTGGG
603. $J_{\mathtt{w14Gw13T}}$: GGAAGCAAACTCCTGCAGGTCAGG
604. $J_{\mathtt{w14Gw13T}}\overline{0}^*$: GGAAGCAAACTCCTGCAGGTCAGG GCTGAGAAGAGG
605. $J_{\mathtt{w14Gw13T}}\overline{2}^*$: GGAAGCAAACTCCTGCAGGTCAGG GGCTTGGAAAGA
606. $0J_{\mathtt{w14Gw13T}}$: CTCATCCTGACC GGAAGCAAACTCCTGCAGGTCAGG
607. $2J_{\mathtt{w14Gw13T}}$: AATGCCACCATT GGAAGCAAACTCCTGCAGGTCAGG
608. $0J_{\mathtt{w14Gw13T}}\overline{0}^*$: CTCATCCTGACC GGAAGCAAACTCCTGCAGGTCAGG GCTGAGAAGAGG
609. $2J_{\mathtt{w14Gw13T}}\overline{2}^*$: AATGCCACCATT GGAAGCAAACTCCTGCAGGTCAGG GGCTTGGAAAGA
610. $0J_{\mathtt{w14Gw13T}}\mathtt{Q}^*$: CTCATCCTGACC GGAAGCAAACTCCTGCAGGTCAGG GGGTAGTGTGGAGAGGTGGG
611. $2J_{\mathtt{w14Gw13T}}\mathtt{Q}^*$: AATGCCACCATT GGAAGCAAACTCCTGCAGGTCAGG GGGTAGTGTGGAGAGGTGGG

#### S10.3 DNA sequences: 25-position programs

##### S10.3.1 624nt Subsequence of the M13 Scaffold DNA sequence

612. scaff-624: $Z^*$ $Y^*$ $X^*$ $W^*$ $V^*$ $U^*$ $T^*$ $S^*$ $R^*$ $Q^*$ $P^*$ $O^*$ $N^*$ $M^*$ $L^*$ $K^*$ $J^*$ $I^*$ $H^*$ $G^*$ $F^*$ $E^*$ $D^*$ $C^*$ $B^*$ $A^*$:
CAGTATTGGACGCTATCCAGTCTA AACATTTTACTATTACCCCCTCTG GCAAAACTTCTTTTGCAAAAGCCT
CTCGCTATTTTGGTTTTTATCGTC GTCTGGTAAACGAGGGTTATGATA GTGTTGCTCTTACTATGCCTCGTA
AATTCCTTTTGGCGAATGTATCTG CATTAGTTGAATGTGGTATTCCTA AATCTCAACTGATGAATCTTTCTA
CCTGTAATAATGTTGTTCCGTTAG TTCGTTTTATTAACGTAGATTTTT CTTCCCAACGTCCTGACTGGTATA
CATGAGCCAGTTCTTAAAATCGCA TAAGGTAATTCACAATGATTAAAG TTGAAATTAAACCATCTCAAGCCC
ATTACTACTCGTTCTGGTGTTTCT CGTCAGGGCAAGCCTTATTCACTG AATGAGCAGCTTTGTTACGTTGAT
TTTGGGTAATGAATATCCGGTTCT TGTCAAGATTTACTCTTGATGAAG GTCAGCCAGCCTATGCGCCTGGTC
TGTACACCGTTCATCTGTCCTCTT TCAAAGTTGGTCAGTTCGGTTCCC TTATGATTGACCGTCTGCGCCTCG
TCCGGCTAAGTAACATGGAGCAGG TCGCGGATTTCGACACAATTTATC

##### S10.3.2 25 position IsoenergeticBitCopy DNA sequences

613. $A0$: GATAAATTGTGTCGAAATCCGCGA CTCATCCTGACC
614. $B0$: CCTGCTCCATGTTACTTAGCCGGA CTCATCCTGACC
615. $C0$: ACGAGGCGCAGACGGTCAATCATA CTCATCCTGACC
616. $D0$: AGGGAACCGAACTGACCAACTTTG CTCATCCTGACC

617. $E0$: AAAGAGGACAGATGAACGGTGTAC CTCATCCTGACC
618. $F0$: AGACCAGGCGCATAGGCTGGCTGA CTCATCCTGACC
619. $G0$: CCTTCATCAAGAGTAATCTTGACA CTCATCCTGACC
620. $H0$: AGAACCGGATATTCATTACCCAAA CTCATCCTGACC
621. $I0$: TCAACGTAACAAAGCTGCTCATTC CTCATCCTGACC
622. $J0$: AGTGAATAAGGCTTGCCCTGACGA CTCATCCTGACC
623. $A\overline{1}$: GATAAATTGTGTCGAAATCCGCGA CATCTCCGATCC
624. $B\overline{1}$: CCTGCTCCATGTTACTTAGCCGGA CATCTCCGATCC
625. $C\overline{1}$: ACGAGGCGCAGACGGTCAATCATA CATCTCCGATCC
626. $D\overline{1}$: AGGGAACCGAACTGACCAACTTTG CATCTCCGATCC
627. $E\overline{1}$: AAAGAGGACAGATGAACGGTGTAC CATCTCCGATCC
628. $F\overline{1}$: AGACCAGGCGCATAGGCTGGCTGA CATCTCCGATCC
629. $G\overline{1}$: CCTTCATCAAGAGTAATCTTGACA CATCTCCGATCC
630. $H\overline{1}$: AGAACCGGATATTCATTACCCAAA CATCTCCGATCC
631. $I\overline{1}$: TCAACGTAACAAAGCTGCTCATTC CATCTCCGATCC
632. $J\overline{1}$: AGTGAATAAGGCTTGCCCTGACGA CATCTCCGATCC
633. $AQ^*$: GATAAATTGTGTCGAAATCCGCGA GGGTAGTGTGGAGAGGTGGG
634. $BQ^*$: CCTGCTCCATGTTACTTAGCCGGA GGGTAGTGTGGAGAGGTGGG
635. $CQ^*$: ACGAGGCGCAGACGGTCAATCATA GGGTAGTGTGGAGAGGTGGG
636. $DQ^*$: AGGGAACCGAACTGACCAACTTTG GGGTAGTGTGGAGAGGTGGG
637. $EQ^*$: AAAGAGGACAGATGAACGGTGTAC GGGTAGTGTGGAGAGGTGGG
638. $FQ^*$: AGACCAGGCGCATAGGCTGGCTGA GGGTAGTGTGGAGAGGTGGG
639. $GQ^*$: CCTTCATCAAGAGTAATCTTGACA GGGTAGTGTGGAGAGGTGGG
640. $HQ^*$: AGAACCGGATATTCATTACCCAAA GGGTAGTGTGGAGAGGTGGG
641. $IQ^*$: TCAACGTAACAAAGCTGCTCATTC GGGTAGTGTGGAGAGGTGGG
642. $JQ^*$: AGTGAATAAGGCTTGCCCTGACGA GGGTAGTGTGGAGAGGTGGG
643. $0^*B\overline{1}$: GGTCAGGATGAG CCTGCTCCATGTTACTTAGCCGGA CATCTCCGATCC
644. $0^*C\overline{1}$: GGTCAGGATGAG ACGAGGCGCAGACGGTCAATCATA CATCTCCGATCC
645. $0^*D\overline{1}$: GGTCAGGATGAG AGGGAACCGAACTGACCAACTTTG CATCTCCGATCC
646. $0^*E\overline{1}$: GGTCAGGATGAG AAAGAGGACAGATGAACGGTGTAC CATCTCCGATCC
647. $0^*F\overline{1}$: GGTCAGGATGAG AGACCAGGCGCATAGGCTGGCTGA CATCTCCGATCC
648. $0^*G\overline{1}$: GGTCAGGATGAG CCTTCATCAAGAGTAATCTTGACA CATCTCCGATCC
649. $0^*H\overline{1}$: GGTCAGGATGAG AGAACCGGATATTCATTACCCAAA CATCTCCGATCC
650. $0^*I\overline{1}$: GGTCAGGATGAG TCAACGTAACAAAGCTGCTCATTC CATCTCCGATCC
651. $0^*J\overline{1}$: GGTCAGGATGAG AGTGAATAAGGCTTGCCCTGACGA CATCTCCGATCC
652. $0^*K\overline{1}$: GGTCAGGATGAG GAAACACCAGAACGAGTAGTAAAT CATCTCCGATCC
653. $0^*L\overline{1}$: GGTCAGGATGAG TGGGCTTGAGATGGTTTAATTTCA CATCTCCGATCC
654. $0^*M\overline{1}$: GGTCAGGATGAG ACTTTAATCATTGTGAATTACCTT CATCTCCGATCC
655. $0^*N\overline{1}$: GGTCAGGATGAG ATGCGATTTTAAGAACTGGCTCAT CATCTCCGATCC
656. $0^*O\overline{1}$: GGTCAGGATGAG TATACCAGTCAGGACGTTGGGAAG CATCTCCGATCC
657. $0^*P\overline{1}$: GGTCAGGATGAG AAAAATCTACGTTAATAAAACGAA CATCTCCGATCC
658. $0^*Q\overline{1}$: GGTCAGGATGAG CTAACGGAACAACATTATTACAGG CATCTCCGATCC
659. $0^*R\overline{1}$: GGTCAGGATGAG TAGAAAGATTCATCAGTTGAGATT CATCTCCGATCC
660. $0^*S\overline{1}$: GGTCAGGATGAG TAGGAATACCACATTCAACTAATG CATCTCCGATCC
661. $0^*T\overline{1}$: GGTCAGGATGAG CAGATACATAACGCCAAAAGGAAT CATCTCCGATCC
662. $0^*U\overline{1}$: GGTCAGGATGAG TACGAGGCATAGTAAGAGCAACAC CATCTCCGATCC
663. $0^*V\overline{1}$: GGTCAGGATGAG TATCATAACCCTCGTTTACCAGAC CATCTCCGATCC
664. $0^*W\overline{1}$: GGTCAGGATGAG GACGATAAAAACCAAAATAGCGAG CATCTCCGATCC
665. $0^*X\overline{1}$: GGTCAGGATGAG AGGCTTTTGCAAAAGAAGTTTTGC CATCTCCGATCC
666. $0^*Y\overline{1}$: GGTCAGGATGAG CAGAGGGGGTAATAGTAAAATGTT CATCTCCGATCC
667. $\overline{1}^*B0$: GGATCGGAGATG CCTGCTCCATGTTACTTAGCCGGA CTCATCCTGACC
668. $\overline{1}^*C0$: GGATCGGAGATG ACGAGGCGCAGACGGTCAATCATA CTCATCCTGACC
669. $\overline{1}^*D0$: GGATCGGAGATG AGGGAACCGAACTGACCAACTTTG CTCATCCTGACC
670. $\overline{1}^*E0$: GGATCGGAGATG AAAGAGGACAGATGAACGGTGTAC CTCATCCTGACC

671. $\overline{1}^*F0$: GGATCGGAGATG AGACCAGGCGCATAGGCTGGCTGA CTCATCCTGACC
672. $\overline{1}^*G0$: GGATCGGAGATG CCTTCATCAAGAGTAATCTTGACA CTCATCCTGACC
673. $\overline{1}^*H0$: GGATCGGAGATG AGAACCGGATATTCATTACCCAAA CTCATCCTGACC
674. $\overline{1}^*I0$: GGATCGGAGATG TCAACGTAACAAAGCTGCTCATTC CTCATCCTGACC
675. $\overline{1}^*J0$: GGATCGGAGATG AGTGAATAAGGCTTGCCCTGACGA CTCATCCTGACC
676. $\overline{1}^*K0$: GGATCGGAGATG GAAACACCAGAACGAGTAGTAAAT CTCATCCTGACC
677. $\overline{1}^*L0$: GGATCGGAGATG TGGGCTTGAGATGGTTTAATTTCA CTCATCCTGACC
678. $\overline{1}^*M0$: GGATCGGAGATG ACTTTAATCATTGTGAATTACCTT CTCATCCTGACC
679. $\overline{1}^*N0$: GGATCGGAGATG ATGCGATTTTAAGAACTGGCTCAT CTCATCCTGACC
680. $\overline{1}^*O0$: GGATCGGAGATG TATACCAGTCAGGACGTTGGGAAG CTCATCCTGACC
681. $\overline{1}^*P0$: GGATCGGAGATG AAAAATCTACGTTAATAAAACGAA CTCATCCTGACC
682. $\overline{1}^*Q0$: GGATCGGAGATG CTAACGGAACAACATTATTACAGG CTCATCCTGACC
683. $\overline{1}^*R0$: GGATCGGAGATG TAGAAAGATTCATCAGTTGAGATT CTCATCCTGACC
684. $\overline{1}^*S0$: GGATCGGAGATG TAGGAATACCACATTCAACTAATG CTCATCCTGACC
685. $\overline{1}^*T0$: GGATCGGAGATG CAGATACATAACGCCAAAAGGAAT CTCATCCTGACC
686. $\overline{1}^*U0$: GGATCGGAGATG TACGAGGCATAGTAAGAGCAACAC CTCATCCTGACC
687. $\overline{1}^*V0$: GGATCGGAGATG TATCATAACCCTCGTTTACCAGAC CTCATCCTGACC
688. $\overline{1}^*W0$: GGATCGGAGATG GACGATAAAAACCAAAATAGCGAG CTCATCCTGACC
689. $\overline{1}^*X0$: GGATCGGAGATG AGGCTTTTGCAAAAGAAGTTTTGC CTCATCCTGACC
690. $\overline{1}^*Y0$: GGATCGGAGATG CAGAGGGGGTAATAGTAAAATGTT CTCATCCTGACC
691. $0^*B$: GGTCAGGATGAG CCTGCTCCATGTTACTTAGCCGGA
692. $0^*C$: GGTCAGGATGAG ACGAGGCGCAGACGGTCAATCATA
693. $0^*D$: GGTCAGGATGAG AGGGAACCGAACTGACCAACTTTG
694. $0^*E$: GGTCAGGATGAG AAAGAGGACAGATGAACGGTGTAC
695. $0^*F$: GGTCAGGATGAG AGACCAGGCGCATAGGCTGGCTGA
696. $0^*G$: GGTCAGGATGAG CCTTCATCAAGAGTAATCTTGACA
697. $0^*H$: GGTCAGGATGAG AGAACCGGATATTCATTACCCAAA
698. $0^*I$: GGTCAGGATGAG TCAACGTAACAAAGCTGCTCATTC
699. $0^*J$: GGTCAGGATGAG AGTGAATAAGGCTTGCCCTGACGA
700. $0^*K$: GGTCAGGATGAG GAAACACCAGAACGAGTAGTAAAT
701. $0^*L$: GGTCAGGATGAG TGGGCTTGAGATGGTTTAATTTCA
702. $0^*M$: GGTCAGGATGAG ACTTTAATCATTGTGAATTACCTT
703. $0^*N$: GGTCAGGATGAG ATGCGATTTTAAGAACTGGCTCAT
704. $0^*O$: GGTCAGGATGAG TATACCAGTCAGGACGTTGGGAAG
705. $0^*P$: GGTCAGGATGAG AAAAATCTACGTTAATAAAACGAA
706. $0^*Q$: GGTCAGGATGAG CTAACGGAACAACATTATTACAGG
707. $0^*R$: GGTCAGGATGAG TAGAAAGATTCATCAGTTGAGATT
708. $0^*S$: GGTCAGGATGAG TAGGAATACCACATTCAACTAATG
709. $0^*T$: GGTCAGGATGAG CAGATACATAACGCCAAAAGGAAT
710. $0^*U$: GGTCAGGATGAG TACGAGGCATAGTAAGAGCAACAC
711. $0^*V$: GGTCAGGATGAG TATCATAACCCTCGTTTACCAGAC
712. $0^*W$: GGTCAGGATGAG GACGATAAAAACCAAAATAGCGAG
713. $0^*X$: GGTCAGGATGAG AGGCTTTTGCAAAAGAAGTTTTGC
714. $0^*Y$: GGTCAGGATGAG CAGAGGGGGTAATAGTAAAATGTT
715. $\overline{1}^*B$: GGATCGGAGATG CCTGCTCCATGTTACTTAGCCGGA
716. $\overline{1}^*C$: GGATCGGAGATG ACGAGGCGCAGACGGTCAATCATA
717. $\overline{1}^*D$: GGATCGGAGATG AGGGAACCGAACTGACCAACTTTG
718. $\overline{1}^*E$: GGATCGGAGATG AAAGAGGACAGATGAACGGTGTAC
719. $\overline{1}^*F$: GGATCGGAGATG AGACCAGGCGCATAGGCTGGCTGA
720. $\overline{1}^*G$: GGATCGGAGATG CCTTCATCAAGAGTAATCTTGACA
721. $\overline{1}^*H$: GGATCGGAGATG AGAACCGGATATTCATTACCCAAA
722. $\overline{1}^*I$: GGATCGGAGATG TCAACGTAACAAAGCTGCTCATTC
723. $\overline{1}^*J$: GGATCGGAGATG AGTGAATAAGGCTTGCCCTGACGA
724. $\overline{1}^*K$: GGATCGGAGATG GAAACACCAGAACGAGTAGTAAAT

725. $\overline{1}^*L$: GGATCGGAGATG TGGGCTTGAGATGGTTTAATTTCA
726. $\overline{1}^*M$: GGATCGGAGATG ACTTTAATCATTGTGAATTACCTT
727. $\overline{1}^*N$: GGATCGGAGATG ATGCGATTTTAAGAACTGGCTCAT
728. $\overline{1}^*O$: GGATCGGAGATG TATACCAGTCAGGACGTTGGGAAG
729. $\overline{1}^*P$: GGATCGGAGATG AAAAATCTACGTTAATAAAACGAA
730. $\overline{1}^*Q$: GGATCGGAGATG CTAACGGAACAACATTATTACAGG
731. $\overline{1}^*R$: GGATCGGAGATG TAGAAAGATTCATCAGTTGAGATT
732. $\overline{1}^*S$: GGATCGGAGATG TAGGAATACCACATTCAACTAATG
733. $\overline{1}^*T$: GGATCGGAGATG CAGATACATAACGCCAAAAGGAAT
734. $\overline{1}^*U$: GGATCGGAGATG TACGAGGCATAGTAAGAGCAACAC
735. $\overline{1}^*V$: GGATCGGAGATG TATCATAACCCTCGTTTACCAGAC
736. $\overline{1}^*W$: GGATCGGAGATG GACGATAAAAACCAAAATAGCGAG
737. $\overline{1}^*X$: GGATCGGAGATG AGGCTTTTGCAAAAGAAGTTTTGC
738. $\overline{1}^*Y$: GGATCGGAGATG CAGAGGGGGTAATAGTAAAATGTT
739. $0^*BQ^*$: GGTCAGGATGAG CCTGCTCCATGTTACTTAGCCGGA GGGTAGTGTGGAGAGGTGGG
740. $0^*CQ^*$: GGTCAGGATGAG ACGAGGCGCAGACGGTCAATCATA GGGTAGTGTGGAGAGGTGGG
741. $0^*DQ^*$: GGTCAGGATGAG AGGGAACCGAACTGACCAACTTTG GGGTAGTGTGGAGAGGTGGG
742. $0^*EQ^*$: GGTCAGGATGAG AAAGAGGACAGATGAACGGTGTAC GGGTAGTGTGGAGAGGTGGG
743. $0^*FQ^*$: GGTCAGGATGAG AGACCAGGCGCATAGGCTGGCTGA GGGTAGTGTGGAGAGGTGGG
744. $0^*GQ^*$: GGTCAGGATGAG CCTTCATCAAGAGTAATCTTGACA GGGTAGTGTGGAGAGGTGGG
745. $0^*HQ^*$: GGTCAGGATGAG AGAACCGGATATTCATTACCCAAA GGGTAGTGTGGAGAGGTGGG
746. $0^*IQ^*$: GGTCAGGATGAG TCAACGTAACAAAGCTGCTCATTC GGGTAGTGTGGAGAGGTGGG
747. $0^*JQ^*$: GGTCAGGATGAG AGTGAATAAGGCTTGCCCTGACGA GGGTAGTGTGGAGAGGTGGG
748. $0^*KQ^*$: GGTCAGGATGAG GAAACACCAGAACGAGTAGTAAAT GGGTAGTGTGGAGAGGTGGG
749. $0^*LQ^*$: GGTCAGGATGAG TGGGCTTGAGATGGTTTAATTTCA GGGTAGTGTGGAGAGGTGGG
750. $0^*MQ^*$: GGTCAGGATGAG ACTTTAATCATTGTGAATTACCTT GGGTAGTGTGGAGAGGTGGG
751. $0^*NQ^*$: GGTCAGGATGAG ATGCGATTTTAAGAACTGGCTCATG GGTAGTGTGGAGAGGTGGG
752. $0^*OQ^*$: GGTCAGGATGAG TATACCAGTCAGGACGTTGGGAAGG GGTAGTGTGGAGAGGTGGG
753. $0^*PQ^*$: GGTCAGGATGAG AAAAATCTACGTTAATAAAACGAAG GGTAGTGTGGAGAGGTGGG
754. $0^*QQ^*$: GGTCAGGATGAG CTAACGGAACAACATTATTACAGGG GGTAGTGTGGAGAGGTGGG
755. $0^*RQ^*$: GGTCAGGATGAG TAGAAAGATTCATCAGTTGAGATTG GGTAGTGTGGAGAGGTGGG
756. $0^*SQ^*$: GGTCAGGATGAG TAGGAATACCACATTCAACTAATGG GGTAGTGTGGAGAGGTGGG
757. $0^*TQ^*$: GGTCAGGATGAG CAGATACATAACGCCAAAAGGAATG GGTAGTGTGGAGAGGTGGG
758. $0^*UQ^*$: GGTCAGGATGAG TACGAGGCATAGTAAGAGCAACACG GGTAGTGTGGAGAGGTGGG
759. $0^*VQ^*$: GGTCAGGATGAG TATCATAACCCTCGTTTACCAGACG GGTAGTGTGGAGAGGTGGG
760. $0^*WQ^*$: GGTCAGGATGAG GACGATAAAAACCAAAATAGCGAGG GGTAGTGTGGAGAGGTGGG
761. $0^*XQ^*$: GGTCAGGATGAG AGGCTTTTGCAAAAGAAGTTTTGCG GGTAGTGTGGAGAGGTGGG
762. $0^*YQ^*$: GGTCAGGATGAG CAGAGGGGGTAATAGTAAAATGTTG GGTAGTGTGGAGAGGTGGG
763. $\overline{1}^*BQ^*$: GGATCGGAGATG CCTGCTCCATGTTACTTAGCCGGA GGGTAGTGTGGAGAGGTGGG
764. $\overline{1}^*CQ^*$: GGATCGGAGATG ACGAGGCGCAGACGGTCAATCATA GGGTAGTGTGGAGAGGTGGG
765. $\overline{1}^*DQ^*$: GGATCGGAGATG AGGGAACCGAACTGACCAACTTTG GGGTAGTGTGGAGAGGTGGG
766. $\overline{1}^*EQ^*$: GGATCGGAGATG AAAGAGGACAGATGAACGGTGTAC GGGTAGTGTGGAGAGGTGGG
767. $\overline{1}^*FQ^*$: GGATCGGAGATG AGACCAGGCGCATAGGCTGGCTGA GGGTAGTGTGGAGAGGTGGG
768. $\overline{1}^*GQ^*$: GGATCGGAGATG CCTTCATCAAGAGTAATCTTGACA GGGTAGTGTGGAGAGGTGGG
769. $\overline{1}^*HQ^*$: GGATCGGAGATG AGAACCGGATATTCATTACCCAAA GGGTAGTGTGGAGAGGTGGG
770. $\overline{1}^*IQ^*$: GGATCGGAGATG TCAACGTAACAAAGCTGCTCATTC GGGTAGTGTGGAGAGGTGGG
771. $\overline{1}^*JQ^*$: GGATCGGAGATG AGTGAATAAGGCTTGCCCTGACGA GGGTAGTGTGGAGAGGTGGG
772. $\overline{1}^*KQ^*$: GGATCGGAGATG GAAACACCAGAACGAGTAGTAAAT GGGTAGTGTGGAGAGGTGGG
773. $\overline{1}^*LQ^*$: GGATCGGAGATG TGGGCTTGAGATGGTTTAATTTCA GGGTAGTGTGGAGAGGTGGG
774. $\overline{1}^*MQ^*$: GGATCGGAGATG ACTTTAATCATTGTGAATTACCTT GGGTAGTGTGGAGAGGTGGG
775. $\overline{1}^*NQ^*$: GGATCGGAGATG ATGCGATTTTAAGAACTGGCTCAT GGGTAGTGTGGAGAGGTGGG
776. $\overline{1}^*OQ^*$: GGATCGGAGATG TATACCAGTCAGGACGTTGGGAAG GGGTAGTGTGGAGAGGTGGG
777. $\overline{1}^*PQ^*$: GGATCGGAGATG AAAAATCTACGTTAATAAAACGAA GGGTAGTGTGGAGAGGTGGG
778. $\overline{1}^*QQ^*$: GGATCGGAGATG CTAACGGAACAACATTATTACAGG GGGTAGTGTGGAGAGGTGGG

779. $\overline{1}^*RQ^*$: GGATCGGAGATG TAGAAAGATTCATCAGTTGAGATT GGGTAGTGTGGAGAGGTGGG
780. $\overline{1}^*SQ^*$: GGATCGGAGATG TAGGAATACCACATTCAACTAATG GGGTAGTGTGGAGAGGTGGG
781. $\overline{1}^*TQ^*$: GGATCGGAGATG CAGATACATAACGCCAAAAGGAAT GGGTAGTGTGGAGAGGTGGG
782. $\overline{1}^*UQ^*$: GGATCGGAGATG TACGAGGCATAGTAAGAGCAACAC GGGTAGTGTGGAGAGGTGGG
783. $\overline{1}^*VQ^*$: GGATCGGAGATG TATCATAACCCTCGTTTACCAGAC GGGTAGTGTGGAGAGGTGGG
784. $\overline{1}^*WQ^*$: GGATCGGAGATG GACGATAAAAACCAAAATAGCGAG GGGTAGTGTGGAGAGGTGGG
785. $\overline{1}^*XQ^*$: GGATCGGAGATG AGGCTTTTGCAAAAGAAGTTTTGC GGGTAGTGTGGAGAGGTGGG
786. $\overline{1}^*YQ^*$: GGATCGGAGATG CAGAGGGGGTAATAGTAAAATGTT GGGTAGTGTGGAGAGGTGGG
787. RF$^*B$: TTGGGATGCGAAGGGATGGT CCTGCTCCATGTTACTTAGCCGGA
788. RF$^*C$: TTGGGATGCGAAGGGATGGT ACGAGGCGCAGACGGTCAATCATA
789. RF$^*D$: TTGGGATGCGAAGGGATGGT AGGGAACCGAACTGACCAACTTTG
790. RF$^*E$: TTGGGATGCGAAGGGATGGT AAAGAGGACAGATGAACGGTGTAC
791. RF$^*F$: TTGGGATGCGAAGGGATGGT AGACCAGGCGCATAGGCTGGCTGA
792. RF$^*G$: TTGGGATGCGAAGGGATGGT CCTTCATCAAGAGTAATCTTGACA
793. RF$^*H$: TTGGGATGCGAAGGGATGGT AGAACCGGATATTCATTACCCAAA
794. RF$^*I$: TTGGGATGCGAAGGGATGGT TCAACGTAACAAAGCTGCTCATTC
795. RF$^*J$: TTGGGATGCGAAGGGATGGT AGTGAATAAGGCTTGCCCTGACGA
796. RF$^*K$: TTGGGATGCGAAGGGATGGT GAAACACCAGAACGAGTAGTAAAT
797. RF$^*L$: TTGGGATGCGAAGGGATGGT TGGGCTTGAGATGGTTTAATTTCA
798. RF$^*M$: TTGGGATGCGAAGGGATGGT ACTTTAATCATTGTGAATTACCTT
799. RF$^*N$: TTGGGATGCGAAGGGATGGT ATGCGATTTTAAGAACTGGCTCAT
800. RF$^*O$: TTGGGATGCGAAGGGATGGT TATACCAGTCAGGACGTTGGGAAG
801. RF$^*P$: TTGGGATGCGAAGGGATGGT AAAAATCTACGTTAATAAAACGAA
802. RF$^*Q$: TTGGGATGCGAAGGGATGGT CTAACGGAACAACATTATTACAGG
803. RF$^*R$: TTGGGATGCGAAGGGATGGT TAGAAAGATTCATCAGTTGAGATT
804. RF$^*S$: TTGGGATGCGAAGGGATGGT TAGGAATACCACATTCAACTAATG
805. RF$^*T$: TTGGGATGCGAAGGGATGGT CAGATACATAACGCCAAAAGGAAT
806. RF$^*U$: TTGGGATGCGAAGGGATGGT TACGAGGCATAGTAAGAGCAACAC
807. RF$^*V$: TTGGGATGCGAAGGGATGGT TATCATAACCCTCGTTTACCAGAC
808. RF$^*W$: TTGGGATGCGAAGGGATGGT GACGATAAAAACCAAAATAGCGAG
809. RF$^*X$: TTGGGATGCGAAGGGATGGT AGGCTTTTGCAAAAGAAGTTTTGC
810. RF$^*Y$: TTGGGATGCGAAGGGATGGT CAGAGGGGGTAATAGTAAAATGTT
811. RF$^*Z$: TTGGGATGCGAAGGGATGGT TAGACTGGATAGCGTCCAATACTG
812. $B$: CCTGCTCCATGTTACTTAGCCGGA
813. $C$: ACGAGGCGCAGACGGTCAATCATA
814. $D$: AGGGAACCGAACTGACCAACTTTG
815. $E$: AAAGAGGACAGATGAACGGTGTAC
816. $F$: AGACCAGGCGCATAGGCTGGCTGA
817. $G$: CCTTCATCAAGAGTAATCTTGACA
818. $H$: AGAACCGGATATTCATTACCCAAA
819. $I$: TCAACGTAACAAAGCTGCTCATTC
820. $J$: AGTGAATAAGGCTTGCCCTGACGA
821. $K$: GAAACACCAGAACGAGTAGTAAAT
822. $L$: TGGGCTTGAGATGGTTTAATTTCA
823. $M$: ACTTTAATCATTGTGAATTACCTT
824. $N$: ATGCGATTTTAAGAACTGGCTCAT
825. $O$: TATACCAGTCAGGACGTTGGGAAG
826. $P$: AAAAATCTACGTTAATAAAACGAA
827. $Q$: CTAACGGAACAACATTATTACAGG
828. $R$: TAGAAAGATTCATCAGTTGAGATT
829. $S$: TAGGAATACCACATTCAACTAATG
830. $T$: CAGATACATAACGCCAAAAGGAAT
831. $U$: TACGAGGCATAGTAAGAGCAACAC
832. $V$: TATCATAACCCTCGTTTACCAGAC

833. $W$: GACGATAAAAACCAAAATAGCGAG
834. $X$: AGGCTTTTGCAAAAGAAGTTTTGC
835. $Y$: CAGAGGGGGTAATAGTAAAATGTT
836. $A$: GATAAATTGTGTCGAAATCCGCGA
837. $Z$: TAGACTGGATAGCGTCCAATACTG
838. $0^*C_{\mathtt{w8G}}\overline{1}$: GGTCAGGATGAG ACGAGGCGGAGACGGTCAATCATA CATCTCCGATCC
839. $0^*F_{\mathtt{w8T}}\overline{1}$: GGTCAGGATGAG AGACCAGGTGCATAGGCTGGCTGA CATCTCCGATCC
840. $0^*H_{\mathtt{w9C}}\overline{1}$: GGTCAGGATGAG AGAACCGGACATTCATTACCCAAA CATCTCCGATCC
841. $0^*I_{\mathtt{w13C}}\overline{1}$: GGTCAGGATGAG TCAACGTAACAAACCTGCTCATTC CATCTCCGATCC
842. $0^*J_{\mathtt{w15T}}\overline{1}$: GGTCAGGATGAG AGTGAATAAGGCTTGTCCTGACGA CATCTCCGATCC
843. $0^*K_{\mathtt{w16A}}\overline{1}$: GGTCAGGATGAG GAAACACCAGAACGAGAAGTAAAT CATCTCCGATCC
844. $0^*L_{\mathtt{w16C}}\overline{1}$: GGTCAGGATGAG TGGGCTTGAGATGGTTCAATTTCA CATCTCCGATCC
845. $\overline{1}^*C_{\mathtt{w8G}}$: GGATCGGAGATG ACGAGGCGGAGACGGTCAATCATA CTCATCCTGACC
846. $\overline{1}^*F_{\mathtt{w8T}}$: GGATCGGAGATG AGACCAGGTGCATAGGCTGGCTGA CTCATCCTGACC
847. $\overline{1}^*H_{\mathtt{w9C}}$: GGATCGGAGATG AGAACCGGACATTCATTACCCAAA CTCATCCTGACC
848. $\overline{1}^*I_{\mathtt{w13C}}$: GGATCGGAGATG TCAACGTAACAAACCTGCTCATTC CTCATCCTGACC
849. $\overline{1}^*J_{\mathtt{w15T}}$: GGATCGGAGATG AGTGAATAAGGCTTGTCCTGACGA CTCATCCTGACC
850. $\overline{1}^*K_{\mathtt{w16A}}$: GGATCGGAGATG GAAACACCAGAACGAGAAGTAAAT CTCATCCTGACC
851. $\overline{1}^*L_{\mathtt{w16C}}$: GGATCGGAGATG TGGGCTTGAGATGGTTCAATTTCA CTCATCCTGACC
852. $0^*N_{\mathtt{w10C}}\overline{1}$: GGTCAGGATGAG ATGCGATTTTCAGAACTGGCTCAT CATCTCCGATCC
853. $0^*O_{\mathtt{w15C}}\overline{1}$: GGTCAGGATGAG TATACCAGTCAGGACCTTGGGAAG CATCTCCGATCC
854. $0^*Q_{\mathtt{w14C}}\overline{1}$: GGTCAGGATGAG CTAACGGAACAACACTATTACAGG CATCTCCGATCC
855. $0^*R_{\mathtt{w16C}}\overline{1}$: GGTCAGGATGAG TAGAAAGATTCATCAGCTGAGATT CATCTCCGATCC
856. $0^*S_{\mathtt{w11A}}\overline{1}$: GGTCAGGATGAG TAGGAATACCAAATTCAACTAATG CATCTCCGATCC
857. $0^*T_{\mathtt{w7C}}\overline{1}$: GGTCAGGATGAG CAGATACCTAACGCCAAAAGGAAT CATCTCCGATCC
858. $0^*U_{\mathtt{w7T}}\overline{1}$: GGTCAGGATGAG TACGAGGTATAGTAAGAGCAACAC CATCTCCGATCC
859. $0^*V_{\mathtt{w13T}}\overline{1}$: GGTCAGGATGAG TATCATAACCCTCTTTTACCAGAC CATCTCCGATCC
860. $0^*W_{\mathtt{w13C}}\overline{1}$: GGTCAGGATGAG GACGATAAAAACCCAAATAGCGAG CATCTCCGATCC
861. $0^*X_{\mathtt{w10G}}\overline{1}$: GGTCAGGATGAG AGGCTTTTGCGAAAGAAGTTTTGC CATCTCCGATCC
862. $Y_{\mathtt{w7Cw8C}}$: GGTCAGGATGAG CAGAGGGGTTAATAGTAAAATGTT CATCTCCGATCC
863. $\overline{1}^*N_{\mathtt{w10C}}0$: GGATCGGAGATG ATGCGATTTTCAGAACTGGCTCAT CTCATCCTGACC
864. $\overline{1}^*O_{\mathtt{w15C}}0$: GGATCGGAGATG TATACCAGTCAGGACCTTGGGAAG CTCATCCTGACC
865. $\overline{1}^*Q_{\mathtt{w14C}}0$: GGATCGGAGATG CTAACGGAACAACACTATTACAGG CTCATCCTGACC
866. $\overline{1}^*R_{\mathtt{w16C}}0$: GGATCGGAGATG TAGAAAGATTCATCAGCTGAGATT CTCATCCTGACC
867. $\overline{1}^*S_{\mathtt{w11A}}0$: GGATCGGAGATG TAGGAATACCAAATTCAACTAATG CTCATCCTGACC
868. $\overline{1}^*T_{\mathtt{w7C}}0$: GGATCGGAGATG CAGATACCTAACGCCAAAAGGAAT CTCATCCTGACC
869. $\overline{1}^*U_{\mathtt{w7T}}0$: GGATCGGAGATG TACGAGGTATAGTAAGAGCAACAC CTCATCCTGACC
870. $\overline{1}^*V_{\mathtt{w13T}}0$: GGATCGGAGATG TATCATAACCCTCTTTTACCAGAC CTCATCCTGACC
871. $\overline{1}^*W_{\mathtt{w13C}}0$: GGATCGGAGATG GACGATAAAAACCCAAATAGCGAG CTCATCCTGACC
872. $\overline{1}^*X_{\mathtt{w10G}}0$: GGATCGGAGATG AGGCTTTTGCGAAAGAAGTTTTGC CTCATCCTGACC
873. $\overline{1}^*Y_{\mathtt{w8T}}0$: GGATCGGAGATG CAGAGGGGTTAATAGTAAAATGTT CTCATCCTGACC
874. $0^*C_{\mathtt{w8G}}$: GGTCAGGATGAG ACGAGGCGGAGACGGTCAATCATA
875. $0^*F_{\mathtt{w8T}}$: GGTCAGGATGAG AGACCAGGTGCATAGGCTGGCTGA
876. $0^*H_{\mathtt{w9C}}$: GGTCAGGATGAG AGAACCGGACATTCATTACCCAAA
877. $0^*I_{\mathtt{w13C}}$: GGTCAGGATGAG TCAACGTAACAAACCTGCTCATTC
878. $0^*J_{\mathtt{w15T}}$: GGTCAGGATGAG AGTGAATAAGGCTTGTCCTGACGA
879. $0^*K_{\mathtt{w16A}}$: GGTCAGGATGAG GAAACACCAGAACGAGAAGTAAAT
880. $0^*L_{\mathtt{w16C}}$: GGTCAGGATGAG TGGGCTTGAGATGGTTCAATTTCA
881. $\overline{1}^*C_{\mathtt{w8G}}$: GGATCGGAGATG ACGAGGCGGAGACGGTCAATCATA
882. $\overline{1}^*F_{\mathtt{w8T}}$: GGATCGGAGATG AGACCAGGTGCATAGGCTGGCTGA
883. $\overline{1}^*H_{\mathtt{w9C}}$: GGATCGGAGATG AGAACCGGACATTCATTACCCAAA
884. $\overline{1}^*I_{\mathtt{w13C}}$: GGATCGGAGATG TCAACGTAACAAACCTGCTCATTC
885. $\overline{1}^*J_{\mathtt{w15T}}$: GGATCGGAGATG AGTGAATAAGGCTTGTCCTGACGA
886. $\overline{1}^*K_{\mathtt{w16A}}$: GGATCGGAGATG GAAACACCAGAACGAGAAGTAAAT

887. $\overline{1}^*L_{w16C}$: GGATCGGAGATG TGGGCTTGAGATGGTTCAATTTCA
888. $0^*C_{w8G}Q^*$: GGTCAGGATGAG ACGAGGCGGAGACGGTCAATCATA GGGTAGTGTGGAGAGGTGGG
889. $0^*F_{w8T}Q^*$: GGTCAGGATGAG AGACCAGGTGCATAGGCTGGCTGA GGGTAGTGTGGAGAGGTGGG
890. $0^*H_{w9C}Q^*$: GGTCAGGATGAG AGAACCGGACATTCATTACCCAAA GGGTAGTGTGGAGAGGTGGG
891. $0^*I_{w13C}Q^*$: GGTCAGGATGAG TCAACGTAACAAACCTGCTCATTC GGGTAGTGTGGAGAGGTGGG
892. $0^*J_{w15T}Q^*$: GGTCAGGATGAG AGTGAATAAGGCTTGTCCTGACGA GGGTAGTGTGGAGAGGTGGG
893. $0^*K_{w16A}Q^*$: GGTCAGGATGAG GAAACACCAGAACGAGAAGTAAAT GGGTAGTGTGGAGAGGTGGG
894. $0^*L_{w16C}Q^*$: GGTCAGGATGAG TGGGCTTGAGATGGTTCAATTTCA GGGTAGTGTGGAGAGGTGGG
895. $\overline{1}^*C_{w8G}Q^*$: GGATCGGAGATG ACGAGGCGGAGACGGTCAATCATA GGGTAGTGTGGAGAGGTGGG
896. $\overline{1}^*F_{w8T}Q^*$: GGATCGGAGATG AGACCAGGTGCATAGGCTGGCTGA GGGTAGTGTGGAGAGGTGGG
897. $\overline{1}^*H_{w9C}Q^*$: GGATCGGAGATG AGAACCGGACATTCATTACCCAAA GGGTAGTGTGGAGAGGTGGG
898. $\overline{1}^*I_{w13C}Q^*$: GGATCGGAGATG TCAACGTAACAAACCTGCTCATTC GGGTAGTGTGGAGAGGTGGG
899. $\overline{1}^*J_{w15T}Q^*$: GGATCGGAGATG AGTGAATAAGGCTTGTCCTGACGA GGGTAGTGTGGAGAGGTGGG
900. $\overline{1}^*K_{w16A}Q^*$: GGATCGGAGATG GAAACACCAGAACGAGAAGTAAAT GGGTAGTGTGGAGAGGTGGG
901. $\overline{1}^*L_{w16C}Q^*$: GGATCGGAGATG TGGGCTTGAGATGGTTCAATTTCA GGGTAGTGTGGAGAGGTGGG
902. $0^*N_{w10C}$: GGTCAGGATGAG ATGCGATTTTCAGAACTGGCTCAT
903. $0^*O_{w15C}$: GGTCAGGATGAG TATACCAGTCAGGACCTTGGGAAG
904. $0^*Q_{w14C}$: GGTCAGGATGAG CTAACGGAACAACACTATTACAGG
905. $0^*R_{w16C}$: GGTCAGGATGAG TAGAAAGATTCATCAGCTGAGATT
906. $0^*S_{w11A}$: GGTCAGGATGAG TAGGAATACCAAATTCAACTAATG
907. $0^*T_{w7C}$: GGTCAGGATGAG CAGATACCTAACGCCAAAAGGAAT
908. $0^*U_{w7T}$: GGTCAGGATGAG TACGAGGTATAGTAAGAGCAACAC
909. $0^*V_{w13T}$: GGTCAGGATGAG TATCATAACCCTCTTTTACCAGAC
910. $0^*W_{w13C}$: GGTCAGGATGAG GACGATAAAAACCCAAATAGCGAG
911. $0^*X_{w10G}$: GGTCAGGATGAG AGGCTTTTGCGAAAGAAGTTTTGC
912. $0^*Y_{w8T}$: GGTCAGGATGAG CAGAGGGGTTAATAGTAAAATGTT
913. $\overline{1}^*N_{w10C}$: GGATCGGAGATG ATGCGATTTTCAGAACTGGCTCAT
914. $\overline{1}^*O_{w15C}$: GGATCGGAGATG TATACCAGTCAGGACCTTGGGAAG
915. $\overline{1}^*Q_{w14C}$: GGATCGGAGATG CTAACGGAACAACACTATTACAGG
916. $\overline{1}^*R_{w16C}$: GGATCGGAGATG TAGAAAGATTCATCAGCTGAGATT
917. $\overline{1}^*S_{w11A}$: GGATCGGAGATG TAGGAATACCAAATTCAACTAATG
918. $\overline{1}^*T_{w7C}$: GGATCGGAGATG CAGATACCTAACGCCAAAAGGAAT
919. $\overline{1}^*U_{w7T}$: GGATCGGAGATG TACGAGGTATAGTAAGAGCAACAC
920. $\overline{1}^*V_{w13T}$: GGATCGGAGATG TATCATAACCCTCTTTTACCAGAC
921. $\overline{1}^*W_{w13C}$: GGATCGGAGATG GACGATAAAAACCCAAATAGCGAG
922. $\overline{1}^*X_{w10G}$: GGATCGGAGATG AGGCTTTTGCGAAAGAAGTTTTGC
923. $\overline{1}^*Y_{w8T}$: GGATCGGAGATG CAGAGGGGTTAATAGTAAAATGTT
924. $0^*N_{w10C}Q^*$: GGTCAGGATGAG ATGCGATTTTCAGAACTGGCTCAT GGGTAGTGTGGAGAGGTGGG
925. $0^*O_{w15C}Q^*$: GGTCAGGATGAG TATACCAGTCAGGACCTTGGGAAG GGGTAGTGTGGAGAGGTGGG
926. $0^*Q_{w14C}Q^*$: GGTCAGGATGAG CTAACGGAACAACACTATTACAGG GGGTAGTGTGGAGAGGTGGG
927. $0^*R_{w16C}Q^*$: GGTCAGGATGAG TAGAAAGATTCATCAGCTGAGATT GGGTAGTGTGGAGAGGTGGG
928. $0^*S_{w11A}Q^*$: GGTCAGGATGAG TAGGAATACCAAATTCAACTAATG GGGTAGTGTGGAGAGGTGGG
929. $0^*T_{w7C}Q^*$: GGTCAGGATGAG CAGATACCTAACGCCAAAAGGAAT GGGTAGTGTGGAGAGGTGGG
930. $0^*U_{w7T}Q^*$: GGTCAGGATGAG TACGAGGTATAGTAAGAGCAACAC GGGTAGTGTGGAGAGGTGGG
931. $0^*V_{w13T}Q^*$: GGTCAGGATGAG TATCATAACCCTCTTTTACCAGAC GGGTAGTGTGGAGAGGTGGG
932. $0^*W_{w13C}Q^*$: GGTCAGGATGAG GACGATAAAAACCCAAATAGCGAG GGGTAGTGTGGAGAGGTGGG
933. $0^*X_{w10G}Q^*$: GGTCAGGATGAG AGGCTTTTGCGAAAGAAGTTTTGC GGGTAGTGTGGAGAGGTGGG
934. $0^*Y_{w8T}Q^*$: GGTCAGGATGAG CAGAGGGGTTAATAGTAAAATGTT GGGTAGTGTGGAGAGGTGGG
935. $\overline{1}^*N_{w10C}Q^*$: GGATCGGAGATG ATGCGATTTTCAGAACTGGCTCAT GGGTAGTGTGGAGAGGTGGG
936. $\overline{1}^*O_{w15C}Q^*$: GGATCGGAGATG TATACCAGTCAGGACCTTGGGAAG GGGTAGTGTGGAGAGGTGGG
937. $\overline{1}^*Q_{w14C}Q^*$: GGATCGGAGATG CTAACGGAACAACACTATTACAGG GGGTAGTGTGGAGAGGTGGG
938. $\overline{1}^*R_{w16C}Q^*$: GGATCGGAGATG TAGAAAGATTCATCAGCTGAGATT GGGTAGTGTGGAGAGGTGGG
939. $\overline{1}^*S_{w11A}Q^*$: GGATCGGAGATG TAGGAATACCAAATTCAACTAATG GGGTAGTGTGGAGAGGTGGG
940. $\overline{1}^*T_{w7C}Q^*$: GGATCGGAGATG CAGATACCTAACGCCAAAAGGAAT GGGTAGTGTGGAGAGGTGGG

941. $\overline{1}^*U_{\mathtt{w7T}}\mathtt{Q}^*$: GGATCGGAGATG TACGAGGTATAGTAAGAGCAACAC GGGTAGTGTGGAGAGGTGGG

942. $\overline{1}^*V_{\mathtt{w13T}}\mathtt{Q}^*$: GGATCGGAGATG TATCATAACCCTCTTTTACCAGAC GGGTAGTGTGGAGAGGTGGG

943. $\overline{1}^*W_{\mathtt{w13C}}\mathtt{Q}^*$: GGATCGGAGATG GACGATAAAAACCCAAATAGCGAG GGGTAGTGTGGAGAGGTGGG

944. $\overline{1}^*X_{\mathtt{w10G}}\mathtt{Q}^*$: GGATCGGAGATG AGGCTTTTGCGAAAGAAGTTTTGC GGGTAGTGTGGAGAGGTGGG

945. $\overline{1}^*Y_{\mathtt{w8T}}\mathtt{Q}^*$: GGATCGGAGATG CAGAGGGGTTAATAGTAAAATGTT GGGTAGTGTGGAGAGGTGGG

946. $0^*Q_{\mathtt{w8T}}\overline{1}$: GGTCAGGATGAG CTAACGGATCAACATTATTACAGG CATCTCCGATCC

947. $0^*R_{\mathtt{w14G}}\overline{1}$: GGTCAGGATGAG TAGAAAGATTCATCGGTTGAGATT CATCTCCGATCC

948. $0^*S_{\mathtt{w15T}}\overline{1}$: GGTCAGGATGAG TAGGAATACCACATTTAACTAATG CATCTCCGATCC

949. $0^*T_{\mathtt{w12A}}\overline{1}$: GGTCAGGATGAG CAGATACATAACACCAAAAGGAAT CATCTCCGATCC

950. $0^*U_{\mathtt{w7Tw8G}}\overline{1}$: GGTCAGGATGAG TACGAGGTGTAGTAAGAGCAACAC CATCTCCGATCC

951. $0^*V_{\mathtt{w9Tw10T}}\overline{1}$: GGTCAGGATGAG TATCATAACTTTCGTTTACCAGAC CATCTCCGATCC

952. $0^*W_{\mathtt{w11Aw12T}}\overline{1}$: GGTCAGGATGAG GACGATAAAAAATAAAATAGCGAG CATCTCCGATCC

953. $0^*X_{\mathtt{w8Cw9T}}\overline{1}$: GGTCAGGATGAG AGGCTTTTCTAAAAGAAGTTTTGC CATCTCCGATCC

954. $0^*Y_{\mathtt{w7Cw8C}}\overline{1}$: GGTCAGGATGAG CAGAGGGCCTAATAGTAAAATGTT CATCTCCGATCC

955. $\overline{1}^*Q_{\mathtt{w8T}}0$: GGATCGGAGATG CTAACGGATCAACATTATTACAGG CTCATCCTGACC

956. $\overline{1}^*R_{\mathtt{w14G}}0$: GGATCGGAGATG TAGAAAGATTCATCGGTTGAGATT CTCATCCTGACC

957. $\overline{1}^*S_{\mathtt{w15T}}0$: GGATCGGAGATG TAGGAATACCACATTTAACTAATG CTCATCCTGACC

958. $\overline{1}^*T_{\mathtt{w12A}}0$: GGATCGGAGATG CAGATACATAACACCAAAAGGAAT CTCATCCTGACC

959. $\overline{1}^*U_{\mathtt{w7Tw8G}}0$: GGATCGGAGATG TACGAGGTGTAGTAAGAGCAACAC CTCATCCTGACC

960. $\overline{1}^*V_{\mathtt{w9Tw10T}}0$: GGATCGGAGATG TATCATAACTTTCGTTTACCAGAC CTCATCCTGACC

961. $\overline{1}^*W_{\mathtt{w11Aw12T}}0$: GGATCGGAGATG GACGATAAAAAATAAAATAGCGAG CTCATCCTGACC

962. $\overline{1}^*X_{\mathtt{w8Cw9T}}0$: GGATCGGAGATG AGGCTTTTCTAAAAGAAGTTTTGC CTCATCCTGACC

963. $\overline{1}^*Y_{\mathtt{w7Cw8C}}0$: GGATCGGAGATG CAGAGGGCCTAATAGTAAAATGTT CTCATCCTGACC

964. $0^*Q_{\mathtt{w8T}}$: GGTCAGGATGAG CTAACGGATCAACATTATTACAGG

965. $0^*R_{\mathtt{w14G}}$: GGTCAGGATGAG TAGAAAGATTCATCGGTTGAGATT

966. $0^*S_{\mathtt{w15T}}$: GGTCAGGATGAG TAGGAATACCACATTTAACTAATG

967. $0^*T_{\mathtt{w12A}}$: GGTCAGGATGAG CAGATACATAACACCAAAAGGAAT

968. $0^*U_{\mathtt{w7Tw8G}}$: GGTCAGGATGAG TACGAGGTGTAGTAAGAGCAACAC

969. $0^*V_{\mathtt{w9Tw10T}}$: GGTCAGGATGAG TATCATAACTTTCGTTTACCAGAC

970. $0^*W_{\mathtt{w11Aw12T}}$: GGTCAGGATGAG GACGATAAAAAATAAAATAGCGAG

971. $0^*X_{\mathtt{w8Cw9T}}$: GGTCAGGATGAG AGGCTTTTCTAAAAGAAGTTTTGC

972. $0^*Y_{\mathtt{w7Cw8C}}$: GGTCAGGATGAG CAGAGGGCCTAATAGTAAAATGTT

973. $\overline{1}^*Q_{\mathtt{w8T}}$: GGATCGGAGATG CTAACGGATCAACATTATTACAGG

974. $\overline{1}^*R_{\mathtt{w14G}}$: GGATCGGAGATG TAGAAAGATTCATCGGTTGAGATT

975. $\overline{1}^*S_{\mathtt{w15T}}$: GGATCGGAGATG TAGGAATACCACATTTAACTAATG

976. $\overline{1}^*T_{\mathtt{w12A}}$: GGATCGGAGATG CAGATACATAACACCAAAAGGAAT

977. $\overline{1}^*U_{\mathtt{w7Tw8G}}$: GGATCGGAGATG TACGAGGTGTAGTAAGAGCAACAC

978. $\overline{1}^*V_{\mathtt{w9Tw10T}}$: GGATCGGAGATG TATCATAACTTTCGTTTACCAGAC

979. $\overline{1}^*W_{\mathtt{w11Aw12T}}$: GGATCGGAGATG GACGATAAAAAATAAAATAGCGAG

980. $\overline{1}^*X_{\mathtt{w8Cw9T}}$: GGATCGGAGATG AGGCTTTTCTAAAAGAAGTTTTGC

981. $\overline{1}^*Y_{\mathtt{w7Cw8C}}$: GGATCGGAGATG CAGAGGGCCTAATAGTAAAATGTT

982. $0^*Q_{\mathtt{w8T}}\mathtt{Q}^*$: GGTCAGGATGAG CTAACGGATCAACATTATTACAGG GGGTAGTGTGGAGAGGTGGG

983. $0^*R_{\mathtt{w14G}}\mathtt{Q}^*$: GGTCAGGATGAG TAGAAAGATTCATCGGTTGAGATT GGGTAGTGTGGAGAGGTGGG

984. $0^*S_{\mathtt{w15T}}\mathtt{Q}^*$: GGTCAGGATGAG TAGGAATACCACATTTAACTAATG GGGTAGTGTGGAGAGGTGGG

985. $0^*T_{\mathtt{w12A}}\mathtt{Q}^*$: GGTCAGGATGAG CAGATACATAACACCAAAAGGAAT GGGTAGTGTGGAGAGGTGGG

986. $0^*U_{\mathtt{w7Tw8G}}\mathtt{Q}^*$: GGTCAGGATGAG TACGAGGTGTAGTAAGAGCAACAC GGGTAGTGTGGAGAGGTGGG

987. $0^*V_{\mathtt{w9Tw10T}}\mathtt{Q}^*$: GGTCAGGATGAG TATCATAACTTTCGTTTACCAGAC GGGTAGTGTGGAGAGGTGGG

988. $0^*W_{\mathtt{w11Aw12T}}\mathtt{Q}^*$: GGTCAGGATGAG GACGATAAAAAATAAAATAGCGAG GGGTAGTGTGGAGAGGTGGG

989. $0^*X_{\mathtt{w8Cw9T}}\mathtt{Q}^*$: GGTCAGGATGAG AGGCTTTTCTAAAAGAAGTTTTGC GGGTAGTGTGGAGAGGTGGG

990. $0^*Y_{\mathtt{w7Cw8C}}\mathtt{Q}^*$: GGTCAGGATGAG CAGAGGGCCTAATAGTAAAATGTT GGGTAGTGTGGAGAGGTGGG

991. $\overline{1}^*Q_{\mathtt{w8T}}\mathtt{Q}^*$: GGATCGGAGATG CTAACGGATCAACATTATTACAGG GGGTAGTGTGGAGAGGTGGG

992. $\overline{1}^*R_{\mathtt{w14G}}\mathtt{Q}^*$: GGATCGGAGATG TAGAAAGATTCATCGGTTGAGATT GGGTAGTGTGGAGAGGTGGG

993. $\overline{1}^*S_{\mathtt{w15T}}\mathtt{Q}^*$: GGATCGGAGATG TAGGAATACCACATTTAACTAATG GGGTAGTGTGGAGAGGTGGG

994. $\overline{1}^*T_{\mathtt{w12A}}\mathtt{Q}^*$: GGATCGGAGATG CAGATACATAACACCAAAAGGAAT GGGTAGTGTGGAGAGGTGGG

995. $\overline{1}^* U_{\texttt{w7Tw8G}}\texttt{Q}^*$: GGATCGGAGATG TACGAGGTGTAGTAAGAGCAACAC GGGTAGTGTGGAGAGGTGGG
996. $\overline{1}^* V_{\texttt{w9Tw10T}}\texttt{Q}^*$: GGATCGGAGATG TATCATAACTTTCGTTTACCAGAC GGGTAGTGTGGAGAGGTGGG
997. $\overline{1}^* W_{\texttt{w11Aw12T}}\texttt{Q}^*$: GGATCGGAGATG GACGATAAAAAATAAAATAGCGAG GGGTAGTGTGGAGAGGTGGG
998. $\overline{1}^* X_{\texttt{w8Cw9T}}\texttt{Q}^*$: GGATCGGAGATG AGGCTTTTCTAAAAGAAGTTTTGC GGGTAGTGTGGAGAGGTGGG
999. $\overline{1}^* Y_{\texttt{w7Cw8C}}\texttt{Q}^*$: GGATCGGAGATG CAGAGGGCCTAATAGTAAAATGTT GGGTAGTGTGGAGAGGTGGG

##### S10.3.3 25 position Addition DNA sequences

1000. $\overline{0}B3^*$: CCTCTTCTCAGC CCTGCTCCATGTTACTTAGCCGGA GACAAGGGTTGT
1001. $\overline{3}B4^*$: TCAATCCTTGCC CCTGCTCCATGTTACTTAGCCGGA TGTTGGGAACAG
1002. $\overline{3}B5^*$: TCAATCCTTGCC CCTGCTCCATGTTACTTAGCCGGA GGACTGGTAGTG
1003. $\overline{2}B3^*$: TCTTTCCAAGCC CCTGCTCCATGTTACTTAGCCGGA GACAAGGGTTGT
1004. $\overline{1}B2^*$: CATCTCCGATCC CCTGCTCCATGTTACTTAGCCGGA AATGGTGGCATT
1005. $\overline{2}B1^*$: TCTTTCCAAGCC CCTGCTCCATGTTACTTAGCCGGA GAACGGAGTTGA
1006. $\overline{4}B3^*$: CACATCCCTGTT CCTGCTCCATGTTACTTAGCCGGA GACAAGGGTTGT
1007. $\overline{7}B4^*$: TCACACTTCGTC CCTGCTCCATGTTACTTAGCCGGA TGTTGGGAACAG
1008. $\overline{7}B5^*$: TCACACTTCGTC CCTGCTCCATGTTACTTAGCCGGA GGACTGGTAGTG
1009. $\overline{6}B7^*$: CAACCAACGTTC CCTGCTCCATGTTACTTAGCCGGA GGACGAAAGTGA
1010. $\overline{5}B6^*$: CCATGTCCCATT CCTGCTCCATGTTACTTAGCCGGA GACAGTGTGTGT
1011. $\overline{6}B5^*$: CAACCAACGTTC CCTGCTCCATGTTACTTAGCCGGA GGACTGGTAGTG
1012. $0C\overline{3}^*$: CTCATCCTGACC ACGAGGCGCAGACGGTCAATCATA GGCAAGGATTGA
1013. $3C\overline{4}^*$: ACAACCCTTGTC ACGAGGCGCAGACGGTCAATCATA AACAGGGATGTG
1014. $3C\overline{5}^*$: ACAACCCTTGTC ACGAGGCGCAGACGGTCAATCATA AATGGGACATGG
1015. $2C\overline{3}^*$: AATGCCACCATT ACGAGGCGCAGACGGTCAATCATA GGCAAGGATTGA
1016. $1C\overline{2}^*$: TCAACTCCGTTC ACGAGGCGCAGACGGTCAATCATA GGCTTGGAAAGA
1017. $2C\overline{1}^*$: AATGCCACCATT ACGAGGCGCAGACGGTCAATCATA GGATCGGAGATG
1018. $4C\overline{3}^*$: CTGTTCCCAACA ACGAGGCGCAGACGGTCAATCATA GGCAAGGATTGA
1019. $7C\overline{4}^*$: TCACTTTCGTCC ACGAGGCGCAGACGGTCAATCATA AACAGGGATGTG
1020. $7C\overline{5}^*$: TCACTTTCGTCC ACGAGGCGCAGACGGTCAATCATA AATGGGACATGG
1021. $6C\overline{7}^*$: ACACACACTGTC ACGAGGCGCAGACGGTCAATCATA GACGAAGTGTGA
1022. $5C\overline{6}^*$: CACTACCAGTCC ACGAGGCGCAGACGGTCAATCATA GAACGTTGGTTG
1023. $6C\overline{5}^*$: ACACACACTGTC ACGAGGCGCAGACGGTCAATCATA AATGGGACATGG
1024. $\overline{0}D3^*$: CCTCTTCTCAGC AGGGAACCGAACTGACCAACTTTG GACAAGGGTTGT
1025. $\overline{3}D4^*$: TCAATCCTTGCC AGGGAACCGAACTGACCAACTTTG TGTTGGGAACAG
1026. $\overline{3}D5^*$: TCAATCCTTGCC AGGGAACCGAACTGACCAACTTTG GGACTGGTAGTG
1027. $\overline{2}D3^*$: TCTTTCCAAGCC AGGGAACCGAACTGACCAACTTTG GACAAGGGTTGT
1028. $\overline{1}D2^*$: CATCTCCGATCC AGGGAACCGAACTGACCAACTTTG AATGGTGGCATT
1029. $\overline{2}D1^*$: TCTTTCCAAGCC AGGGAACCGAACTGACCAACTTTG GAACGGAGTTGA
1030. $\overline{4}D3^*$: CACATCCCTGTT AGGGAACCGAACTGACCAACTTTG GACAAGGGTTGT
1031. $\overline{7}D4^*$: TCACACTTCGTC AGGGAACCGAACTGACCAACTTTG TGTTGGGAACAG
1032. $\overline{7}D5^*$: TCACACTTCGTC AGGGAACCGAACTGACCAACTTTG GGACTGGTAGTG
1033. $\overline{6}D7^*$: CAACCAACGTTC AGGGAACCGAACTGACCAACTTTG GGACGAAAGTGA
1034. $\overline{5}D6^*$: CCATGTCCCATT AGGGAACCGAACTGACCAACTTTG GACAGTGTGTGT
1035. $\overline{6}D5^*$: CAACCAACGTTC AGGGAACCGAACTGACCAACTTTG GGACTGGTAGTG
1036. $0E\overline{3}^*$: CTCATCCTGACC AAAGAGGACAGATGAACGGTGTAC GGCAAGGATTGA
1037. $3E\overline{4}^*$: ACAACCCTTGTC AAAGAGGACAGATGAACGGTGTAC AACAGGGATGTG
1038. $3E\overline{5}^*$: ACAACCCTTGTC AAAGAGGACAGATGAACGGTGTAC AATGGGACATGG
1039. $2E\overline{3}^*$: AATGCCACCATT AAAGAGGACAGATGAACGGTGTAC GGCAAGGATTGA
1040. $1E\overline{2}^*$: TCAACTCCGTTC AAAGAGGACAGATGAACGGTGTAC GGCTTGGAAAGA
1041. $2E\overline{1}^*$: AATGCCACCATT AAAGAGGACAGATGAACGGTGTAC GGATCGGAGATG
1042. $4E\overline{3}^*$: CTGTTCCCAACA AAAGAGGACAGATGAACGGTGTAC GGCAAGGATTGA
1043. $7E\overline{4}^*$: TCACTTTCGTCC AAAGAGGACAGATGAACGGTGTAC AACAGGGATGTG
1044. $7E\overline{5}^*$: TCACTTTCGTCC AAAGAGGACAGATGAACGGTGTAC AATGGGACATGG
1045. $6E\overline{7}^*$: ACACACACTGTC AAAGAGGACAGATGAACGGTGTAC GACGAAGTGTGA

1046. $5E\overline{6}^*$: CACTACCAGTCC AAAGAGGACAGATGAACGGTGTAC GAACGTTGGTTG
1047. $6E\overline{5}^*$: ACACACACTGTC AAAGAGGACAGATGAACGGTGTAC AATGGGACATGG
1048. $\overline{0}F3^*$: CCTCTTCTCAGC AGACCAGGCGCATAGGCTGGCTGA GACAAGGGTTGT
1049. $\overline{3}F4^*$: TCAATCCTTGCC AGACCAGGCGCATAGGCTGGCTGA TGTTGGGAACAG
1050. $\overline{3}F5^*$: TCAATCCTTGCC AGACCAGGCGCATAGGCTGGCTGA GGACTGGTAGTG
1051. $\overline{2}F3^*$: TCTTTCCAAGCC AGACCAGGCGCATAGGCTGGCTGA GACAAGGGTTGT
1052. $\overline{1}F2^*$: CATCTCCGATCC AGACCAGGCGCATAGGCTGGCTGA AATGGTGGCATT
1053. $\overline{2}F1^*$: TCTTTCCAAGCC AGACCAGGCGCATAGGCTGGCTGA GAACGGAGTTGA
1054. $\overline{4}F3^*$: CACATCCCTGTT AGACCAGGCGCATAGGCTGGCTGA GACAAGGGTTGT
1055. $\overline{7}F4^*$: TCACACTTCGTC AGACCAGGCGCATAGGCTGGCTGA TGTTGGGAACAG
1056. $\overline{7}F5^*$: TCACACTTCGTC AGACCAGGCGCATAGGCTGGCTGA GGACTGGTAGTG
1057. $\overline{6}F7^*$: CAACCAACGTTC AGACCAGGCGCATAGGCTGGCTGA GGACGAAAGTGA
1058. $\overline{5}F6^*$: CCATGTCCCATT AGACCAGGCGCATAGGCTGGCTGA GACAGTGTGTGT
1059. $\overline{6}F5^*$: CAACCAACGTTC AGACCAGGCGCATAGGCTGGCTGA GGACTGGTAGTG
1060. $0G\overline{3}^*$: CTCATCCTGACC CCTTCATCAAGAGTAATCTTGACA GGCAAGGATTGA
1061. $3G\overline{4}^*$: ACAACCCTTGTC CCTTCATCAAGAGTAATCTTGACA AACAGGGATGTG
1062. $3G\overline{5}^*$: ACAACCCTTGTC CCTTCATCAAGAGTAATCTTGACA AATGGGACATGG
1063. $2G\overline{3}^*$: AATGCCACCATT CCTTCATCAAGAGTAATCTTGACA GGCAAGGATTGA
1064. $1G\overline{2}^*$: TCAACTCCGTTC CCTTCATCAAGAGTAATCTTGACA GGCTTGGAAAGA
1065. $2G\overline{1}^*$: AATGCCACCATT CCTTCATCAAGAGTAATCTTGACA GGATCGGAGATG
1066. $4G\overline{3}^*$: CTGTTCCCAACA CCTTCATCAAGAGTAATCTTGACA GGCAAGGATTGA
1067. $7G\overline{4}^*$: TCACTTTCGTCC CCTTCATCAAGAGTAATCTTGACA AACAGGGATGTG
1068. $7G\overline{5}^*$: TCACTTTCGTCC CCTTCATCAAGAGTAATCTTGACA AATGGGACATGG
1069. $6G\overline{7}^*$: ACACACACTGTC CCTTCATCAAGAGTAATCTTGACA GACGAAGTGTGA
1070. $5G\overline{6}^*$: CACTACCAGTCC CCTTCATCAAGAGTAATCTTGACA GAACGTTGGTTG
1071. $6G\overline{5}^*$: ACACACACTGTC CCTTCATCAAGAGTAATCTTGACA AATGGGACATGG
1072. $\overline{0}H3^*$: CCTCTTCTCAGC AGAACCGGATATTCATTACCCAAA GACAAGGGTTGT
1073. $\overline{3}H4^*$: TCAATCCTTGCC AGAACCGGATATTCATTACCCAAA TGTTGGGAACAG
1074. $\overline{3}H5^*$: TCAATCCTTGCC AGAACCGGATATTCATTACCCAAA GGACTGGTAGTG
1075. $\overline{2}H3^*$: TCTTTCCAAGCC AGAACCGGATATTCATTACCCAAA GACAAGGGTTGT
1076. $\overline{1}H2^*$: CATCTCCGATCC AGAACCGGATATTCATTACCCAAA AATGGTGGCATT
1077. $\overline{2}H1^*$: TCTTTCCAAGCC AGAACCGGATATTCATTACCCAAA GAACGGAGTTGA
1078. $\overline{4}H3^*$: CACATCCCTGTT AGAACCGGATATTCATTACCCAAA GACAAGGGTTGT
1079. $\overline{7}H4^*$: TCACACTTCGTC AGAACCGGATATTCATTACCCAAA TGTTGGGAACAG
1080. $\overline{7}H5^*$: TCACACTTCGTC AGAACCGGATATTCATTACCCAAA GGACTGGTAGTG
1081. $\overline{6}H7^*$: CAACCAACGTTC AGAACCGGATATTCATTACCCAAA GGACGAAAGTGA
1082. $\overline{5}H6^*$: CCATGTCCCATT AGAACCGGATATTCATTACCCAAA GACAGTGTGTGT
1083. $\overline{6}H5^*$: CAACCAACGTTC AGAACCGGATATTCATTACCCAAA GGACTGGTAGTG
1084. $0I\overline{3}^*$: CTCATCCTGACC TCAACGTAACAAAGCTGCTCATTC GGCAAGGATTGA
1085. $3I\overline{4}^*$: ACAACCCTTGTC TCAACGTAACAAAGCTGCTCATTC AACAGGGATGTG
1086. $3I\overline{5}^*$: ACAACCCTTGTC TCAACGTAACAAAGCTGCTCATTC AATGGGACATGG
1087. $2I\overline{3}^*$: AATGCCACCATT TCAACGTAACAAAGCTGCTCATTC GGCAAGGATTGA
1088. $1I\overline{2}^*$: TCAACTCCGTTC TCAACGTAACAAAGCTGCTCATTC GGCTTGGAAAGA
1089. $2I\overline{1}^*$: AATGCCACCATT TCAACGTAACAAAGCTGCTCATTC GGATCGGAGATG
1090. $4I\overline{3}^*$: CTGTTCCCAACA TCAACGTAACAAAGCTGCTCATTC GGCAAGGATTGA
1091. $7I\overline{4}^*$: TCACTTTCGTCC TCAACGTAACAAAGCTGCTCATTC AACAGGGATGTG
1092. $7I\overline{5}^*$: TCACTTTCGTCC TCAACGTAACAAAGCTGCTCATTC AATGGGACATGG
1093. $6I\overline{7}^*$: ACACACACTGTC TCAACGTAACAAAGCTGCTCATTC GACGAAGTGTGA
1094. $5I\overline{6}^*$: CACTACCAGTCC TCAACGTAACAAAGCTGCTCATTC GAACGTTGGTTG
1095. $6I\overline{5}^*$: ACACACACTGTC TCAACGTAACAAAGCTGCTCATTC AATGGGACATGG
1096. $\overline{0}J3^*$: CCTCTTCTCAGC AGTGAATAAGGCTTGCCCTGACGA GACAAGGGTTGT
1097. $\overline{3}J4^*$: TCAATCCTTGCC AGTGAATAAGGCTTGCCCTGACGA TGTTGGGAACAG
1098. $\overline{3}J5^*$: TCAATCCTTGCC AGTGAATAAGGCTTGCCCTGACGA GGACTGGTAGTG
1099. $\overline{2}J3^*$: TCTTTCCAAGCC AGTGAATAAGGCTTGCCCTGACGA GACAAGGGTTGT

1100. $\overline{1}J2^*$: CATCTCCGATCC AGTGAATAAGGCTTGCCCTGACGA AATGGTGGCATT

1101. $\overline{2}J1^*$: TCTTTCCAAGCC AGTGAATAAGGCTTGCCCTGACGA GAACGGAGTTGA

1102. $\overline{4}J3^*$: CACATCCCTGTT AGTGAATAAGGCTTGCCCTGACGA GACAAGGGTTGT

1103. $\overline{7}J4^*$: TCACACTTCGTC AGTGAATAAGGCTTGCCCTGACGA TGTTGGGAACAG

1104. $\overline{7}J5^*$: TCACACTTCGTC AGTGAATAAGGCTTGCCCTGACGA GGACTGGTAGTG

1105. $\overline{6}J7^*$: CAACCAACGTTC AGTGAATAAGGCTTGCCCTGACGA GGACGAAAGTGA

1106. $\overline{5}J6^*$: CCATGTCCCATT AGTGAATAAGGCTTGCCCTGACGA GACAGTGTGTGT

1107. $\overline{6}J5^*$: CAACCAACGTTC AGTGAATAAGGCTTGCCCTGACGA GGACTGGTAGTG

1108. $0K\overline{3}^*$: CTCATCCTGACC GAAACACCAGAACGAGTAGTAAAT GGCAAGGATTGA

1109. $3K\overline{4}^*$: ACAACCCTTGTC GAAACACCAGAACGAGTAGTAAAT AACAGGGATGTG

1110. $3K\overline{5}^*$: ACAACCCTTGTC GAAACACCAGAACGAGTAGTAAAT AATGGGACATGG

1111. $2K\overline{3}^*$: AATGCCACCATT GAAACACCAGAACGAGTAGTAAAT GGCAAGGATTGA

1112. $1K\overline{2}^*$: TCAACTCCGTTC GAAACACCAGAACGAGTAGTAAAT GGCTTGGAAAGA

1113. $2K\overline{1}^*$: AATGCCACCATT GAAACACCAGAACGAGTAGTAAAT GGATCGGAGATG

1114. $4K\overline{3}^*$: CTGTTCCCAACA GAAACACCAGAACGAGTAGTAAAT GGCAAGGATTGA

1115. $7K\overline{4}^*$: TCACTTTCGTCC GAAACACCAGAACGAGTAGTAAAT AACAGGGATGTG

1116. $7K\overline{5}^*$: TCACTTTCGTCC GAAACACCAGAACGAGTAGTAAAT AATGGGACATGG

1117. $6K\overline{7}^*$: ACACACACTGTC GAAACACCAGAACGAGTAGTAAAT GACGAAGTGTGA

1118. $5K\overline{6}^*$: CACTACCAGTCC GAAACACCAGAACGAGTAGTAAAT GAACGTTGGTTG

1119. $6K\overline{5}^*$: ACACACACTGTC GAAACACCAGAACGAGTAGTAAAT AATGGGACATGG

1120. $\overline{0}L3^*$: CCTCTTCTCAGC TGGGCTTGAGATGGTTTAATTTCA GACAAGGGTTGT

1121. $\overline{3}L4^*$: TCAATCCTTGCC TGGGCTTGAGATGGTTTAATTTCA TGTTGGGAACAG

1122. $\overline{3}L5^*$: TCAATCCTTGCC TGGGCTTGAGATGGTTTAATTTCA GGACTGGTAGTG

1123. $\overline{2}L3^*$: TCTTTCCAAGCC TGGGCTTGAGATGGTTTAATTTCA GACAAGGGTTGT

1124. $\overline{1}L2^*$: CATCTCCGATCC TGGGCTTGAGATGGTTTAATTTCA AATGGTGGCATT

1125. $\overline{2}L1^*$: TCTTTCCAAGCC TGGGCTTGAGATGGTTTAATTTCA GAACGGAGTTGA

1126. $\overline{4}L3^*$: CACATCCCTGTT TGGGCTTGAGATGGTTTAATTTCA GACAAGGGTTGT

1127. $\overline{7}L4^*$: TCACACTTCGTC TGGGCTTGAGATGGTTTAATTTCA TGTTGGGAACAG

1128. $\overline{7}L5^*$: TCACACTTCGTC TGGGCTTGAGATGGTTTAATTTCA GGACTGGTAGTG

1129. $\overline{6}L7^*$: CAACCAACGTTC TGGGCTTGAGATGGTTTAATTTCA GGACGAAAGTGA

1130. $\overline{5}L6^*$: CCATGTCCCATT TGGGCTTGAGATGGTTTAATTTCA GACAGTGTGTGT

1131. $\overline{6}L5^*$: CAACCAACGTTC TGGGCTTGAGATGGTTTAATTTCA GGACTGGTAGTG

1132. $0M\overline{3}^*$: CTCATCCTGACC ACTTTAATCATTGTGAATTACCTT GGCAAGGATTGA

1133. $3M\overline{4}^*$: ACAACCCTTGTC ACTTTAATCATTGTGAATTACCTT AACAGGGATGTG

1134. $3M\overline{5}^*$: ACAACCCTTGTC ACTTTAATCATTGTGAATTACCTT AATGGGACATGG

1135. $2M\overline{3}^*$: AATGCCACCATT ACTTTAATCATTGTGAATTACCTT GGCAAGGATTGA

1136. $1M\overline{2}^*$: TCAACTCCGTTC ACTTTAATCATTGTGAATTACCTT GGCTTGGAAAGA

1137. $2M\overline{1}^*$: AATGCCACCATT ACTTTAATCATTGTGAATTACCTT GGATCGGAGATG

1138. $4M\overline{3}^*$: CTGTTCCCAACA ACTTTAATCATTGTGAATTACCTT GGCAAGGATTGA

1139. $7M\overline{4}^*$: TCACTTTCGTCC ACTTTAATCATTGTGAATTACCTT AACAGGGATGTG

1140. $7M\overline{5}^*$: TCACTTTCGTCC ACTTTAATCATTGTGAATTACCTT AATGGGACATGG

1141. $6M\overline{7}^*$: ACACACACTGTC ACTTTAATCATTGTGAATTACCTT GACGAAGTGTGA

1142. $5M\overline{6}^*$: CACTACCAGTCC ACTTTAATCATTGTGAATTACCTT GAACGTTGGTTG

1143. $6M\overline{5}^*$: ACACACACTGTC ACTTTAATCATTGTGAATTACCTT AATGGGACATGG

1144. $\overline{0}N3^*$: CCTCTTCTCAGC ATGCGATTTTAAGAACTGGCTCAT GACAAGGGTTGT

1145. $\overline{3}N4^*$: TCAATCCTTGCC ATGCGATTTTAAGAACTGGCTCAT TGTTGGGAACAG

1146. $\overline{3}N5^*$: TCAATCCTTGCC ATGCGATTTTAAGAACTGGCTCAT GGACTGGTAGTG

1147. $\overline{2}N3^*$: TCTTTCCAAGCC ATGCGATTTTAAGAACTGGCTCAT GACAAGGGTTGT

1148. $\overline{1}N2^*$: CATCTCCGATCC ATGCGATTTTAAGAACTGGCTCAT AATGGTGGCATT

1149. $\overline{2}N1^*$: TCTTTCCAAGCC ATGCGATTTTAAGAACTGGCTCAT GAACGGAGTTGA

1150. $\overline{4}N3^*$: CACATCCCTGTT ATGCGATTTTAAGAACTGGCTCAT GACAAGGGTTGT

1151. $\overline{7}N4^*$: TCACACTTCGTC ATGCGATTTTAAGAACTGGCTCAT TGTTGGGAACAG

1152. $\overline{7}N5^*$: TCACACTTCGTC ATGCGATTTTAAGAACTGGCTCAT GGACTGGTAGTG

1153. $\overline{6}N7^*$: CAACCAACGTTC ATGCGATTTTAAGAACTGGCTCAT GGACGAAAGTGA

1154. $\overline{5}N6^*$: CCATGTCCCATT ATGCGATTTTAAGAACTGGCTCAT GACAGTGTGTGT
1155. $\overline{6}N5^*$: CAACCAACGTTC ATGCGATTTTAAGAACTGGCTCAT GGACTGGTAGTG
1156. $0O\overline{3}^*$: CTCATCCTGACC TATACCAGTCAGGACGTTGGGAAG GGCAAGGATTGA
1157. $3O\overline{4}^*$: ACAACCCTTGTC TATACCAGTCAGGACGTTGGGAAG AACAGGGATGTG
1158. $3O\overline{5}^*$: ACAACCCTTGTC TATACCAGTCAGGACGTTGGGAAG AATGGGACATGG
1159. $2O\overline{3}^*$: AATGCCACCATT TATACCAGTCAGGACGTTGGGAAG GGCAAGGATTGA
1160. $1O\overline{2}^*$: TCAACTCCGTTC TATACCAGTCAGGACGTTGGGAAG GGCTTGGAAAGA
1161. $2O\overline{1}^*$: AATGCCACCATT TATACCAGTCAGGACGTTGGGAAG GGATCGGAGATG
1162. $4O\overline{3}^*$: CTGTTCCCAACA TATACCAGTCAGGACGTTGGGAAG GGCAAGGATTGA
1163. $7O\overline{4}^*$: TCACACTTCGTC TATACCAGTCAGGACGTTGGGAAG AACAGGGATGTG
1164. $7O\overline{5}^*$: TCACACTTCGTC TATACCAGTCAGGACGTTGGGAAG AATGGGACATGG
1165. $6O\overline{7}^*$: ACACACACTGTC TATACCAGTCAGGACGTTGGGAAG GACGAAGTGTGA
1166. $5O\overline{6}^*$: CACTACCAGTCC TATACCAGTCAGGACGTTGGGAAG GAACGTTGGTTG
1167. $6O\overline{5}^*$: ACACACACTGTC TATACCAGTCAGGACGTTGGGAAG AATGGGACATGG
1168. $\overline{0}P3^*$: CCTCTTCTCAGC AAAAATCTACGTTAATAAAACGAA GACAAGGGTTGT
1169. $\overline{3}P4^*$: TCAATCCTTGCC AAAAATCTACGTTAATAAAACGAA TGTTGGGAACAG
1170. $\overline{3}P5^*$: TCAATCCTTGCC AAAAATCTACGTTAATAAAACGAA GGACTGGTAGTG
1171. $\overline{2}P3^*$: TCTTTCCAAGCC AAAAATCTACGTTAATAAAACGAA GACAAGGGTTGT
1172. $\overline{1}P2^*$: CATCTCCGATCC AAAAATCTACGTTAATAAAACGAA AATGGTGGCATT
1173. $\overline{2}P1^*$: TCTTTCCAAGCC AAAAATCTACGTTAATAAAACGAA GAACGGAGTTGA
1174. $\overline{4}P3^*$: CACATCCCTGTT AAAAATCTACGTTAATAAAACGAA GACAAGGGTTGT
1175. $\overline{7}P4^*$: TCACACTTCGTC AAAAATCTACGTTAATAAAACGAA TGTTGGGAACAG
1176. $\overline{7}P5^*$: TCACACTTCGTC AAAAATCTACGTTAATAAAACGAA GGACTGGTAGTG
1177. $\overline{6}P7^*$: CAACCAACGTTC AAAAATCTACGTTAATAAAACGAA GGACGAAAGTGA
1178. $\overline{5}P6^*$: CCATGTCCCATT AAAAATCTACGTTAATAAAACGAA GACAGTGTGTGT
1179. $\overline{6}P5^*$: CAACCAACGTTC AAAAATCTACGTTAATAAAACGAA GGACTGGTAGTG
1180. $0Q\overline{3}^*$: CTCATCCTGACC CTAACGGAACAACATTATTACAGG GGCAAGGATTGA
1181. $3Q\overline{4}^*$: ACAACCCTTGTC CTAACGGAACAACATTATTACAGG AACAGGGATGTG
1182. $3Q\overline{5}^*$: ACAACCCTTGTC CTAACGGAACAACATTATTACAGG AATGGGACATGG
1183. $2Q\overline{3}^*$: AATGCCACCATT CTAACGGAACAACATTATTACAGG GGCAAGGATTGA
1184. $1Q\overline{2}^*$: TCAACTCCGTTC CTAACGGAACAACATTATTACAGG GGCTTGGAAAGA
1185. $2Q\overline{1}^*$: AATGCCACCATT CTAACGGAACAACATTATTACAGG GGATCGGAGATG
1186. $4Q\overline{3}^*$: CTGTTCCCAACA CTAACGGAACAACATTATTACAGG GGCAAGGATTGA
1187. $7Q\overline{4}^*$: TCACTTTCGTCC CTAACGGAACAACATTATTACAGG AACAGGGATGTG
1188. $7Q\overline{5}^*$: TCACTTTCGTCC CTAACGGAACAACATTATTACAGG AATGGGACATGG
1189. $6Q\overline{7}^*$: ACACACACTGTC CTAACGGAACAACATTATTACAGG GACGAAGTGTGA
1190. $5Q\overline{6}^*$: CACTACCAGTCC CTAACGGAACAACATTATTACAGG GAACGTTGGTTG
1191. $6Q\overline{5}^*$: ACACACACTGTC CTAACGGAACAACATTATTACAGG AATGGGACATGG
1192. $\overline{0}R3^*$: CCTCTTCTCAGC TAGAAAGATTCATCAGTTGAGATT GACAAGGGTTGT
1193. $\overline{3}R4^*$: TCAATCCTTGCC TAGAAAGATTCATCAGTTGAGATT TGTTGGGAACAG
1194. $\overline{3}R5^*$: TCAATCCTTGCC TAGAAAGATTCATCAGTTGAGATT GGACTGGTAGTG
1195. $\overline{2}R3^*$: TCTTTCCAAGCC TAGAAAGATTCATCAGTTGAGATT GACAAGGGTTGT
1196. $\overline{1}R2^*$: CATCTCCGATCC TAGAAAGATTCATCAGTTGAGATT AATGGTGGCATT
1197. $\overline{2}R1^*$: TCTTTCCAAGCC TAGAAAGATTCATCAGTTGAGATT GAACGGAGTTGA
1198. $\overline{4}R3^*$: CACATCCCTGTT TAGAAAGATTCATCAGTTGAGATT GACAAGGGTTGT
1199. $\overline{7}R4^*$: TCACACTTCGTC TAGAAAGATTCATCAGTTGAGATT TGTTGGGAACAG
1200. $\overline{7}R5^*$: TCACACTTCGTC TAGAAAGATTCATCAGTTGAGATT GGACTGGTAGTG
1201. $\overline{6}R7^*$: CAACCAACGTTC TAGAAAGATTCATCAGTTGAGATT GGACGAAAGTGA
1202. $\overline{5}R6^*$: CCATGTCCCATT TAGAAAGATTCATCAGTTGAGATT GACAGTGTGTGT
1203. $\overline{6}R5^*$: CAACCAACGTTC TAGAAAGATTCATCAGTTGAGATT GGACTGGTAGTG
1204. $0S\overline{3}^*$: CTCATCCTGACC TAGGAATACCACATTCAACTAATG GGCAAGGATTGA
1205. $3S\overline{4}^*$: ACAACCCTTGTC TAGGAATACCACATTCAACTAATG AACAGGGATGTG
1206. $3S\overline{5}^*$: ACAACCCTTGTC TAGGAATACCACATTCAACTAATG AATGGGACATGG
1207. $2S\overline{3}^*$: AATGCCACCATT TAGGAATACCACATTCAACTAATG GGCAAGGATTGA

1208. $1S\overline{2}^*$: TCAACTCCGTTC TAGGAATACCACATTCAACTAATG GGCTTGGAAAGA
1209. $2S\overline{1}^*$: AATGCCACCATT TAGGAATACCACATTCAACTAATG GGATCGGAGATG
1210. $4S\overline{3}^*$: CTGTTCCCAACA TAGGAATACCACATTCAACTAATG GGCAAGGATTGA
1211. $7S\overline{4}^*$: TCACTTTCGTCC TAGGAATACCACATTCAACTAATG AACAGGGATGTG
1212. $7S\overline{5}^*$: TCACTTTCGTCC TAGGAATACCACATTCAACTAATG AATGGGACATGG
1213. $6S\overline{7}^*$: ACACACACTGTC TAGGAATACCACATTCAACTAATG GACGAAGTGTGA
1214. $5S\overline{6}^*$: CACTACCAGTCC TAGGAATACCACATTCAACTAATG GAACGTTGGTTG
1215. $6S\overline{5}^*$: ACACACACTGTC TAGGAATACCACATTCAACTAATG AATGGGACATGG
1216. $\overline{0}T3^*$: CCTCTTCTCAGC CAGATACATAACGCCAAAAGGAAT GACAAGGGTTGT
1217. $\overline{3}T4^*$: TCAATCCTTGCC CAGATACATAACGCCAAAAGGAAT TGTTGGGAACAG
1218. $\overline{3}T5^*$: TCAATCCTTGCC CAGATACATAACGCCAAAAGGAAT GGACTGGTAGTG
1219. $\overline{2}T3^*$: TCTTTCCAAGCC CAGATACATAACGCCAAAAGGAAT GACAAGGGTTGT
1220. $\overline{1}T2^*$: CATCTCCGATCC CAGATACATAACGCCAAAAGGAAT AATGGTGGCATT
1221. $\overline{2}T1^*$: TCTTTCCAAGCC CAGATACATAACGCCAAAAGGAAT GAACGGAGTTGA
1222. $\overline{4}T3^*$: CACATCCCTGTT CAGATACATAACGCCAAAAGGAAT GACAAGGGTTGT
1223. $\overline{7}T4^*$: TCACACTTCGTC CAGATACATAACGCCAAAAGGAAT TGTTGGGAACAG
1224. $\overline{7}T5^*$: TCACACTTCGTC CAGATACATAACGCCAAAAGGAAT GGACTGGTAGTG
1225. $\overline{6}T7^*$: CAACCAACGTTC CAGATACATAACGCCAAAAGGAAT GGACGAAAGTGA
1226. $\overline{5}T6^*$: CCATGTCCCATT CAGATACATAACGCCAAAAGGAAT GACAGTGTGTGT
1227. $\overline{6}T5^*$: CAACCAACGTTC CAGATACATAACGCCAAAAGGAAT GGACTGGTAGTG
1228. $0U\overline{3}^*$: CTCATCCTGACC TACGAGGCATAGTAAGAGCAACAC GGCAAGGATTGA
1229. $3U\overline{4}^*$: ACAACCCTTGTC TACGAGGCATAGTAAGAGCAACAC AACAGGGATGTG
1230. $3U\overline{5}^*$: ACAACCCTTGTC TACGAGGCATAGTAAGAGCAACAC AATGGGACATGG
1231. $2U\overline{3}^*$: AATGCCACCATT TACGAGGCATAGTAAGAGCAACAC GGCAAGGATTGA
1232. $1U\overline{2}^*$: TCAACTCCGTTC TACGAGGCATAGTAAGAGCAACAC GGCTTGGAAAGA
1233. $2U\overline{1}^*$: AATGCCACCATT TACGAGGCATAGTAAGAGCAACAC GGATCGGAGATG
1234. $4U\overline{3}^*$: CTGTTCCCAACA TACGAGGCATAGTAAGAGCAACAC GGCAAGGATTGA
1235. $7U\overline{4}^*$: TCACTTTCGTCC TACGAGGCATAGTAAGAGCAACAC AACAGGGATGTG
1236. $7U\overline{5}^*$: TCACTTTCGTCC TACGAGGCATAGTAAGAGCAACAC AATGGGACATGG
1237. $6U\overline{7}^*$: ACACACACTGTC TACGAGGCATAGTAAGAGCAACAC GACGAAGTGTGA
1238. $5U\overline{6}^*$: CACTACCAGTCC TACGAGGCATAGTAAGAGCAACAC GAACGTTGGTTG
1239. $6U\overline{5}^*$: ACACACACTGTC TACGAGGCATAGTAAGAGCAACAC AATGGGACATGG
1240. $\overline{0}V3*^*$: CCTCTTCTCAGC TATCATAACCCTCGTTTACCAGAC GACAAGGGTTGT
1241. $\overline{3}V4*^*$: TCAATCCTTGCC TATCATAACCCTCGTTTACCAGAC TGTTGGGAACAG
1242. $\overline{3}V5*^*$: TCAATCCTTGCC TATCATAACCCTCGTTTACCAGAC GGACTGGTAGTG
1243. $\overline{2}V3*^*$: TCTTTCCAAGCC TATCATAACCCTCGTTTACCAGAC GACAAGGGTTGT
1244. $\overline{1}V2*^*$: CATCTCCGATCC TATCATAACCCTCGTTTACCAGAC AATGGTGGCATT
1245. $\overline{2}V1*^*$: TCTTTCCAAGCC TATCATAACCCTCGTTTACCAGAC GAACGGAGTTGA
1246. $\overline{4}V3*^*$: CACATCCCTGTT TATCATAACCCTCGTTTACCAGAC GACAAGGGTTGT
1247. $\overline{7}V4*^*$: TCACACTTCGTC TATCATAACCCTCGTTTACCAGAC TGTTGGGAACAG
1248. $\overline{7}V5*^*$: TCACACTTCGTC TATCATAACCCTCGTTTACCAGAC GGACTGGTAGTG
1249. $\overline{6}V7*^*$: CAACCAACGTTC TATCATAACCCTCGTTTACCAGAC GGACGAAAGTGA
1250. $\overline{5}V6*^*$: CCATGTCCCATT TATCATAACCCTCGTTTACCAGAC GACAGTGTGTGT
1251. $\overline{6}V5*^*$: CAACCAACGTTC TATCATAACCCTCGTTTACCAGAC GGACTGGTAGTG
1252. $0W\overline{3}^*$: CTCATCCTGACC GACGATAAAAACCAAAATAGCGAG GGCAAGGATTGA
1253. $3W\overline{4}^*$: ACAACCCTTGTC GACGATAAAAACCAAAATAGCGAG AACAGGGATGTG
1254. $3W\overline{5}^*$: ACAACCCTTGTC GACGATAAAAACCAAAATAGCGAG AATGGGACATGG
1255. $2W\overline{3}^*$: AATGCCACCATT GACGATAAAAACCAAAATAGCGAG GGCAAGGATTGA
1256. $1W\overline{2}^*$: TCAACTCCGTTC GACGATAAAAACCAAAATAGCGAG GGCTTGGAAAGA
1257. $2W\overline{1}^*$: AATGCCACCATT GACGATAAAAACCAAAATAGCGAG GGATCGGAGATG
1258. $4W\overline{3}^*$: CTGTTCCCAACA GACGATAAAAACCAAAATAGCGAG GGCAAGGATTGA
1259. $7W\overline{4}^*$: TCACTTTCGTCC GACGATAAAAACCAAAATAGCGAG AACAGGGATGTG
1260. $7W\overline{5}^*$: TCACTTTCGTCC GACGATAAAAACCAAAATAGCGAG AATGGGACATGG
1261. $6W\overline{7}^*$: ACACACACTGTC GACGATAAAAACCAAAATAGCGAG GACGAAGTGTGA

1262. $5W\overline{6}^*$: CACTACCAGTCC GACGATAAAAACCAAAATAGCGAG GAACGTTGGTTG
1263. $6W\overline{5}^*$: ACACACACTGTC GACGATAAAAACCAAAATAGCGAG AATGGGACATGG
1264. $\overline{0}X3{*}^*$: CCTCTTCTCAGC AGGCTTTTGCAAAAGAAGTTTTGC GACAAGGGTTGT
1265. $\overline{3}X4{*}^*$: TCAATCCTTGCC AGGCTTTTGCAAAAGAAGTTTTGC TGTTGGGAACAG
1266. $\overline{3}X5{*}^*$: TCAATCCTTGCC AGGCTTTTGCAAAAGAAGTTTTGC GGACTGGTAGTG
1267. $\overline{2}X3{*}^*$: TCTTTCCAAGCC AGGCTTTTGCAAAAGAAGTTTTGC GACAAGGGTTGT
1268. $\overline{1}X2{*}^*$: CATCTCCGATCC AGGCTTTTGCAAAAGAAGTTTTGC AATGGTGGCATT
1269. $\overline{2}X1{*}^*$: TCTTTCCAAGCC AGGCTTTTGCAAAAGAAGTTTTGC GAACGGAGTTGA
1270. $\overline{4}X3{*}^*$: CACATCCCTGTT AGGCTTTTGCAAAAGAAGTTTTGC GACAAGGGTTGT
1271. $\overline{7}X4{*}^*$: TCACACTTCGTC AGGCTTTTGCAAAAGAAGTTTTGC TGTTGGGAACAG
1272. $\overline{7}X5{*}^*$: TCACACTTCGTC AGGCTTTTGCAAAAGAAGTTTTGC GGACTGGTAGTG
1273. $\overline{6}X7{*}^*$: CAACCAACGTTC AGGCTTTTGCAAAAGAAGTTTTGC GGACGAAAGTGA
1274. $\overline{5}X6{*}^*$: CCATGTCCCATT AGGCTTTTGCAAAAGAAGTTTTGC GACAGTGTGTGT
1275. $\overline{6}X5{*}^*$: CAACCAACGTTC AGGCTTTTGCAAAAGAAGTTTTGC GGACTGGTAGTG
1276. $0Y\overline{3}^*$: CTCATCCTGACC CAGAGGGGGTAATAGTAAAATGTT GGCAAGGATTGA
1277. $3Y\overline{4}^*$: ACAACCCTTGTC CAGAGGGGGTAATAGTAAAATGTT AACAGGGATGTG
1278. $3Y\overline{5}^*$: ACAACCCTTGTC CAGAGGGGGTAATAGTAAAATGTT AATGGGACATGG
1279. $2Y\overline{3}^*$: AATGCCACCATT CAGAGGGGGTAATAGTAAAATGTT GGCAAGGATTGA
1280. $1Y\overline{2}^*$: TCAACTCCGTTC CAGAGGGGGTAATAGTAAAATGTT GGCTTGGAAAGA
1281. $2Y\overline{1}^*$: AATGCCACCATT CAGAGGGGGTAATAGTAAAATGTT GGATCGGAGATG
1282. $4Y\overline{3}^*$: CTGTTCCCAACA CAGAGGGGGTAATAGTAAAATGTT GGCAAGGATTGA
1283. $7Y\overline{4}^*$: TCACTTTCGTCC CAGAGGGGGTAATAGTAAAATGTT AACAGGGATGTG
1284. $7Y\overline{5}^*$: TCACTTTCGTCC CAGAGGGGGTAATAGTAAAATGTT AATGGGACATGG
1285. $6Y\overline{7}^*$: ACACACACTGTC CAGAGGGGGTAATAGTAAAATGTT GACGAAGTGTGA
1286. $5Y\overline{6}^*$: CACTACCAGTCC CAGAGGGGGTAATAGTAAAATGTT GAACGTTGGTTG
1287. $6Y\overline{5}^*$: ACACACACTGTC CAGAGGGGGTAATAGTAAAATGTT AATGGGACATGG
1288. $\overline{0}B\mathtt{Q}^*$: CCTCTTCTCAGC CCTGCTCCATGTTACTTAGCCGGA GGGTAGTGTGGAGAGGTGGG
1289. $\overline{3}B\mathtt{Q}^*$: TCAATCCTTGCC CCTGCTCCATGTTACTTAGCCGGA GGGTAGTGTGGAGAGGTGGG
1290. $\overline{5}B\mathtt{Q}^*$: CCATGTCCCATT CCTGCTCCATGTTACTTAGCCGGA GGGTAGTGTGGAGAGGTGGG
1291. $\overline{6}B\mathtt{Q}^*$: CAACCAACGTTC CCTGCTCCATGTTACTTAGCCGGA GGGTAGTGTGGAGAGGTGGG
1292. $\overline{4}B$: CACATCCCTGTT CCTGCTCCATGTTACTTAGCCGGA
1293. $\overline{7}B$: TCACACTTCGTC CCTGCTCCATGTTACTTAGCCGGA
1294. $\overline{1}B$: CATCTCCGATCC CCTGCTCCATGTTACTTAGCCGGA
1295. $\overline{2}B$: TCTTTCCAAGCC CCTGCTCCATGTTACTTAGCCGGA
1296. $A$: GATAAATTGTGTCGAAATCCGCGA
1297. $B$: CCTGCTCCATGTTACTTAGCCGGA
1298. $A\overline{0}^*$: GATAAATTGTGTCGAAATCCGCGA GCTGAGAAGAGG
1299. $B0^*$: CCTGCTCCATGTTACTTAGCCGGA GGTCAGGATGAG
1300. $0CQ^*$: CTCATCCTGACC ACGAGGCGCAGACGGTCAATCATA GGGTAGTGTGGAGAGGTGGG
1301. $3CQ^*$: ACAACCCTTGTC ACGAGGCGCAGACGGTCAATCATA GGGTAGTGTGGAGAGGTGGG
1302. $5CQ^*$: CACTACCAGTCC ACGAGGCGCAGACGGTCAATCATA GGGTAGTGTGGAGAGGTGGG
1303. $6CQ^*$: ACACACACTGTC ACGAGGCGCAGACGGTCAATCATA GGGTAGTGTGGAGAGGTGGG
1304. $4C$: CTGTTCCCAACA ACGAGGCGCAGACGGTCAATCATA
1305. $7C$: TCACTTTCGTCC ACGAGGCGCAGACGGTCAATCATA
1306. $1C$: TCAACTCCGTTC ACGAGGCGCAGACGGTCAATCATA
1307. $2C$: AATGCCACCATT ACGAGGCGCAGACGGTCAATCATA
1308. $AQ^*$: GATAAATTGTGTCGAAATCCGCGA GGGTAGTGTGGAGAGGTGGG
1309. $BQ^*$: CCTGCTCCATGTTACTTAGCCGGA GGGTAGTGTGGAGAGGTGGG
1310. $A\overline{1}^*$: GATAAATTGTGTCGAAATCCGCGA GGATCGGAGATG
1311. $B1^*$: CCTGCTCCATGTTACTTAGCCGGA GAACGGAGTTGA
1312. $\overline{0}D\mathtt{Q}^*$: CCTCTTCTCAGC AGGGAACCGAACTGACCAACTTTG GGGTAGTGTGGAGAGGTGGG
1313. $\overline{3}D\mathtt{Q}^*$: TCAATCCTTGCC AGGGAACCGAACTGACCAACTTTG GGGTAGTGTGGAGAGGTGGG
1314. $\overline{5}D\mathtt{Q}^*$: CCATGTCCCATT AGGGAACCGAACTGACCAACTTTG GGGTAGTGTGGAGAGGTGGG
1315. $\overline{6}D\mathtt{Q}^*$: CAACCAACGTTC AGGGAACCGAACTGACCAACTTTG GGGTAGTGTGGAGAGGTGGG

1316. $\overline{4}D$: CACATCCCTGTT AGGGAACCGAACTGACCAACTTTG
1317. $\overline{7}D$: TCACACTTCGTC AGGGAACCGAACTGACCAACTTTG
1318. $\overline{1}D$: CATCTCCGATCC AGGGAACCGAACTGACCAACTTTG
1319. $\overline{2}D$: TCTTTCCAAGCC AGGGAACCGAACTGACCAACTTTG
1320. $A\overline{2}^*$: GATAAATTGTGTCGAAATCCGCGA GGCTTGGAAAGA
1321. $B2^*$: CCTGCTCCATGTTACTTAGCCGGA AATGGTGGCATT
1322. $0EQ^*$: CTCATCCTGACC AAAGAGGACAGATGAACGGTGTAC GGGTAGTGTGGAGAGGTGGG
1323. $3EQ^*$: ACAACCCTTGTC AAAGAGGACAGATGAACGGTGTAC GGGTAGTGTGGAGAGGTGGG
1324. $5EQ^*$: CACTACCAGTCC AAAGAGGACAGATGAACGGTGTAC GGGTAGTGTGGAGAGGTGGG
1325. $6EQ^*$: ACACACACTGTC AAAGAGGACAGATGAACGGTGTAC GGGTAGTGTGGAGAGGTGGG
1326. $4E$: CTGTTCCCAACA AAAGAGGACAGATGAACGGTGTAC
1327. $7E$: TCACACTTCGTC AAAGAGGACAGATGAACGGTGTAC
1328. $1E$: TCAACTCCGTTC AAAGAGGACAGATGAACGGTGTAC
1329. $2E$: AATGCCACCATT AAAGAGGACAGATGAACGGTGTAC
1330. $A\overline{3}^*$: GATAAATTGTGTCGAAATCCGCGA GGCAAGGATTGA
1331. $B3^*$: CCTGCTCCATGTTACTTAGCCGGA GACAAGGGTTGT
1332. $\overline{0}F\mathtt{Q}^*$: CCTCTTCTCAGC AGACCAGGCGCATAGGCTGGCTGA GGGTAGTGTGGAGAGGTGGG
1333. $\overline{3}F\mathtt{Q}^*$: TCAATCCTTGCC AGACCAGGCGCATAGGCTGGCTGA GGGTAGTGTGGAGAGGTGGG
1334. $\overline{5}F\mathtt{Q}^*$: CCATGTCCCATT AGACCAGGCGCATAGGCTGGCTGA GGGTAGTGTGGAGAGGTGGG
1335. $\overline{6}F\mathtt{Q}^*$: CAACCAACGTTC AGACCAGGCGCATAGGCTGGCTGA GGGTAGTGTGGAGAGGTGGG
1336. $\overline{4}F$: CACATCCCTGTT AGACCAGGCGCATAGGCTGGCTGA
1337. $\overline{7}F$: TCACACTTCGTC AGACCAGGCGCATAGGCTGGCTGA
1338. $\overline{1}F$: CATCTCCGATCC AGACCAGGCGCATAGGCTGGCTGA
1339. $\overline{2}F$: TCTTTCCAAGCC AGACCAGGCGCATAGGCTGGCTGA
1340. $A\overline{4}^*$: GATAAATTGTGTCGAAATCCGCGA AACAGGGATGTG
1341. $B4^*$: CCTGCTCCATGTTACTTAGCCGGA TGTTGGGAACAG
1342. $0GQ^*$: CTCATCCTGACC CCTTCATCAAGAGTAATCTTGACA GGGTAGTGTGGAGAGGTGGG
1343. $3GQ^*$: ACAACCCTTGTC CCTTCATCAAGAGTAATCTTGACA GGGTAGTGTGGAGAGGTGGG
1344. $5GQ^*$: CACTACCAGTCC CCTTCATCAAGAGTAATCTTGACA GGGTAGTGTGGAGAGGTGGG
1345. $6GQ^*$: ACACACACTGTC CCTTCATCAAGAGTAATCTTGACA GGGTAGTGTGGAGAGGTGGG
1346. $4G$: CTGTTCCCAACA CCTTCATCAAGAGTAATCTTGACA
1347. $7G$: TCACACTTCGTC CCTTCATCAAGAGTAATCTTGACA
1348. $1G$: TCAACTCCGTTC CCTTCATCAAGAGTAATCTTGACA
1349. $2G$: AATGCCACCATT CCTTCATCAAGAGTAATCTTGACA
1350. $A\overline{5}^*$: GATAAATTGTGTCGAAATCCGCGA AATGGGACATGG
1351. $B5^*$: CCTGCTCCATGTTACTTAGCCGGA GGACTGGTAGTG
1352. $\overline{0}H\mathtt{Q}^*$: CCTCTTCTCAGC AGAACCGGATATTCATTACCCAAA GGGTAGTGTGGAGAGGTGGG
1353. $\overline{3}H\mathtt{Q}^*$: TCAATCCTTGCC AGAACCGGATATTCATTACCCAAA GGGTAGTGTGGAGAGGTGGG
1354. $\overline{5}H\mathtt{Q}^*$: CCATGTCCCATT AGAACCGGATATTCATTACCCAAA GGGTAGTGTGGAGAGGTGGG
1355. $\overline{6}H\mathtt{Q}^*$: CAACCAACGTTC AGAACCGGATATTCATTACCCAAA GGGTAGTGTGGAGAGGTGGG
1356. $\overline{4}H$: CACATCCCTGTT AGAACCGGATATTCATTACCCAAA
1357. $\overline{7}H$: TCACACTTCGTC AGAACCGGATATTCATTACCCAAA
1358. $\overline{1}H$: CATCTCCGATCC AGAACCGGATATTCATTACCCAAA
1359. $\overline{2}H$: TCTTTCCAAGCC AGAACCGGATATTCATTACCCAAA
1360. $A\overline{6}^*$: GATAAATTGTGTCGAAATCCGCGA GAACGTTGGTTG
1361. $B6^*$: CCTGCTCCATGTTACTTAGCCGGA GACAGTGTGTGT
1362. $0IQ^*$: CTCATCCTGACC TCAACGTAACAAAGCTGCTCATTC GGGTAGTGTGGAGAGGTGGG
1363. $3IQ^*$: ACAACCCTTGTC TCAACGTAACAAAGCTGCTCATTC GGGTAGTGTGGAGAGGTGGG
1364. $5IQ^*$: CACTACCAGTCC TCAACGTAACAAAGCTGCTCATTC GGGTAGTGTGGAGAGGTGGG
1365. $6IQ^*$: ACACACACTGTC TCAACGTAACAAAGCTGCTCATTC GGGTAGTGTGGAGAGGTGGG
1366. $4I$: CTGTTCCCAACA TCAACGTAACAAAGCTGCTCATTC
1367. $7I$: TCACACTTCGTC TCAACGTAACAAAGCTGCTCATTC
1368. $1I$: TCAACTCCGTTC TCAACGTAACAAAGCTGCTCATTC
1369. $2I$: AATGCCACCATT TCAACGTAACAAAGCTGCTCATTC

1370. $A\overline{7}^*$: GATAAATTGTGTCGAAATCCGCGA GACGAAGTGTGA
1371. $B7^*$: CCTGCTCCATGTTACTTAGCCGGA GGACGAAAGTGA
1372. $\overline{0}J\mathtt{Q}^*$: CCTCTTCTCAGC AGTGAATAAGGCTTGCCCTGACGA GGGTAGTGTGGAGAGGTGGG
1373. $\overline{3}J\mathtt{Q}^*$: TCAATCCTTGCC AGTGAATAAGGCTTGCCCTGACGA GGGTAGTGTGGAGAGGTGGG
1374. $\overline{5}J\mathtt{Q}^*$: CCATGTCCCATT AGTGAATAAGGCTTGCCCTGACGA GGGTAGTGTGGAGAGGTGGG
1375. $\overline{6}J\mathtt{Q}^*$: CAACCAACGTTC AGTGAATAAGGCTTGCCCTGACGA GGGTAGTGTGGAGAGGTGGG
1376. $\overline{4}J$: CACATCCCTGTT AGTGAATAAGGCTTGCCCTGACGA
1377. $\overline{7}J$: TCACACTTCGTC AGTGAATAAGGCTTGCCCTGACGA
1378. $\overline{1}J$: CATCTCCGATCC AGTGAATAAGGCTTGCCCTGACGA
1379. $\overline{2}J$: TCTTTCCAAGCC AGTGAATAAGGCTTGCCCTGACGA
1380. $I$: TCAACGTAACAAAGCTGCTCATTC
1381. $J$: AGTGAATAAGGCTTGCCCTGACGA
1382. $I\overline{0}^*$: TCAACGTAACAAAGCTGCTCATTC GCTGAGAAGAGG
1383. $J0^*$: AGTGAATAAGGCTTGCCCTGACGA GGTCAGGATGAG
1384. $0K\mathtt{Q}^*$: CTCATCCTGACC GAAACACCAGAACGAGTAGTAAAT GGGTAGTGTGGAGAGGTGGG
1385. $3K\mathtt{Q}^*$: ACAACCCTTGTC GAAACACCAGAACGAGTAGTAAAT GGGTAGTGTGGAGAGGTGGG
1386. $5K\mathtt{Q}^*$: CACTACCAGTCC GAAACACCAGAACGAGTAGTAAAT GGGTAGTGTGGAGAGGTGGG
1387. $6K\mathtt{Q}^*$: ACACACACTGTC GAAACACCAGAACGAGTAGTAAAT GGGTAGTGTGGAGAGGTGGG
1388. $4K$: CTGTTCCCAACA GAAACACCAGAACGAGTAGTAAAT
1389. $7K$: TCACTTTCGTCC GAAACACCAGAACGAGTAGTAAAT
1390. $1K$: TCAACTCCGTTC GAAACACCAGAACGAGTAGTAAAT
1391. $2K$: AATGCCACCATT GAAACACCAGAACGAGTAGTAAAT
1392. $I\mathtt{Q}^*$: TCAACGTAACAAAGCTGCTCATTC GGGTAGTGTGGAGAGGTGGG
1393. $J\mathtt{Q}^*$: AGTGAATAAGGCTTGCCCTGACGA GGGTAGTGTGGAGAGGTGGG
1394. $I\overline{1}^*$: TCAACGTAACAAAGCTGCTCATTC GGATCGGAGATG
1395. $J1^*$: AGTGAATAAGGCTTGCCCTGACGA GAACGGAGTTGA
1396. $\overline{0}L\mathtt{Q}^*$: CCTCTTCTCAGC TGGGCTTGAGATGGTTTAATTTCA GGGTAGTGTGGAGAGGTGGG
1397. $\overline{3}L\mathtt{Q}^*$: TCAATCCTTGCC TGGGCTTGAGATGGTTTAATTTCA GGGTAGTGTGGAGAGGTGGG
1398. $\overline{5}L\mathtt{Q}^*$: CCATGTCCCATT TGGGCTTGAGATGGTTTAATTTCA GGGTAGTGTGGAGAGGTGGG
1399. $\overline{6}L\mathtt{Q}^*$: CAACCAACGTTC TGGGCTTGAGATGGTTTAATTTCA GGGTAGTGTGGAGAGGTGGG
1400. $\overline{4}L$: CACATCCCTGTT TGGGCTTGAGATGGTTTAATTTCA
1401. $\overline{7}L$: TCACACTTCGTC TGGGCTTGAGATGGTTTAATTTCA
1402. $\overline{1}L$: CATCTCCGATCC TGGGCTTGAGATGGTTTAATTTCA
1403. $\overline{2}L$: TCTTTCCAAGCC TGGGCTTGAGATGGTTTAATTTCA
1404. $I\overline{2}^*$: TCAACGTAACAAAGCTGCTCATTC GGCTTGGAAAGA
1405. $J2^*$: AGTGAATAAGGCTTGCCCTGACGA AATGGTGGCATT
1406. $0M\mathtt{Q}^*$: CTCATCCTGACC ACTTTAATCATTGTGAATTACCTT GGGTAGTGTGGAGAGGTGGG
1407. $3M\mathtt{Q}^*$: ACAACCCTTGTC ACTTTAATCATTGTGAATTACCTT GGGTAGTGTGGAGAGGTGGG
1408. $5M\mathtt{Q}^*$: CACTACCAGTCC ACTTTAATCATTGTGAATTACCTT GGGTAGTGTGGAGAGGTGGG
1409. $6M\mathtt{Q}^*$: ACACACACTGTC ACTTTAATCATTGTGAATTACCTT GGGTAGTGTGGAGAGGTGGG
1410. $4M$: CTGTTCCCAACA ACTTTAATCATTGTGAATTACCTT
1411. $7M$: TCACTTTCGTCC ACTTTAATCATTGTGAATTACCTT
1412. $1M$: TCAACTCCGTTC ACTTTAATCATTGTGAATTACCTT
1413. $2M$: AATGCCACCATT ACTTTAATCATTGTGAATTACCTT
1414. $I\overline{3}^*$: TCAACGTAACAAAGCTGCTCATTC GGCAAGGATTGA
1415. $J3^*$: AGTGAATAAGGCTTGCCCTGACGA GACAAGGGTTGT
1416. $\overline{0}N\mathtt{Q}^*$: CCTCTTCTCAGC ATGCGATTTTAAGAACTGGCTCAT GGGTAGTGTGGAGAGGTGGG
1417. $\overline{3}N\mathtt{Q}^*$: TCAATCCTTGCC ATGCGATTTTAAGAACTGGCTCAT GGGTAGTGTGGAGAGGTGGG
1418. $\overline{5}N\mathtt{Q}^*$: CCATGTCCCATT ATGCGATTTTAAGAACTGGCTCAT GGGTAGTGTGGAGAGGTGGG
1419. $\overline{6}N\mathtt{Q}^*$: CAACCAACGTTC ATGCGATTTTAAGAACTGGCTCAT GGGTAGTGTGGAGAGGTGGG
1420. $\overline{4}N$: CACATCCCTGTT ATGCGATTTTAAGAACTGGCTCAT
1421. $\overline{7}N$: TCACACTTCGTC ATGCGATTTTAAGAACTGGCTCAT
1422. $\overline{1}N$: CATCTCCGATCC ATGCGATTTTAAGAACTGGCTCAT
1423. $\overline{2}N$: TCTTTCCAAGCC ATGCGATTTTAAGAACTGGCTCAT

1424. $I\overline{4}^*$: TCAACGTAACAAAGCTGCTCATTC AACAGGGATGTG
1425. $J4^*$: AGTGAATAAGGCTTGCCCTGACGA TGTTGGGAACAG
1426. $0O\mathtt{Q}^*$: CTCATCCTGACC TATACCAGTCAGGACGTTGGGAAG GGGTAGTGTGGAGAGGTGGG
1427. $3O\mathtt{Q}^*$: ACAACCCTTGTC TATACCAGTCAGGACGTTGGGAAG GGGTAGTGTGGAGAGGTGGG
1428. $5O\mathtt{Q}^*$: CACTACCAGTCC TATACCAGTCAGGACGTTGGGAAG GGGTAGTGTGGAGAGGTGGG
1429. $6O\mathtt{Q}^*$: ACACACACTGTC TATACCAGTCAGGACGTTGGGAAG GGGTAGTGTGGAGAGGTGGG
1430. $4O$: CTGTTCCCAACA TATACCAGTCAGGACGTTGGGAAG
1431. $7O$: TCACACTTCGTC TATACCAGTCAGGACGTTGGGAAG
1432. $1O$: TCAACTCCGTTC TATACCAGTCAGGACGTTGGGAAG
1433. $2O$: AATGCCACCATT TATACCAGTCAGGACGTTGGGAAG
1434. $I\overline{5}^*$: TCAACGTAACAAAGCTGCTCATTC AATGGGACATGG
1435. $J5^*$: AGTGAATAAGGCTTGCCCTGACGA GGACTGGTAGTG
1436. $\overline{0}P\mathtt{Q}^*$: CCTCTTCTCAGC AAAAATCTACGTTAATAAAACGAA GGGTAGTGTGGAGAGGTGGG
1437. $\overline{3}P\mathtt{Q}^*$: TCAATCCTTGCC AAAAATCTACGTTAATAAAACGAA GGGTAGTGTGGAGAGGTGGG
1438. $\overline{5}P\mathtt{Q}^*$: CCATGTCCCATT AAAAATCTACGTTAATAAAACGAA GGGTAGTGTGGAGAGGTGGG
1439. $\overline{6}P\mathtt{Q}^*$: CAACCAACGTTC AAAAATCTACGTTAATAAAACGAA GGGTAGTGTGGAGAGGTGGG
1440. $\overline{4}P$: CACATCCCTGTT AAAAATCTACGTTAATAAAACGAA
1441. $\overline{7}P$: TCACACTTCGTC AAAAATCTACGTTAATAAAACGAA
1442. $\overline{1}P$: CATCTCCGATCC AAAAATCTACGTTAATAAAACGAA
1443. $\overline{2}P$: TCTTTCCAAGCC AAAAATCTACGTTAATAAAACGAA
1444. $I\overline{6}^*$: TCAACGTAACAAAGCTGCTCATTC GAACGTTGGTTG
1445. $J6^*$: AGTGAATAAGGCTTGCCCTGACGA GACAGTGTGTGT
1446. $0Q\mathtt{Q}^*$: CTCATCCTGACC CTAACGGAACAACATTATTACAGG GGGTAGTGTGGAGAGGTGGG
1447. $3Q\mathtt{Q}^*$: ACAACCCTTGTC CTAACGGAACAACATTATTACAGG GGGTAGTGTGGAGAGGTGGG
1448. $5Q\mathtt{Q}^*$: CACTACCAGTCC CTAACGGAACAACATTATTACAGG GGGTAGTGTGGAGAGGTGGG
1449. $6Q\mathtt{Q}^*$: ACACACACTGTC CTAACGGAACAACATTATTACAGG GGGTAGTGTGGAGAGGTGGG
1450. $4Q$: CTGTTCCCAACA CTAACGGAACAACATTATTACAGG
1451. $7Q$: TCACACTTCGTC CTAACGGAACAACATTATTACAGG
1452. $1Q$: TCAACTCCGTTC CTAACGGAACAACATTATTACAGG
1453. $2Q$: AATGCCACCATT CTAACGGAACAACATTATTACAGG
1454. $I\overline{7}^*$: TCAACGTAACAAAGCTGCTCATTC GACGAAGTGTGA
1455. $J7^*$: AGTGAATAAGGCTTGCCCTGACGA GGACGAAAGTGA
1456. $\overline{0}R\mathtt{Q}^*$: CCTCTTCTCAGC TAGAAAGATTCATCAGTTGAGATT GGGTAGTGTGGAGAGGTGGG
1457. $\overline{3}R\mathtt{Q}^*$: TCAATCCTTGCC TAGAAAGATTCATCAGTTGAGATT GGGTAGTGTGGAGAGGTGGG
1458. $\overline{5}R\mathtt{Q}^*$: CCATGTCCCATT TAGAAAGATTCATCAGTTGAGATT GGGTAGTGTGGAGAGGTGGG
1459. $\overline{6}R\mathtt{Q}^*$: CAACCAACGTTC TAGAAAGATTCATCAGTTGAGATT GGGTAGTGTGGAGAGGTGGG
1460. $\overline{4}R$: CACATCCCTGTT TAGAAAGATTCATCAGTTGAGATT
1461. $\overline{7}R$: TCACACTTCGTC TAGAAAGATTCATCAGTTGAGATT
1462. $\overline{1}R$: CATCTCCGATCC TAGAAAGATTCATCAGTTGAGATT
1463. $\overline{2}R$: TCTTTCCAAGCC TAGAAAGATTCATCAGTTGAGATT
1464. $Q$: CTAACGGAACAACATTATTACAGG
1465. $R$: TAGAAAGATTCATCAGTTGAGATT
1466. $Q\overline{0}^*$: CTAACGGAACAACATTATTACAGG GCTGAGAAGAGG
1467. $R0^*$: TAGAAAGATTCATCAGTTGAGATT GGTCAGGATGAG
1468. $0S\mathtt{Q}^*$: CTCATCCTGACC TAGGAATACCACATTCAACTAATG GGGTAGTGTGGAGAGGTGGG
1469. $3S\mathtt{Q}^*$: ACAACCCTTGTC TAGGAATACCACATTCAACTAATG GGGTAGTGTGGAGAGGTGGG
1470. $5S\mathtt{Q}^*$: CACTACCAGTCC TAGGAATACCACATTCAACTAATG GGGTAGTGTGGAGAGGTGGG
1471. $6S\mathtt{Q}^*$: ACACACACTGTC TAGGAATACCACATTCAACTAATG GGGTAGTGTGGAGAGGTGGG
1472. $4S$: CTGTTCCCAACA TAGGAATACCACATTCAACTAATG
1473. $7S$: TCACTTTCGTCC TAGGAATACCACATTCAACTAATG
1474. $1S$: TCAACTCCGTTC TAGGAATACCACATTCAACTAATG
1475. $2S$: AATGCCACCATT TAGGAATACCACATTCAACTAATG
1476. $Q\mathtt{Q}^*$: CTAACGGAACAACATTATTACAGG GGGTAGTGTGGAGAGGTGGG
1477. $R\mathtt{Q}^*$: TAGAAAGATTCATCAGTTGAGATT GGGTAGTGTGGAGAGGTGGG

1478. $Q\overline{1}^*$: CTAACGGAACAACATTATTACAGG GGATCGGAGATG
1479. $R1^*$: TAGAAAGATTCATCAGTTGAGATT GAACGGAGTTGA
1480. $\overline{0}T\mathtt{Q}^*$: CCTCTTCTCAGC CAGATACATAACGCCAAAAGGAAT GGGTAGTGTGGAGAGGTGGG
1481. $\overline{3}T\mathtt{Q}^*$: TCAATCCTTGCC CAGATACATAACGCCAAAAGGAAT GGGTAGTGTGGAGAGGTGGG
1482. $\overline{5}T\mathtt{Q}^*$: CCATGTCCCATT CAGATACATAACGCCAAAAGGAAT GGGTAGTGTGGAGAGGTGGG
1483. $\overline{6}T\mathtt{Q}^*$: CAACCAACGTTC CAGATACATAACGCCAAAAGGAAT GGGTAGTGTGGAGAGGTGGG
1484. $\overline{4}T$: CACATCCCTGTT CAGATACATAACGCCAAAAGGAAT
1485. $\overline{7}T$: TCACACTTCGTC CAGATACATAACGCCAAAAGGAAT
1486. $\overline{1}T$: CATCTCCGATCC CAGATACATAACGCCAAAAGGAAT
1487. $\overline{2}T$: TCTTTCCAAGCC CAGATACATAACGCCAAAAGGAAT
1488. $Q\overline{2}^*$: CTAACGGAACAACATTATTACAGG GGCTTGGAAAGA
1489. $R2^*$: TAGAAAGATTCATCAGTTGAGATT AATGGTGGCATT
1490. $0U\mathtt{Q}^*$: CTCATCCTGACC TACGAGGCATAGTAAGAGCAACAC GGGTAGTGTGGAGAGGTGGG
1491. $3U\mathtt{Q}^*$: ACAACCCTTGTC TACGAGGCATAGTAAGAGCAACAC GGGTAGTGTGGAGAGGTGGG
1492. $5U\mathtt{Q}^*$: CACTACCAGTCC TACGAGGCATAGTAAGAGCAACAC GGGTAGTGTGGAGAGGTGGG
1493. $6U\mathtt{Q}^*$: ACACACACTGTC TACGAGGCATAGTAAGAGCAACAC GGGTAGTGTGGAGAGGTGGG
1494. $4U$: CTGTTCCCAACA TACGAGGCATAGTAAGAGCAACAC
1495. $7U$: TCACACTTCGTC TACGAGGCATAGTAAGAGCAACAC
1496. $1U$: TCAACTCCGTTC TACGAGGCATAGTAAGAGCAACAC
1497. $2U$: AATGCCACCATT TACGAGGCATAGTAAGAGCAACAC
1498. $Q\overline{3}^*$: CTAACGGAACAACATTATTACAGG GGCAAGGATTGA
1499. $R3^*$: TAGAAAGATTCATCAGTTGAGATT GACAAGGGTTGT
1500. $\overline{0}V\mathtt{Q}^*$: CCTCTTCTCAGC TATCATAACCCTCGTTTACCAGAC GGGTAGTGTGGAGAGGTGGG
1501. $\overline{3}V\mathtt{Q}^*$: TCAATCCTTGCC TATCATAACCCTCGTTTACCAGAC GGGTAGTGTGGAGAGGTGGG
1502. $\overline{5}V\mathtt{Q}^*$: CCATGTCCCATT TATCATAACCCTCGTTTACCAGAC GGGTAGTGTGGAGAGGTGGG
1503. $\overline{6}V\mathtt{Q}^*$: CAACCAACGTTC TATCATAACCCTCGTTTACCAGAC GGGTAGTGTGGAGAGGTGGG
1504. $\overline{4}V$: CACATCCCTGTT TATCATAACCCTCGTTTACCAGAC
1505. $\overline{7}V$: TCACACTTCGTC TATCATAACCCTCGTTTACCAGAC
1506. $\overline{1}V$: CATCTCCGATCC TATCATAACCCTCGTTTACCAGAC
1507. $\overline{2}V$: TCTTTCCAAGCC TATCATAACCCTCGTTTACCAGAC
1508. $Q\overline{4}^*$: CTAACGGAACAACATTATTACAGG AACAGGGATGTG
1509. $R4^*$: TAGAAAGATTCATCAGTTGAGATT TGTTGGGAACAG
1510. $\overline{0}W\mathtt{Q}^*$: CTCATCCTGACC GACGATAAAAACCAAAATAGCGAG GGGTAGTGTGGAGAGGTGGG
1511. $\overline{3}W\mathtt{Q}^*$: ACAACCCTTGTC GACGATAAAAACCAAAATAGCGAG GGGTAGTGTGGAGAGGTGGG
1512. $\overline{5}W\mathtt{Q}^*$: CACTACCAGTCC GACGATAAAAACCAAAATAGCGAG GGGTAGTGTGGAGAGGTGGG
1513. $\overline{6}W\mathtt{Q}^*$: ACACACACTGTC GACGATAAAAACCAAAATAGCGAG GGGTAGTGTGGAGAGGTGGG
1514. $\overline{4}W$: CTGTTCCCAACA GACGATAAAAACCAAAATAGCGAG
1515. $\overline{7}W$: TCACACTTCGTC GACGATAAAAACCAAAATAGCGAG
1516. $\overline{1}W$: TCAACTCCGTTC GACGATAAAAACCAAAATAGCGAG
1517. $\overline{2}W$: AATGCCACCATT GACGATAAAAACCAAAATAGCGAG
1518. $Q\overline{5}^*$: CTAACGGAACAACATTATTACAGG AATGGGACATGG
1519. $R5^*$: TAGAAAGATTCATCAGTTGAGATT GGACTGGTAGTG
1520. $\overline{0}X\mathtt{Q}^*$: CCTCTTCTCAGC AGGCTTTTGCAAAAGAAGTTTTGC GGGTAGTGTGGAGAGGTGGG
1521. $\overline{3}X\mathtt{Q}^*$: TCAATCCTTGCC AGGCTTTTGCAAAAGAAGTTTTGC GGGTAGTGTGGAGAGGTGGG
1522. $\overline{5}X\mathtt{Q}^*$: CCATGTCCCATT AGGCTTTTGCAAAAGAAGTTTTGC GGGTAGTGTGGAGAGGTGGG
1523. $\overline{6}X\mathtt{Q}^*$: CAACCAACGTTC AGGCTTTTGCAAAAGAAGTTTTGC GGGTAGTGTGGAGAGGTGGG
1524. $\overline{4}X$: CACATCCCTGTT AGGCTTTTGCAAAAGAAGTTTTGC
1525. $\overline{7}X$: TCACACTTCGTC AGGCTTTTGCAAAAGAAGTTTTGC
1526. $\overline{1}X$: CATCTCCGATCC AGGCTTTTGCAAAAGAAGTTTTGC
1527. $\overline{2}X$: TCTTTCCAAGCC AGGCTTTTGCAAAAGAAGTTTTGC
1528. $Q\overline{6}^*$: CTAACGGAACAACATTATTACAGG GAACGTTGGTTG
1529. $R6^*$: TAGAAAGATTCATCAGTTGAGATT GACAGTGTGTGT
1530. $0Y\mathtt{Q}^*$: CTCATCCTGACC CAGAGGGGGTAATAGTAAAATGTT GGGTAGTGTGGAGAGGTGGG
1531. $3Y\mathtt{Q}^*$: ACAACCCTTGTC CAGAGGGGGTAATAGTAAAATGTT GGGTAGTGTGGAGAGGTGGG

1532. $5Y\mathtt{Q}^*$: CACTACCAGTCC CAGAGGGGGTAATAGTAAAATGTT GGGTAGTGTGGAGAGGTGGG
1533. $\overline{6}Y\mathtt{Q}^*$: ACACACACTGTC CAGAGGGGGTAATAGTAAAATGTT GGGTAGTGTGGAGAGGTGGG
1534. $4Y$: CTGTTCCCAACA CAGAGGGGGTAATAGTAAAATGTT
1535. $7Y$: TCACACTTCGTC CAGAGGGGGTAATAGTAAAATGTT
1536. $1Y$: TCAACTCCGTTC CAGAGGGGGTAATAGTAAAATGTT
1537. $2Y$: AATGCCACCATT CAGAGGGGGTAATAGTAAAATGTT
1538. $Q\overline{7}^*$: CTAACGGAACAACATTATTACAGG GACGAAGTGTGA
1539. $R7^*$: TAGAAAGATTCATCAGTTGAGATT GGACGAAAGTGA
